# Programmable Self-Assembly of RNA Nanostructures with >100 Unique Components

**DOI:** 10.64898/2026.09.02.748990

**Authors:** Liangxiao Chen, Zhishang Li, Jun Yan, Swarup Dey, Cong Li, Alexandra Petrova, Abhay Prasad, Deeksha Satyabola, Kira DeVore, Xinyi Tu, Yanzhe Qu, Shiyumou Wang, Gengshi Wu, Po-Lin Chiu, Di Wen, Joseph Che-Yen Wang, Hao Yan, Peng Yin, Di Liu

## Abstract

Sophisticated biomolecular functions often arise from large, precisely organized architectures, motivating efforts to construct increasingly complex structures through programmed nucleic acid self-assembly. Although RNA offers a richer repertoire of structural motifs and biological functions than DNA, RNA nanostructures constructed to date have remained substantially less complex and less scalable than their DNA counterparts, most notably DNA origami and DNA bricks, which rely on large libraries of synthetic single-stranded (ss) DNA strands. Directly adapting these strategies to RNA, however, faces two major barriers: (i) the high cost of producing large libraries of distinct synthetic ssRNA strands, and (ii) the limited stability of ssRNA under conditions commonly used for assembling multicomponent nucleic acid nanostructures. Here we present the double-stranded RNA (dsRNA) bricks approach, which addresses both barriers by adopting principles inspired by natural RNA systems: (i) many distinct RNA components (“bricks”) are encoded within a single precursor transcript and released by enzymatic processing, (ii) each brick’s predominantly double-stranded nature enhances stability. The resulting dsRNA bricks self-assemble through programmable branched kissing-loop interactions. Using this approach, we constructed complex two- and three-dimensional RNA nanostructures with more than 100 distinct components, representing, to our knowledge, the largest and most compositionally complex RNA nanoarchitectures reported to date. Overall, the dsRNA bricks approach establishes a scalable and robust framework for constructing complex RNA nanostructures with compositional and architectural sophistication approaching that of DNA-based systems, thereby opening new opportunities for programmable RNA materials and RNA-based devices.

## Introduction

In nature, self-assembly of biomolecules such as proteins and nucleic acids produces large and complex architectures that fulfill sophisticated functions within biological systems^1,2^. Inspired by these systems, engineered biomolecular self-assembly has become a powerful route for creating functional biomaterials with applications in biotechnology and biomedicine^3–6^. Among biomolecular building blocks, nucleic acids—DNA and RNA—are uniquely attractive because their sequence-dictated base pairing and well-defined helical geometry provide a reliable molecular grammar for programmable nanoscale construction. Over the past four decades, these features have driven rapid progress in nucleic acid nanotechnology, enabling the construction of structures with increasing size, complexity, and functional integration^7–10^. A notable advance in the field is DNA origami^11,12^, in which a long scaffold strand is folded by hundreds of short synthetic staple strands into prescribed geometries. In parallel, DNA bricks, also known as single-stranded tiles (SSTs)^13–15^, established a scaffold-free and highly modular strategy for self-assembling hundreds, and in some cases tens of thousands, of distinct ssDNA strands, each serving as an individual brick or tile, much like a molecular LEGO-like construction system.

Compared with DNA, RNA offers a broader design palette: RNA supports diverse secondary and tertiary structural motifs^16^ and can encode biological functions such as ligand binding, catalysis, and protein recognition, making it an especially compelling material for building biologically functional nanoscale systems^17,18^. In addition, several design principles and structural motifs originally developed in DNA nanotechnology have been adapted for RNA nanostructure construction, including three-way junction^19^, T-junction^20^ and several double-crossover motifs^21,22^. Despite this promise, RNA nanostructures^9,18,23,24^ remain less complex and less scalable compared to their DNA counterparts with respect to both total component number and total nucleotide count (Supplementary Fig. 1). Directly translating mature DNA nanotechnology strategies, such as DNA origami and DNA bricks, into RNA is not straightforward, because both strategies critically rely on large libraries of chemically synthesized ssDNA components. In an RNA context, this reliance creates two major obstacles: first, large libraries of distinct RNA strands are substantially more costly to produce by chemical synthesis^25^; second, ssRNA is more vulnerable to enzymatic degradation and chemical cleavage^26^, particularly during prolonged exposure to heating or divalent cations^27^ under conditions commonly used for multicomponent self-assembly^28^.

In contrast to phosphoramidite-based solid-phase chemical synthesis, RNA, particularly for long transcripts, can be prepared more readily by enzymatic in vitro transcription (IVT). Accordingly, efforts to overcome the scalability limitations of RNA nanotechnology have led to the development of ingenious methods based on one or a small number of IVT transcripts, which undergo hierarchical folding and assembly: predefined secondary structures form first, followed by higher-order organization mediated by natural or artificial interaction motifs, including kissing loops (KLs)^29–31^, branched kissing loops^32–35^ (bKLs), and parallel crossover cohesions^36,37^. While powerful, architectures built from these methods typically face at least one of two important limitations: first, repeated use of identical components sacrifices full addressability because individual sites cannot be independently specified or functionalized; and second, designs based on a single long strand impose substantial strand-routing constraints and topological or kinetic barriers, which become especially severe for space-filling three-dimensional (3D) RNA architectures. Therefore, a desirable framework for highly complex RNA nanostructure construction should ideally integrate three key features: (i) compatibility with long RNA transcripts generated by IVT, (ii) stable RNA building blocks with minimal single-stranded regions, and (iii) assembly from many distinct components without the routing constraints imposed by a single long strand.

Here we present such a framework—the double-stranded RNA (dsRNA) bricks approach—inspired by two general principles in natural RNA biogenesis: (i) the enzymatic maturation of long precursor transcripts into multiple functional RNA components and (ii) the stabilization by extensive base pairing. In this dsRNA bricks approach, a single IVT-generated precursor RNA encodes many distinct dsRNA bricks, separated by linker regions that hybridize with a guide DNA to enable RNase H-mediated cleavage and release of the individual bricks. These predominantly double-stranded bricks contain only minimal single-stranded regions, confined to loops and bulges that mediate programmable bKL interactions, allowing the bricks to self-assemble into prescribed geometries without the strand-routing constraints that would otherwise be imposed by a single long RNA strand. We further developed an expanded 9-bp bKL design, building on our previous experience in designing 6-bp bKL^32^, to expand the orthogonal sequence space for more complex assemblies. Combining the dsRNA bricks approach with the expanded 9-bp bKL design, we have constructed finite-sized 2D and 3D RNA nanostructures comprising more than 100 distinct components, representing, to our knowledge, the largest and most compositionally complex discrete RNA nanostructures reported so far. Together, these results establish the dsRNA bricks strategy as a scalable and robust platform for constructing complex RNA nanostructures with a degree of compositional and architectural sophistication approaching that of DNA-based systems.

### Concept and design of dsRNA bricks

A common principle in RNA biology is that multiple functional RNA components can be generated from a single polycistronic precursor transcript through enzymatic processing. Ribosomal RNA (rRNA) maturation^38,39^ provides a canonical example of this strategy. In bacteria such as *E. coli*, rRNAs are first synthesized as a single precursor transcript that is subsequently processed to release mature rRNAs, which then assemble with ribosomal proteins into functional ribosomes^40^ (Fig. 1a). This natural mechanism exemplifies an efficient strategy for coordinated production of multiple RNA components from a single precursor transcript, while also helping maintain defined stoichiometry for downstream assembly^41^. Inspired by this principle, we developed the dsRNA bricks approach for constructing high-complexity RNA nanostructures (Fig. 1b). In this approach, a DNA template encodes a long precursor RNA containing multiple RNA building blocks, separated by a universal linker sequence. These RNA building blocks are termed “dsRNA bricks” because they fold into predominantly double-stranded structures, a feature expected to enhance their stability. Following IVT, the long precursor RNA is hybridized at the linker regions with a complementary guide DNA, creating RNA-DNA hybrids that direct RNase H cleavage to release the individual dsRNA bricks for subsequent self-assembly into higher-order structures. As shown in the inset of Fig. 1b, a typical dsRNA brick contains three double-stranded helical domains (two beam helices and one strut helix) and four single-stranded regions (two bulges and two hairpin loops). These bulges and loops mediate the inter-brick bKL interactions^32^ (Fig. 1c), in which kissing helices form through programmable Watson–Crick base pairing. In the assembled structure, the resulting extended, coaxially stacked pseudo-continuous helices are termed ‘rails’ (four rails run along the x axis in the assembled structure shown in Fig. 1b). By modulating beam and strut lengths together with the interaction patterns of the bKLs, the geometry of the resulting dsRNA-brick assemblies can be precisely programmed.

**Fig. 1.**
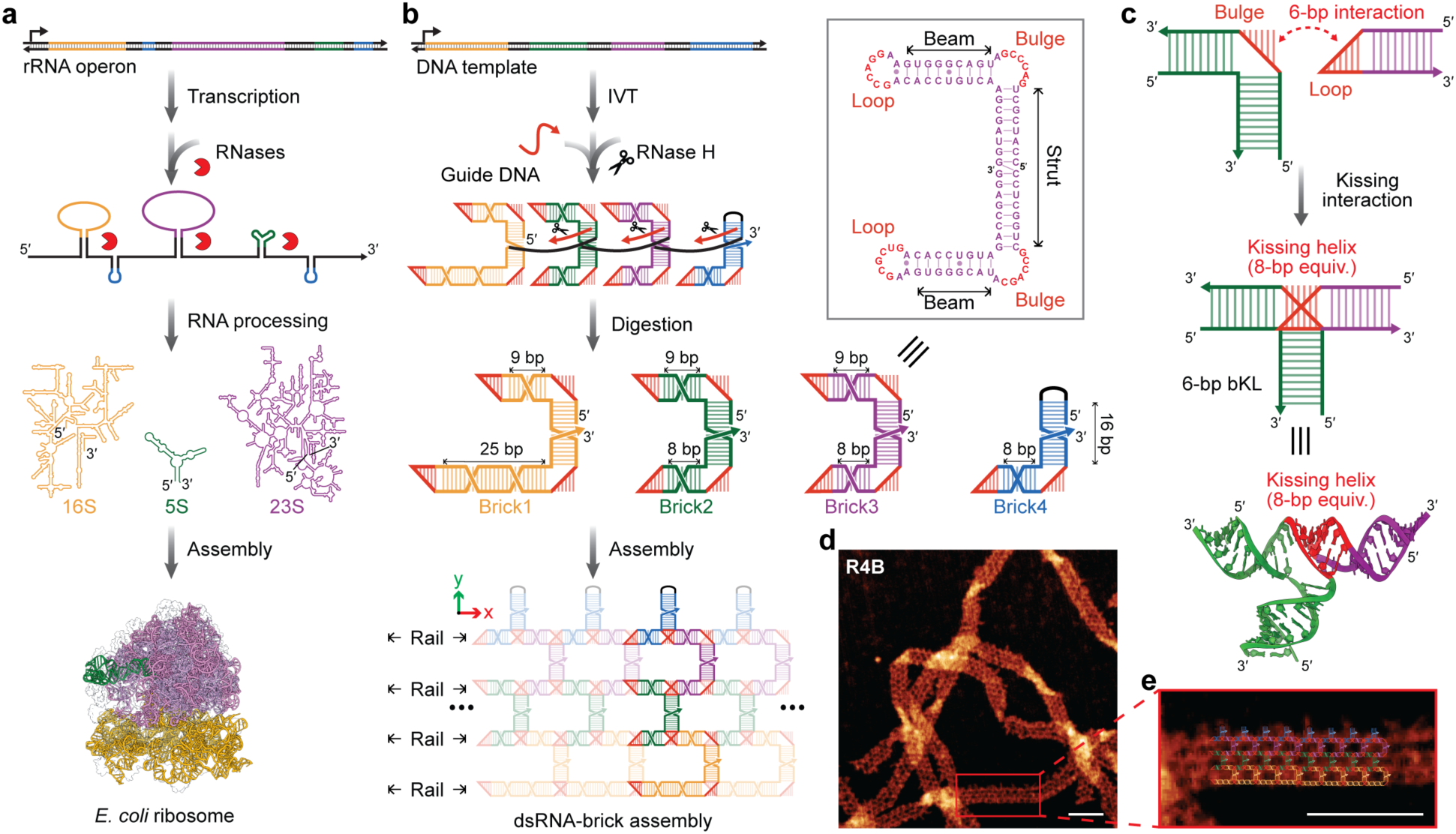
Nature-inspired design and validation of the dsRNA bricks approach. a, Natural processing pathway of bacterial rRNAs. In a typical rRNA operon, such as that of *E. coli*, 16S, 23S, and 5S rRNAs, often together with tRNAs, are transcribed as a single precursor RNA containing spacer sequences. The precursor undergoes successive enzymatic processing steps to generate mature rRNAs, which assemble with ribosomal proteins into a functional ribosome. Different colors denote distinct rRNA components (orange, green, and purple) and tRNA sequences (blue). **b,** Schematic of the dsRNA bricks workflow using the example of a four-brick ribbon, **R4B**. Sequences encoding multiple RNA bricks are incorporated within a single DNA template and separated by copies of a universal linker sequence. Following IVT, the precursor RNA is hybridized with a guide DNA complementary to the linker regions. RNase H-mediated cleavage of the resulting RNA-DNA hybrids releases individual dsRNA bricks, which subsequently self-assemble through programmed inter-brick interactions. In the assembled ribbon, coaxially stacked pseudo-continuous helices running along the x axis are termed rails (four in **R4B**); ellipses denote periodic propagation along the x axis. The inset shows the sequence and secondary structure of a representative dsRNA brick. Each canonical brick contains two beam helices and one strut helix, together with two loops and two bulges that serve as programmable interaction sites. **c**, Formation and structure of an established 6-bp bKL interaction used to connect neighboring dsRNA bricks. Complementary loop and bulge sequences (top) interact to form a kissing helix (red) of a bKL interaction (middle and bottom, 2D and 3D models, respectively). The 6-bp bKL contributes to a helical twist equivalent to that of an 8-bp helix (8 bp equiv.) as estimated previously^32^. **d**, **e**, Zoomed-out (**d**) and zoomed-in (**e**) AFM images of **R4B** ribbons. In **e**, the **R4B** schematic, colored as in **b**, is superimposed on part of the imaged ribbon, showing the expected periodic lattice. Scale bar, 50 nm.

### Validation of the dsRNA bricks method

To validate the dsRNA bricks method, we first designed a four-brick ribbon (**R4B**) structure using the original 6-bp bKL motif^32^. **R4B** comprises four distinct dsRNA bricks (Fig. 1b and Supplementary Fig. 2). The central two (**R4B-brick2** and **R4B-brick3**) are generic C-shaped bricks (C-bricks), each containing a 16-bp strut helix, corresponding to approximately 1.5 helical turns. Their two beam helices were designed to be 8 and 9 bp respectively so that each C-brick contributes an effective twist equivalent to a 16- or 17-bp helix (∼1.5 helical turns) along the rails, assuming that each 6-bp bKL contributes a twist equivalent to an 8-bp helix (8-bp equiv., Fig. 1c), as estimated previously^32^. This design is expected to produce an approximately 180° rotation between adjacent C-bricks and to minimize the net twist (3 helical turns) over the repeating unit. **R4B-brick1** and **R4B-brick4** are boundary bricks derived from the generic C-bricks: **R4B-brick1** has one beam extended to 25 bp along the bottom rail, while the other beam retains the standard length, and this brick serves as the potential nucleation site that promotes self-assembly and improves assembly kinetics^42–44^; **R4B-brick4** is a half-C-brick, containing only one beam and serves as the terminating brick, with the strut capped by a tetraloop.

The **R4B** precursor RNA contains linker regions separating the four brick-encoding sequences. To optimize the RNase H-mediated cleavage of the precursor, we evaluated a series of guide DNA designs and found that a 14-nt all-DNA guide produced the highest cleavage efficiency among the tested designs (Supplementary Fig. 3). The released dsRNA bricks were then annealed in 1× TAE buffer containing 1 mM free Mg^2+^ by cooling from 70 °C to 4 °C over 3 h. Atomic force microscopy (AFM) imaging revealed micrometer-long ribbons with the expected grid-like pattern, consistent with **R4B** assembly through the dsRNA bricks method (Fig. 1d, e and Supplementary Fig. 4).

### Geometric control of periodic dsRNA-brick assemblies

By adjusting the number of bricks and programming the inter-brick bKL sequences, we designed different dsRNA-brick assemblies in which the bricks within each repeating unit are arranged along either the y axis or the x axis (Fig. 2a). To increase the number of bricks along the y axis within each repeating unit, we added two or four C-bricks to the **R4B** design, generating a six-brick ribbon (**R6B**, Fig. 2b and Supplementary Fig. 5 and 6) and an eight-brick ribbon (**R8B**, Fig. 2c and Supplementary Fig. 7 and 8), respectively. To extend the bricks along the x axis within a repeating unit, we arranged three consecutive four-brick modules with distinct bKL sequences to produce **R4×3**, a twelve-brick ribbon whose repeating unit comprises four bricks along the y axis and three repeats along the x axis (Fig. 2d and Supplementary Fig. 9).

**Fig. 2.**
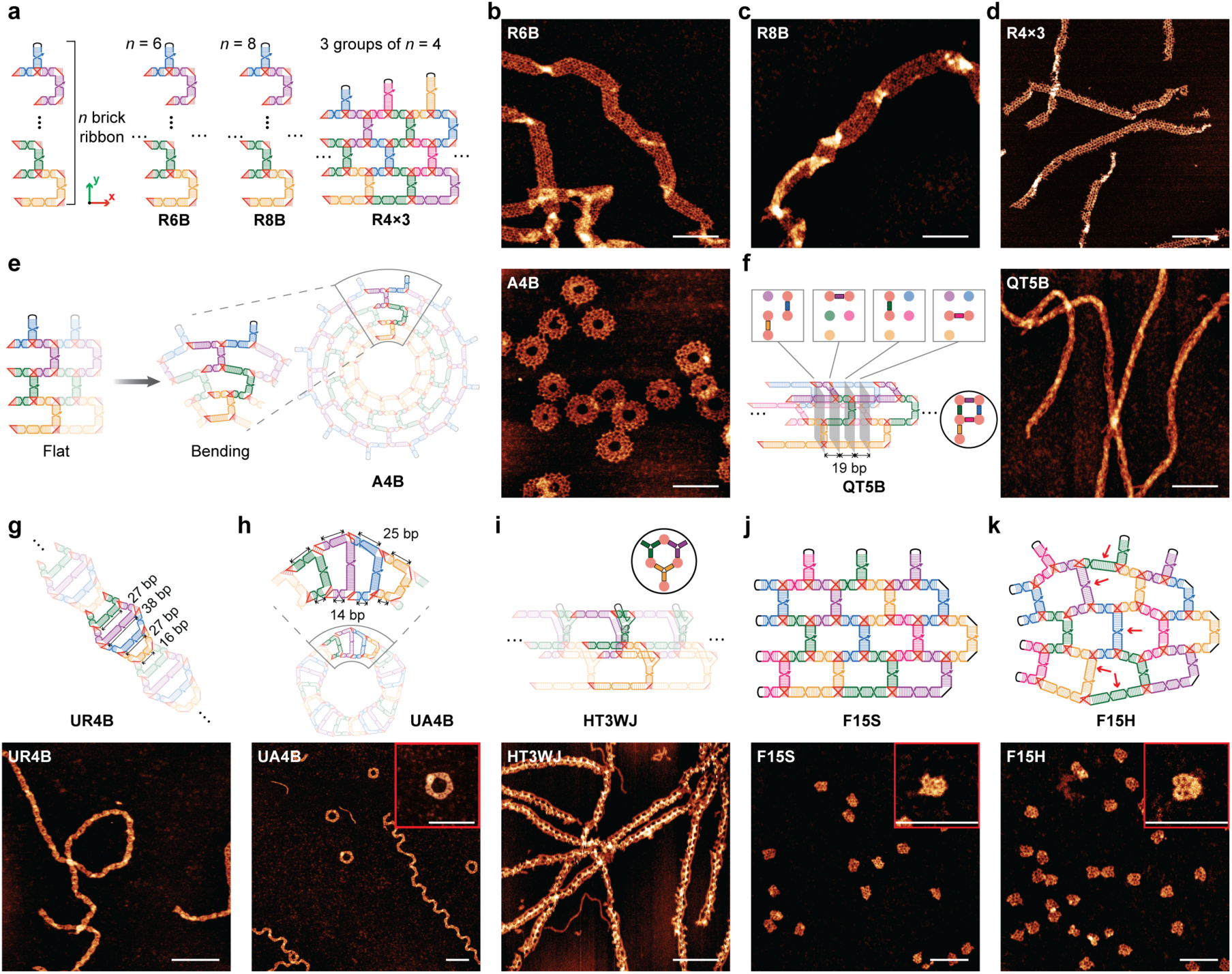
Programmable RNA architectures assembled from dsRNA bricks. a, Designs of periodic ribbon assemblies derived from the generic *n*-brick ribbon architecture. **R6B** and **R8B** comprise six and eight distinct dsRNA bricks along the y axis, respectively. **R4×3** extends the **R4B** architecture along the x axis by combining three consecutive four-brick modules with distinct bKL sequences, generating a twelve-brick repeating unit. **b–d**, AFM images of **R6B** (**b**), **R8B** (**c**) and **R4×3** (**d**) ribbon assemblies. **e**, Design principle and AFM image for in-plane curvature of **A4B**. Unequal rail lengths produce a trapezoidal repeating unit that bends to form the annular four-brick assembly, **A4B**. **f**, Design and AFM image for the five-brick square-tube assembly, **QT5B**. Programmed beam lengths produce defined inter-brick dihedral angles and a square tube cross-section. Boxed schematics show the brick arrangement at the four marked grey planes; the circled inset shows the resulting square cross-section of the assembly. **g**, Design and AFM image of the undulated ribbon **UR4B,** containing varied strut lengths. **h**, Design and AFM images of the undulated annulus **UA4B**. Extending one beam of each **UR4B** brick introduces in-plane curvature, producing annular and extended ribbon-like assembly products. **i**, Design and AFM image of the 3WJ-based hexagonal tube **HT3WJ**, assembled from claw-like dsRNA bricks containing three-way junctions. The circled inset shows the resulting hexagonal cross-section of the assembly. **j**, Design and AFM images of fully addressable finite-sized shield-shaped assemblies, **F15S**. Non-interacting loops and bulges (black) terminate growth at the boundaries. **k**, Design and AFM images of fully addressable finite-sized heart-shaped assemblies, **F15H**. **F15H** is derived from **F15S** by extending selected beams and struts, indicated by red arrows, to broaden and round the lateral edges. Scale bars, 100 nm.

The geometries of dsRNA-brick assemblies can be further tuned by varying beam and strut helix lengths within C-bricks or by modifying the brick architecture beyond the C-brick shape. First, differences in repeating-unit lengths along individual rails can introduce in-plane bending during self-assembly^32,45^. In the four-brick annulus (**A4B**) design, the repeating unit is trapezoidal, spanning 22, 33, 44, and 55 bp, respectively, along the four rails, causing the rails to curve into concentric rings (Fig. 2e). Geometric analysis predicted that a flat **A4B** annulus would contain 12 or 13 repeating units (Supplementary Fig. 10), broadly consistent with AFM analysis showing that most closed annuli contained 12–15 repeating units (Fig. 2e and Supplementary Fig. 11 and 12). Second, the effective along-rail spacing between adjacent bricks can be tuned to introduce torsion about the x axis^46,47^ (Supplementary Fig. 13). For example, in the design of the five-brick square tube (**QT5B**), the effective along-rail spacing is 19 bp, which is 3 bp shorter than two full helical turns and is expected to produce an approximately 270° helical rotation between neighboring bricks (Supplementary Fig. 14), yielding a tube with a square cross-section (Fig. 2f and Supplementary Fig. 15). Similarly, the seven-brick hexagonal tube (**HT7B**) design employs an 18-bp effective along-rail spacing and is expected to produce an approximately 240° helical rotation between neighboring bricks (Supplementary Fig. 16 and 17).

Third, we further explored the geometric design space of C-brick assemblies by varying strut lengths. In the four-brick undulated ribbon (**UR4B**) design, adjacent dsRNA bricks are separated by two helical turns along the ribbon, whereas the strut lengths of the four dsRNA bricks were set to 16, 27, 38, and 27 bp, respectively, producing an undulated ribbon with periodically varying width (Fig. 2g and Supplementary Fig. 18 and 19). Additionally, we increased one beam of each **UR4B** brick to 25 bp, with the elongated beams placed on the same rail of the assembly to introduce in-plane bending, yielding the undulated annulus (**UA4B**) design (Supplementary Fig. 20). Notably, the **UA4B** design produced two morphologically distinct classes of structures: the expected closed annuli and unexpected long, undulated ribbons (Fig. 2h and Supplementary Fig. 21). The latter structures may arise from geometric frustration in the design, which disfavors annulus closure and instead promotes continued growth along the ribbon axis.

Finally, in addition to C-bricks, we also explored claw-like dsRNA bricks by connecting three half-C-bricks through a three-way junction (3WJ)^32^. In the 3WJ-based three-brick hexagonal tube (**HT3WJ**) design, the 3WJs lie in the y–z plane (the beams extending along the x axis). The 18-bp effective spacing along the x axis is expected to impose an approximately 120° dihedral angle between adjacent dsRNA bricks, and together with the ∼120° angles formed by the branches of each 3WJ, this arrangement generates a hexagonal cross-section, in which alternating vertices are defined by the 3WJs and the inter-brick dihedral angles (Fig. 2i and Supplementary Fig. 22). AFM imaging revealed **HT3WJ** tubes with features consistent with a 3D tube morphology (Supplementary Fig. 23). These results show that the dsRNA bricks method can be readily extended to multivalent building blocks with intrinsic 3D geometries.

### Fully addressable finite-sized dsRNA-brick assemblies

In addition to periodic assemblies, finite-sized dsRNA-brick assemblies, in which each brick is uniquely addressable, can be constructed by assigning distinct interaction patterns to individual bricks and terminating growth at boundaries. We designed two finite-sized assemblies with a shield-shaped structure (**F15S**) and a heart-shaped structure (**F15H**), respectively, each containing 15 bricks (Fig. 2j, k). In **F15S**, the design principle for the inner bricks is similar to that of the above 4-layered ribbon designs, such as **R4B** and **R4×3**. To prevent periodic extension of the assembly along the x axis, terminating bricks are incorporated at both vertical edges by replacing active bKL loops and bulges with non-interacting tetraloops (UUCG or GUAA) and non-interacting bulges (AAAUAAA or AAUAAUA). **F15H** shares the same overall frame architecture as **F15S**, but the beams and struts of selected bricks (indicated by red arrows in Fig. 2k) have been elongated by one helical turn so that the two lateral edges broaden and curve outward. The successful construction of **F15S** and **F15H** (Fig. 2j, k and Supplementary Fig. 24-26) demonstrates that the dsRNA bricks method supports finite-sized RNA nanostructures with programmable global geometry and individually addressable component positions.

### Expanding the orthogonal sequence space of bKLs

The structural complexity of the dsRNA-brick assemblies described thus far (up to 15 distinct bricks with 23 pairs of bKL interactions in **F15H**) still falls short of that routinely achieved by DNA origami and DNA bricks. Indeed, when we attempted to construct a periodic 15-brick ribbon (**R15B**) with 29 pairs of bKLs, only a small fraction of the products showed the expected assembly morphology (Supplementary Fig. 27). Although multiple factors may contribute to this limitation, crosstalk among bKL interactions is likely to impair correct assembly. Accordingly, the number of mutually orthogonal bKL pairs imposes a practical upper limit on the number of distinct dsRNA bricks that can be incorporated into an assembly, and thus on its overall structural complexity. However, the orthogonal sequence space of the widely used original 6-bp bKL design (on which all of the above designs are based) is limited, especially under our requirement for relatively strong and comparable interaction strengths with a fixed GC-rich composition of five G-C pairs and one A-U pair.

Because longer kissing helices can expand the potential orthogonal sequence space, we explored whether bKL motifs with extended kissing helices would remain structurally compatible with dsRNA-brick self-assembly (Fig. 3a). The original 6-bp bKL design^32^ was generated by motif fusion from the HIV-1 dimerization initiation site (DIS) KL complex^48,49^. In this design, one hairpin loop of the KL complex was converted into a bulge by replacing two unpaired 5ʹ adenines with an A-form helical stem (Supplementary Fig. 28). This yielded a kissing interaction between a bulge (with 6N-1A sequence) and a loop (2A-6N-1A; identical to the parent KL), whose complementary 6N segments together form the kissing helix. We then extended the 6N kissing segments to 7N, 8N, and 9N to generate the 7-, 8- and 9-bp bKL designs, respectively. This extension required structure-guided adjustment of the flanking unpaired adenines that connect the kissing helix to the three adjoining A-form helical branches (Fig. 3b; see also Supplementary Fig. 29 for the design principle and Supplementary Fig. 30 for exemplar sequence and predicted 3D models). For each of the 7-, 8-, and 9-bp bKL designs, we introduced two variants based on whether the loop contained the clamping 5ʹ and 3ʹ adenines. These two adenines are predicted to form a noncanonical A·A pair, based on the previous crystallographic study of the 6-bp bKL^50^ and NMR^51^ and cryo-EM^52^ studies of the HIV-1 DIS KL. Type A designs retain this A·A pair, whereas type B designs do not (Fig. 3b and Supplementary Fig. 30). For both 9-bp bKL designs, we further introduced an A·A pair connecting the non-coaxial branch helix to the bulge to accommodate the widened spacing across the major groove of the 9-bp kissing helix.

**Fig. 3.**
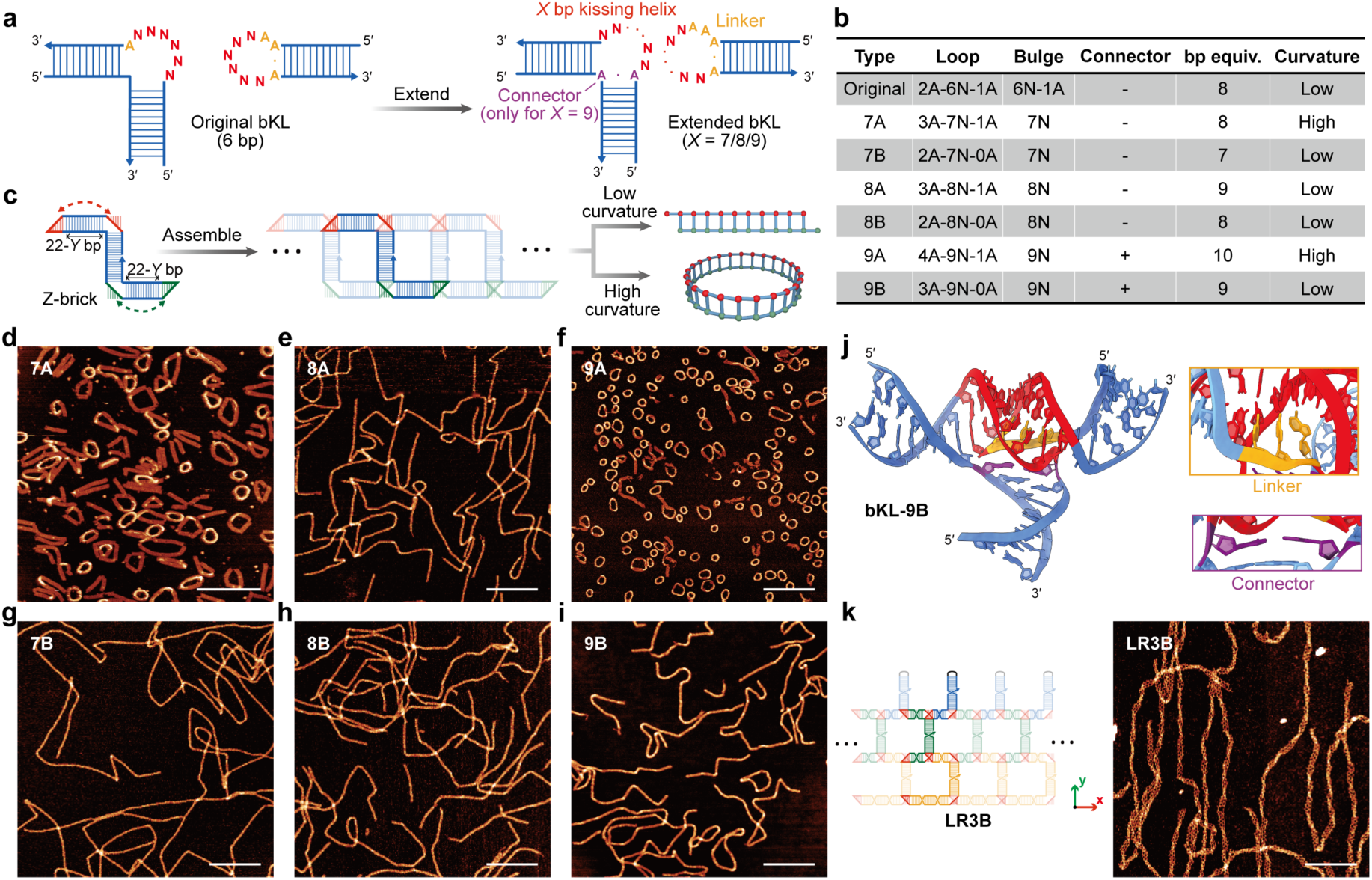
Extended bKL designs expand the orthogonal sequence space for the dsRNA bricks method. a, Design strategy for extending the original 6-bp bKL motif to *X*-bp bKL variants (*X* = 7, 8, or 9; the number of bp in the kissing helix). To accommodate the different lengths of the kissing helices in different variants, we adjusted the flanking adenines and polyA linkers that connect the kissing helix to the adjoining helical branches. For the 9-bp designs, an additional A·A connector was introduced between the bulge and the branched helix to accommodate the increased separation across the major groove of the kissing helix. **b,** Sequence patterns of the original and extended bKL designs. Type A and type B variants differ in the adenine pattern flanking the loop sequence and the resulting bp equivalents (Y) of the bKL motifs are X + 1 and X, respectively. **c,** Self-assembly of **Z-bricks** for evaluating the intrinsic curvature of bKL motifs. Beam lengths (22 − Y) are tuned to keep the effective along-rail spacing at 22 bp (exactly two helical turns) for every variant, so that **Z-bricks** assemble into extended ladder-like ribbons when the bKL motif has low intrinsic curvature, and into closed rings when it is more flexible or intrinsically curved. Red and green interaction sites indicate the two intermolecular bKL interfaces. **d–i,** AFM images of assembled products from **Z-bricks** based on **bKL-7A** (**d**), **bKL-8A** (**e**), **bKL-9A** (**f**), **bKL-7B** (**g**), **bKL-8B** (**h**) and **bKL-9B** (**i**). **j,** SimRNA model of the **bKL-9B**. The kissing helix (red) stacks coaxially with the adjacent helical stems. Insets show the polyA linker (orange box) and A·A connector (magenta box) regions, respectively. **k,** Design and AFM image of a three-brick ribbon **LR3B** incorporating **bKL-9B**. Scale bars, 200 nm.

To evaluate the six extended bKL motifs, we incorporated each motif into a Z-shaped brick, or **Z-brick** (Supplementary Fig. 31). The **Z-brick** contains a strut corresponding to an integer number of helical turns and is designed to self-assemble into ladder-like ribbons, allowing intrinsic out-of-plane curvature of the bKL motif to accumulate along the assembly (Fig. 3c). AFM imaging showed that all six **Z-brick** designs produced assembled products (Fig. 3d–i and Supplementary Fig. 32-37). **Z-bricks** incorporating **bKL-7A** (Fig. 3d) or **bKL-9A** (Fig. 3f) motifs predominantly formed closed rings, similar to the original 6-bp bKL^32^, implying substantial flexibility and/or intrinsic curvature of these bKL motifs. By contrast, the other four **Z-brick** designs formed extended ladder-like structures, suggesting reduced intrinsic curvature and greater rigidity of their respective bKL motifs.

Among the extended bKL candidates, 9-bp bKL was particularly attractive because its longer kissing helix substantially expands the orthogonal sequence space. We therefore implemented a graph-based workflow to select mutually orthogonal 9-bp bKL interaction pairs (see Supplementary Note I for sequence selection criteria and algorithm). Candidate 9-nt kissing sequences were generated with fixed base composition constraints, then filtered to remove self-complementary sequences. Pairwise off-target interactions were encoded as a conflict graph, in which nodes represent candidate sequences and edges represent predicted crosstalk. Selection of a large non-conflicting subset from this graph yielded more than 200 orthogonal 9-bp bKL sequences with 4 G/C and 5 A/U composition (Supplementary Fig. 38). This expanded interaction pool enabled the design of dsRNA-brick systems containing more than 100 standard bricks, with each brick carrying two loops and two bulges.

Coarse-grained simulations of the 9-bp bKL using SimRNA^53,54^ suggested that the 9-bp kissing helix adopts an A-form dsRNA helical geometry (Fig. 3j) and stacks coaxially with adjacent stems. To establish the compatibility of 9-bp bKL with the dsRNA bricks method, we incorporated it into a three-brick ribbon (**LR3B**) system. **LR3B** contains C-shaped bricks with 27-bp struts, corresponding to 2.5 helical turns, which are longer than the struts used in the previous architectures. This design produces a 55-bp repeating unit along each rail, corresponding to 5 helical turns (Fig. 3k and Supplementary Fig. 39). AFM imaging revealed well-formed **LR3B** assemblies, supporting the compatibility of 9-bp bKL with dsRNA brick assembly (Fig. 3k and Supplementary Fig. 40). Together, these results laid the foundation for constructing dsRNA-brick assemblies with substantially greater structural complexity.

### Finite-sized 2D assemblies containing up to 107 dsRNA bricks

To explore the assembly complexity enabled by the expanded pool of orthogonal 9-bp bKLs, we designed and constructed a finite-sized 2D assembly, **2D_107**, containing 107 distinct dsRNA bricks (Fig. 4a). Similar to the **LR3B** system, each dsRNA brick in **2D_107** is a C-brick with a 27-bp strut and beam lengths chosen to maintain an effective 55-bp along-rail spacing (Fig. 4b). The 107 dsRNA bricks were divided into three classes^44^ (Fig. 4c): core bricks (cBs), nucleating bricks (nBs), and boundary bricks (bBs). The cBs constitute the interior of the structure and contain beams of 18 or 19 bp. The nBs are derived from cBs by elongating one beam to 46 bp while retaining the other beam at the standard 18-bp length; this design allows two nBs together with one cB to form a closed ring that serves as a nucleation site for self-assembly (Supplementary Fig. 41). This feature was designed to lower the nucleation barrier and improve assembly kinetics^43^. Following the strategy used for the finite-sized assemblies described above, the bBs terminate growth at the edges of the structure: half-C-bricks are placed at the top boundary to block growth along the y direction, whereas for the left- and right-boundary bBs, non-interacting loop and bulge sequences replace the active bKL loops and bulges to prevent extension along the x-direction.

**Fig. 4.**
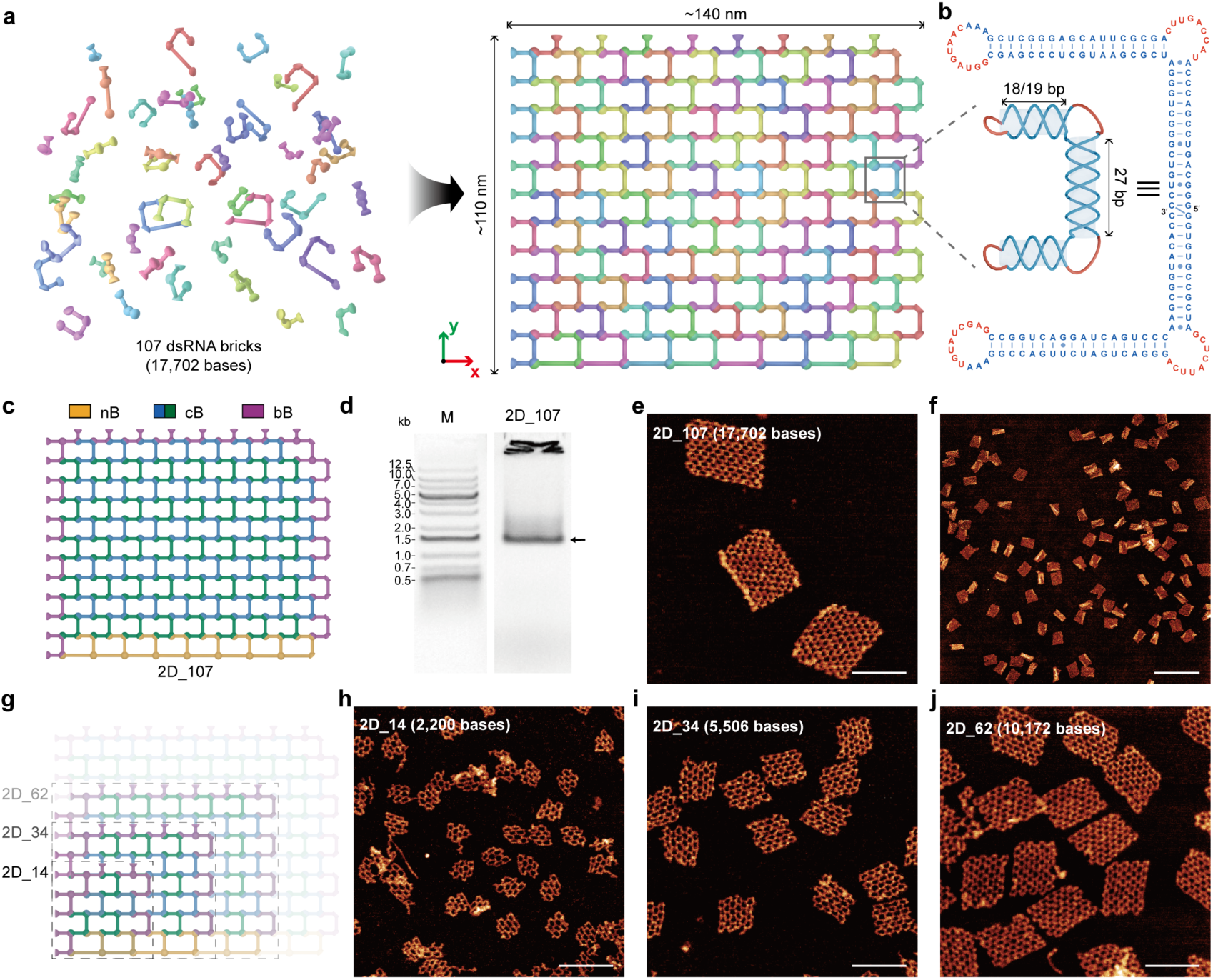
Finite-sized 2D assembly of dsRNA bricks containing up to 107 distinct components. **a**, Schematic of the assembly of **2D_107** from 107 distinct dsRNA bricks. Individual bricks, each carrying programmed bKL interaction sites, self-assemble into a finite rectangular lattice. The **2D_107** design contains 17,702 nucleotides, with expected dimensions of approximately 140 × 110 nm. **b**, Design and secondary structure of a representative dsRNA brick of **2D_107**. It contains 18- or 19-bp beams and a 27-bp strut, with four bKL-participating sequences (red) that mediate finite-sized self-assembly. **c**, Blueprint of **2D_107** colored according to tile types: nucleating bricks (nBs; orange), core bricks (cBs; blue and green distinguish bricks in alternating positions) and boundary bricks (bBs; purple). **d**, Native agarose gel analysis of **2D_107** assembly. The arrow indicates the target assembly band. Lane M, DNA ladder. **e**, **f**, Zoomed-in (**e**) and zoomed-out (**f**) AFM images of **2D_107** assemblies, showing well-defined finite rectangular lattices. **g**, Modular subset assemblies derived from the **2D_107** design, comprising 14, 34 and 62 bricks (**2D_14**, **2D_34** and **2D_62**, respectively, rendered with increasing transparencies). **h–j**, Representative AFM images of **2D_14** (**h**), **2D_34** (**i**) and **2D_62** (**j**). Scale bars, 100 nm (**e**, **h–j**) and 500 nm (**f**).

To facilitate the design of such finite-sized 2D assemblies, we developed a computational workflow for dsRNA-brick architecture generation (Supplementary Note II). Using this workflow, arbitrary 2D brick layouts can be created; for the 107-brick structure described here, a 9×12 rectangular blueprint was generated (Fig. 4a). Orthogonal bKL sequences were then assigned to the bulge and loop regions of all bricks within the blueprint (Supplementary Fig. 42), and full sequences for individual bricks were generated using NUPACK^55^ based on these bKL assignments and the prescribed secondary-structure specifications. In total, the complete **2D_107** assembly contains 17,702 nucleotides and forms an approximately 140 nm × 110 nm rectangular structure. To simplify RNA synthesis and purification, and to test progressively increasing structural complexity, we used a module strategy in which the full sequence map was partitioned into distinct regions^13^ and encoded in multiple plasmids (Supplementary Fig. 43). This modular design also enabled the construction of smaller rectangular assemblies containing 14, 34, and 62 dsRNA bricks (**2D_14**, **2D_34** and **2D_62**, respectively) by combining selected modules from the **2D_107** design with the corresponding bBs (Supplementary Fig. 44-46).

Plasmids encoding brick sequences from different modules were used for IVT to produce long RNA precursors ranging from 600 to 1,800 nt. After gel purification (see Methods for more details), the polycistronic RNA precursors were mixed according to the design blueprint and processed by RNase H using a universal guide DNA to release individual dsRNA bricks. The released dsRNA bricks were then annealed from 65 °C to 4 °C over 11 h. Native agarose gel electrophoresis showed a predominant target band for **2D_107**, suggesting efficient formation of the intended assembly (Fig. 4d). AFM imaging further confirmed the formation of well-defined finite-sized 2D lattices (Fig. 4e, f and Supplementary Fig. 47). Smaller finite-sized 2D nanostructures also assembled with high fidelity to the prescribed designs and programmed dimensions (Fig. 4g–j and Supplementary Fig. 48-50). These results highlight key intended advantages of the dsRNA bricks method, including improved structural stability of RNA building blocks and coordinated generation of near-stoichiometric dsRNA brick components from precursor transcripts. The assembly of **2D_107** further demonstrates the robustness of 9-bp bKL, as this structure requires 194 orthogonal bKL interactions to operate collectively within a single nanostructure.

Optimizing assembly conditions was crucial for maximizing the yield of large finite-sized 2D assemblies. For smaller structures, efficient self-assembly was achieved with 40 nM per dsRNA brick in 40 mM Tris buffer containing 1 mM free Mg^2+^. However, **2D_107** had a low yield under these conditions. Adding Na^+^ and increasing the Mg^2+^ concentration substantially improved the yield up to 58% (Supplementary Fig. 51), suggesting that Na^+^ and elevated Mg^2+^ concentrations reduce electrostatic repulsion and stabilize the bKL-mediated interactions, thereby promoting the formation of larger assembly products. Brick concentration was also an important parameter. Low brick concentrations led to aggregation products, as observed by agarose gel electrophoresis (Supplementary Fig. 52). This result suggests that sufficiently high brick concentrations are required to promote complete assembly, whereas low concentrations may allow partially assembled intermediates with exposed interaction sites to accumulate and aggregate with one another.

### Finite-sized 3D assemblies containing up to 118 dsRNA bricks

We next explored the dsRNA bricks method for assembling finite-sized 3D nanostructures (Fig. 5a). Similar to the 2D design, the cBs for 3D assembly retain a C-brick architecture with a 27-bp strut. To enable extension along the z direction, we adjusted the beam length to 15 or 16 bp, allowing neighboring bricks to assemble through bKL interactions with an approximately 90° dihedral angle (inset of Fig. 5a). Accordingly, we designed a 118-brick 3D structure, designated **3D_118**, containing 18,104 nucleotides in total and with expected dimensions of around 77 nm × 62 nm × 36 nm (Fig. 5b and Supplementary Fig. 53-54). The bricks in **3D_118** are arranged in four layers and fall into two orientation classes: bricks oriented in the x–y plane (x–y bricks) and bricks oriented in the x–z plane (x–z bricks). The bottom layer contains nBs with elongated beams to facilitate assembly nucleation. A computational workflow analogous to that used for the 2D designs was used to generate the architecture and sequences of the individual dsRNA bricks (Supplementary Note III).

**Fig. 5.**
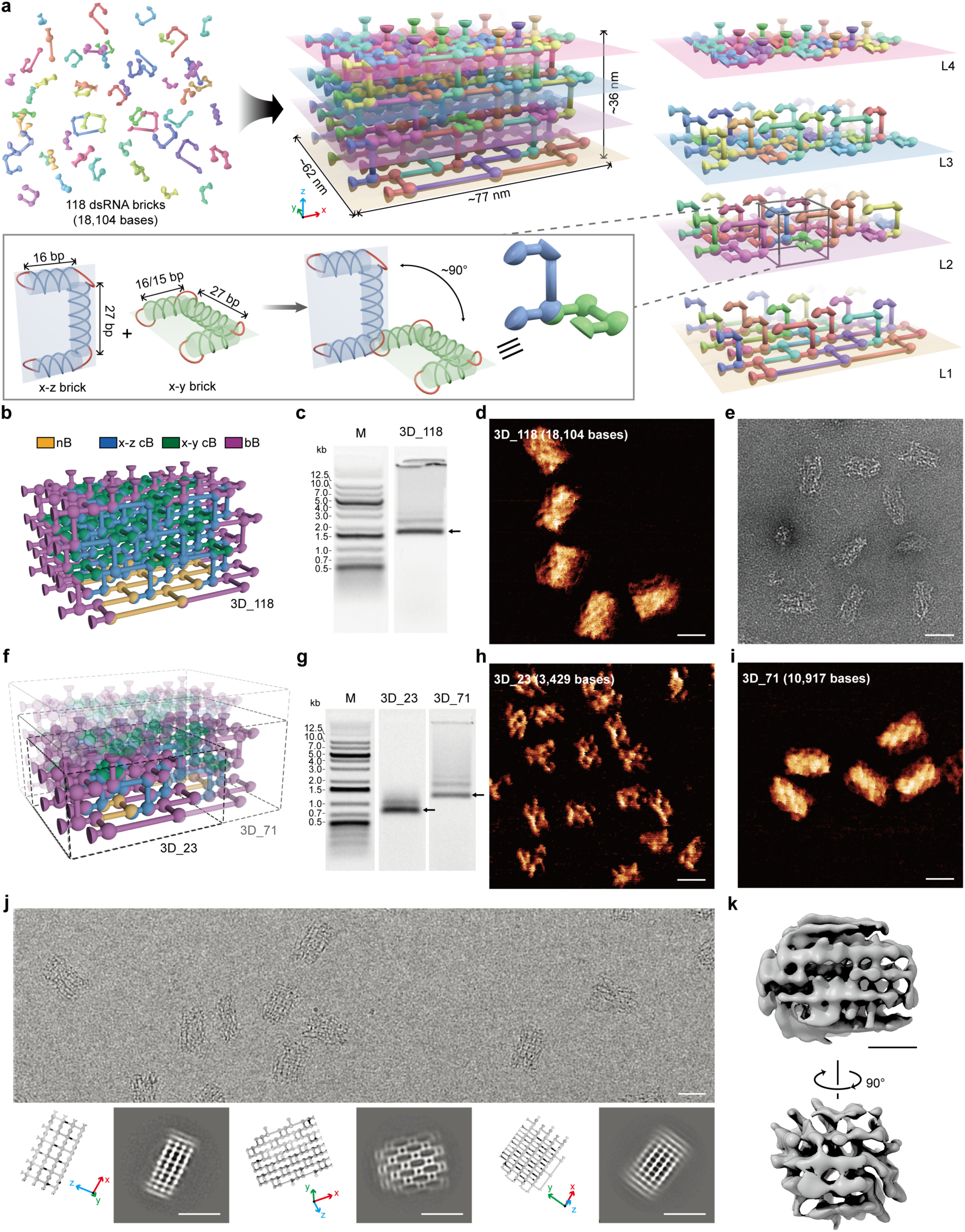
Finite-sized 3D assembly of dsRNA bricks containing up to 118 distinct components. **a**, Schematic of the assembly of **3D_118** from 118 distinct dsRNA bricks. **3D_118** is a four-layer nanostructure containing 18,104 nucleotides, with designed dimensions of approximately 77 × 62 × 36 nm. The four layers are shown individually at right, labeled L1 (bottom) to L4 (top). The inset shows the design parameters of the dsRNA bricks in **3D_118**. Orthogonal interactions between x–z and x–y bricks are mediated by bKLs positioned on 15- or 16-bp beams, producing an approximately 90° inter-brick dihedral angle. **b**, Blueprint of **3D_118** colored according to tile types: nucleating bricks (nBs; orange), core bricks oriented in the x–z and x–y planes (x–z cBs, blue and x–y cBs, green) and boundary bricks (bBs; purple). **c**, Native agarose gel analysis of **3D_118** assembly. The arrow indicates the target assembly band. **d, e**, AFM (**d**) and negative-stain TEM (**e**) images of **3D_118** assemblies. **f**, Modular subset assemblies derived from the **3D_118** design, comprising 23 and 71 bricks (**3D_23** and **3D_71**, respectively, rendered with increasing transparencies). **g**, Native agarose gel analysis of the **3D_23** and **3D_71** assemblies. **h, i,** AFM images of **3D_23** (**h**) and **3D_71** (**i**) assemblies. **j,** A representative cryo-EM micrograph of **3D_118** (top) and reference-free 2D class averages (bottom), each shown alongside a projection of the design model in the matching orientation. **k**, Two orthogonal views of the 3D reconstruction of **3D_118**. The reconstruction reveals multilayered reticular features consistent with the designed wireframe architecture. Scale bars, 50 nm (d, e, h–j) and 20 nm (**k**).

Similar to **2D_107**, the final target 3D structure, **3D_118**, was designed using a modular strategy in which the full 118-brick blueprint was partitioned into multiple plasmid-encoded modules. Using selected subsets of these modules together with the corresponding bBs, we also designed smaller intermediate structures, including a two-layer structure containing 23 bricks (**3D_23**, Supplementary Fig. 55) and a three-layer structure containing 71 bricks (**3D_71**, Supplementary Fig. 56). Native agarose gel electrophoresis and AFM imaging confirmed the formation of **3D_118** assemblies with high yield (Fig. 5c, d; see Supplementary Fig. 57-59 for assembly-condition optimization). Negative-stain transmission electron microscopy (TEM) further revealed reticular 3D architectures consistent with the design (Fig. 5e and Supplementary Fig. 60). Smaller finite-sized 3D nanostructures likewise formed well-defined assemblies with dimensions consistent with the programmed architectures (Fig. 5f–i, Supplementary Fig. 61-64).

To further validate the assembly of **3D_118**, we performed single-particle cryo-electron microscopy (cryo-EM; see Supplementary Fig. 65 for a representative micrograph). The resulting 2D class averages revealed features consistent with the design model in multiple orientations (Fig. 5j and Supplementary Fig. 66). Although the 3D reconstruction of **3D_118** reached only a modest overall resolution of 39.5 Å (Supplementary Fig. 67), likely due to the flexibility of its wireframe architecture, in which each edge is formed by a single RNA duplex, the map still revealed multilayered reticular features consistent with the intended structure (Fig. 5k). To our knowledge, **3D_118** represents the largest and most compositionally complex fully addressable finite-sized RNA nanostructure reported to date, demonstrating the potential of the dsRNA bricks method to construct sophisticated RNA architectures with addressable complexity^56^ approaching that of established DNA-based platforms.

## Discussion

The dsRNA bricks method provides a robust framework for the synthesis of complex RNA nanostructures. For decades, a central challenge in RNA nanotechnology has been achieving the structural complexity and scalability already demonstrated in DNA-based systems, particularly because large libraries of distinct synthetic ssRNA strands are costly to produce and insufficiently stable under multicomponent assembly conditions. Inspired by natural RNA maturation pathways, we adopted a polycistronic precursor strategy, in which many distinct dsRNA bricks are encoded within a single IVT-generated transcript and released by enzymatic processing. This strategy enables the scalable production of diverse RNA components while allowing each component to retain a predominantly double-stranded architecture that enhances stability during assembly and storage (Supplementary Fig. 68). Using this approach, we constructed **3D_118**, a programmable RNA nanostructure comprising 118 distinct dsRNA bricks and 18,104 nucleotides. To our knowledge, this represents the largest and most complex programmable RNA nanostructure reported to date (Supplementary Fig. 1), matching the complexity of conventional DNA origami^11^.

Our work also introduces a distinct design strategy for nucleic acid nanostructures. Self-folding and self-assembly have long been the defining principles of nucleic acid-based nanostructure design. Two strategies lie at the extremes of this spectrum: single-stranded nucleic acid origami^29,36^ and DNA bricks^14,15^. The former relies on the self-folding of a long single strand to form nanostructures; the latter depends on the self-assembly of many short, initially unstructured DNA strands. Scaffolded DNA origami, lying between these extremes, can be viewed as a self-assembly-assisted self-folding process. In contrast, our dsRNA bricks method represents a different intermediate strategy that is particularly well suited to RNA: self-folding-facilitated self-assembly, in which intramolecular folding (or secondary-structure formation) first yields the dsRNA bricks that then self-assemble through inter-brick programmable bKL interactions. Furthermore, each individual brick is itself a programmable unit: its shape, size, and valence can be controlled by secondary structure design, allowing the dsRNA brick concept to be extended beyond simple duplex-based motifs to more sophisticated folded RNA architectures, including single-stranded RNA origami^29,36,52^ and natural RNA-only nanocages^57–60^.

One of the key advantages of RNA over DNA is its potential for intracellular production and function. Recent studies have reported the formation of single-stranded RNA origami^61,62^ or RNA nanostar condensates^63^ in cells, but the intracellular expression and assembly of complex RNA structures remain challenging. Our dsRNA bricks exhibit high stability, and their local secondary structure does not rely on thermal annealing, offering additional flexibility in potential applications (Supplementary Fig. 69). By integrating the dsRNA bricks strategy with cell-compatible RNA expression and processing modules such as tRNA scaffolds^64,65^ and Tornado^66^, artificial RNA nanoarchitectures or devices may be generated, stabilized and programmed to self-assemble into functional architectures inside cells. Future work will need to improve control over intracellular processing, optimize nucleation and assembly under cellular conditions, and evaluate how predominantly dsRNA architectures interact with innate immune sensors^67^. Addressing these challenges will help transform dsRNA bricks from a robust in vitro assembly platform into a generalizable framework for programmable RNA materials^68,69^ and RNA-based biological devices^70,71^.

## Methods

### Design of dsRNA bricks

Sequences of individual dsRNA bricks were designed using NUPACK based on prescribed secondary structures while minimizing unintended intramolecular folding^55^. RNA secondary structures were drawn using VARNA^72^. ChimeraX^73^ was used to visualize bKL structures. Orthogonal bKL sequences were selected using custom Python scripts according to the criteria described in Supplementary Note I. Finite-sized 2D and 3D dsRNA-brick assemblies were designed using the custom software workflow described in Supplementary Note II and III.

### Preparation of RNA

DNA templates were purchased either as gBlocks Gene Fragments from Integrated DNA Technologies (IDT) or as plasmids ordered from WuXi Qinglan Biotech. All DNA oligonucleotide primers were purchased from IDT. DNA templates were amplified using Platinum™ SuperFi II PCR Master Mix (Invitrogen) according to the recommended protocol provided by the manufacturer.

RNAs were synthesized by in vitro transcription using the HiScribe® T7 High Yield RNA Synthesis Kit (New England Biolabs). IVT products were optionally treated with Turbo DNase (Invitrogen™) to remove DNA templates for 2D and 3D dsRNA bricks assembly. Precursor RNAs were purified primarily by denaturing PAGE containing 7 M urea, followed by ethanol precipitation and resuspension in nuclease-free water. Long RNAs used for finite-sized 2D and 3D assemblies were purified by denaturing agarose gel electrophoresis using 1.5% Certified RCR Agarose (Bio-Rad) containing 2% linear polyacrylamide and 7 M urea followed by extraction and cleanup using RNA Clean and Concentrator kits (Zymo).

For RNase H-mediated processing, 50 μl 1 μM RNA precursors were mixed with the universal guide DNA at a 1:2.5 molar ratio in 1× RNase H buffer. The mixtures were annealed from 65 °C to 4 °C at a ramp rate of −1 °C per min. Then, 1 μl RNase H (New England Biolabs) was added to cleave the RNAs at the linker regions at 37 °C for 1 hour. The processed products were purified using 10-kDa molecular-weight-cutoff Amicon Ultra centrifugal filters.

### RNA nanostructure assembly

For periodic brick assembly, processed dsRNA bricks were diluted to 100 nM per brick in 1× TA buffer (40 mM Tris, 20 mM acetic acid at pH 7.0) with 1 mM MgCl_2_. Then, the samples were annealed from 65 °C to 4 °C over 3 h.

For the finite-sized 2D assemblies **2D_14**, **2D_34** and **2D_62**, processed dsRNA bricks were diluted to 40 nM per brick in 40 mM Tris buffer (pH 7.0) containing 1 mM MgCl_2_. For **2D_107**, processed dsRNA bricks were diluted to 100 nM per brick in 40 mM Tris buffer (pH 7.0) containing 100 mM NaCl and 30 mM MgCl_2_. For the finite-sized 3D assemblies **3D_23** and **3D_71**, processed dsRNA bricks were diluted to 40 nM per brick in 40 mM Tris buffer (pH 7.0) containing 100 mM NaCl and 10 mM MgCl_2_. For **3D_118**, processed dsRNA bricks were diluted to 40 nM per brick in 40 mM Tris buffer (pH 7.0) containing 100 mM NaCl and 30 mM MgCl_2_. Then, samples were annealed from 65 °C to 4 °C over 11 h. Brick concentration and buffer composition were further optimized for individual systems.

### Denaturing polyacrylamide gel electrophoresis

Denaturing polyacrylamide gel electrophoresis (PAGE) was performed using 8% polyacrylamide gels prepared from a 29:1 acrylamide/bisacrylamide solution in 0.5× TBE buffer containing 7 M urea. Gels were run at 50 °C for 2 h at 200 V and then stained with GelRed. Gel images were acquired using a Bio-Rad Gel Doc XR+ imaging system.

### Agarose gel electrophoresis

Agarose gels (0.8%) were prepared with Certified PCR Low-Melt Agarose (Bio-Rad) in 0.5× TBE buffer supplemented with 10 mM Mg^2+^ and prestained with GelRed. The gels were run for 2.5 h at 80 V in 0.5× TBE buffer supplemented with 10 mM Mg^2+^. For characterization, the gels were imaged using a Bio-Rad Gel Doc XR+ Imaging System. For purification, the major target band was excised under ultraviolet illumination and recovered using a freeze’N’Squeeze DNA gel extraction spin column (Bio-Rad). Gel images were processed and analyzed using ImageJ. Yield was calculated as the background-corrected intensity of the main product band divided by the total lane intensity below the wells.

### AFM imaging

AFM images were collected in air. The RNA assemblies were diluted to 5 nM in 1× TAE buffer supplemented with 10 mM Mg^2+^. A 10 μl aliquot of the diluted sample was deposited onto freshly cleaved mica (Ted Pella) and incubated for 1 min. The specimen was dried with compressed air. The mica surface was then washed with 40 μl of 3 mM magnesium acetate solution and dried again with compressed air. Samples were imaged in “ScanAsyst in Air” mode using SNL-10 probes (tip C; Bruker) on a Dimension FastScan AFM (Bruker). AFM images were processed and analyzed using Gwyddion.

### Coarse-grained RNA structure modeling

Coarse-grained simulations of bKLs were performed using SimRNAweb v2.0^53,54^. The three RNA strands directly involved in each bKL motif were included in the simulations, comprising one loop-containing strand and two strands forming the adjacent bulged helical segment. Secondary structures predicted from the corresponding RNA sequences were supplied to SimRNAweb as soft secondary-structure restraints. All simulations were performed using the default SimRNAweb parameters. The resulting coarse-grained conformational models were reconstructed as all-atom RNA structures by the server and used to assess the local geometric compatibility of the designed bKL interactions, including the relative orientation of the connected helical segments and the accessibility of the bKL interface.

### TEM imaging

Samples were negatively stained for TEM imaging. The annealing mixture was diluted to 40 nM per brick in 40 mM Tris buffer (pH 7.0) containing 100 mM NaCl and 30 mM MgCl_2_. The assembly product was purified by agarose gel electrophoresis and recovered using freeze’N’Squeeze spin columns. The purified RNA nanostructure was then diluted to 6 nM per brick. A 3 μl aliquot of the purified sample was adsorbed onto ultrathin carbon-coated 400-mesh copper grids that had been glow-discharged for 1 min at 15 mA using a Pelco easiGlow glow-discharge system. The sample was stained with 5 μl of freshly prepared 2% aqueous uranyl formate containing 25 mM NaOH. Excess liquid was wicked away with Whatman filter paper. Images were acquired on an ARM200F TEM (JEOL) operated at an accelerating voltage of 120 kV using a charge-coupled device camera at ×50,000 magnification.

### Cryo-EM imaging and single-particle reconstruction

For cryo-EM analysis of the **3D_118** RNA nanostructure, the annealing mixture was diluted to 40 nM per brick in 40 mM Tris buffer (pH 7.0) containing 100 mM NaCl and 30 mM MgCl2. The assembly product was purified by agarose gel electrophoresis and recovered using freeze’N’Squeeze spin columns. The purified RNA nanostructure was then diluted to 6 nM per brick. Next, 3 μl of sample solution was applied to ultrathin carbon on Cu grids (CF300-Cu-UL, EMS), blotted for 10 s, immediately frozen in liquid ethane and subsequently cooled in liquid nitrogen. Cryo-EM data were collected using a Thermo Fisher Scientific Titan Krios G3i Cryo-TEM at 300 kV. Approximately 4,565 micrographs were collected at a nominal magnification of ×64,000 using a Gatan K3 camera with the energy slit set to 30 eV.

Single-particle reconstruction was performed using RELION^74^. In general, cryo-EM data were processed following a standard single-particle analysis workflow^75^. Movie frames were first motion-corrected and dose-weighted, followed by CTF estimation from the corrected micrographs. Micrographs with poor ice quality, strong drift, or poor CTF estimation were removed. Particles were then picked, extracted, and cleaned by iterative 2D classification. High-quality 2D classes were selected for initial model generation, 3D classification, and 3D refinement. A total of 32,285 particles were selected for the final 3D reconstruction, and the resolution was estimated using the gold-standard FSC criterion. The reconstructed density map was visualized using UCSF ChimeraX^73^.

## Data availability

Plasmids generated for this study have been deposited in the Addgene database at https://www.addgene.org/Di_Liu/.

## Code availability

Custom code developed for this study, along with future updates, is available via GitHub at https://github.com/DiLiuLab/dsRNA-brick.

## Acknowledgements

This work was supported by NSF grants (CMMI-1333215, CMMI-1344915 and CBET-1729397), AFOSR grant (MURI FATE, FA9550-15-1-0514), NIH grant (5DP1GM133052) and Molecular Robotics Initiative fund from the Wyss Institute to P.Y. D.L. and H.Y. acknowledge financial support from Arizona State University.

## Contributions

D.L., P.Y. and L.C. conceived and designed the study. L.C. and D.L. designed, prepared and characterized the RNA nanostructures and drafted the manuscript. Z.L., J.Y., S.D., C.L., G.W., X.T. and Al. P. participated in the preparation of RNA nanostructures. K.D., Ab.P., D.S. and P.-L.C. prepared the cryo-EM samples. J.C.-Y.W. performed the cryo-EM experiments and carried out 3D reconstruction. L.C., D.L., Y.Q. and S.W. developed software. D.L., P.Y., H.Y. and D.W. supervised the project. All authors analyzed the data and commented on the manuscript.

