## Supplemental information for "Programmable Self-Assembly of RNA Nanostructures with >100 Unique Components"

### Supplementary Information

### Figures

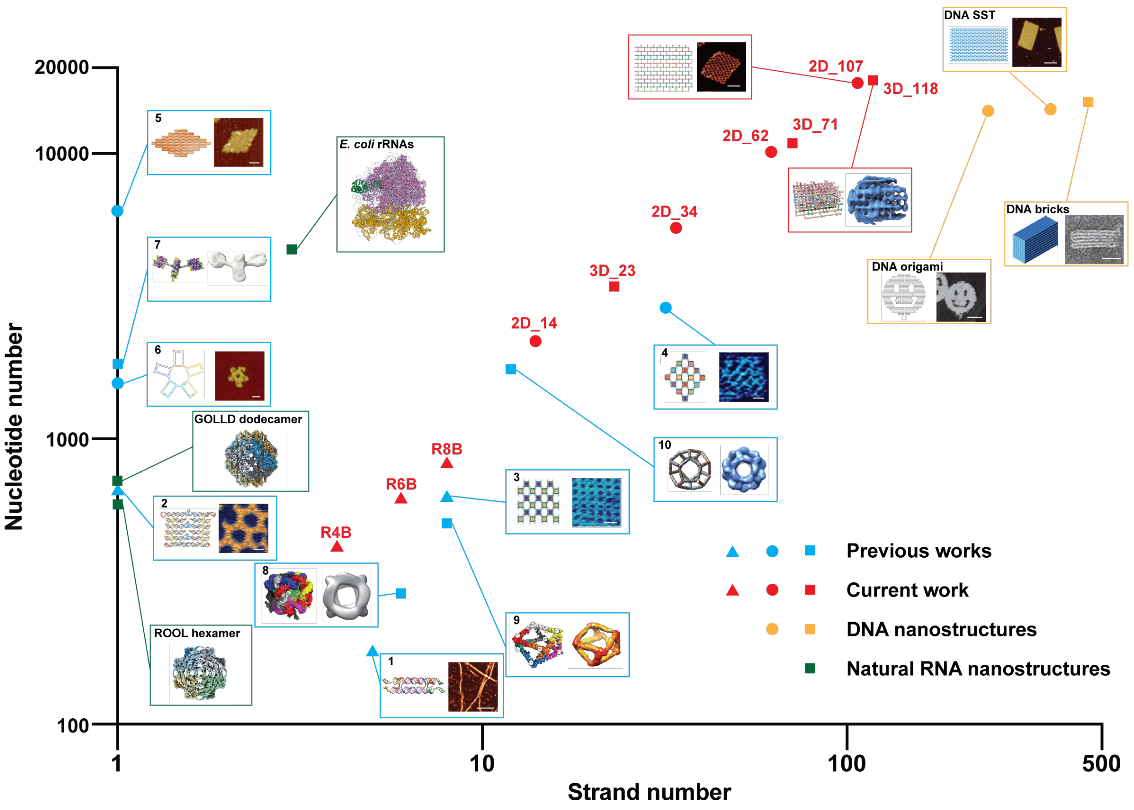

**Supplementary Fig. 1 Comparison of previously reported nucleic acid nanostructures with those presented in this work.** ▲, ●, and ■ represent periodic self-assembly, finite-sized 2D self-assembly, and finite-sized 3D self-assembly, respectively. Blue symbols denote previously reported RNA nanostructures, while red symbols indicate the nanostructures achieved in this study. Orange symbols denote the DNA nanostructures, including smiley-shaped DNA origami[1], 24H×28T DNA single-stranded tiles (SSTs)[2] and 6H×10H×128B DNA bricks[3]. Green symbols denote for natural RNA nanostructure, including rRNAs[4] (3 strands, 4566-nt) from *E. coli*., ROOL hexamer[5] (1 strand, 580-nt) from *Enterococcus faecalis* (*Efa*) and GOLLD dodecamer[5] (1 strand, 700-nt) from *Streptococcus agalactiae* (*Sag*). Structures: 1, RNA double crossover tile[6] (5 strands, 180-nt; scale bar, 200 nm); 2, ssRNA origami[7] (1 strand, 660-nt; scale bar, 20 nm); 3, tectoRNA lattice[8] (8 strands, 624-nt; scale bar, 20 nm); 4, 4×4 tectoRNA lattice[8] (32 strands, 2899-nt; scale bar, 20 nm); 5, rhombus-shaped ssRNA origami[9] (1 strand, 6337-nt; scale bar, 20 nm); 6, RNA 5-petal flower[10] (1 strand, 1571-nt; scale bar, 20 nm); 7, 16-helix satellite ssRNA origami[11] (1 strand, 1832-nt); 8, RNA cube[12] (6 strands, 288-nt); 9, RNA square antiprism[13] (8 strands, 506-nt); 10, RNA dodecahedron[14] (12 strands, 1760-nt). Scale bars for 2D_107, DNA origami and DNA SST, 50 nm. Scale bar for DNA brick, 20 nm.

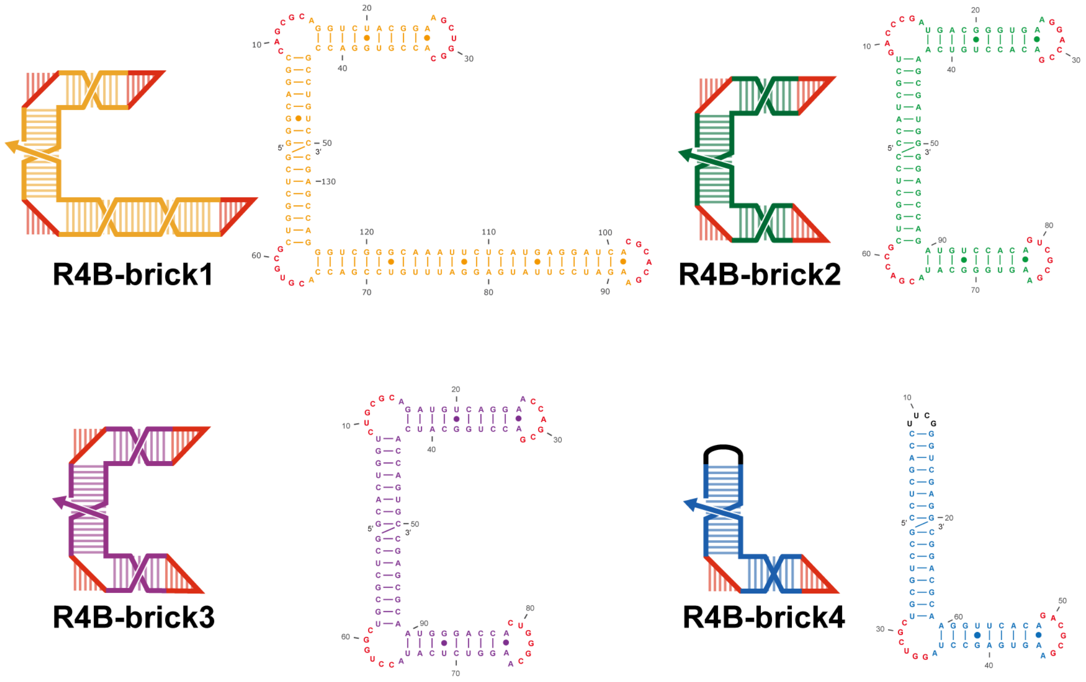

**Supplementary Fig. 2 Schematic of individual dsRNA bricks in R4B.**

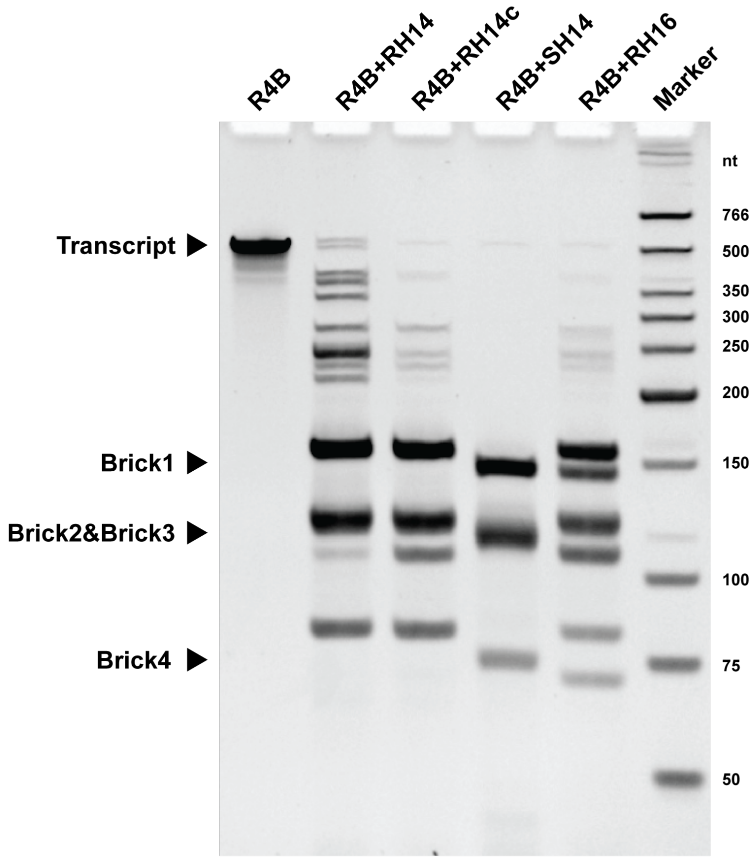

**Supplementary Fig. 3 Gel analysis for RNase H cleavage with different guide DNA sequences.**The transcript precursor (R4B) is mixed with different types of guide DNA sequences for RNase H cleavage. (RH14: 14-nt long complementary sequence with 8-nt 2'-O-methyl ribonucleotides in the middle. RH14c: Higher concentration of RH14. SH14: 14-nt long complementary sequence with all deoxyribonucleotides. RH16: 16-nt long complementary sequence with 8-nt 2'-O-methyl ribonucleotides in the middle.) The 14-nt guide DNA SH14 shows the highest cleavage efficiency and gives clean bands for individual bricks.

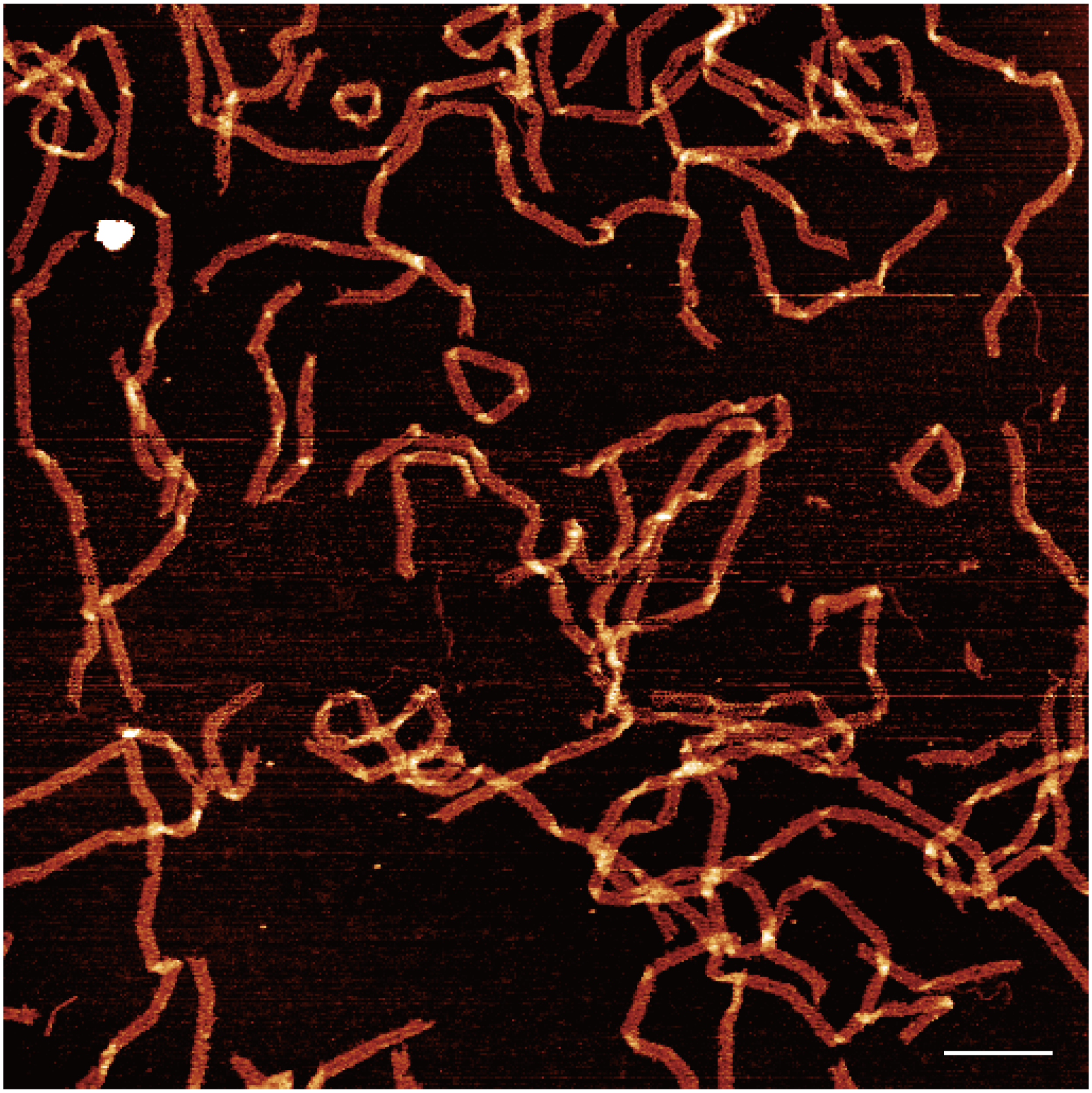

**Supplementary Fig. 4 Zoomed-out AFM image of R4B.** Scale bar, 200 nm.

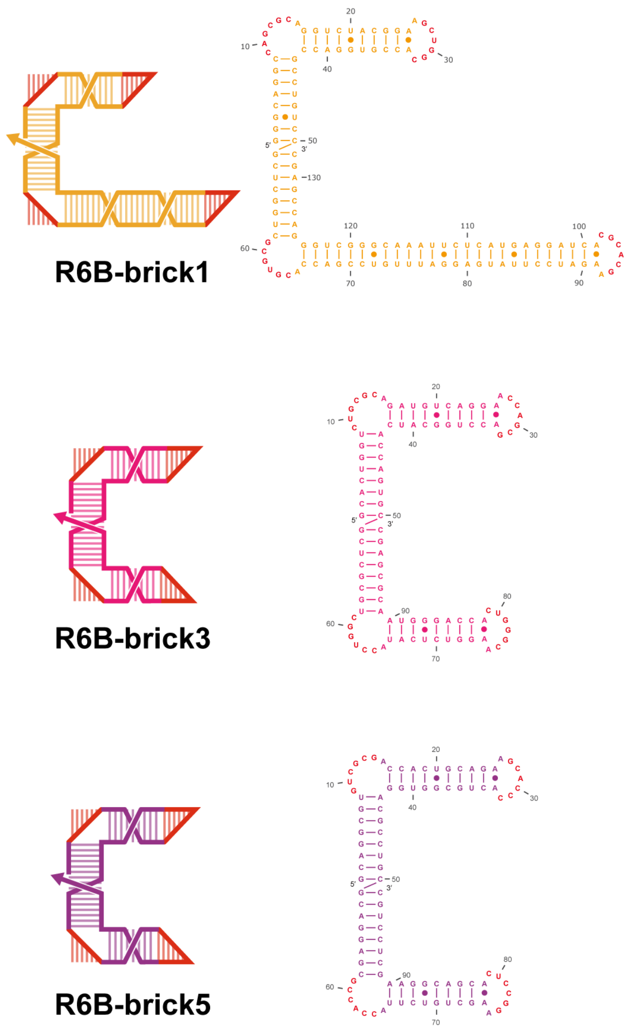

**Supplementary Fig. 5 Schematic of individual dsRNA bricks in R6B.**

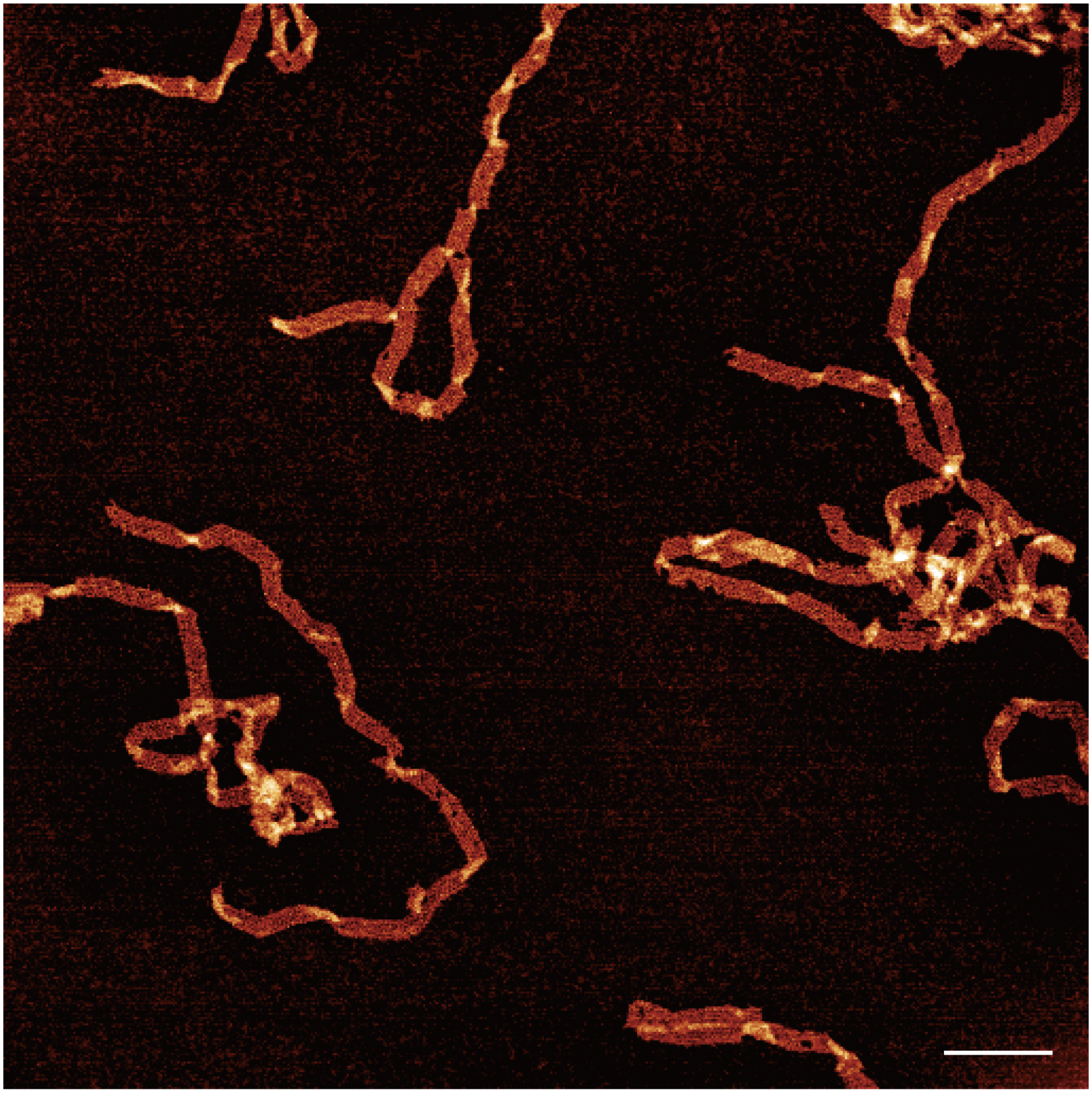

**Supplementary Fig. 6 Zoomed-out AFM image of R6B.** Scale bar, 200 nm.

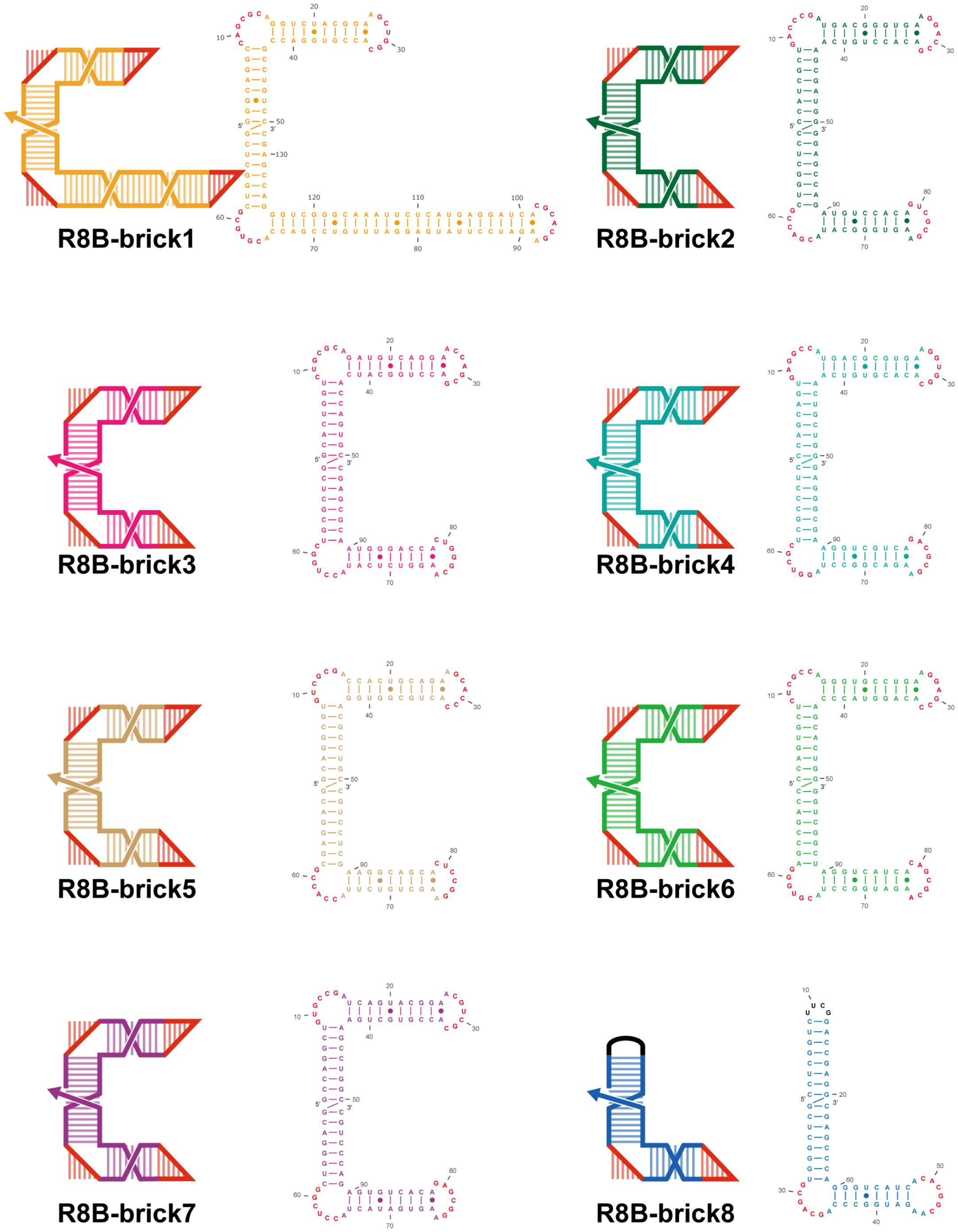

**Supplementary Fig. 7 Schematic of individual dsRNA bricks in R8B.**

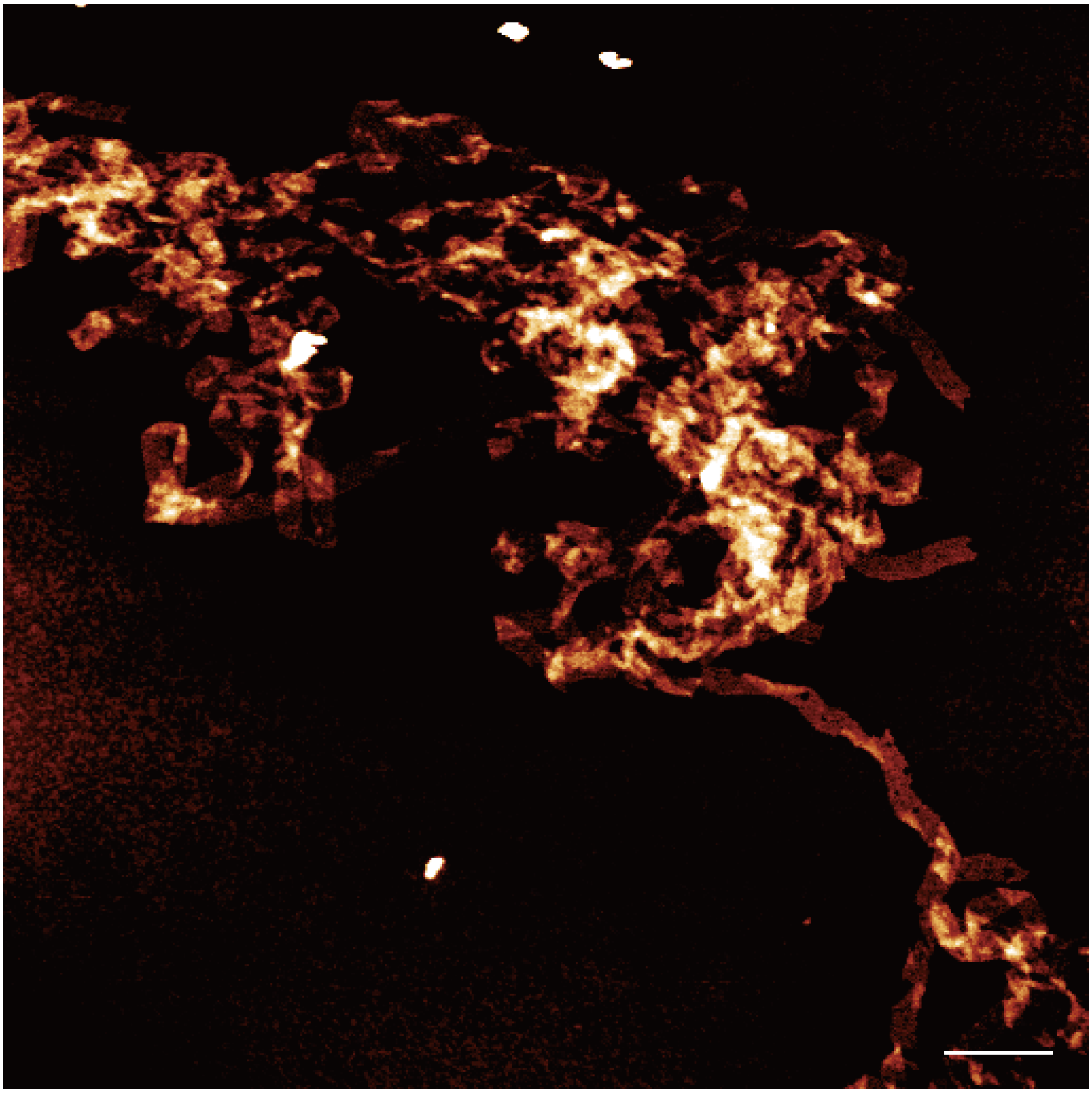

**Supplementary Fig. 8 Zoomed-out AFM image of R8B.** Scale bar, 200 nm.

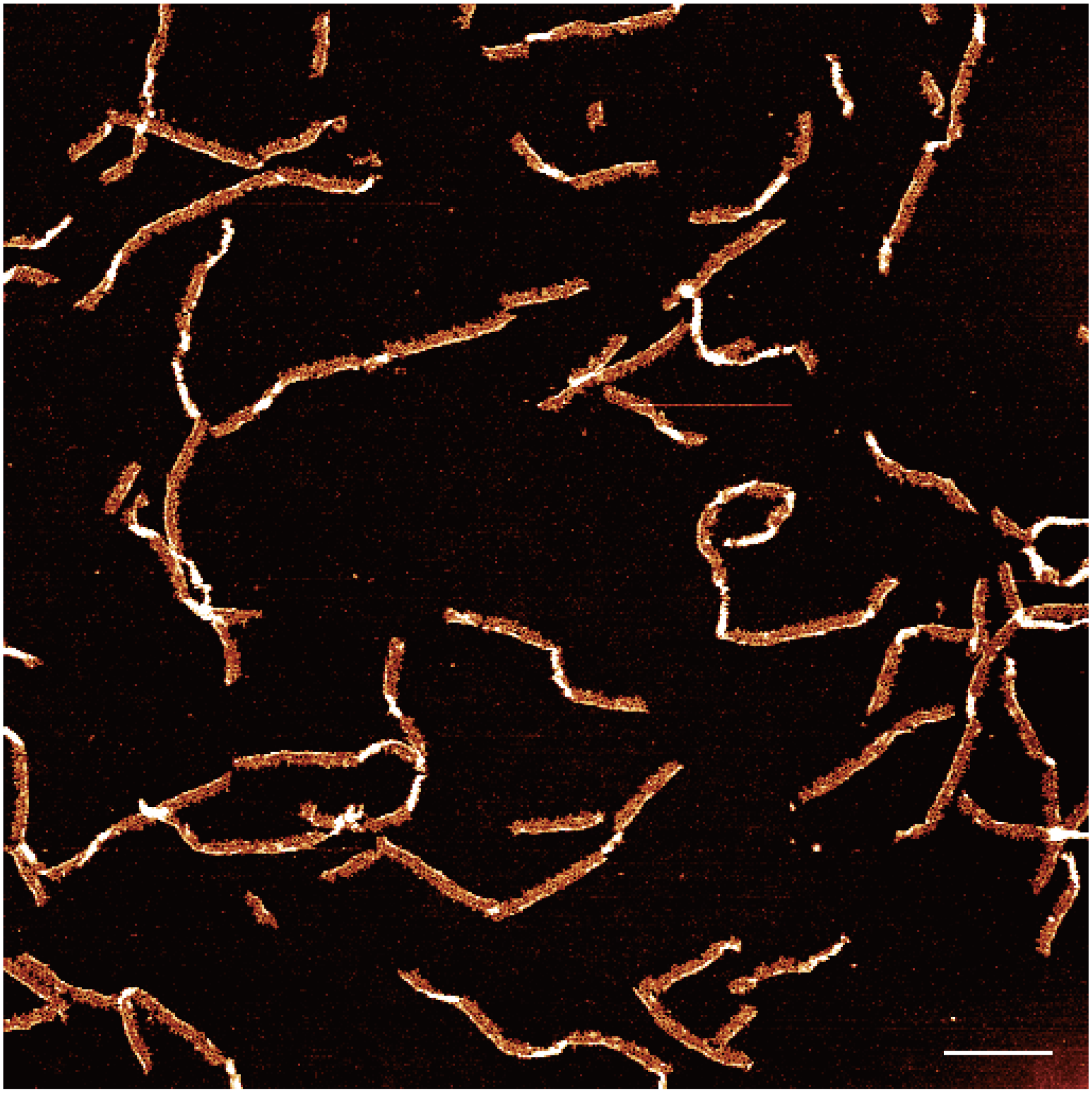

**Supplementary Fig. 9 Zoomed-out AFM image of R4×3.** Scale bar, 200 nm.

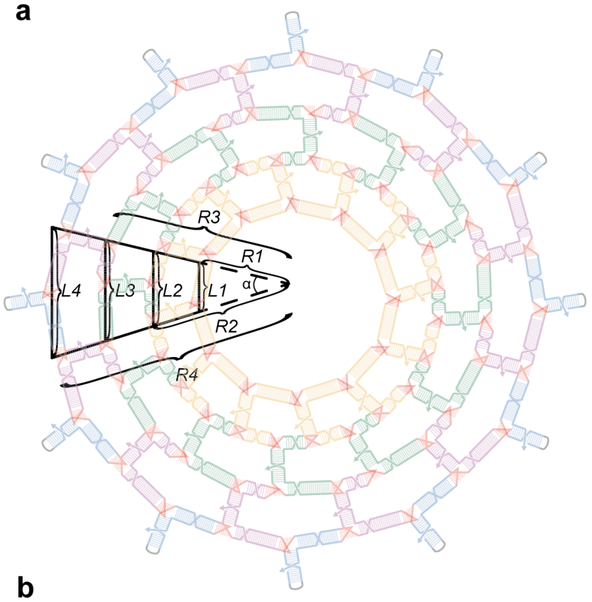

$$\frac{L_1}{R_1} = \frac{L_2}{R_2} = \frac{L_3}{R_3} = \frac{L_4}{R_4} = \frac{\Delta L}{\Delta R} = \frac{\alpha}{180^0}\pi$$

$$L_4 - L_3 = L_3 - L_2 = L_2 - L_1 = 11 \text{ bp}$$

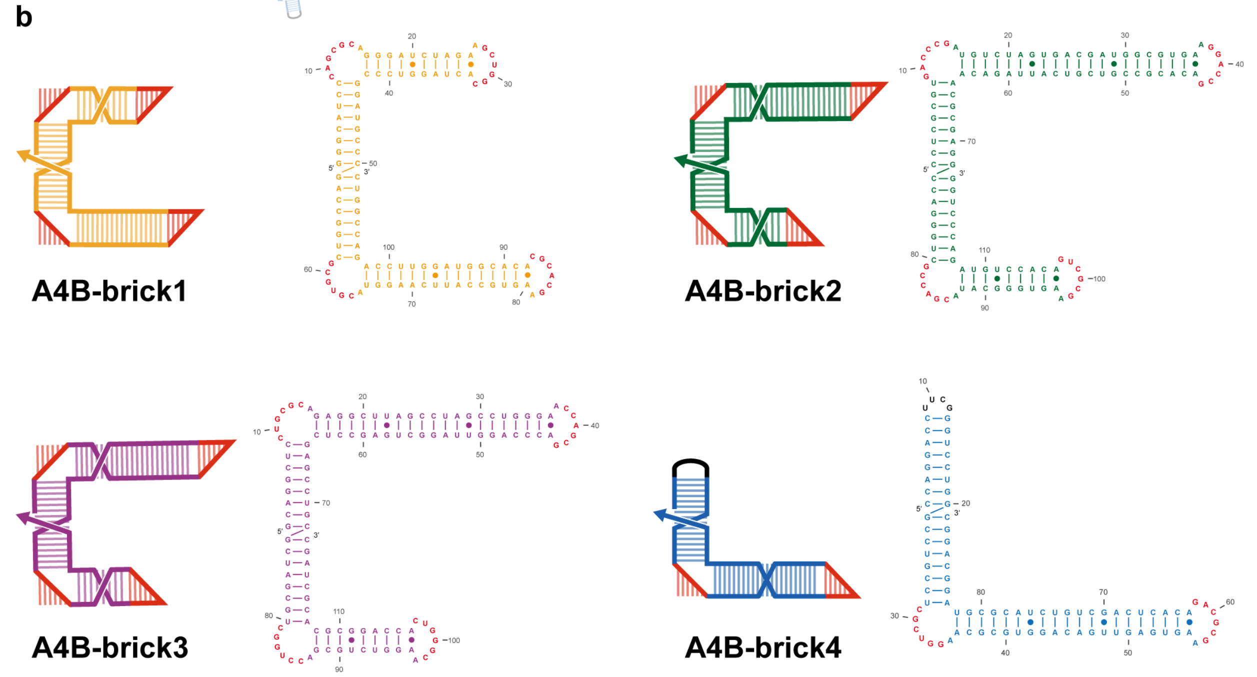

**Supplementary Fig. 10 Schematic of individual dsRNA bricks in A4B.** The length of repeating units in each concentric layer ($L_i$) is defined by the sum of the adjacent beam segments and bKLs (each bKLs is equivalent to an 8-bp duplex). Accordingly, $L_1 = 14 + 8 = 22$bp (one 14-bp beam + one bKLs), $L_2 = 9 + 8 + 8 + 8 = 33$bp (one 9-bp beam from brick1, one 8-bp beam from brick2, and two bKLs), $L_3 = 20 + 8 + 8 + 8 = 44$bp (one 20-bp beam from brick2, one 8-bp beam from brick3, and two bKLs), and $L_4 = 20 + 19 + 8 + 8 = 55$bp (one 20-bp beam from brick3, one 19-bp beam from brick4, and two bKLs). Target curvature is imposed by maintaining a constant arc-length-to-radius ratio across layers, with a fixed inter-layer arc-length increment $\Delta L = 11$bp. The radial spacing is set by the strut length (16 bp) plus the helix diameter expressed in base pairs, $\Delta R \approx 20.3$ bp, corresponding to an angular increment of $\alpha \approx 31°$per repeating unit. This predicts $\sim 360°/31° \approx 11.6$units per full turn, indicating an approximately dodecagon in theory.

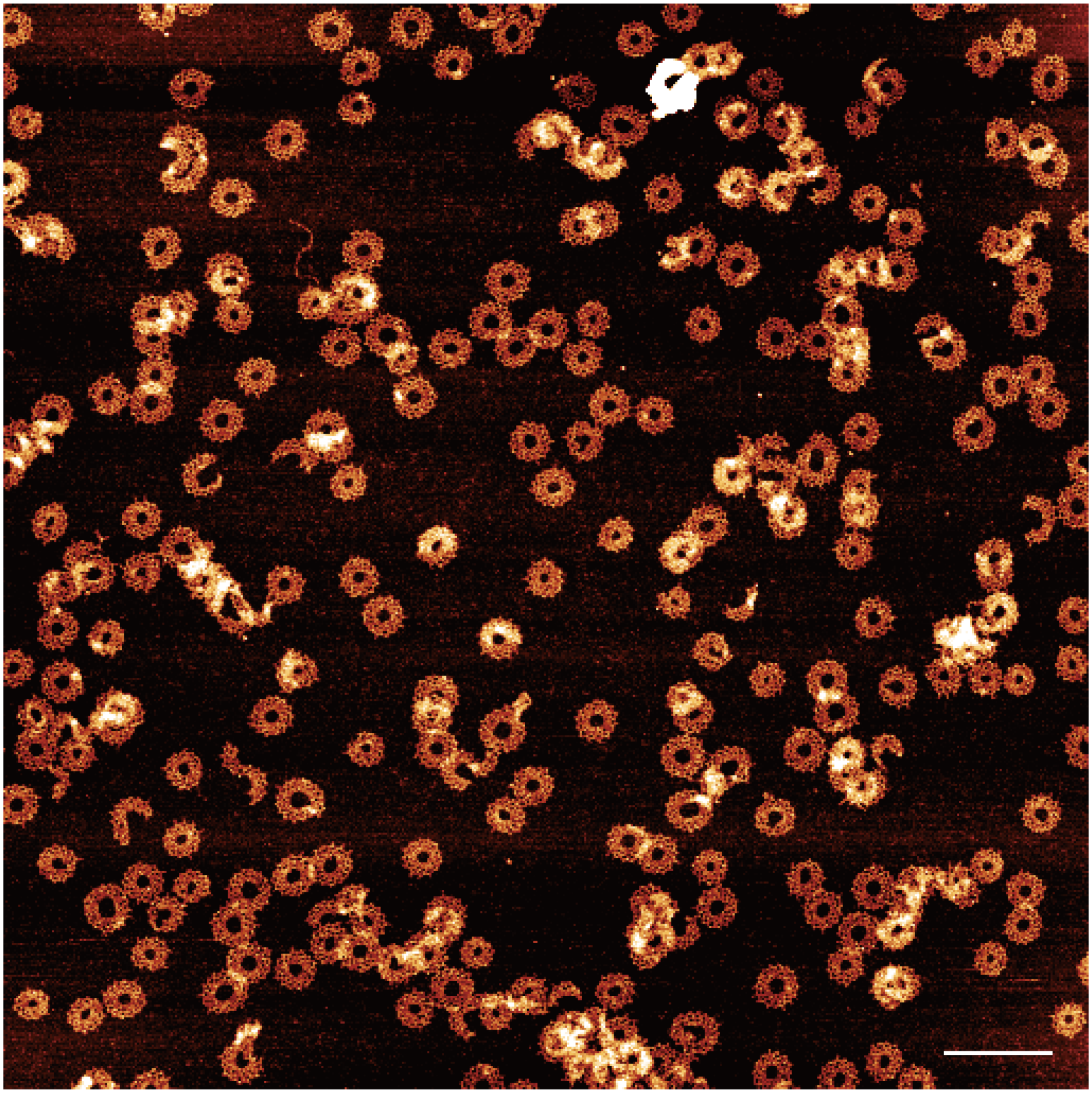

**Supplementary Fig. 11 Zoomed-out AFM image of A4B.** Scale bar, 200 nm.

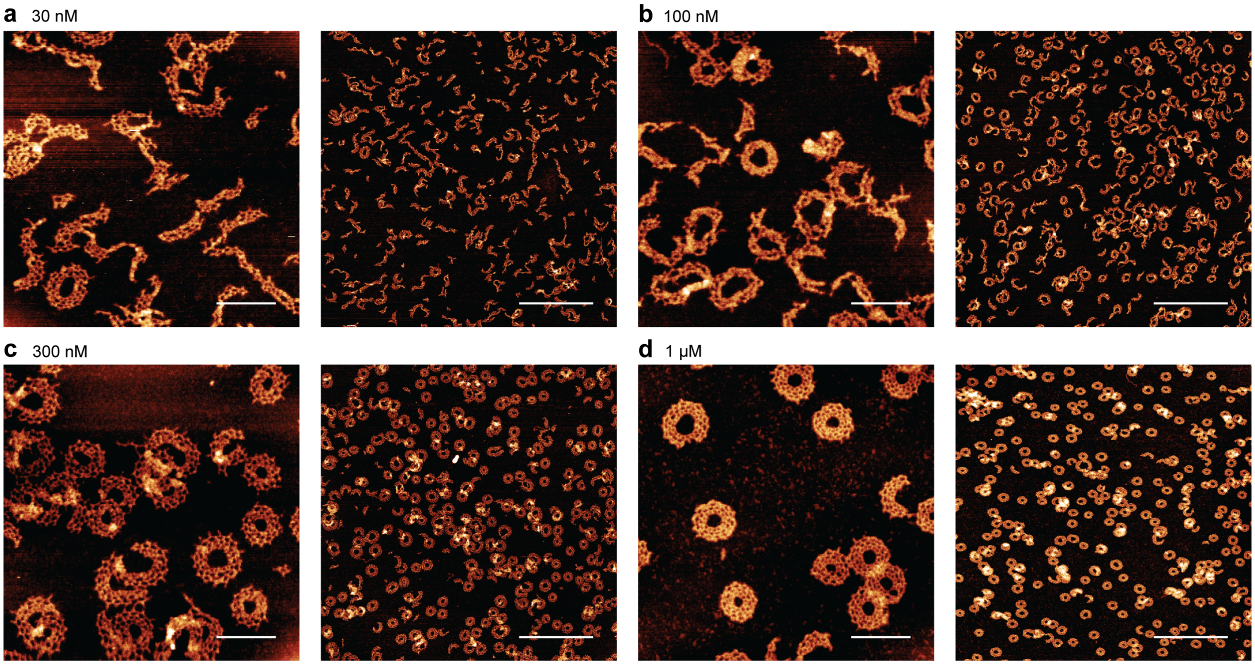

**Supplementary Fig. 12 Effect of brick concentration on A4B annulus assembly.** AFM images of A4B assemblies formed at brick concentrations of 30 nM (**a**), 100 nM (**b**), 300 nM (**c**) and 1 μM (**d**). At lower brick concentrations, the assembly products predominantly consisted of partially closed or extended ribbon-like structures. Increasing brick concentration promoted the formation of closed annuli, with more complete ring-like assemblies observed at higher concentrations. Scale bars, 100 nm (left) and 500 nm (right).

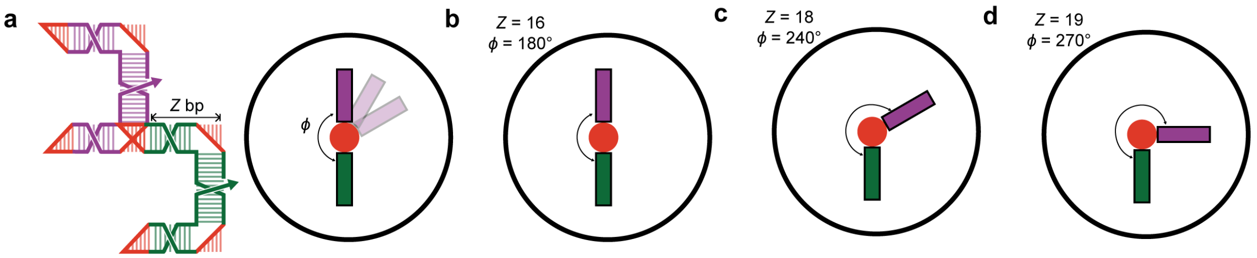

**Supplementary Fig. 13 Programming inter-brick geometry through the effective repeat length along a rail. a,** Definition of the effective repeat length along a rail, *Z*, which is determined by the combined length of a beam segment and the intervening bKL interaction between adjacent dsRNA bricks. The relative helical phase associated with *Z* determines the relative orientation of neighboring bricks. **b-d,** Predicted relative orientations of adjacent bricks for effective repeat lengths of 16 bp (**b**), 18 bp (**c**) and 19 bp (**d**). These repeat lengths correspond to relative helical rotations of approximately 180°, 240° and 270°, respectively, equivalent to inter-brick dihedral angles of approximately 180°, 120° and 90°. The 18-bp and 19-bp repeat lengths were used for the hexagonal tube **HT7B** and square tube **QT5B**, respectively.

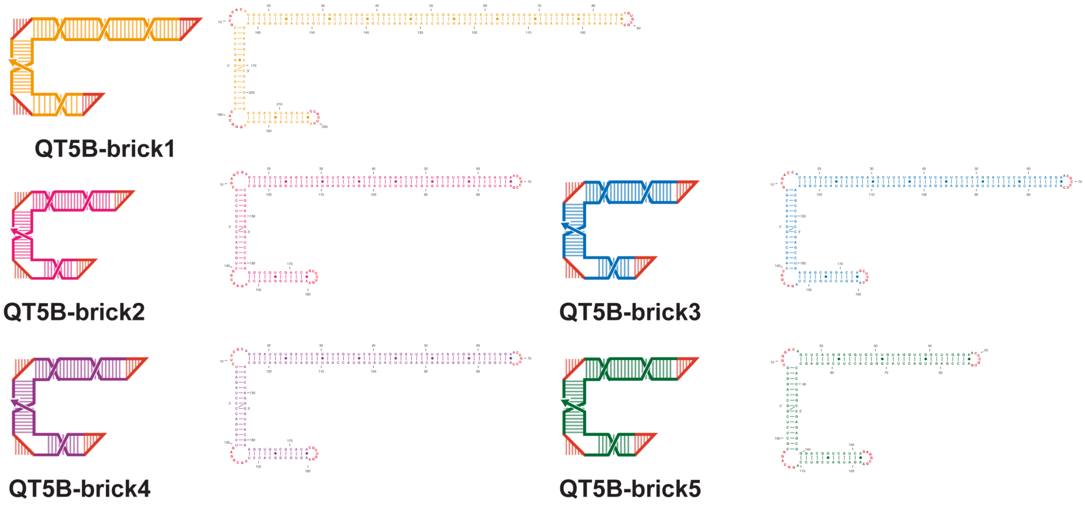

**Supplementary Fig. 14 Schematic of individual dsRNA bricks in QT5B.**

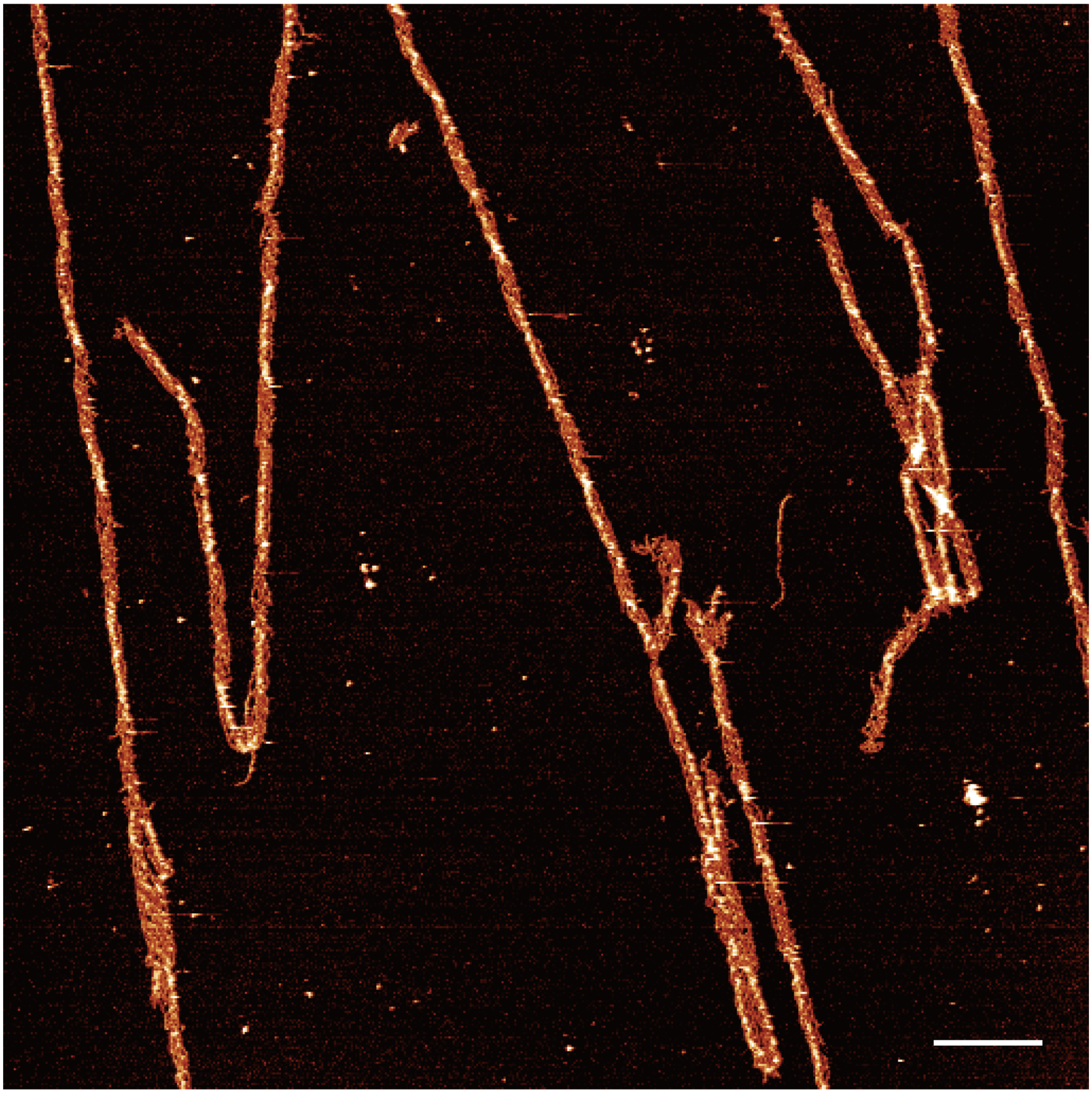

**Supplementary Fig. 15 Zoomed-out AFM image of QT5B.** Scale bar, 200 nm.

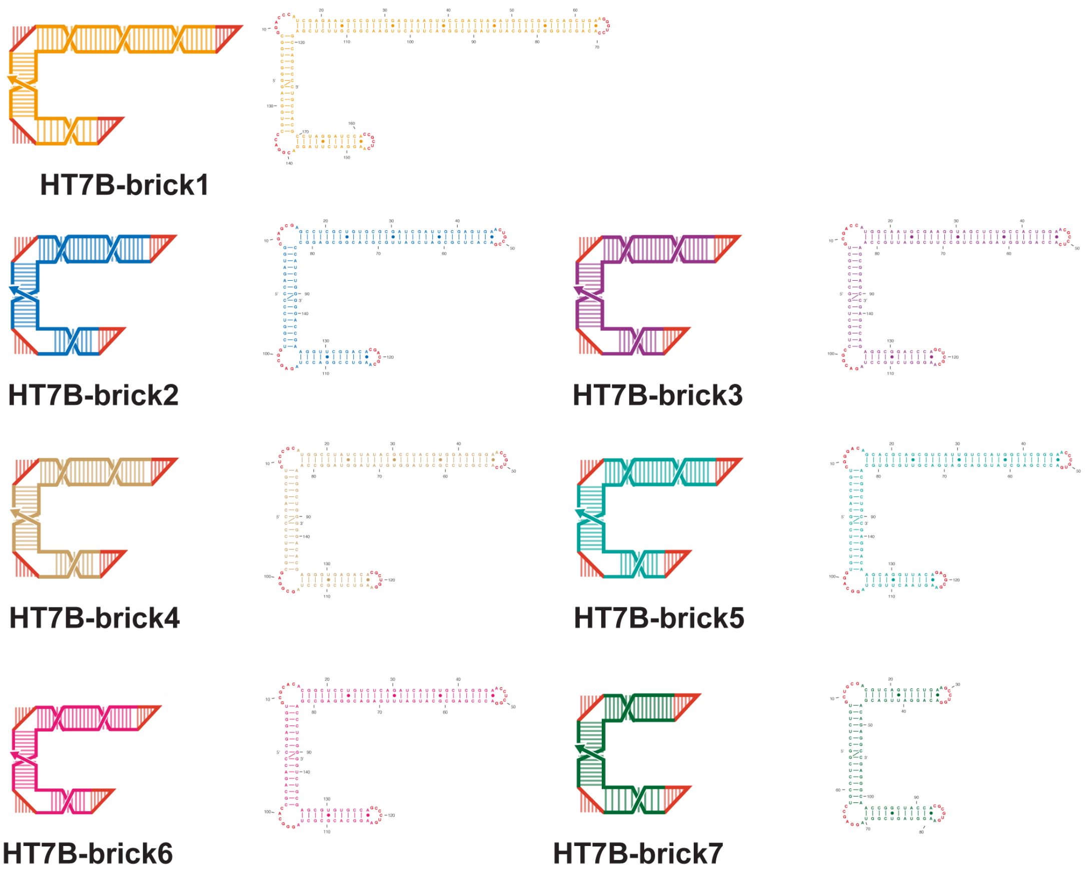

**Supplementary Fig. 16 Schematic of individual dsRNA bricks in HT7B.**

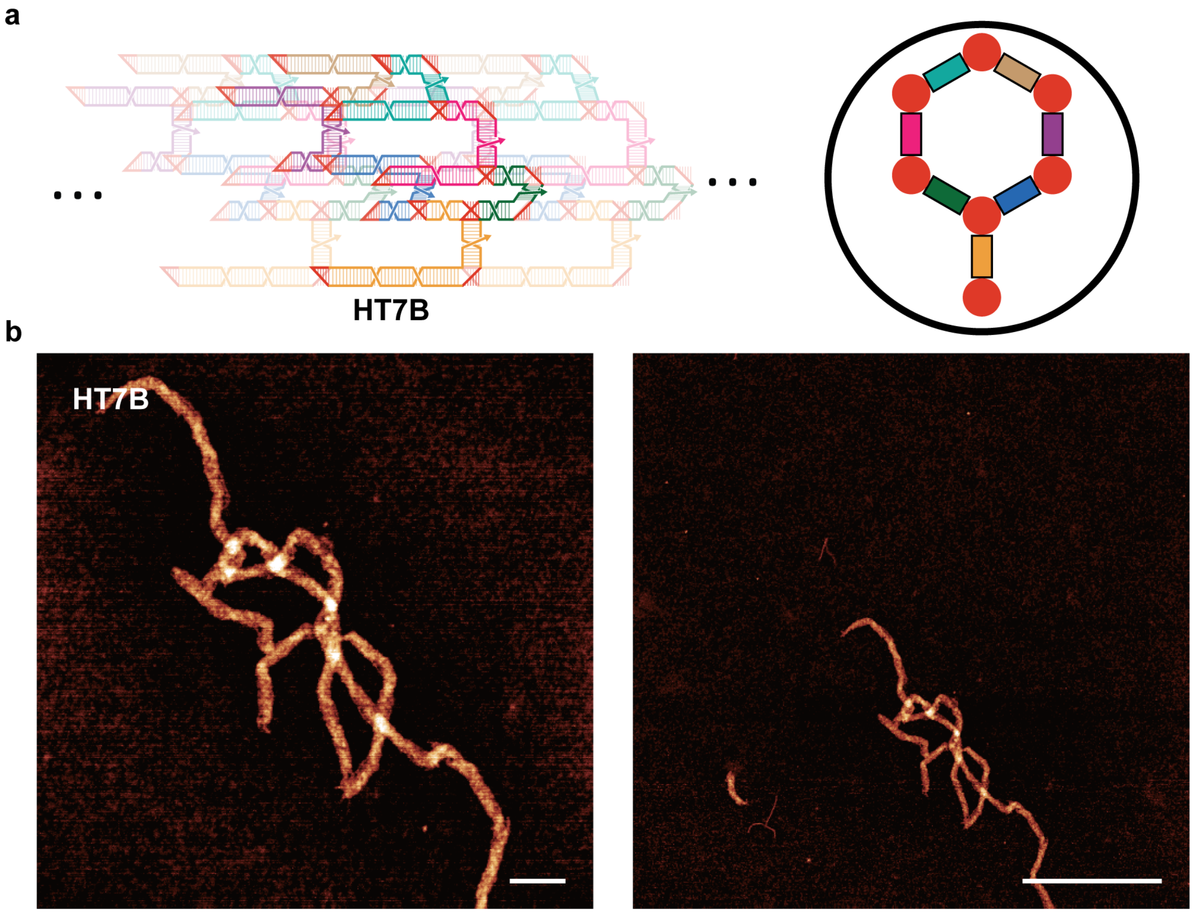

**Supplementary Fig. 17 Design and AFM images of HT7B. a,** Schematics of the formation of hexagonal tubes with HT7B dsRNA bricks. **b,** Zoomed-in (left, scale bar, 100 nm) and zoomed-out (right, scale bar, 500 nm) AFM images of HT7B.

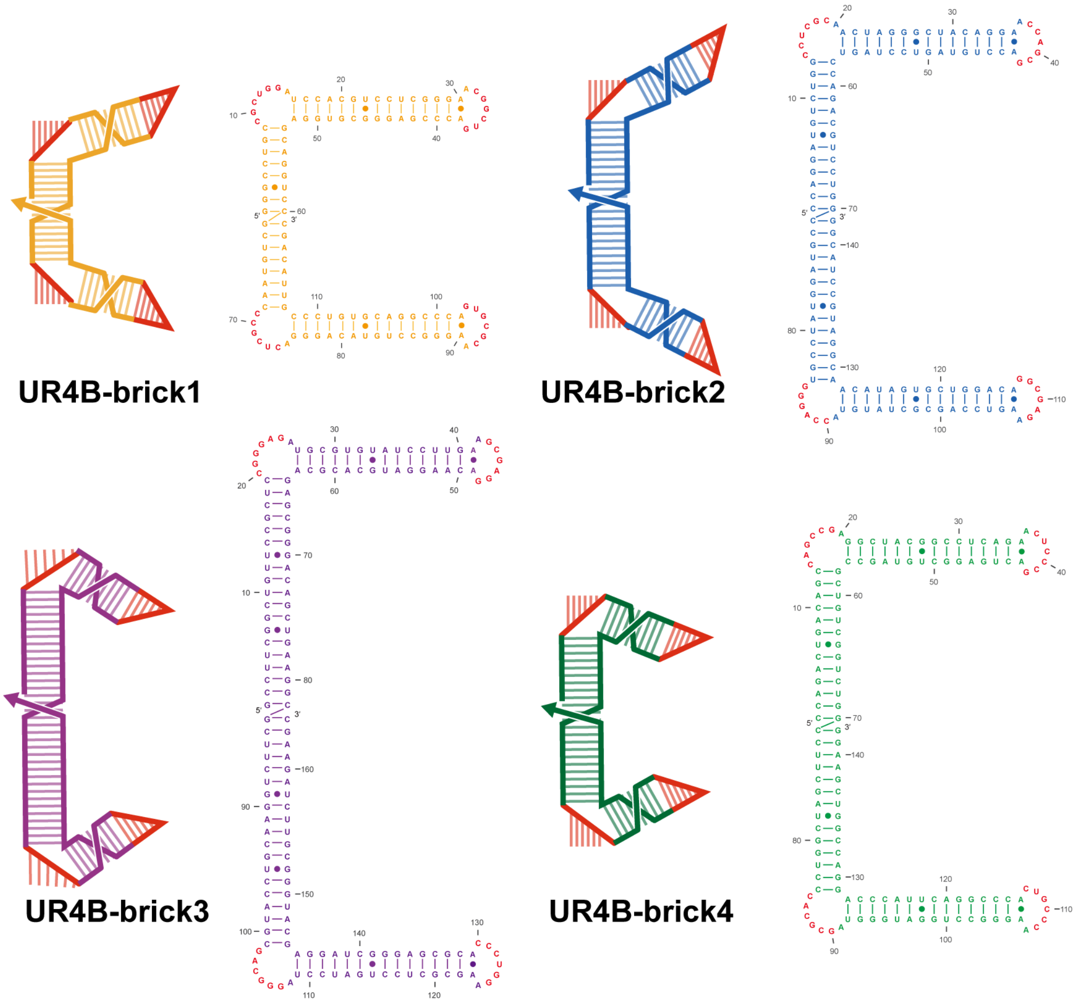

**Supplementary Fig. 18 Schematic of individual dsRNA bricks in UR4B.**

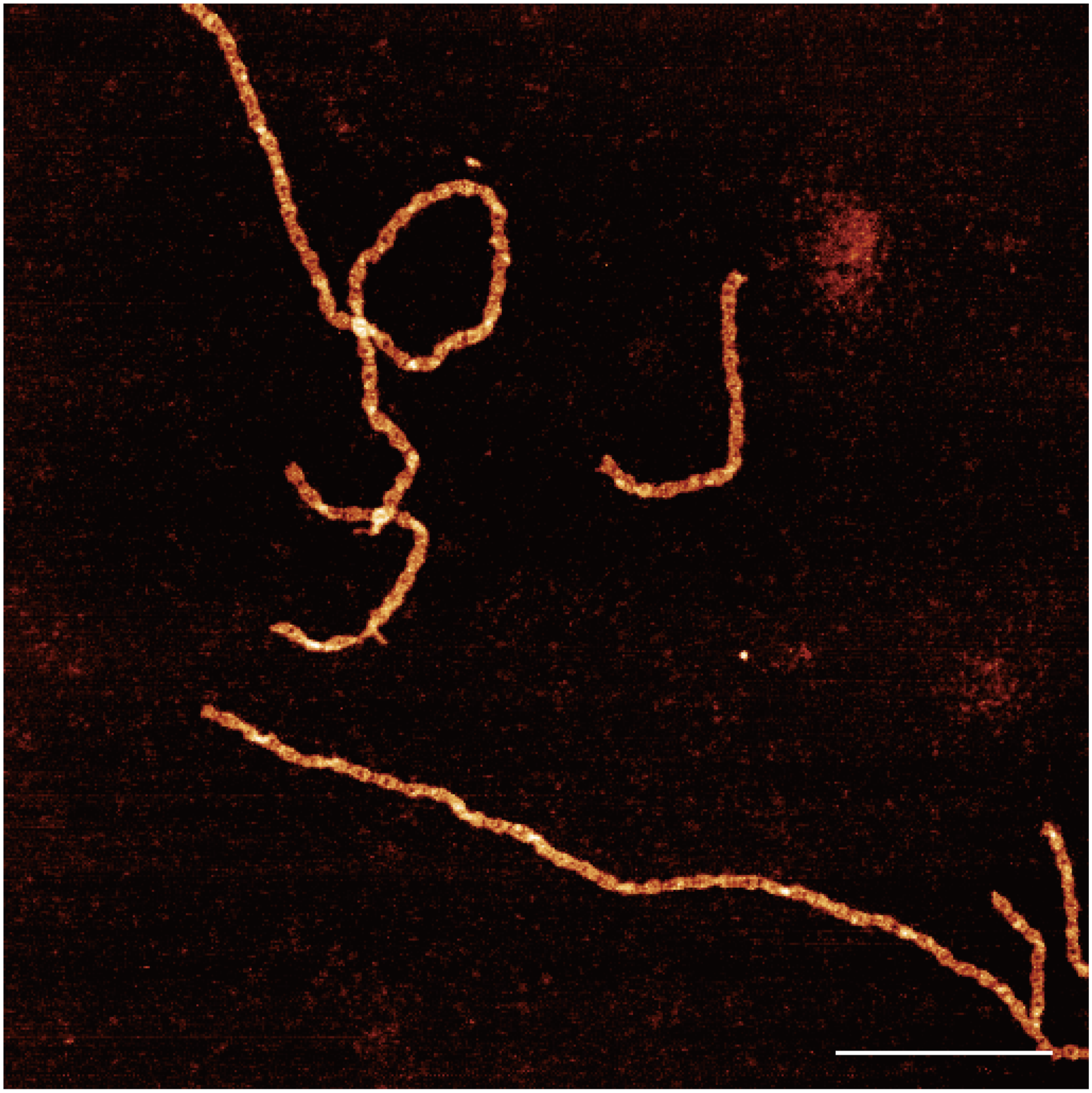

**Supplementary Fig. 19 Zoomed-out AFM image of UR4B.** Scale bar, 200 nm.

**a**

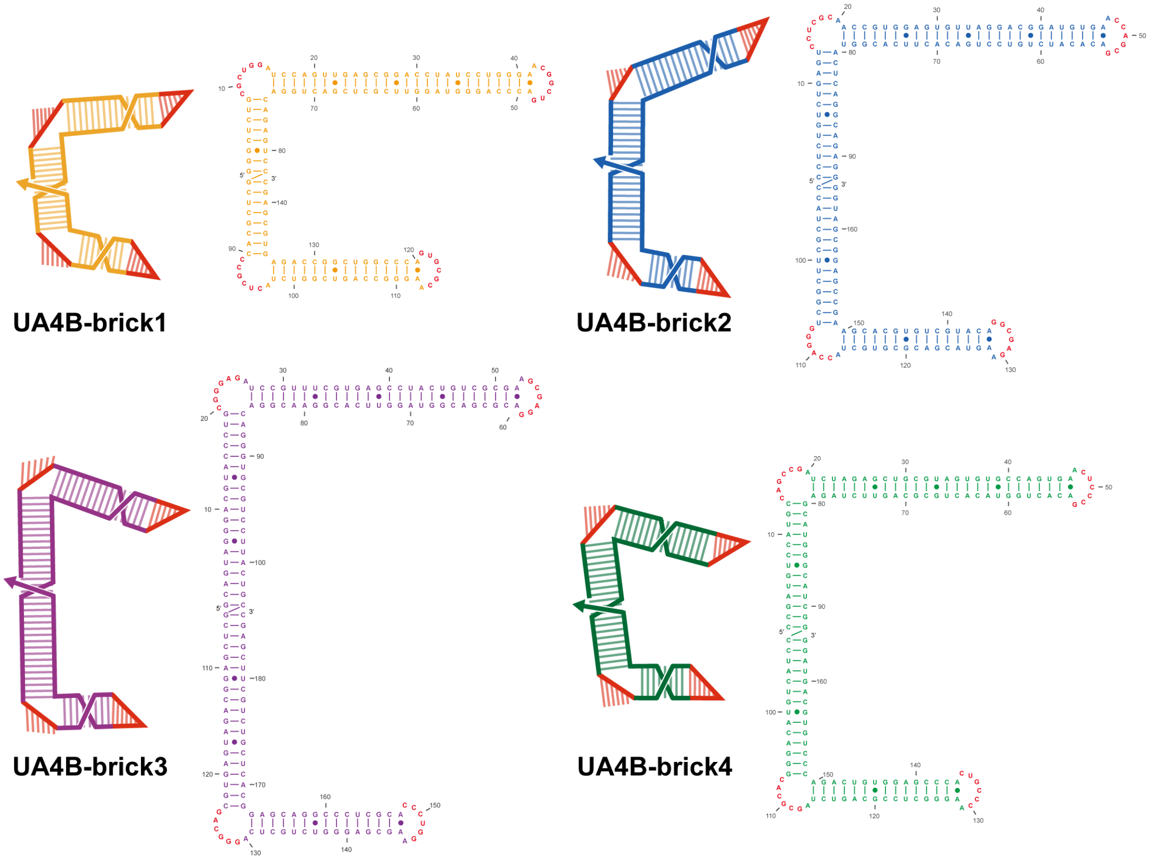

**b**

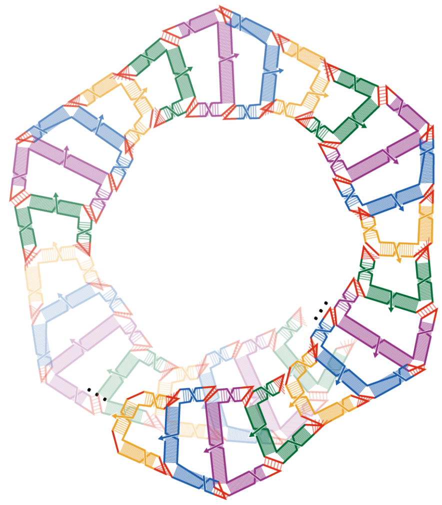

**Supplementary Fig. 20 Schematic of self-assembly and sequence of individual dsRNA bricks in UA4B. a,** The sequences and secondary structures of UA4B bricks. The length of struts and beams in different bricks are optimized to create curvature in the self-assembly. **b,** Schematics of the formation of undulated ladder with UA4B dsRNA bricks.

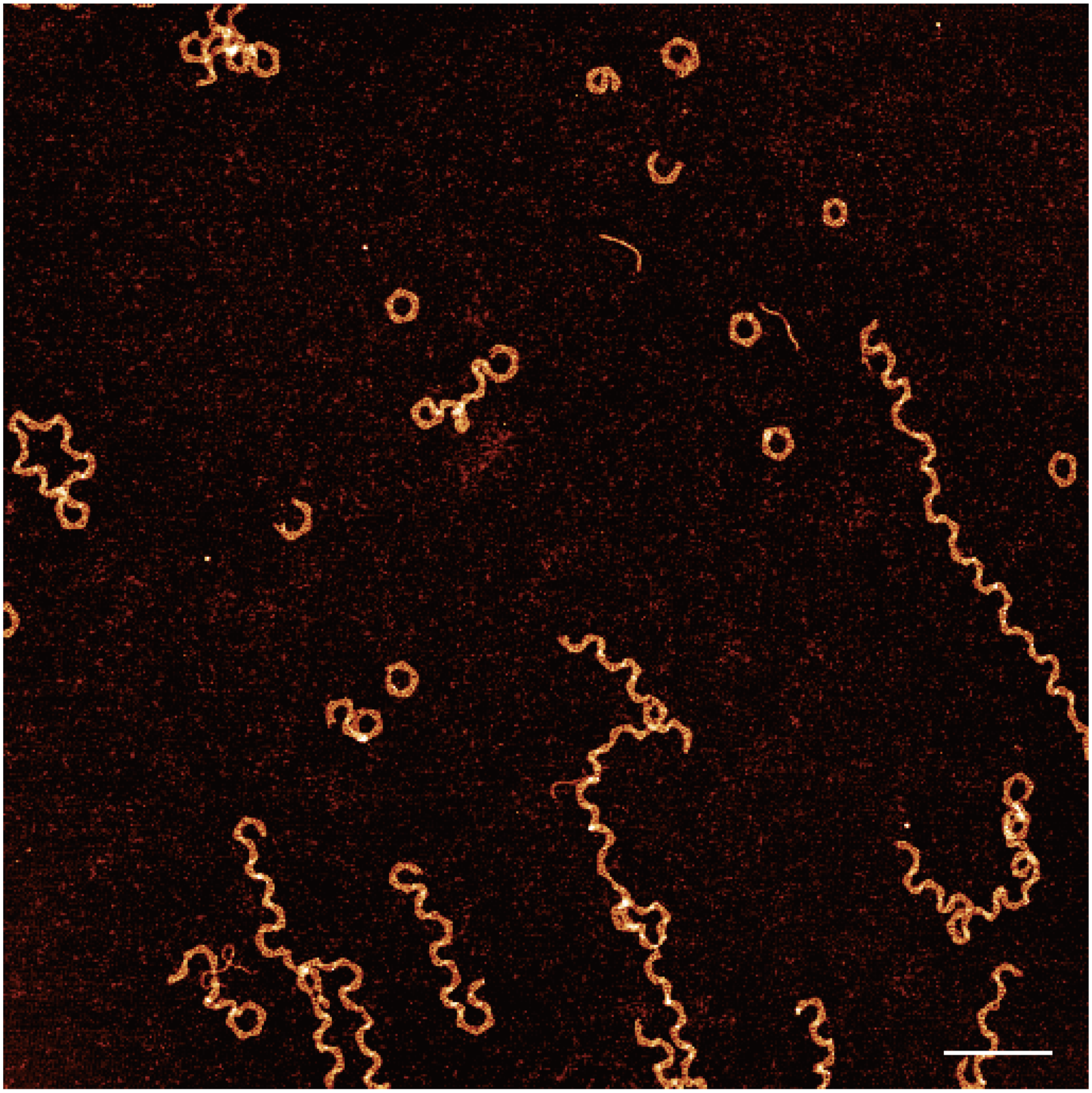

**Supplementary Fig. 21 Zoomed-out AFM image of UA4B.** In the AFM, we can see two types of self-assembly products. One type is the pentagonal or quadrilateral annulus with 20 or 16 bricks respectively. The other type is undulated ladder structures, which indicates the existence of curvature in the self-assembly and avoids the formation of closed structures. Scale bar, 200 nm.

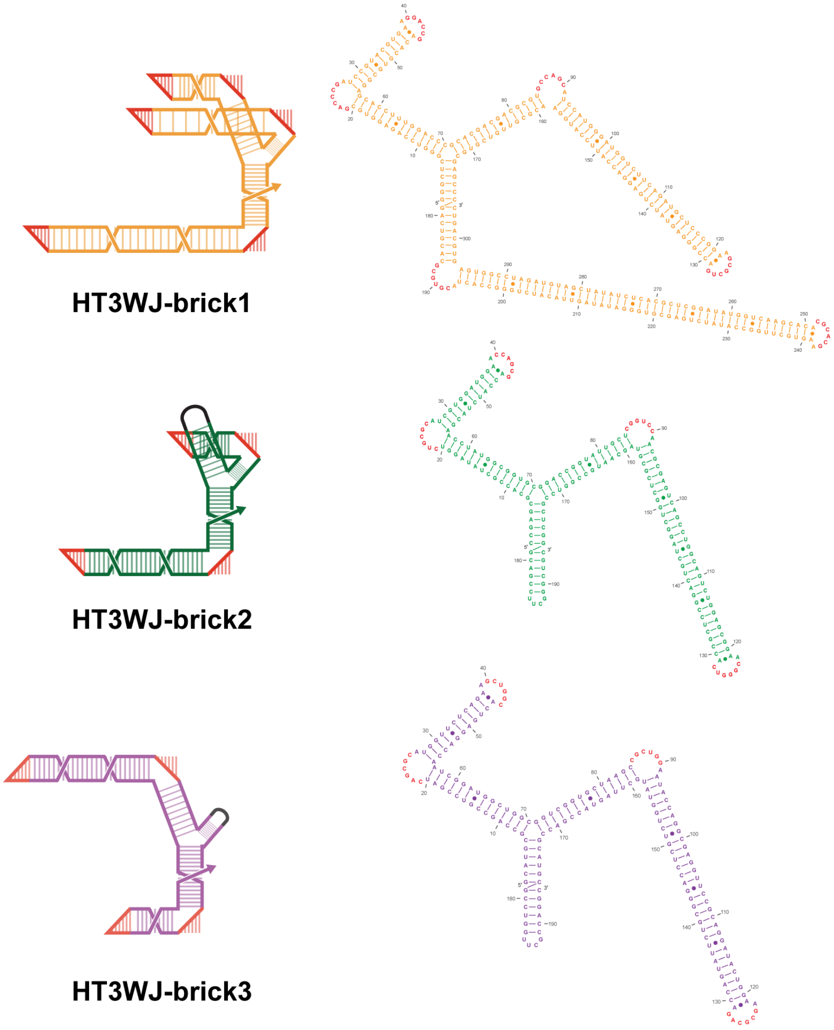

**Supplementary Fig. 22 Schematic of individual bricks in HT3WJ.** HT3WJ-brick1 contains an elongated third beam that serves as nucleation site, whereas the other two bricks terminate in tetraloops. The length of repeating unit along x-axis remains 55-bp to achieve a straight rail.

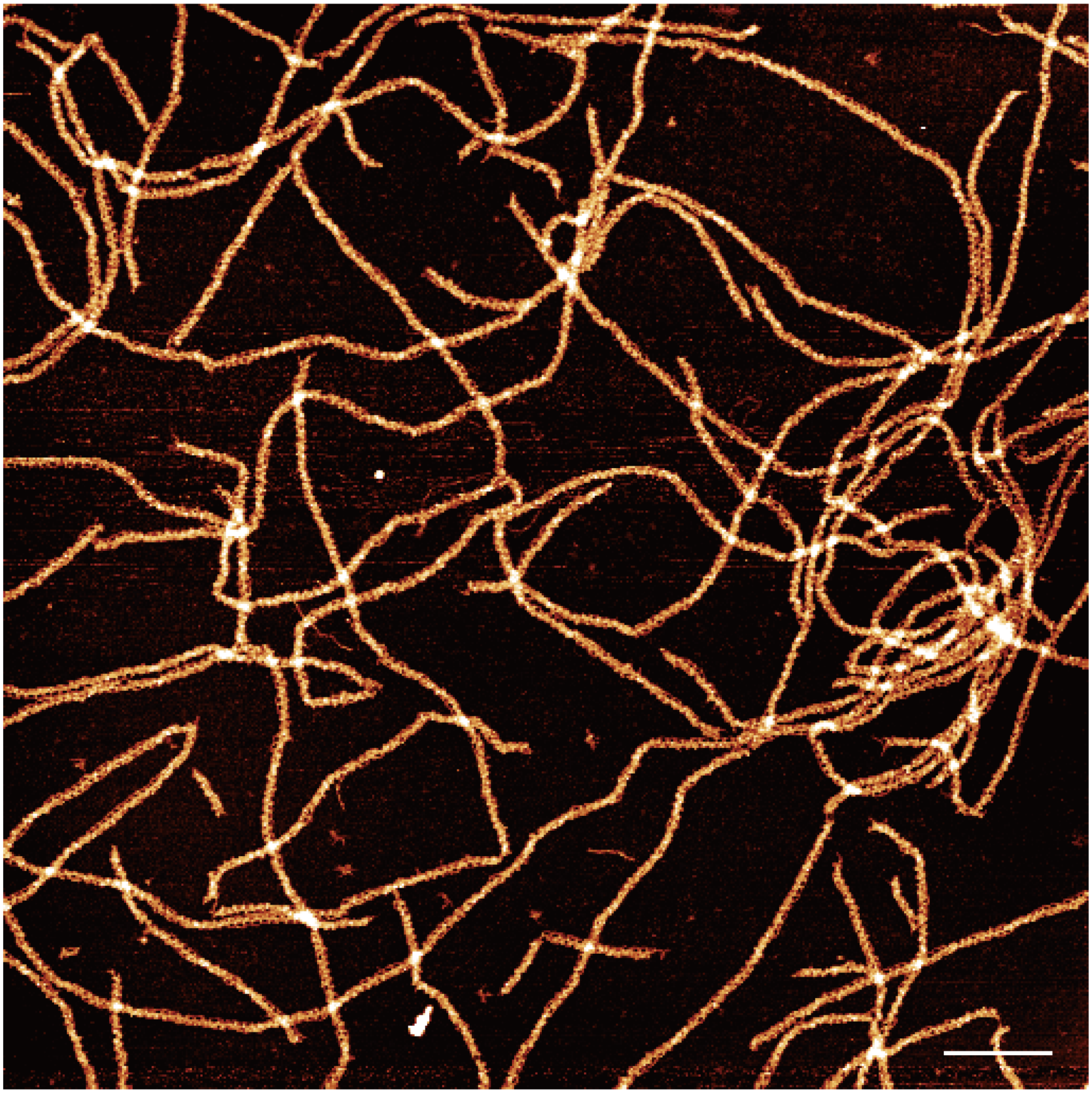

**Supplementary Fig. 23 Zoomed-out AFM image of HT3WJ.** Scale bar, 200 nm.

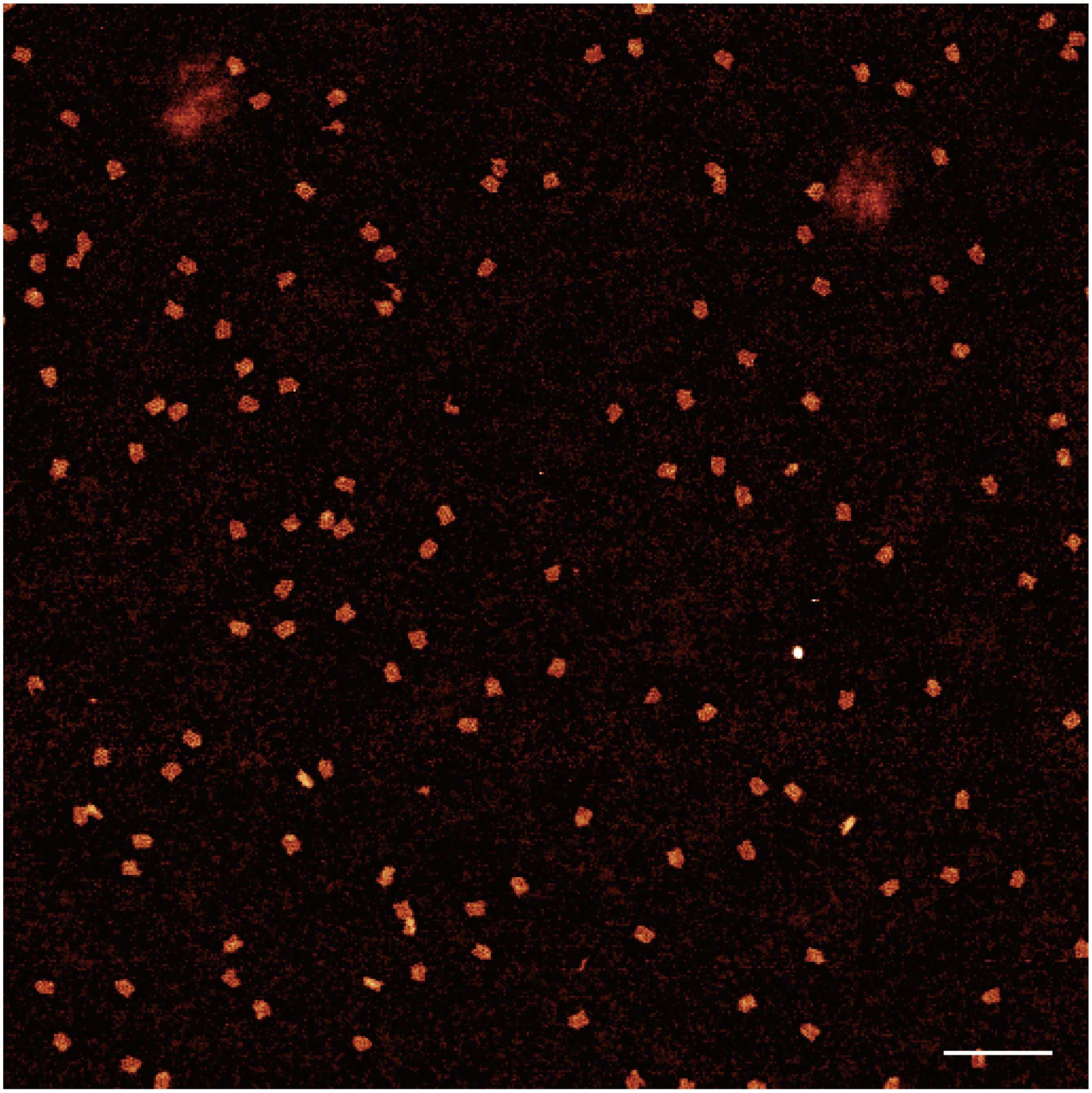

**Supplementary Fig. 24 Zoomed-out AFM image of F15S.** Scale bar, 200 nm.

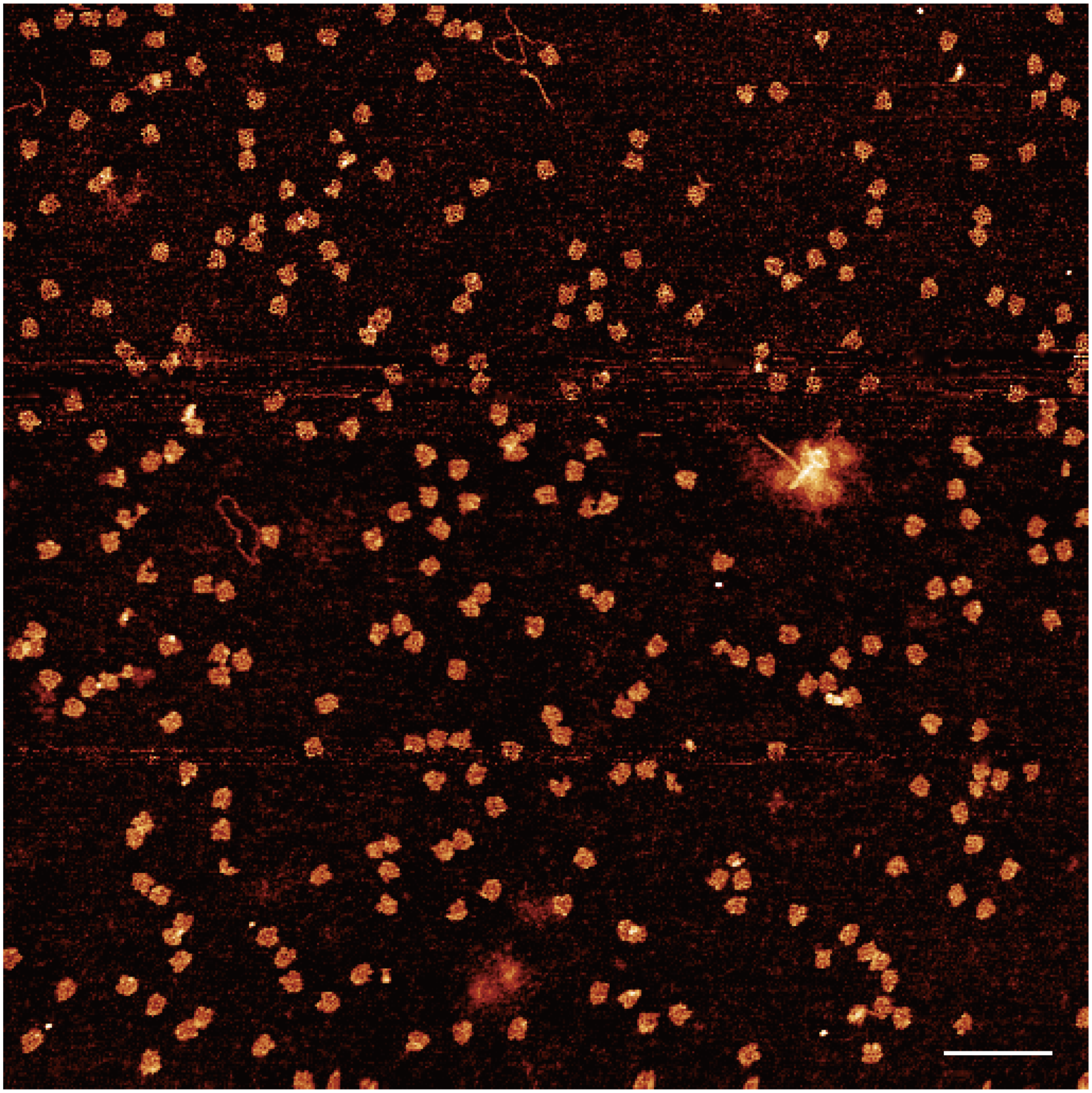

**Supplementary Fig. 25 Zoomed-out AFM image of F15H.** Scale bar, 200 nm.

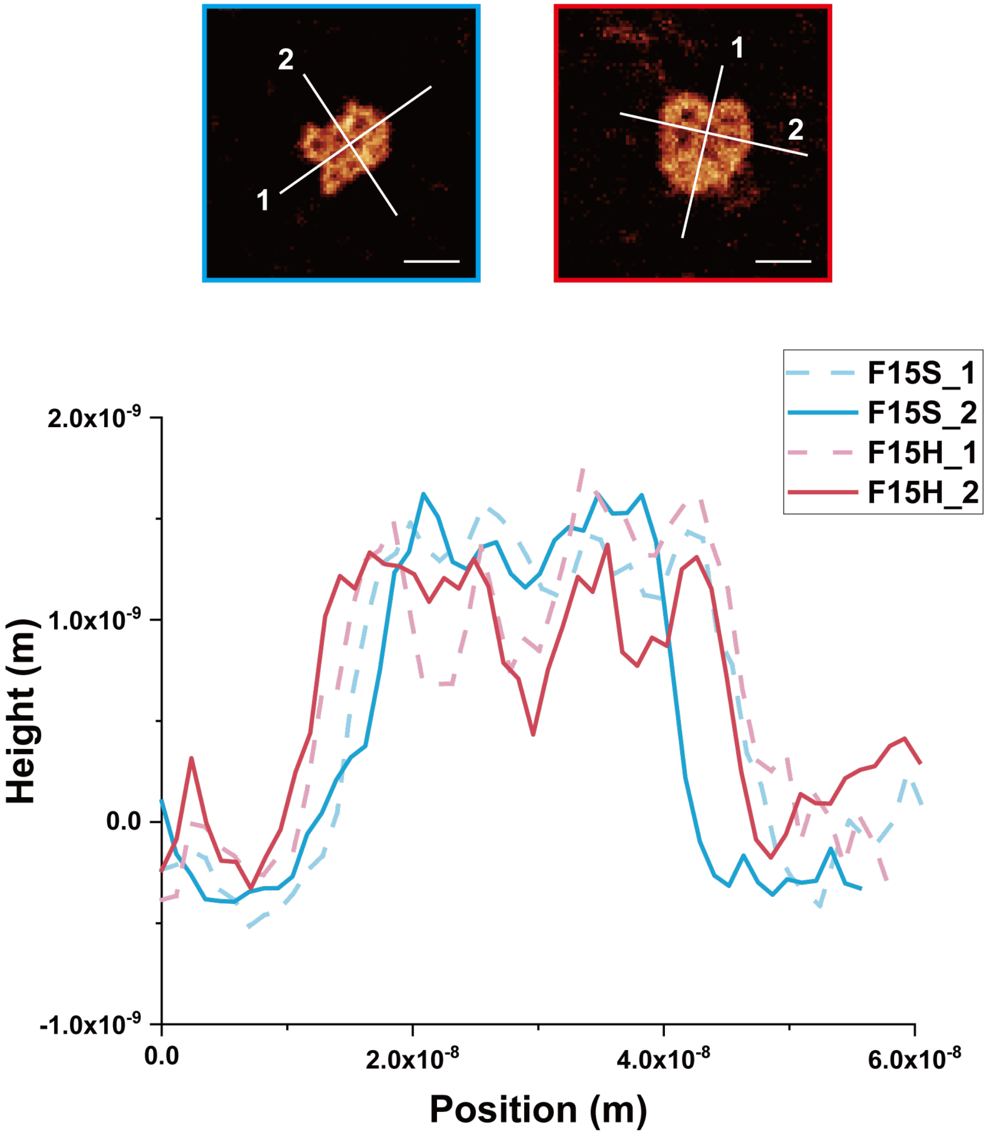

**Supplementary Fig. 26 Height measurements of F15S and F15H by AFM.** F15S and F15H have similar lengths, but the width of F15H is larger than that of F15S, which is corresponding to the elongated beams and struts of F15H. Scale bars, 20 nm.

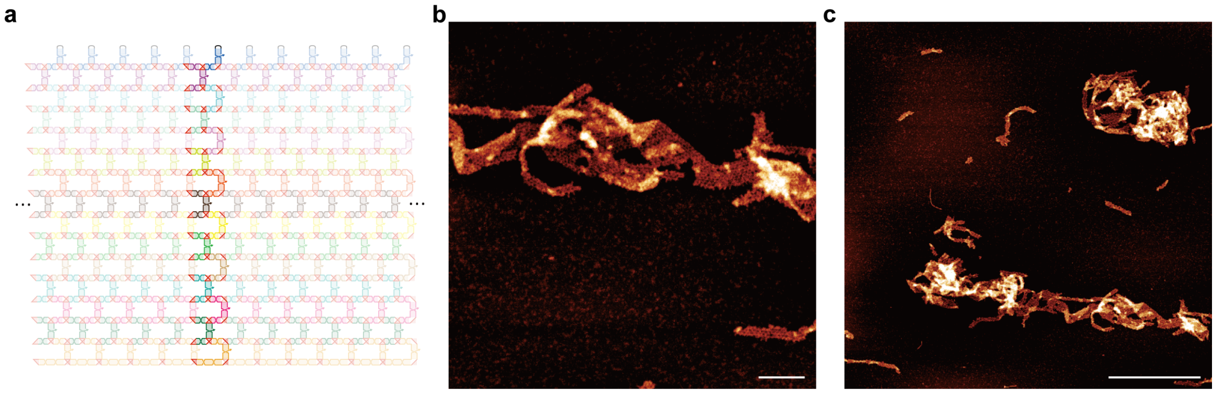

**Supplementary Fig. 27 Design of R15B with more bKL interactions. a,** Schematics of design of **R15B**. **b,** Zoomed-in AFM image of **R15B** assembly products. Scale bar, 100 nm. **c,** Zoomed-out AFM image of **R15B** assembly products. Scale bar, 500 nm. Although partially ribbon-like structures were observed, complete **R15B** ribbons were rare, suggesting limited assembly fidelity. This limitation may arise from the restricted orthogonal sequence space of 6-bp bKLs and potential crosstalk among bKL interactions.

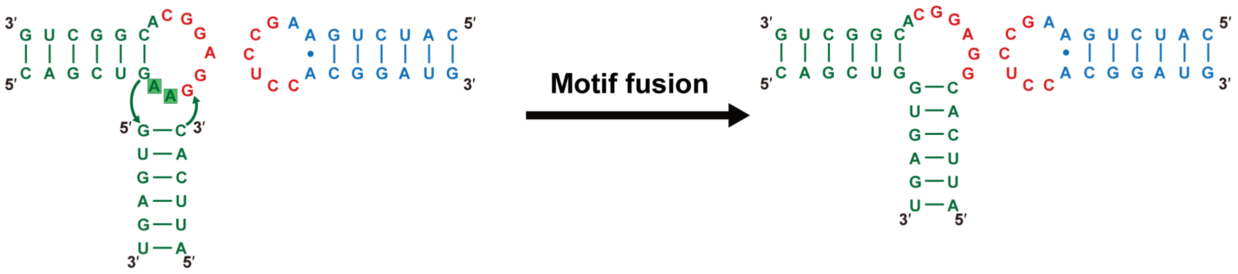

**Supplementary Fig. 28 Design of bKL interaction by motif fusion.** The sequence details of the motif fusion process used to design bKL from a KL and an RNA helix.

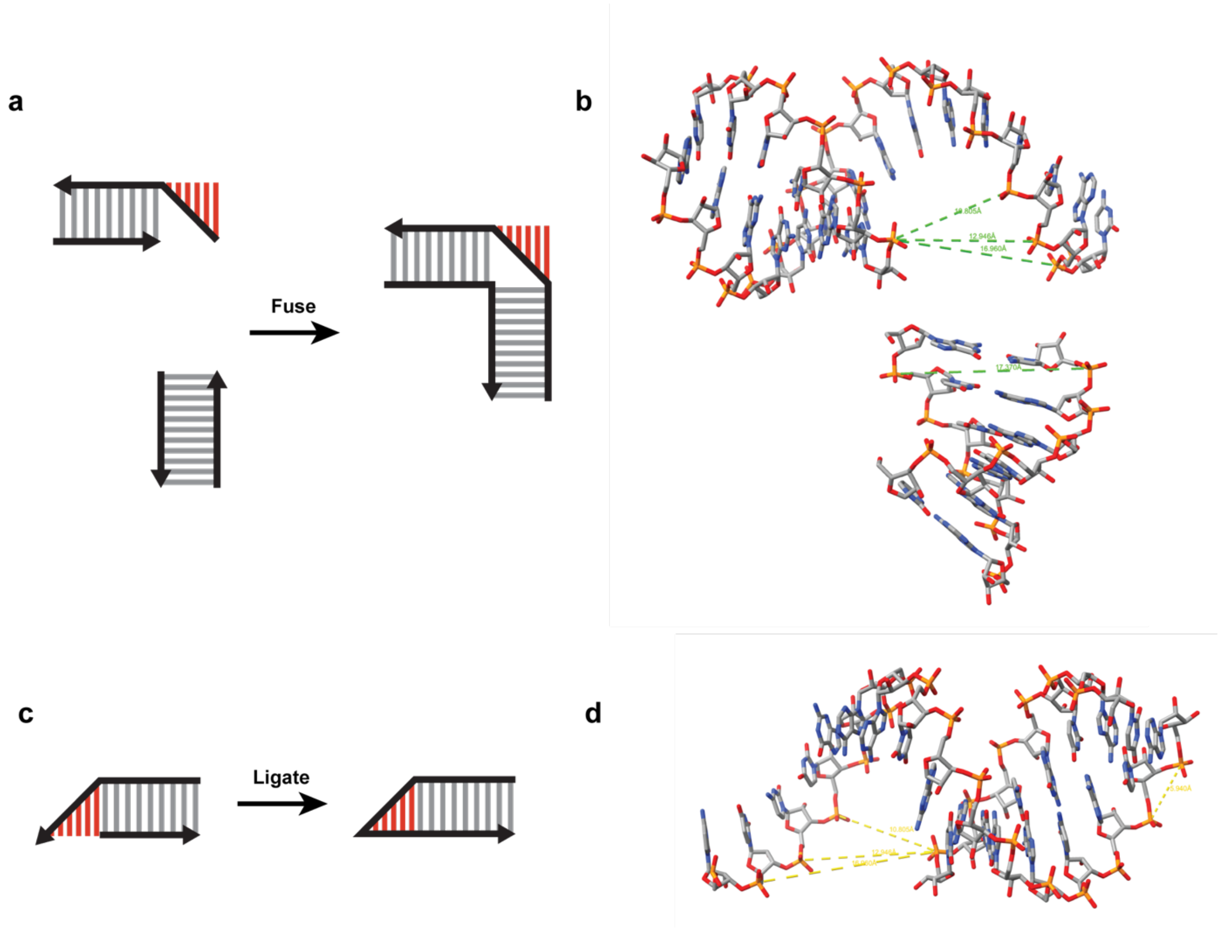

**Supplementary Fig. 29 Design of bKL interaction with longer kissing helix. a,** Schematics design of the bulge side of the bKL. One of the helices in x-axis has the extension on the 3' end for the kissing helix and fuses with the other perpendicular helix to form the loop. **b,** Model of RNA helix shows that the distance between two ends of 7-bp, 8-bp and 9-bp extension is 10.8 Å, 12.9 Å, and 17.0 Å respectively. The distance between two paired nucleotides in the perpendicular helix is 17.4 Å, which implies 9-bp design needs an A·A pair to increase the flexibility. **c,** The schematics of design of loop in the bKLs. The helix in x-axis has the extension on the 3' end for the kissing helix and ligates with the 5' end of the complementary strand with a linker. **d,** Model of RNA helix shows the distance between two ends of 7-bp, 8-bp and 9-bp extension is 10.8 Å, 12.9 Å, and 17.0 Å, respectively. The spacer between two nucleotides is 5.9 Å. Thus, 7-bp, 8-bp and 9-bp bKLs need 2, 2, 3-nt polyA spacer respectively.

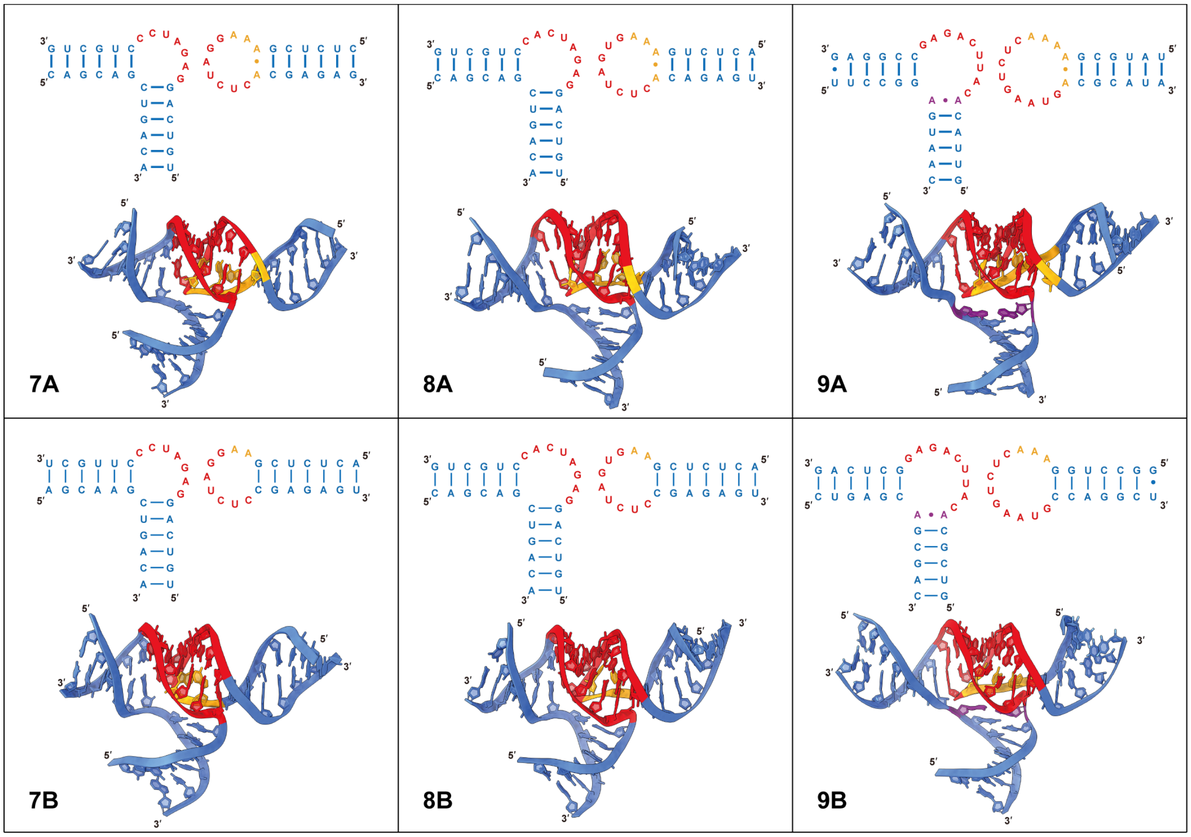

**Supplementary Fig. 30 Sequences and models of different bKLs.** In type A designs, there is an A·A pair as a linker between loop and stem. In type B designs, this A·A pair is absent and the loop is directly connected to the stem. Meanwhile, the 9-bp bKL, an extra A·A pair was introduced between branched helix and the bulge to provide additional flexibility. The structures of different bKLs were simulated by SimRNA[15,16]. All the six bKLs shows coaxial stacking of base pairs within stems and kissing helix.

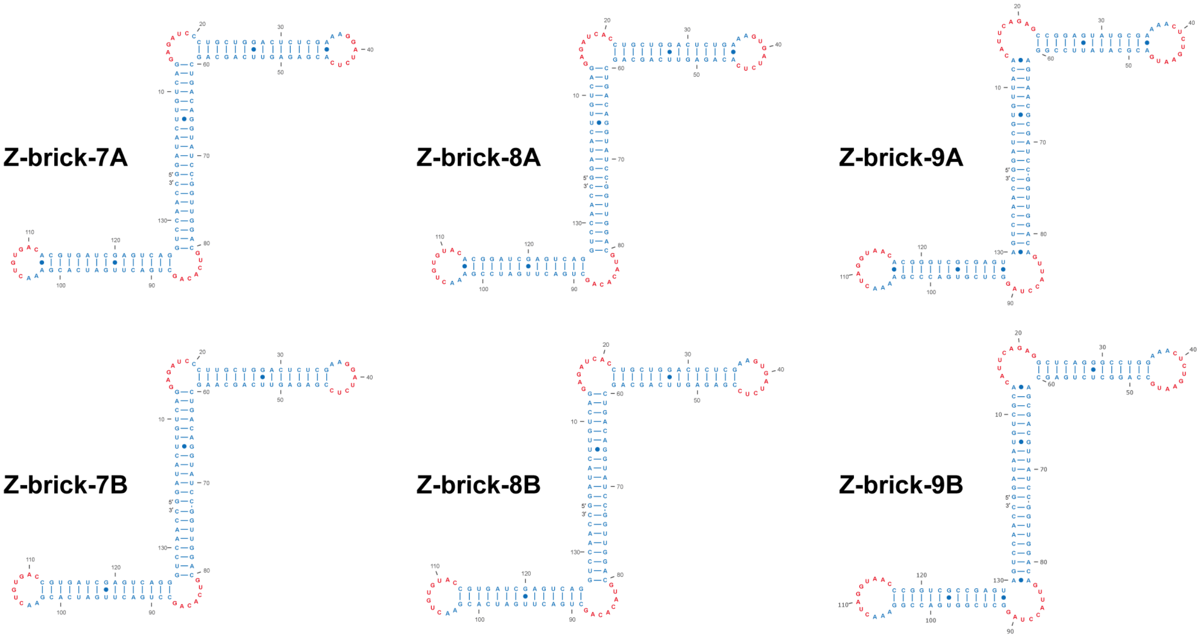

**Supplementary Fig. 31 Sequences of Z-brick with different types of bKLs.** All Z-bricks contain a 21-bp strut (corresponding approximately to two helical turns), which enforces an antiparallel arrangement such that the two beams extend in opposite directions. The beam length of each Z-brick is tuned so that the effective length of each repeating unit, defined as one beam segment plus one bKL, is fixed at 22 bp, thereby minimizing intrinsic curvature.

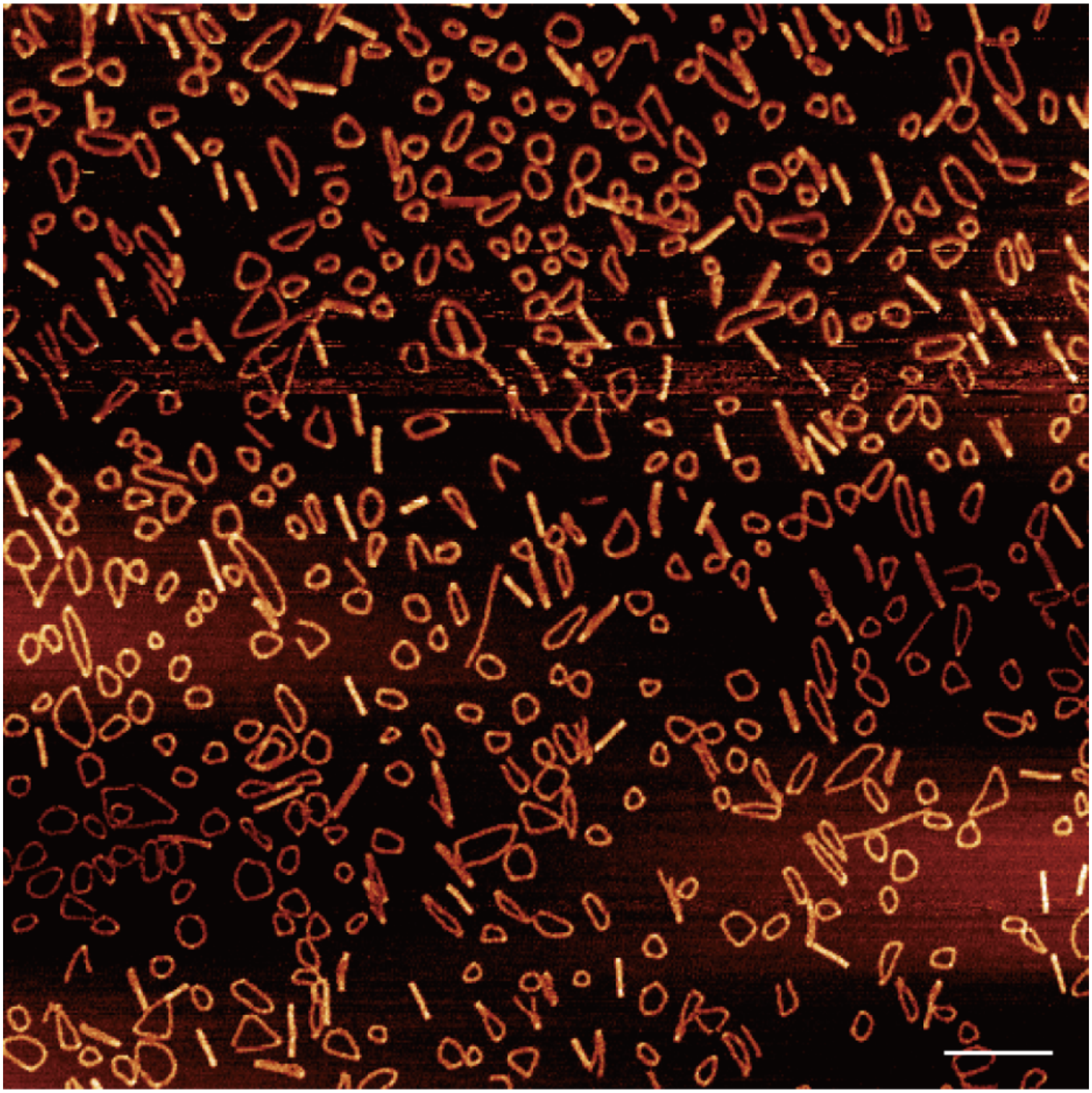

**Supplementary Fig. 32 Zoomed-out AFM image of Z-brick self-assembly with bKL-7As.** Scale bar, 200 nm.

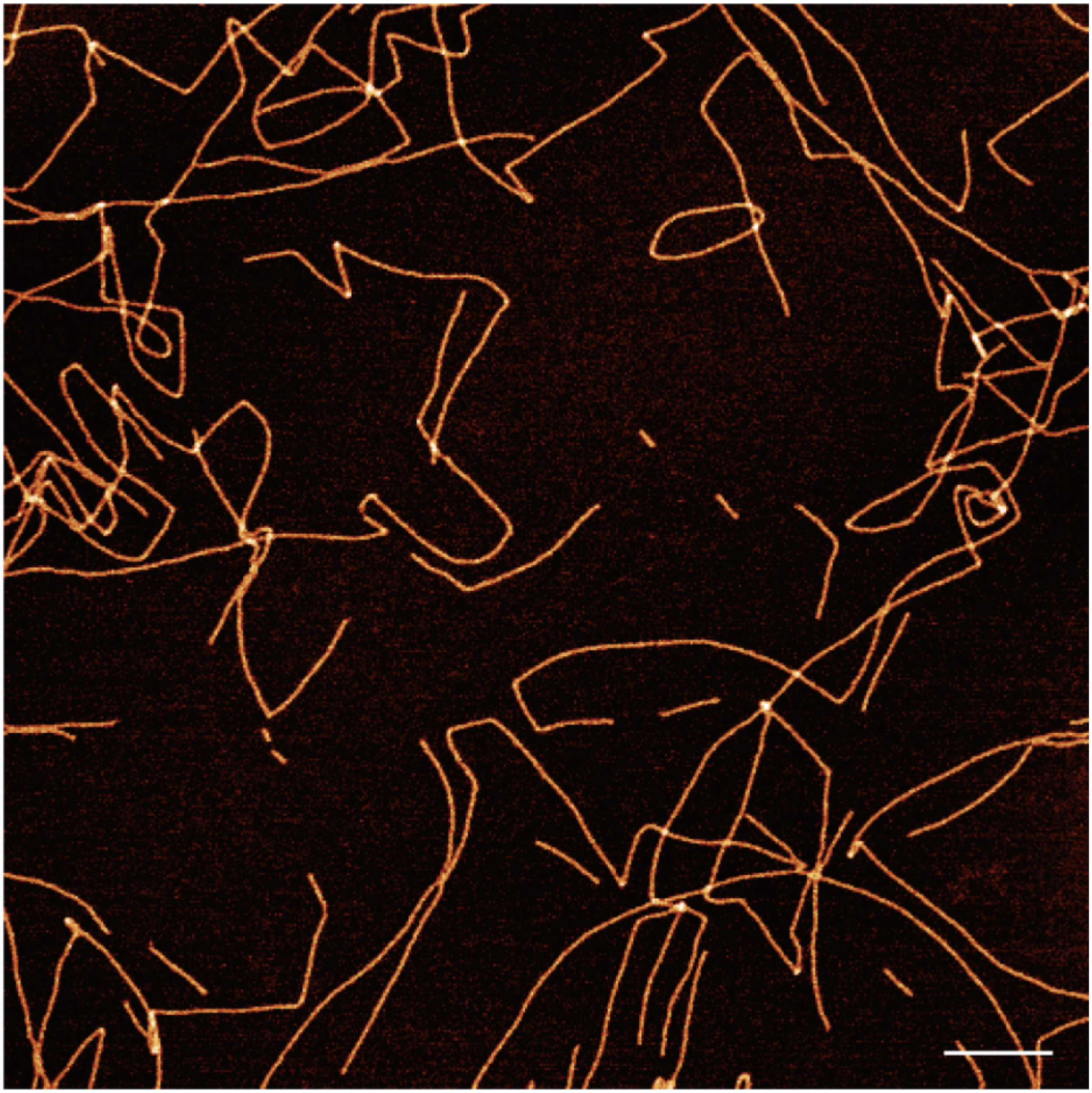

**Supplementary Fig. 33 Zoomed-out AFM image of Z-brick self-assembly with bKL-7Bs.** Scale bar, 200 nm.

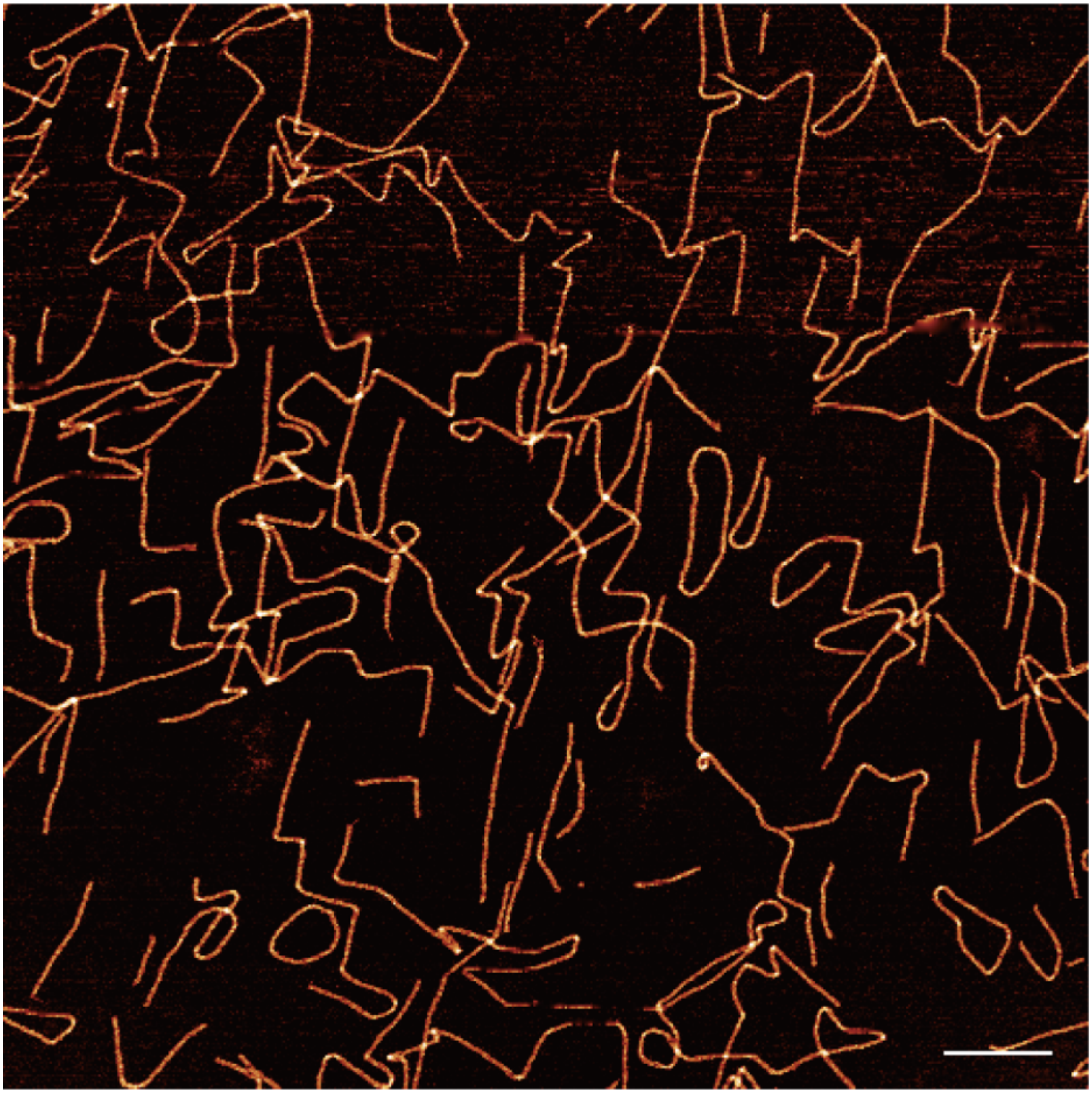

**Supplementary Fig. 34 Zoomed-out AFM image of Z-brick self-assembly with bKL-8As.** Scale bar, 200 nm.

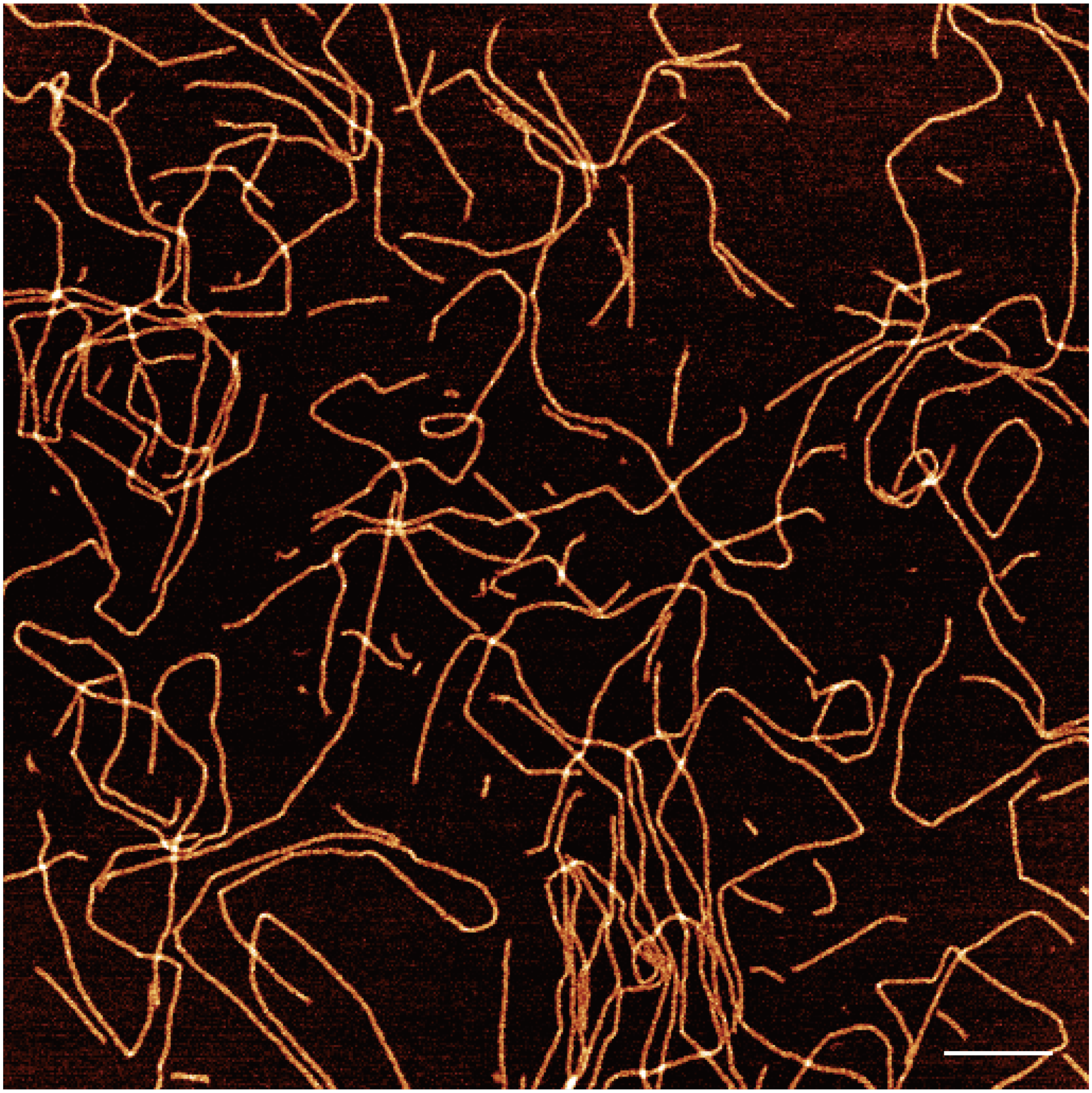

**Supplementary Fig. 35 Zoomed-out AFM image of Z-brick self-assembly with bKL-8Bs.** Scale bar, 200 nm.

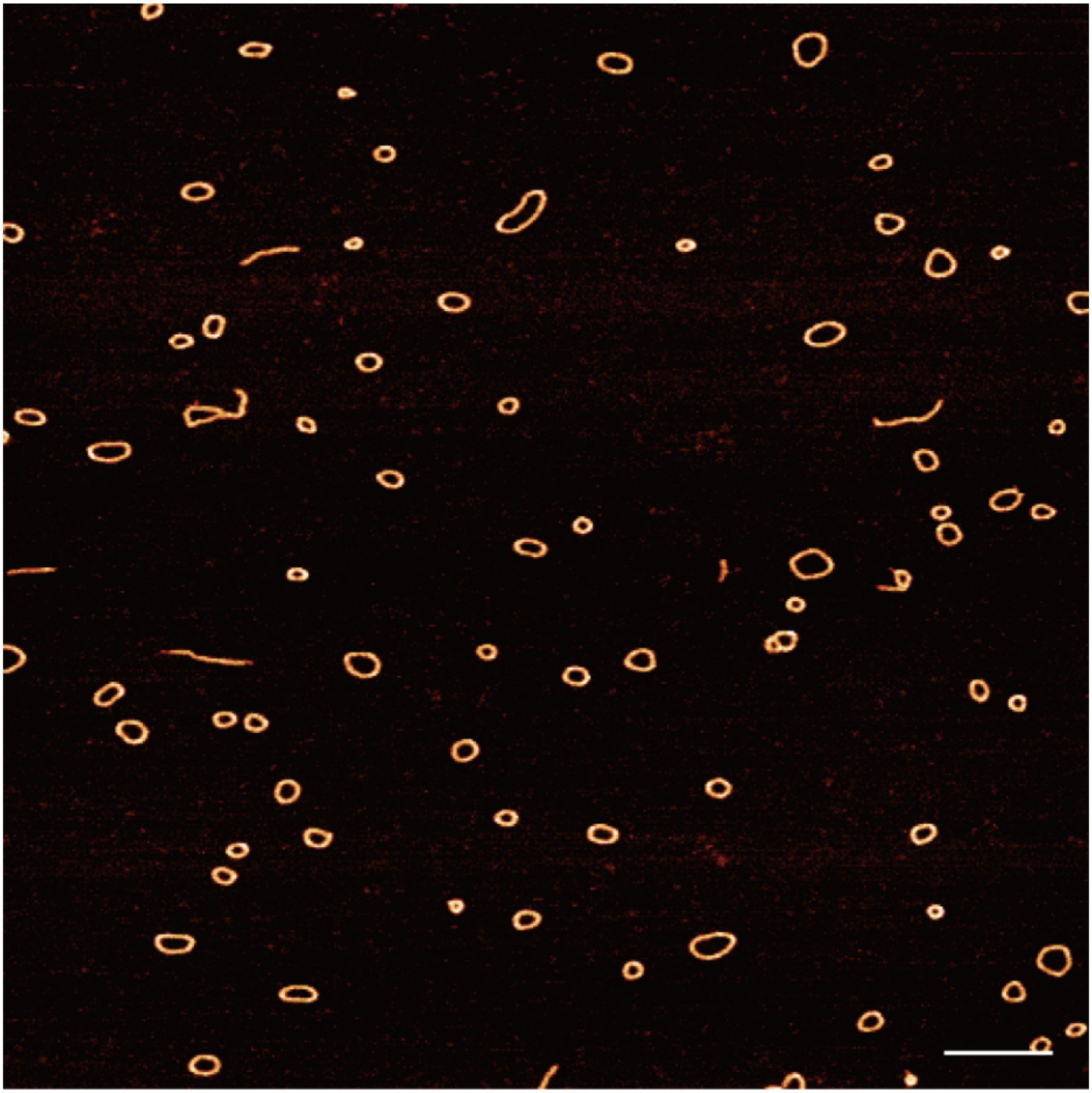

**Supplementary Fig. 36 Zoomed-out AFM image of Z-brick self-assembly with bKL-9As.** Scale bar, 200 nm.

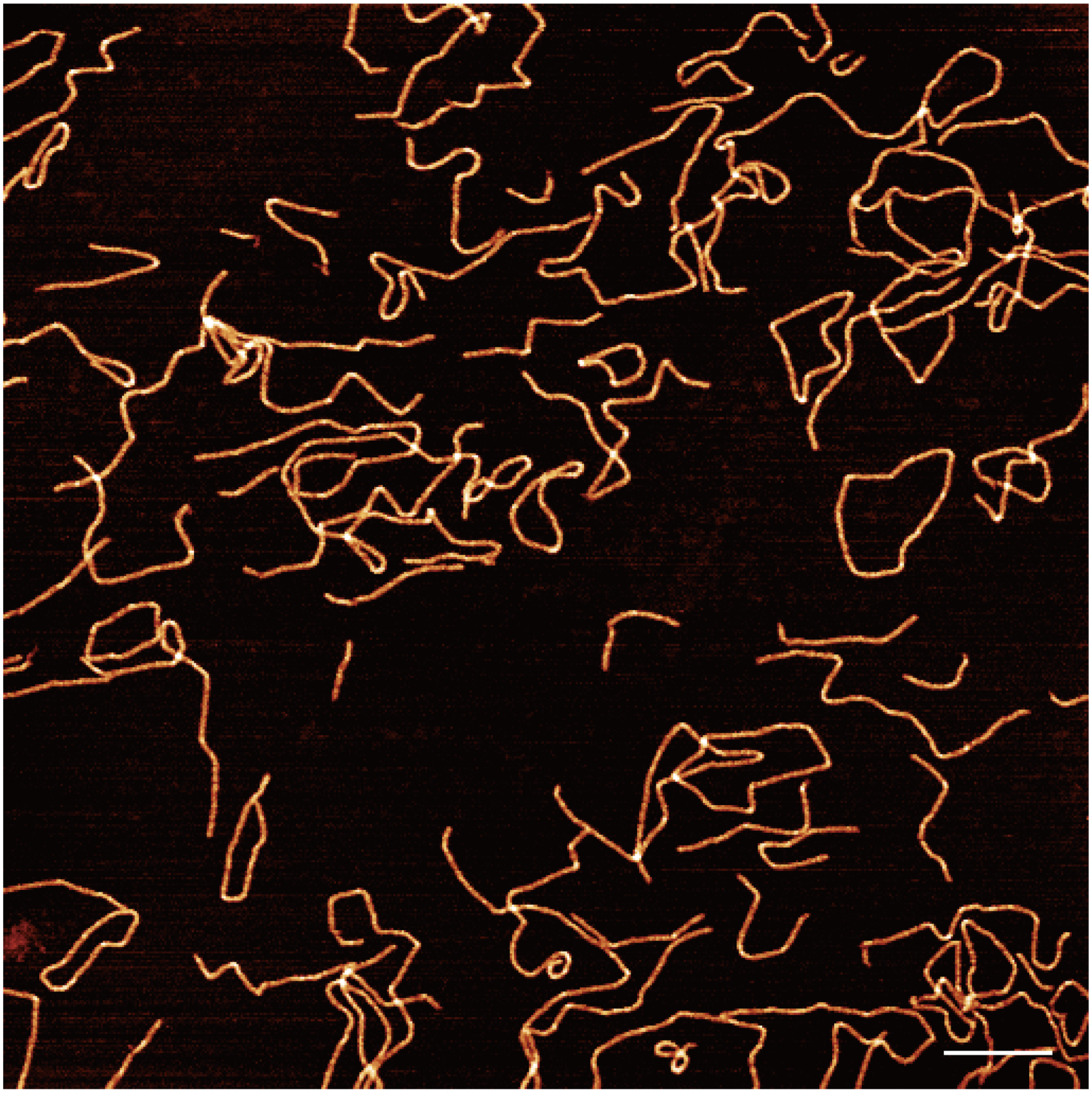

**Supplementary Fig. 37 Zoomed-out AFM image of Z-brick self-assembly with bKL-9Bs.** Scale bar, 200 nm.

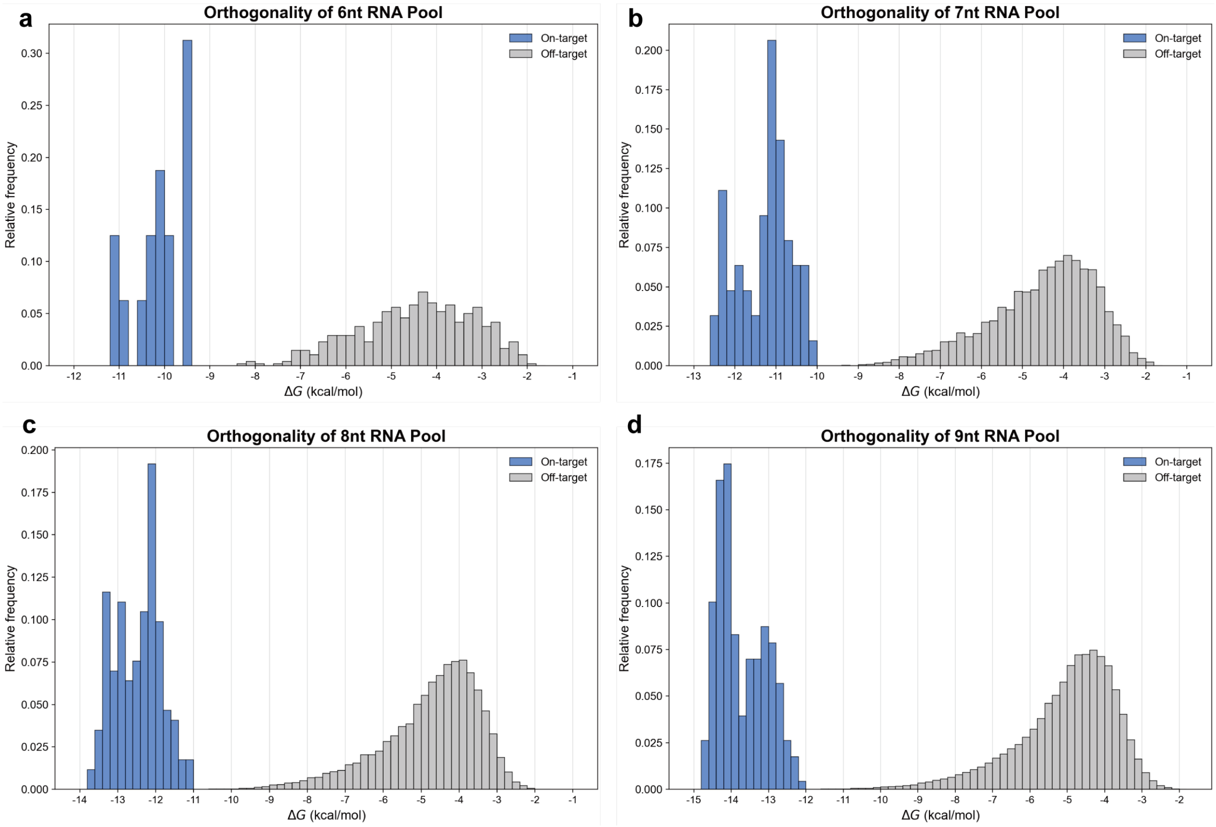

**Supplementary Fig. 38 Orthogonality analysis of selected bKL interaction pairs.** On-target energy indicates the ensemble free energy of paired bKL sequences. Off-target energy indicates the ensemble free energy of crosstalk among all the bKL sequences in the selected pool. The selected pools contain 16, 63, 172 and 229 mutually orthogonal bKL interaction pairs for the 6-, 7-, 8- and 9-bp designs, respectively. The ensemble free energy is calculated by NUPACK.

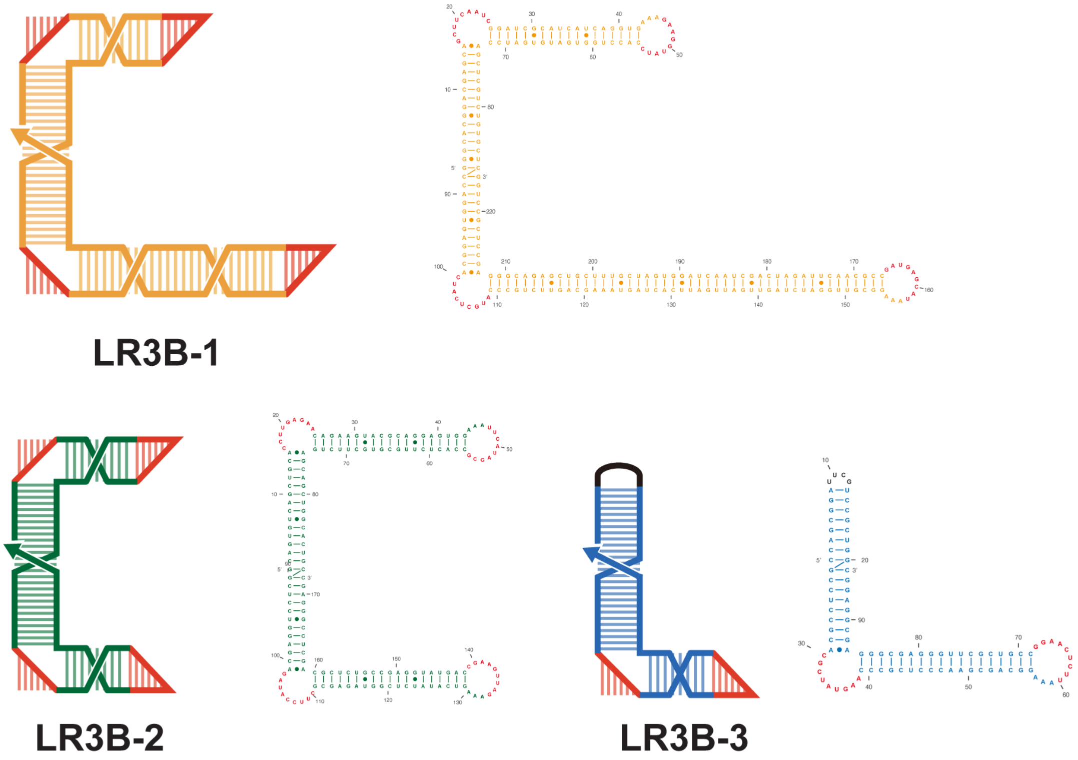

**Supplementary Fig. 39 Schematic of individual dsRNA bricks in LR3B.**

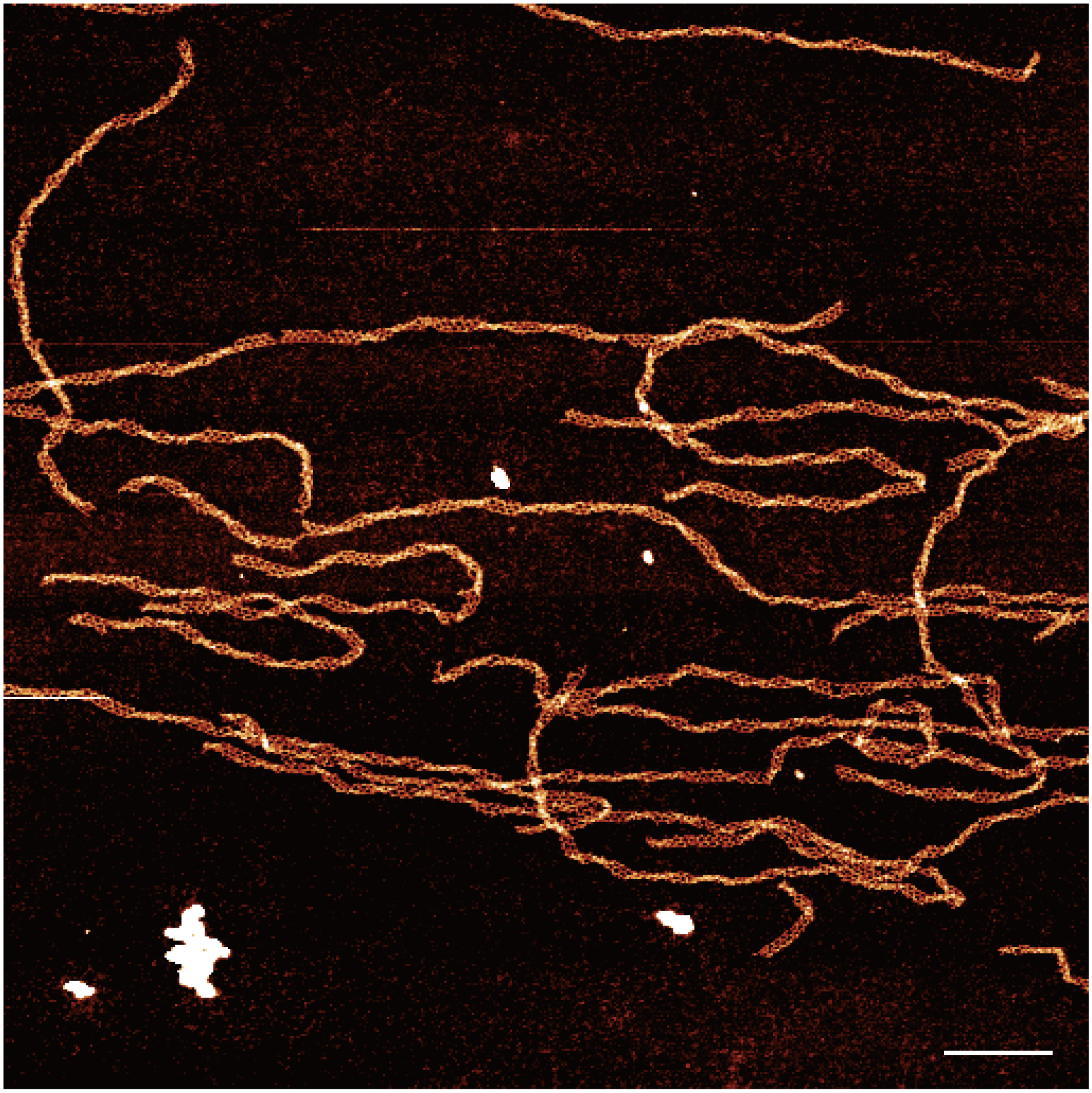

**Supplementary Fig. 40 Zoomed-out AFM image of LR3B.** Scale bar, 200 nm.

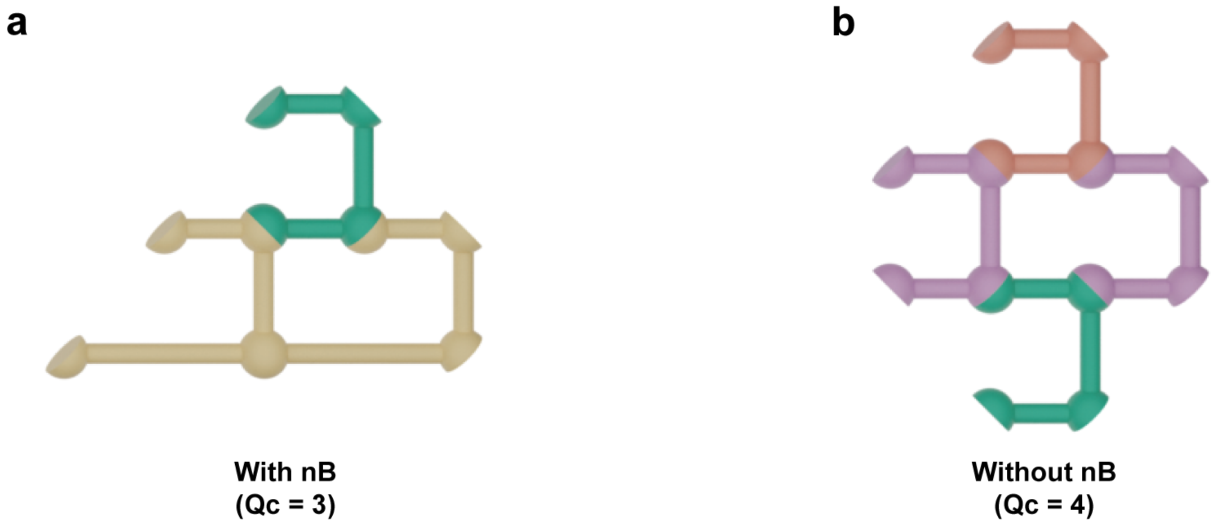

**Supplementary Fig. 41 Demonstration of nucleation with nBs in 2D_107. a**. The demonstration of nucleation with nBs. Only 3 bricks are needed to form a stable closed ring to initialize the growth in 2D. **b**. Without nBs, the 4 cBs are required to initialize the growth. Qc: coordination number.

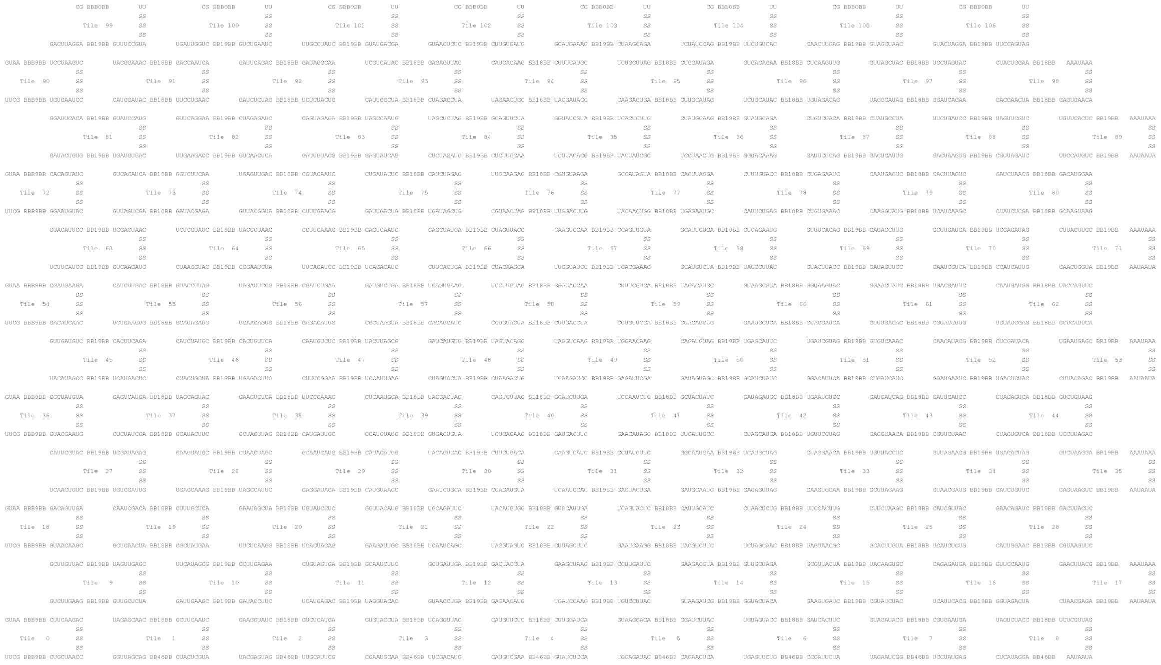

**Supplementary Fig. 42 Sequence map of finite-sized 2D self-assembly of dsRNA bricks.** The generated 9-bp bKL pairs were aligned into different dsRNA bricks and complemented with each other to form the 2D self-assembly. The struts are 27-bp long and marked as “SS”. The beams have different lengths and are marked as “BB”. The numbers in the “BB” indicate the length of the beams. Notably, the length of the beams in the lowest layer are 46-bp, which will act as the seed for the structure formation.

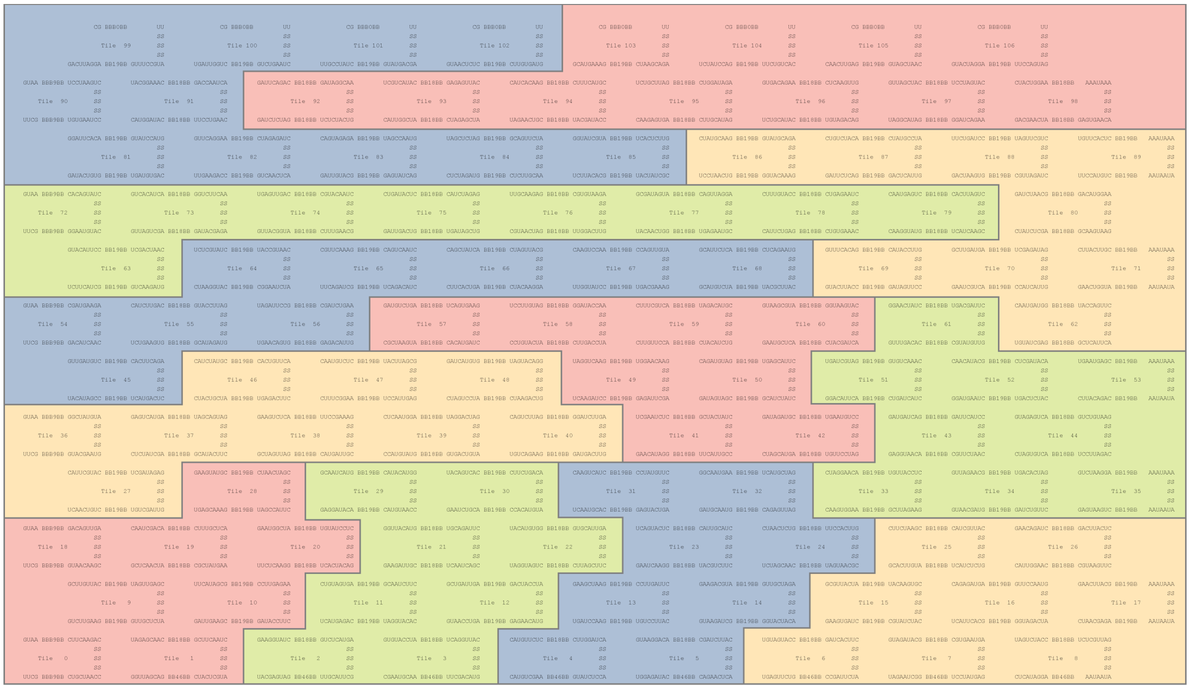

**Supplementary Fig. 43 Sequence map of finite-sized 2D self-assembly of dsRNA bricks with 107 bricks.**In the sequence map, the whole structure is split into 12 different modules. Each module contains different numbers of bricks and is marked with different colors. Each module will be encoded into one plasmid for order and IVT. Different modules can be used in different designs.

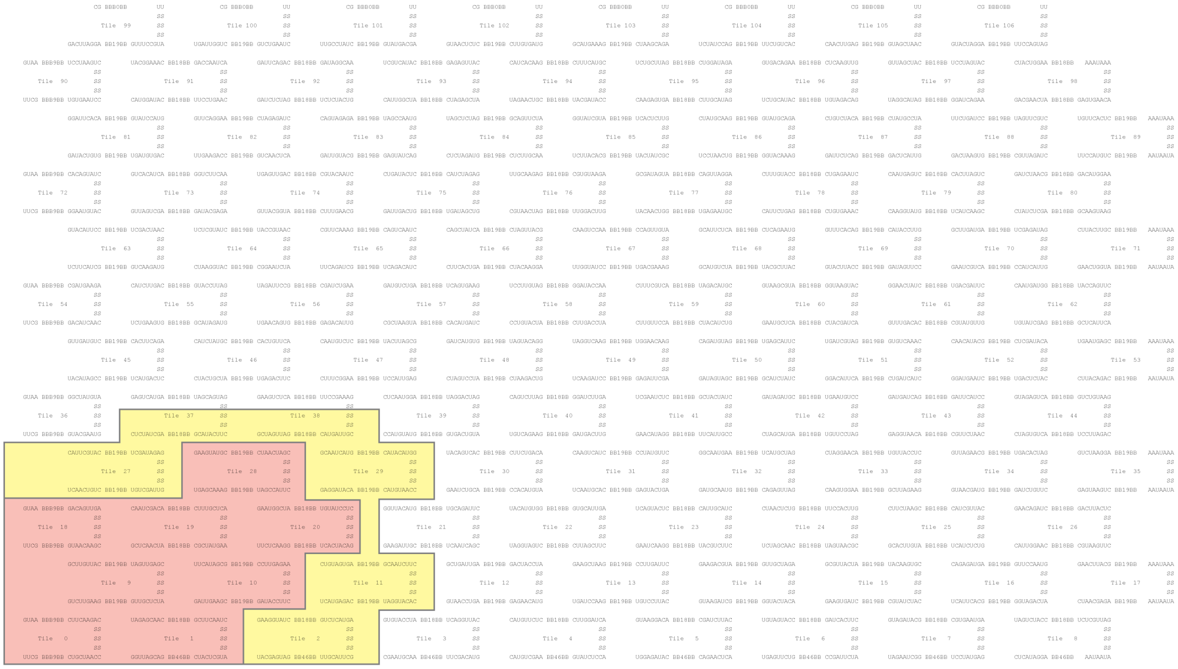

**Supplementary Fig. 44 Sequence map of finite-sized 2D self-assembly of dsRNA bricks with 14 bricks.** One module is used for the synthesis of 2D_14. The yellow area indicates the design for the new bBs for 2D_14.

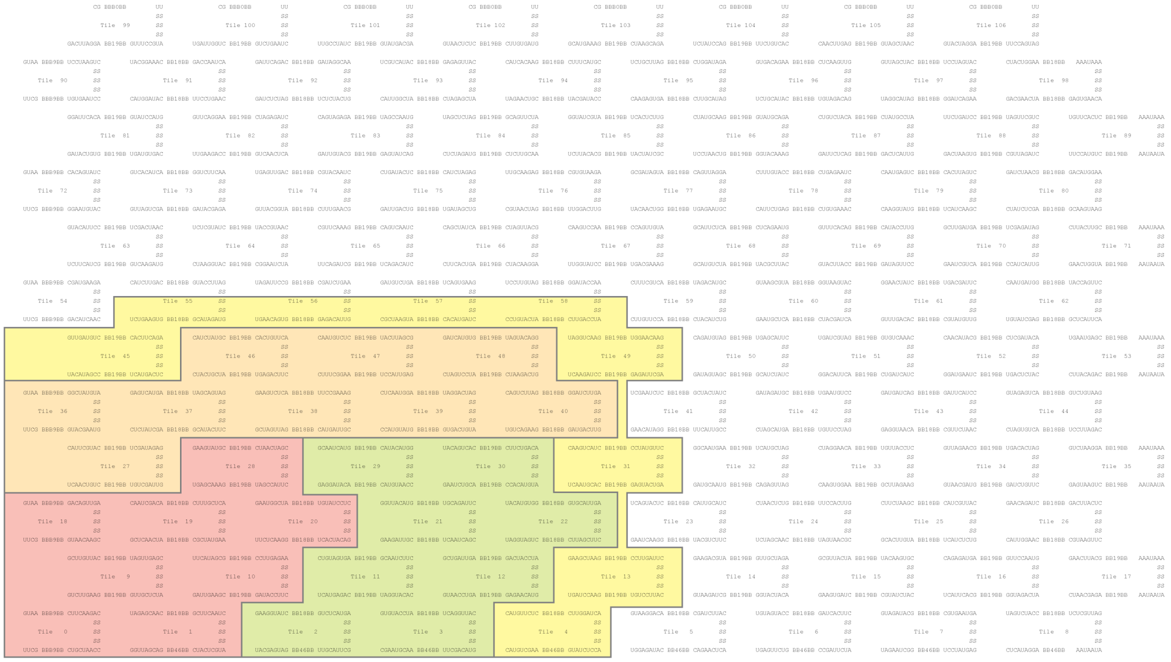

**Supplementary Fig. 45 Sequence map of finite-sized 2D self-assembly of dsRNA bricks with 34 bricks.** Three modules are used for the synthesis of 2D_34. The yellow area indicates the design for the new bBs for 2D_34.

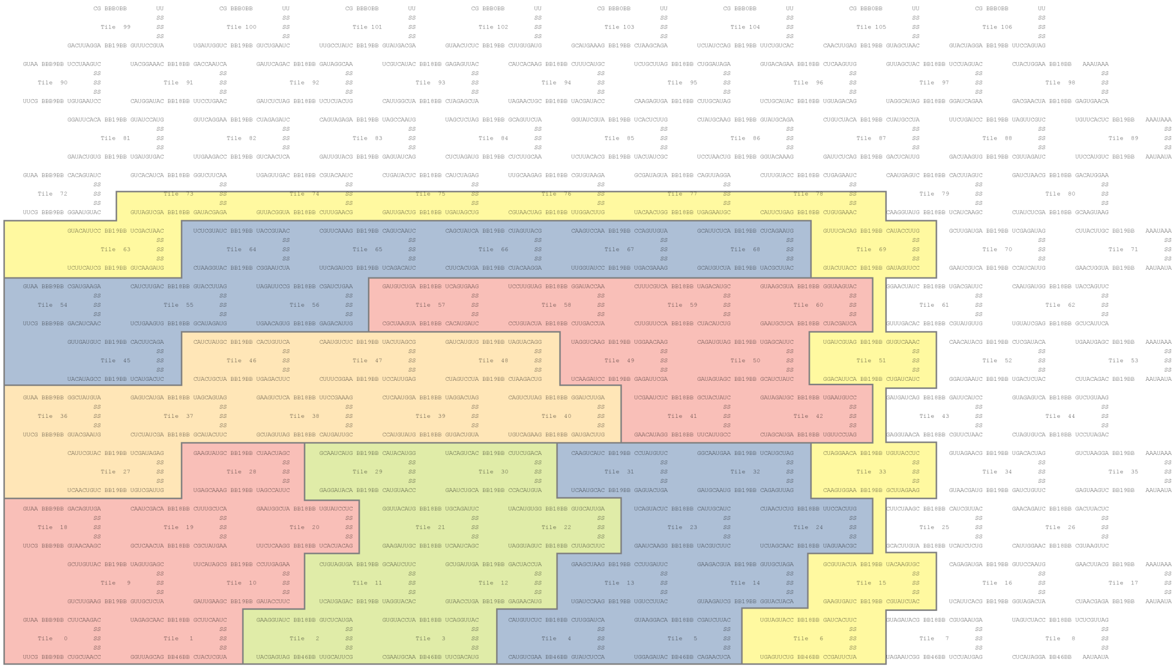

**Supplementary Fig. 46 Sequence map of finite-sized 2D self-assembly of dsRNA bricks with 62 bricks.** Six modules are used for the synthesis of 2D_62. The yellow area indicates the design for the new bBs for 2D_62.

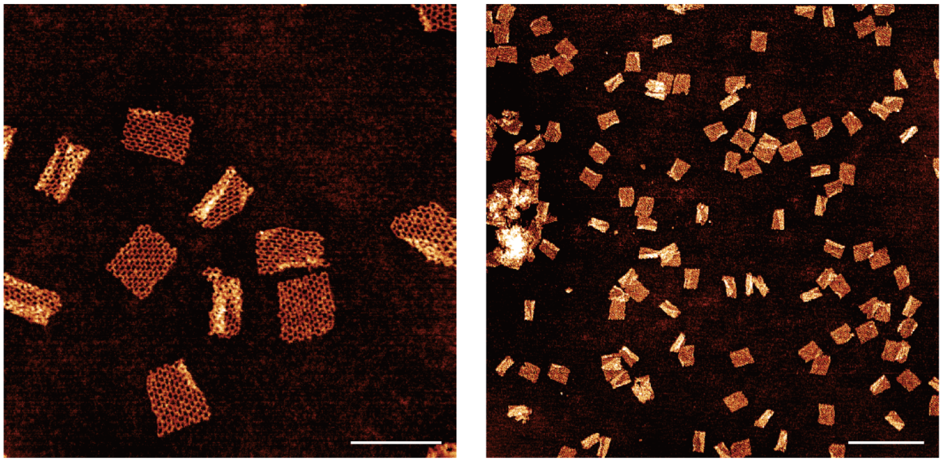

**Supplementary Fig. 47 Zoomed-in and zoomed-out AFM images of 2D_107.** The sample was prepared using 100 nM bricks in 1× TAE buffer with 300 mM $Na^+$ and 30 mM free $Mg^{2+}$. Scale bars: 200 nm (left) and 500 nm (right).

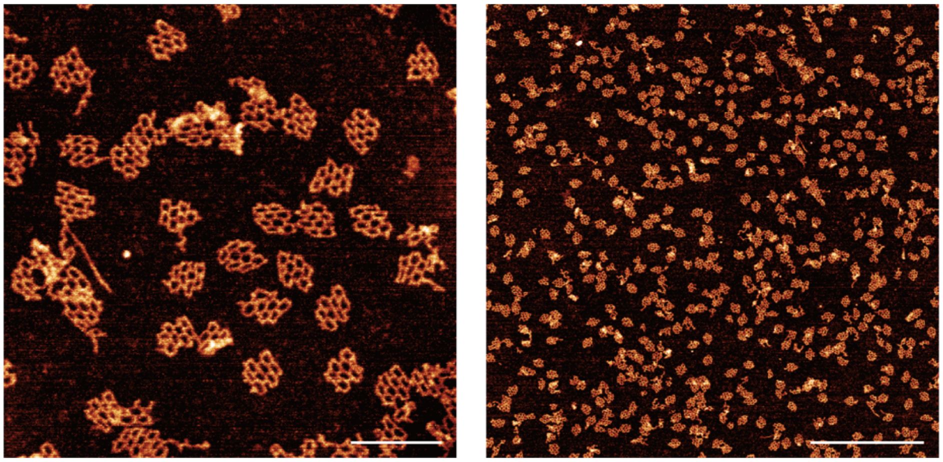

**Supplementary Fig. 48 Zoomed-in and zoomed-out AFM images of 2D_14.** The sample was prepared using 40 nM bricks in 1× TAE buffer with 1 mM free $Mg^{2+}$. Scale bars: 100 nm (left) and 500 nm (right).

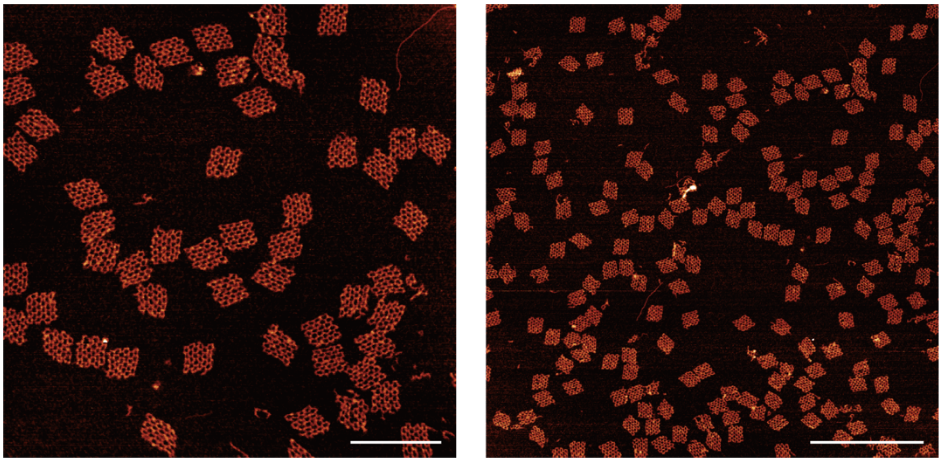

**Supplementary Fig. 49 Zoomed-in and zoomed-out AFM images of 2D_34.** The sample was prepared using 40 nM bricks in 1× TAE buffer with 1 mM free $Mg^{2+}$. Scale bars: 200 nm (left) and 500 nm (right).

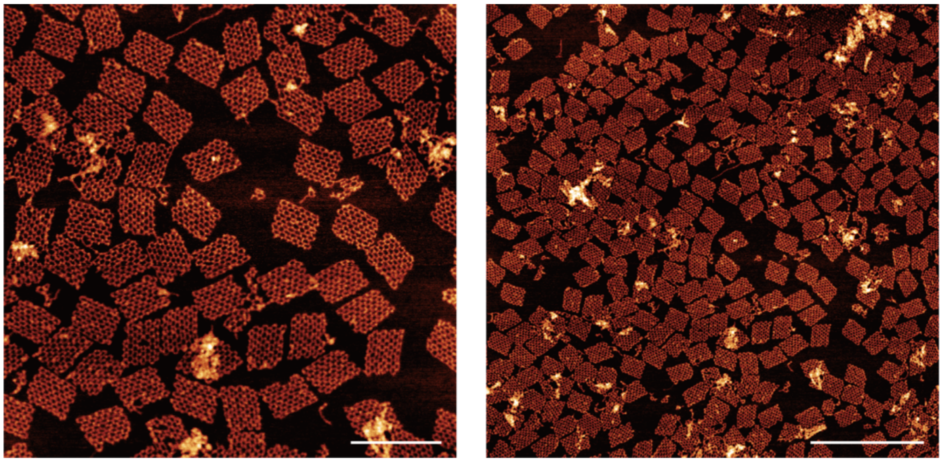

**Supplementary Fig. 50 Zoomed-in and zoomed-out AFM images of 2D_62.** The sample was prepared using 40 nM bricks in 1× TAE buffer with 1 mM free $Mg^{2+}$. Scale bars: 200 nm (left) and 500 nm (right).

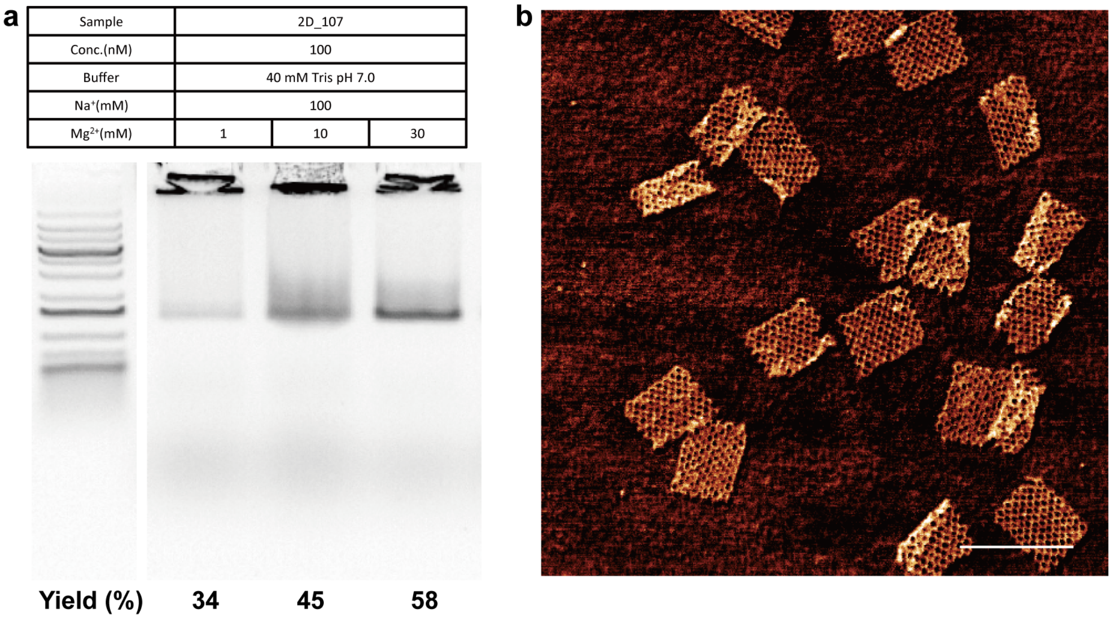

**Supplementary Fig. 51 Synthesis of 2D self-assembly of dsRNA bricks with 107 bricks.a,** Agarose gel analysis of 2D_107 with different $Mg^{2+}$ concentrations. The higher $Mg^{2+}$ concentration gives higher yield of 2D_107 structure. Apparent yield was calculated by dividing the background-corrected intensity of the main product band by the total corrected intensity in the migrated portion of each lane. Signal remaining in the loading wells was excluded. Material was retained in some wells, although AFM images of the corresponding samples did not reveal obvious large aggregates. The measured main-band fraction may also have been reduced by structural destabilization or partial disassembly during electrophoresis. **b,** AFM images of 2D_107 with 100 nM bricks in 40 mM Tris, 100 mM $Na^+$ and 30 mM $Mg^{2+}$. Scale bar, 200 nm.

| Sample | 2D_107 | | | |
| --- | --- | --- | --- | --- |
| Conc. (nM) | 10 | 16 | 40 | 80 |
| Buffer | 40 mM Tris, pH 7.0 | | | |
| $Na^{+}$ (mM) | 200 | | | |
| $Mg^{2+}$ (mM) | 1 | | | |

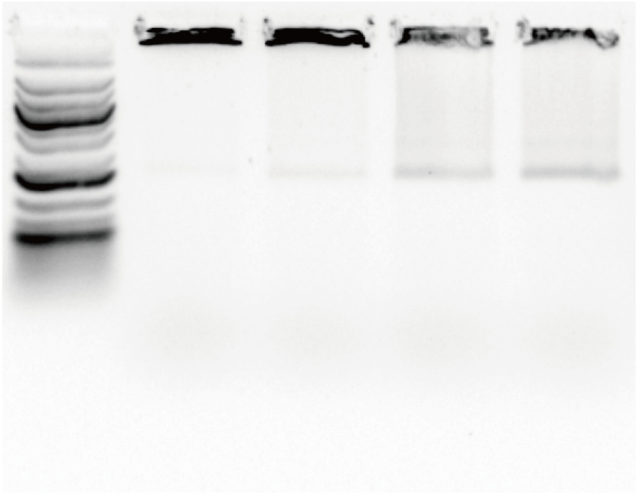

**Yield (%)** **16** **23** **31** **37**

**Supplementary Fig. 52 Agarose gel analysis of 2D_107 with different brick concentrations.**In the low dsRNA brick concentration, no band for 2D_107 was observed. Higher brick concentration gives higher yield.

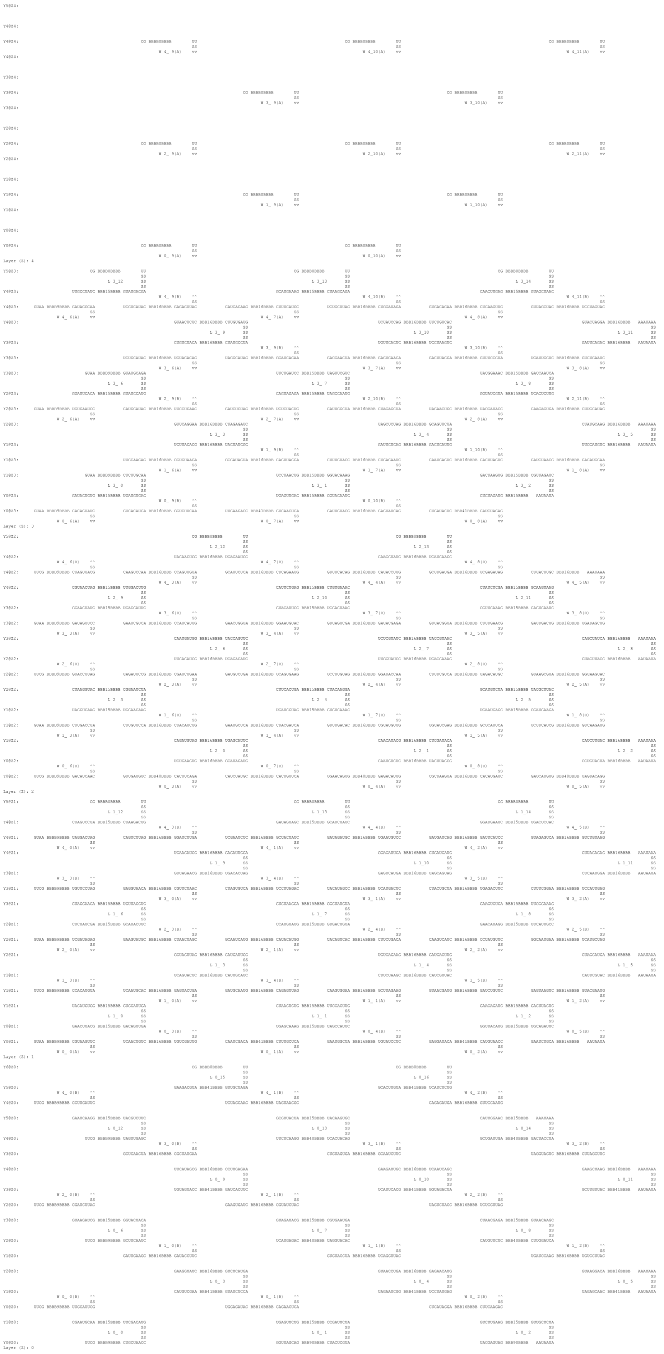

**Supplementary Fig. 53 Sequence map of finite-sized 3D self-assembly of dsRNA bricks.** This sequence map shows how dsRNA bricks self-assemble to form the 3D nanostructures. The struts are 27-bp long and marked as “SS”. The beams have different lengths and are marked as “BB”. The numbers in the “BB” indicate the length of the beams. On the left, the position of the helix is marked as Ya@Zb. For example, Y2@Z0 indicates the third helix in the y-direction on the first layer. There are two types of bricks in the map: x-y bricks are denoted L, and x-z bricks are denoted W. The x-z bricks are split into two parts in the map: the half interact with lower x-y plane named as B, and the half interact with upper x-y plane named as U. The position is marked on the left of the map. For example, the W 0_1 (B) in the Y0@z0 and W 0_1 (U) in the Y0@z1 are from the same brick.

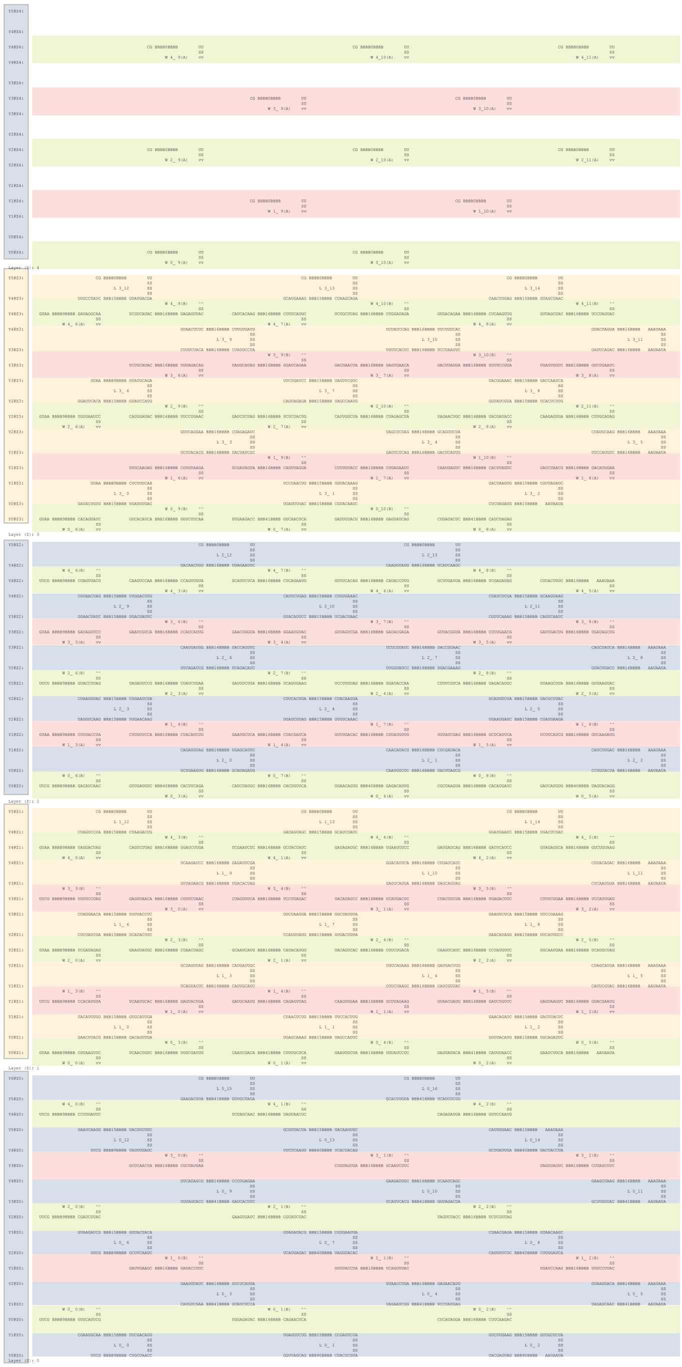

**Supplementary Fig. 54 Sequence map of finite-sized 3D self-assembly of dsRNA bricks with 118 bricks.** The blue and yellow color denote the x-y bricks within the same layer. The red and green color denote the x-z bricks that span between layers within the same y-section.

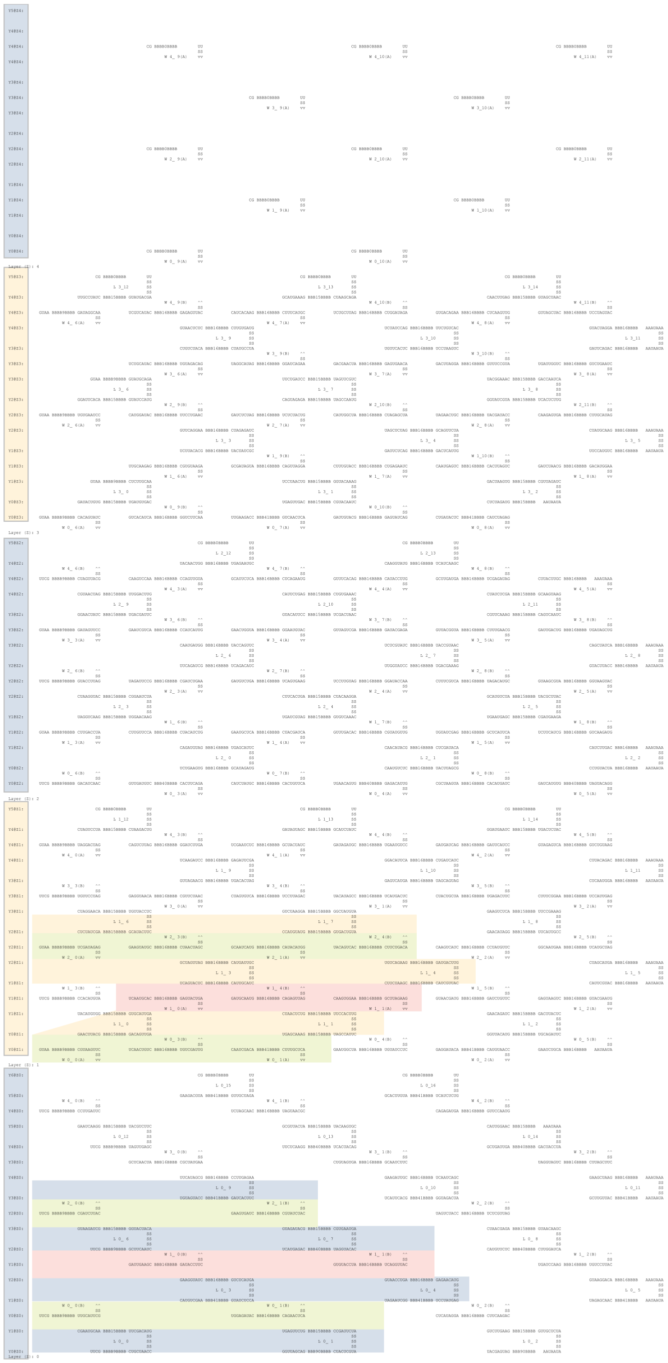

**Supplementary Fig. 55 Sequence map of finite-sized 3D self-assembly of dsRNA bricks with 23 bricks.**

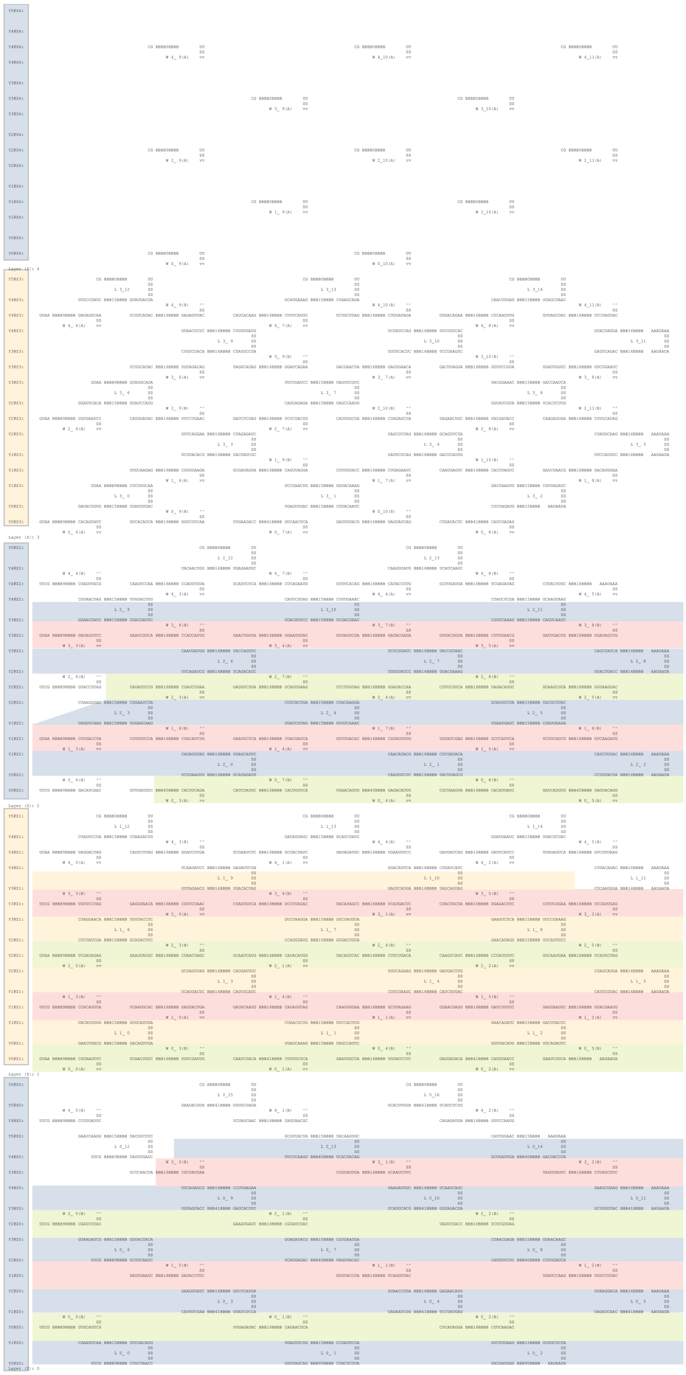

**Supplementary Fig. 56 Sequence map of finite-sized 3D self-assembly of dsRNA bricks with 71 bricks.**

| Sample | 3D_118 | | | | | | | | | | | |
|---|---|---|---|---|---|---|---|---|---|---|---|---|
| Concentration (nM) | 5 | | | | 10 | | | | 16 | | | |
| Buffer | 40 mM Tris, pH7.0 | | | | | | | | | | | |
| $Na^+$ (mM) | 100 | | | | | | | | | | | |
| $Mg^{2+}$ (mM) | 5 | 10 | 20 | 30 | 5 | 10 | 20 | 30 | 5 | 10 | 20 | 30 |

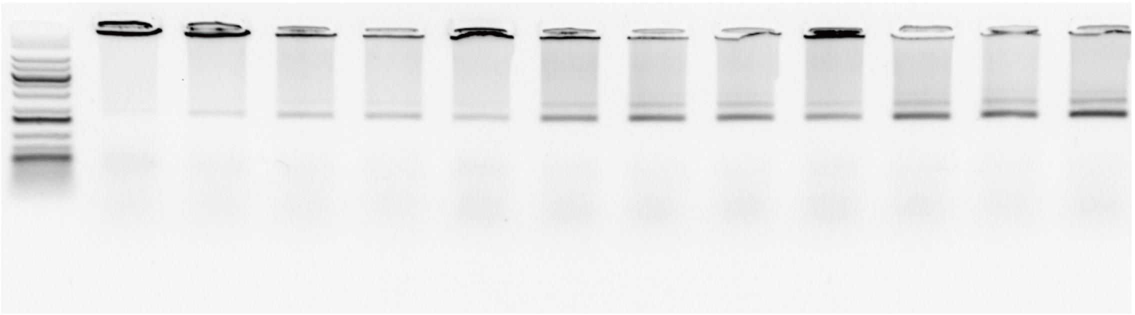

**Yield (%) 15 20 24 27 22 28 33 35 28 36 42 43**

**Supplementary Fig. 57 Assembly optimization of 3D self-assembly of dsRNA bricks with 118 bricks in different $Mg^{2+}$ concentrations.**Similar to the 2D self-assembly, a high $Mg^{2+}$ concentration (30 mM) remains important for the formation of 3D self-assembly of dsRNA bricks.

a

| Sample | 3D_118 | | | |
|---|---|---|---|---|
| Conc.(nM) | 10 | 16 | 40 | 80 |
| Buffer | 40 mM Tris pH 7.0 | | | |
| $Na^+$(mM) | 100 | | | |
| $Mg^{2+}$(mM) | 30 | | | |

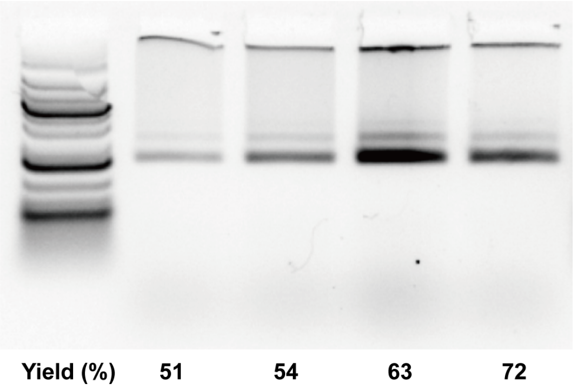

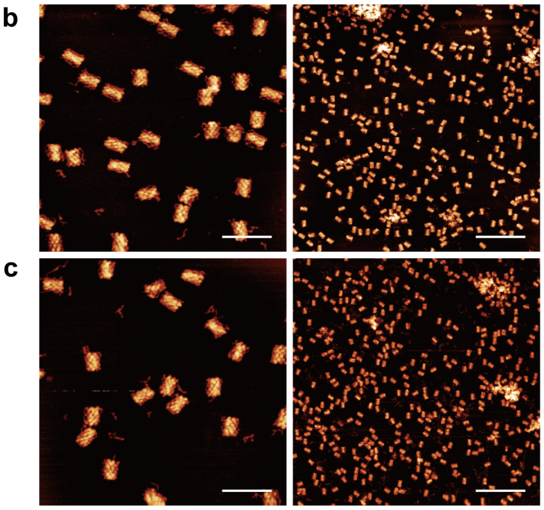

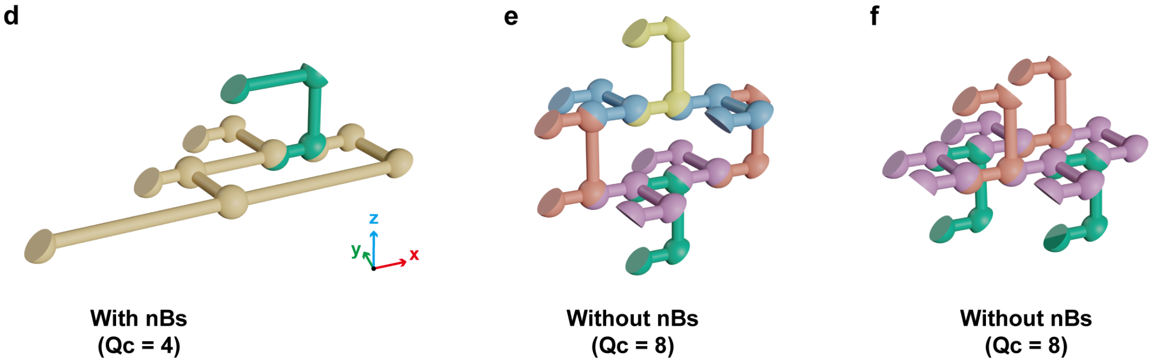

**Supplementary Fig. 58 Assembly optimization of 3D_118 in different brick concentrations. a,** The agarose gel analysis of **3D_118** with different brick concentrations. **b,** The zoomed-in (left, scale bar: 200 nm) and zoomed-out (right, scale bar: 500 nm) AFM images of **3D_118** with 40 nM bricks in 40 mM Tris, 100 mM $Na^+$ and 30 mM $Mg^{2+}$. **c,** The zoomed-in (left, scale bar: 200 nm) and zoomed-out (right, scale bar: 500 nm) AFM images of **3D_118** with 80 nM bricks in 40 mM Tris, 100 mM $Na^+$ and 30 mM $Mg^{2+}$. **d**, The demonstration of nucleation with nBs. Only 4 bricks are needed to form a stable closed ring to initialize the growth in 3D. **e-f**, Without nBs, the 8 cBs are required to initialize the growth in the x-z plane (**e**) and x-y plane (**f**). Qc: coordination number. Interestingly, the **3D_118** exhibited a much higher yield (72%) at higher brick concentration (80 nM). This improved efficiency is attributed to the nucleating brick design. The stable nucleation site for 3D architecture is inherently more challenging, making nonspecific nucleation events far less probable. Our specially designed nBs can substantially reduce the nucleation barrier, thereby enabling high yields.

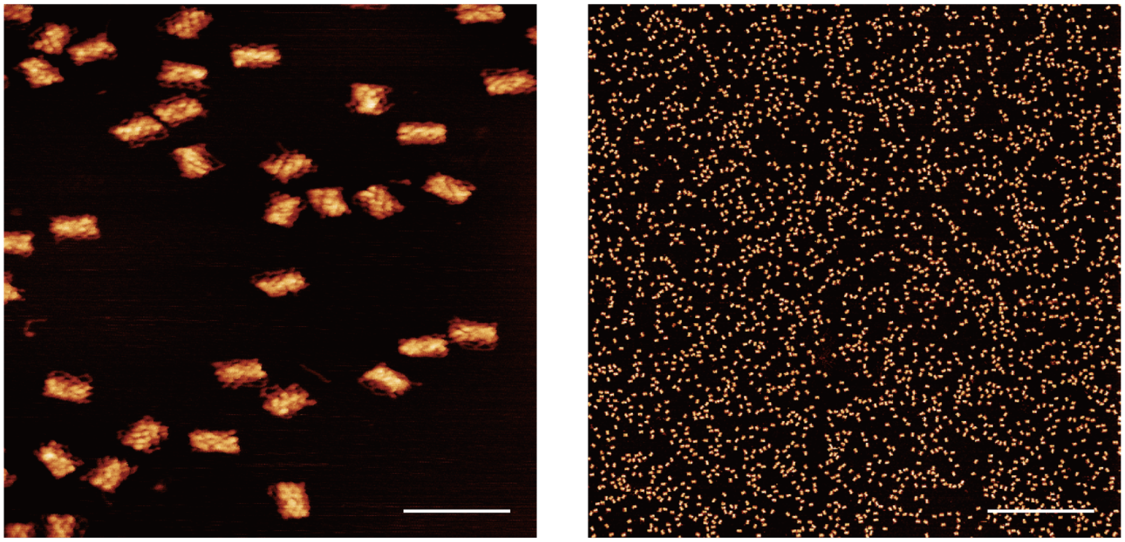

**Supplementary Fig. 59 Zoomed-in and zoomed-out AFM images of purified 3D_118.** The sample was prepared using 80 nM bricks in 40 mM Tris with 100 mM $Na^+$ and 30 mM $Mg^{2+}$. The sample is purified by cutting agarose gel. Scale bars: 200 nm (left) and 2 μm (right).

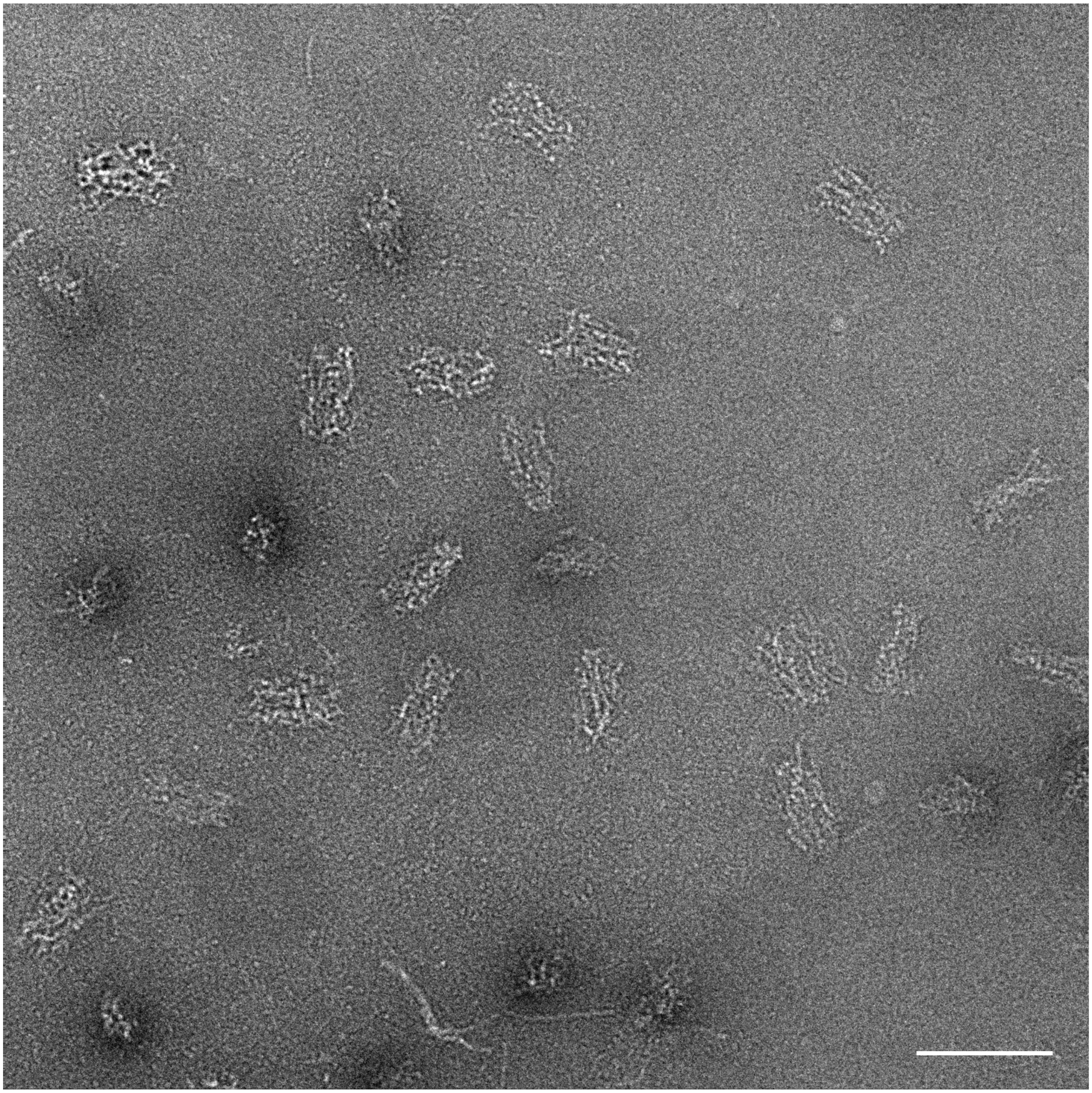

**Supplementary Fig. 60 Zoomed-out TEM image of 3D_118.** Scale bar, 100 nm.

| Sample | 3D_71 | | | 3D_23 | | |
|---|---|---|---|---|---|---|
| Conc. (nM) | 40 | | | | | |
| Buffer | 40 mM Tris, pH 7.0 | | | | | |
| $Na^+$ (mM) | 100 | | | | | |
| $Mg^{2+}$ (mM) | 10 | 20 | 30 | 10 | 20 | 30 |

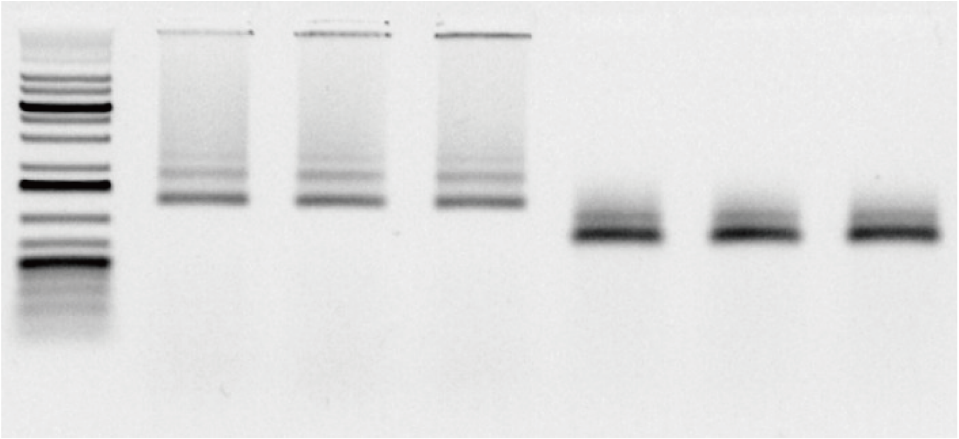

**Yield (%)** **33** **35** **38** **72** **74** **72**

**Supplementary Fig. 61 Assembly optimization of 3D_71 and 3D_23 in different $Mg^{2+}$ concentrations.** Agarose gel analysis of 3D_71 and 3D_23 with 40 nM brick concentrations in 40 mM Tris with 100 mM $Na^+$ and 10/20/30 mM $Mg^{2+}$. 10 mM $Mg^{2+}$ is enough to mediate the formation of the 3D RNA nanostructures.

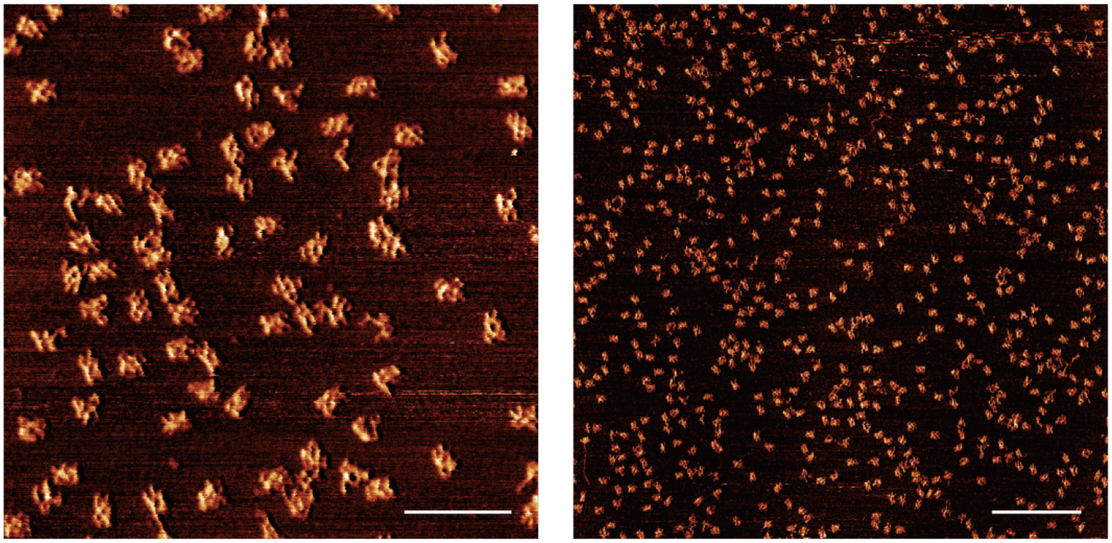

**Supplementary Fig. 62 Zoomed-in and zoomed-out AFM images of 3D_23.** The sample was prepared using 40 nM bricks in 40 mM Tris with 100 mM $Na^+$ and 10 mM $Mg^{2+}$. Scale bars: 200 nm (left) and 500 nm (right).

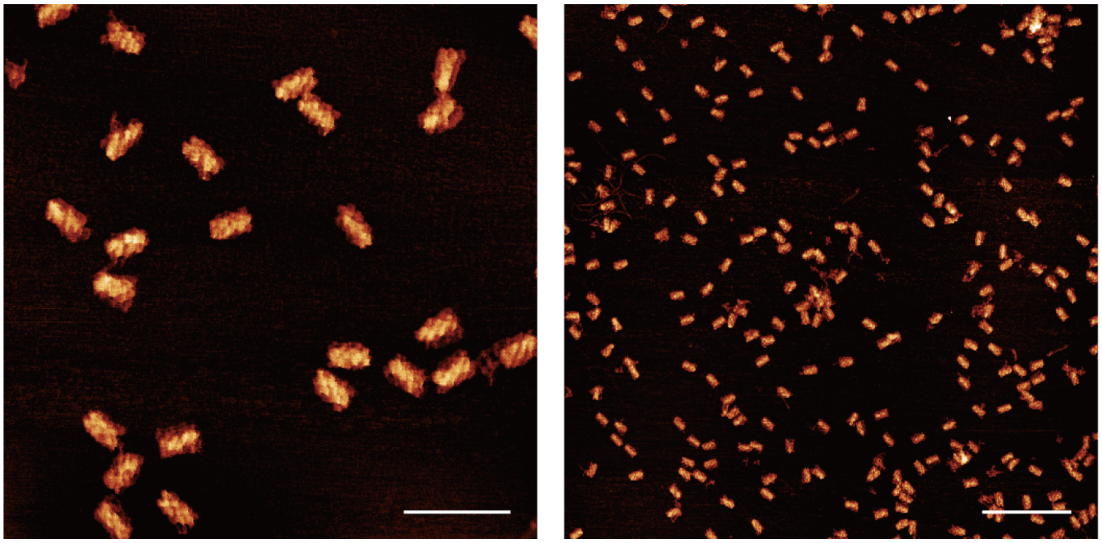

**Supplementary Fig. 63 Zoomed-in and zoomed-out AFM images of 3D_71.** The sample was prepared using 40 nM bricks in 40 mM Tris with 100 mM $Na^+$ and 10 mM $Mg^{2+}$. Scale bars: 200 nm (left) and 500 nm (right).

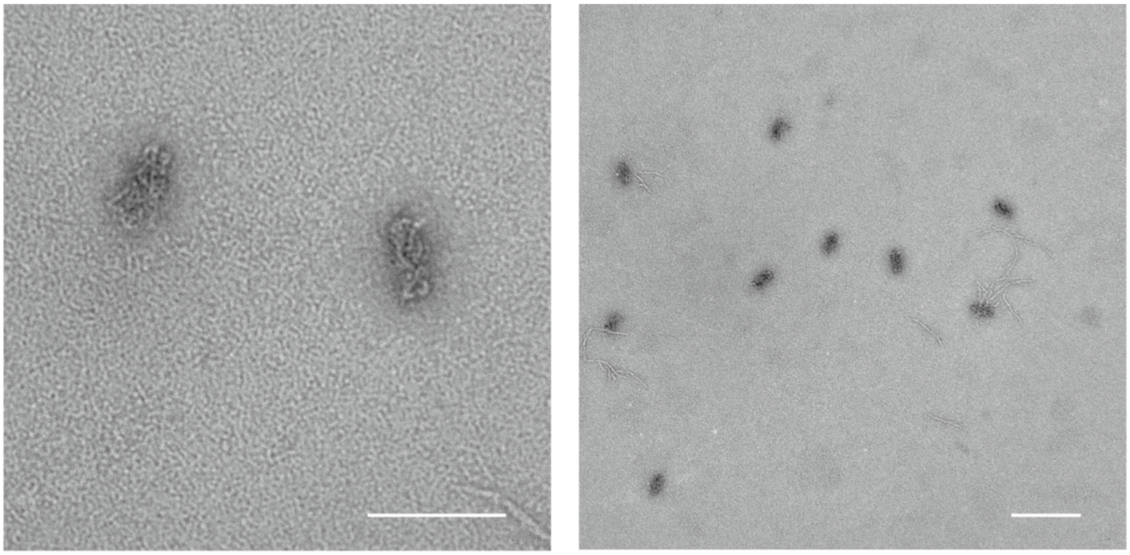

**Supplementary Fig. 64 Zoomed-in and zoomed-out TEM images of 3D_71.** The sample was prepared using 40 nM bricks in 40 mM Tris with 100 mM $Na^+$ and 10 mM $Mg^{2+}$. Scale bars: 100 nm (left) and 200 nm (right).

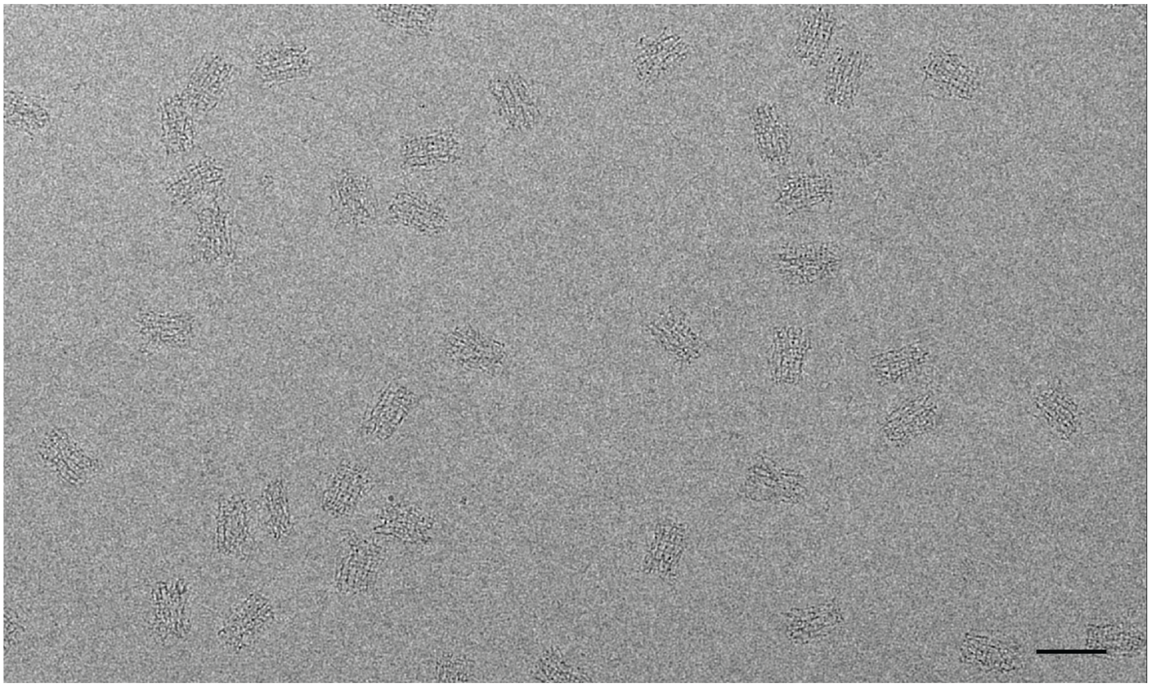

**Supplementary Fig. 65 Representative EM micrograph of 3D_118.** Scale bar, 100 nm.

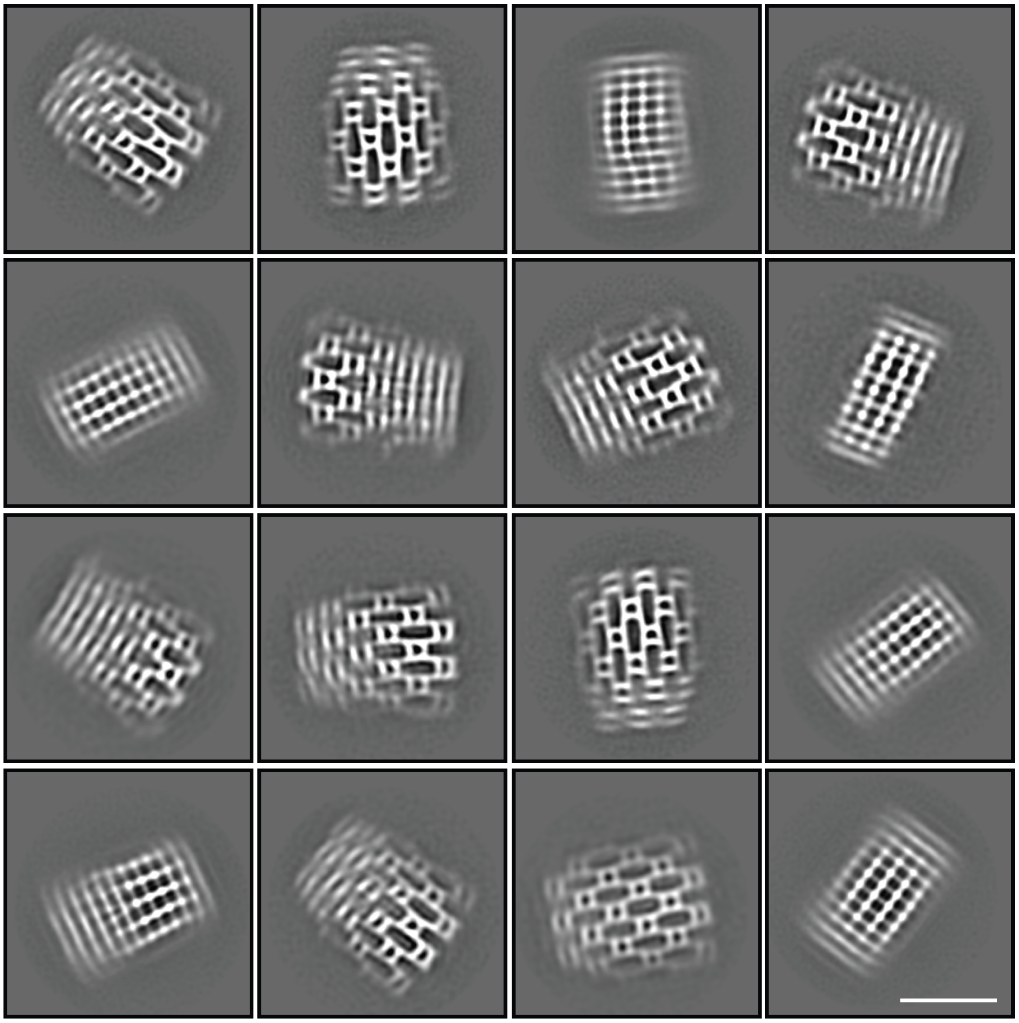

**Supplementary Fig. 66 Representative two-dimensional class averages of 3D_118.** Scale bar, 50 nm.

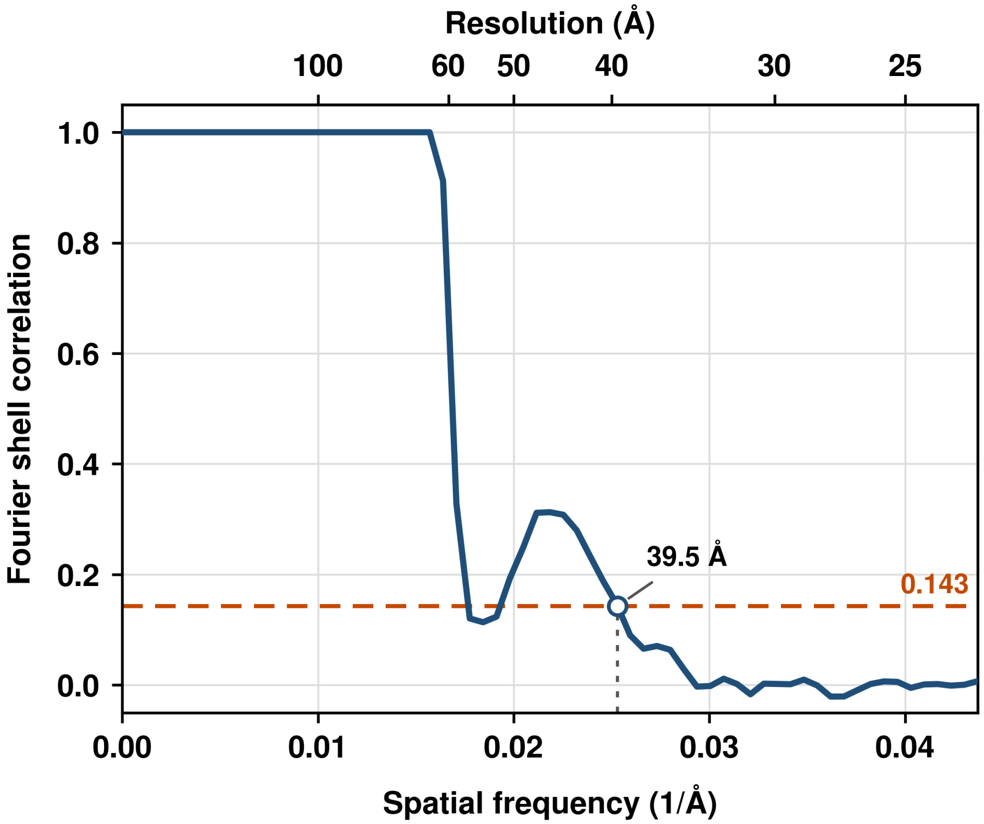

**Supplementary Fig. 67 Fourier shell correlation (FSC) curve of the cryo-EM reconstruction of 3D_118.** The nominal resolution was estimated to be 39.5 Å using the FSC = 0.143 criterion. Spatial frequency and the corresponding real-space resolution are indicated on the lower and upper x axes, respectively.

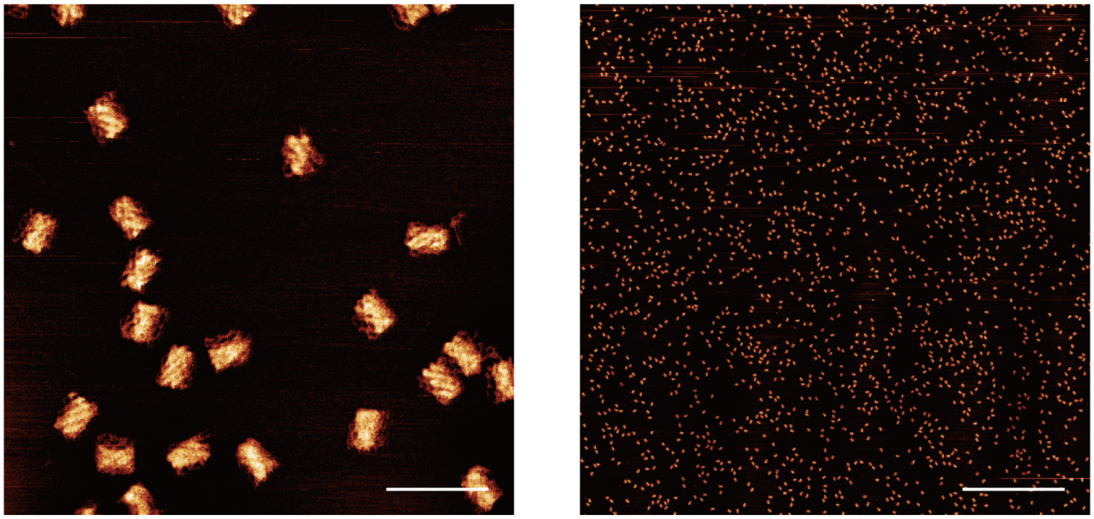

**Supplementary Fig. 68 Zoomed-in and zoomed-out AFM image of purified 3D_118 stored at 4 °C for two weeks.** After storage at 4 °C for two weeks, purified 3D_118 retained its overall morphology, as confirmed by AFM, with no apparent morphological degradation or deformation. This stability is consistent with the predominantly double-stranded architecture of the dsRNA-brick assembly. Scale bars: 200 nm (left) and 2 μm (right).

**a**

| Sample | 2D_14 | | | | | |
|---|---|---|---|---|---|---|
| Conc.(nM) | 40 | | | | | |
| Buffer | 40 mM Tris pH 7.0 | | | | | |
| Cations | 100 mM Na+, 1mMMg | | | | | |
| Preparation | ANL | I32.5 | I34.2 | I37 | I38.9 | I42.3 |

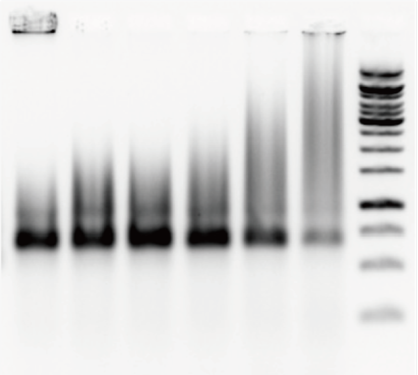

**b**

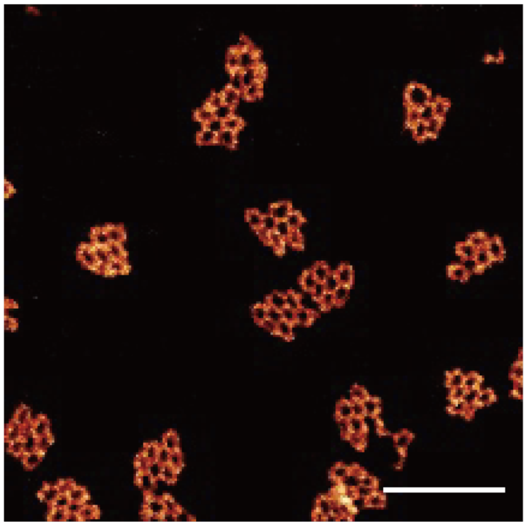

**Supplementary Fig. 69 Isothermal assembly of 2D_14.** **a,** The agarose gel analysis of isothermal assembly of **2D_14** at different temperatures. ANL denotes annealing, which is the same as previously described method. I32.5, I34.2, I37, I38.9 and I42.3 denote the isothermal incubation in 32.5, 34.2, 37, 38.9 and 42.3°C for 1 h, respectively. **b,** AFM image of **2D_14** with 40 nM bricks in 40 mM Tris, 100 mM $Na^+$ and 1 mM $Mg^{2+}$ after isothermal incubation at 37 °C for 1 h. Scale bar, 100 nm.

### Notes

#### I. Selection of orthogonal kissing loop sequences

##### Outline

For `N` (default `9`), the workflow has 4 steps:

1. Generate candidate RNA pool

- Output: `RNA_Pool_Nnt.txt`

2. Filter candidates and build conflict graph

- Outputs: `FilteredPool_Nnt.txt`, `ConflictGraph_Nnt.txt`

3. Select a large non-conflicting set (independent set)

- Output: `SelectedPool_from_graph.txt`

4. Generate full complementary pairs and orthogonality figure

- Outputs: `Orthogonal_RNA_Pool_Nnt.txt`, `orthogonality_Nnt_RNA_Pool.png`

All files are written into a new folder:

- `Orthogonal_RNA_Pool_Nnt` (Here `N` is replaced by your `--num-nt` value, e.g. `9`.)

##### Requirements

- Python 3.9+.
- Step 1–3: standard Python library only.
- Step 4 figure (`orthogonality_Nnt_RNA_Pool.png`): requires `nupack` (and `matplotlib`, `numpy`).
- If `nupack` is not installed, the pipeline still completes and still writes:
  - `Orthogonal_RNA_Pool_Nnt.txt`
  - all step 1–3 files
  - plus a warning: `No NUPACK` (figure is skipped).

##### How To Run

~~~~~~~~~~~~~~~~~~~~~~~~~~~~~~~~~~~~~~~~~~~~~~~~~~~~~~~~~~~~~~~~~~~~~~~~~~~~~~~~~~~~~~~~~~~~

```
python run_orthogonal_selection.py -N 9
```

~~~~~~~~~~~~~~~~~~~~~~~~~~~~~~~~~~~~~~~~~~~~~~~~~~~~~~~~~~~~~~~~~~~~~~~~~~~~~~~~~~~~~~~~~~~~

Example output folder:

- `Orthogonal_RNA_Pool_9nt/`

Example files inside:

- `RNA_Pool_9nt.txt`
- `FilteredPool_9nt.txt`
- `ConflictGraph_9nt.txt`
- `SelectedPool_from_graph.txt`
- `Orthogonal_RNA_Pool_9nt.txt`
- `orthogonality_9nt_RNA_Pool.png`

**Supported Options**

- `-N`, `--num-nt`
  - Meaning: sequence length `N`.
  - Default: `9`.
- `-R`, `--rounds`
  - Meaning: number of randomized attempts in step 3.
  - Default: `1000`.
- `-G`, `--guide`
  - Meaning: guide sequence used in filtering (step 2).
  - Default: `CACGAAGUCAAUAC`.
  - Multiple guides are allowed by repeating `-G`.
- `-O`, `--output`
  - Meaning: filename of step-4 text output.
  - Default: `Orthogonal_RNA_Pool_Nnt.txt`.
- `-U`, `--User`
  - Meaning: show built-in usage instructions.

**Step 1 – `generate_pool_nnt.py`**

**Goal:** enumerate "nice" N-nt KL candidates (for us N = 9) with controlled GC content, no obvious hairpins, and reasonable end patterns.

**How to run it (9-nt example)**

From the project directory:

~~~~~~~~~~~~~~~~~~~~~~~~~~~~~~~~~~~~~~~~~~~~~~~~~~~~~~~~~~~~~~~~~~~~~~~~~~~~~~~~~~~~~~~~

```
python generate_pool_nnt.py 9
```

~~~~~~~~~~~~~~~~~~~~~~~~~~~~~~~~~~~~~~~~~~~~~~~~~~~~~~~~~~~~~~~~~~~~~~~~~~~~~~~~~~~~~~~~

- **Positional argument**: `N` (here 9)
- **Optional**: `--output <filename>`

If you don't give `--output`, it will write to: `RNAPool_9nt.txt` (one sequence per line, all uppercase RNA).

**What does it mean by "4S and (N–4) W"**

It works at two levels:

- **S/W pattern** level: "strong" vs "weak"
  - **S = strong** = G or C
  - **W = weak** = A or U

  **Base assignment** level: choose actual A/G/U/C consistent with that S/W pattern.

For N = 9 the script enforces **exactly 4 S and 5 W**, so **every 9-mer has 4 G/C and 5 A/U**.

**S/W pattern generation (function `gen_pattern_sw`)**

For each 9-nt pattern, it:

- Picks positions of the 4 S's (indices Sa, Sb, Sc, Sd) and fills the rest with W.
- Rejects patterns that:
  - Have **4 consecutive S** (avoids long GC runs),
  - Have a **long run of W at the 5′ end** (avoids very A/U-rich 5′ segments, following the original C++ logic).

Result: a list of strings like "`SWWSWWSWW`" that tell you where G/C vs A/U are allowed.

**Assigning A/G/U/C (function `assign_seq_for_pattern`)**

For each S/W pattern, it enumerates all 2^N assignments:

- At each W position: choose **A or U**
- At each S position: choose **G or C**

For each resulting sequence `s` (length 9), it applies filters:

- **All four nucleotides must appear at least once**
  - `s` must contain **A, G, U, and C** somewhere.

  **Hairpin exclusion (for N ≥ 9, so yes for 9-mers)**
  - It looks at the would-be stem formed by the outer 3 pairs:
    - Pairs `s[0]–s[-1]`, `s[1]–s[-2]`, `s[2]–s[-3]`.

- Two scores:
  - `score` using normal base pairs (AU, UA, GC, CG, GU, UG),
  - `score_comp` using a slightly different "complement" rule (allows AC/CA).
- If **all three** outer positions look pairable in either sense (`score ≥ 3` or `score_comp ≥ 3`), the sequence is **discarded** – it's too hairpin-like.

**No internal "UUU" or mirrored "AAA"**

- For positions `i = 2 ... N-3` (internal):
- Reject if there is `U U U` at `s[i:i+3]`.
- Reject if there is `A A A` at the **mirrored** positions at the 5′ side.
- So we avoid long internal **U-tracts** and **A-tracts**, which are structurally problematic.

Sequences passing these tests are collected into a pool.

**Final refinement filters (function `refine_seq`)**

After pooling all candidates from all patterns, `refine_seq` applies four more rules:

- `if 6_kmry` **– avoid 6-long degenerate runs**

  - K = [G, U]
  - M = [A, C]
  - R = [A, G]
  - Y = [C, U]
  - It tracks the longest consecutive run for each group (K, M, R, Y). If **any group has length ≥ 6**, the sequence is **rejected**. → Roughly: no "6 bases in a row that all behave like purines, pyrimidines, etc."

  `if_ends_ww` **– both ends weak reject**

  - If both 5′ and 3′ ends are A/U (in any combination A/U vs A/U) the sequence is rejected.
  - Ensures **at least one end has a G or C**, which stabilises the designed KL and reduces problematic 3′/5′ fluff.

  `if_ends_3w` **– 3 weak bases at either end**

  - If positions 0–2 or positions N-3–N-1 are all A/U, reject.

  `if_ends_pair_u` **– more detailed end hairpin checks**

  - For N ≥ 9, it looks for patterns like:
  - 5′ A at the start or 3′ U at the end,
  - short A-rich segments at 5′ that could strongly pair with 3′,
  - 3′ terminal U or UU that could form a little end-stem with the 5′ side.
  - If these patterns indicate likely terminal pairing, the sequence is rejected.

**Step 2 – `build_rna_conflict_graphV2.py`**

**Goal:** from the raw pool, throw away self-complementary and guide-complementary sequences, then build a **graph of pairwise conflicts** based on strong complementarity.

**How to run it (9-nt example)**

Basic usage with defaults:

~~~~~~~~~~~~~~~~~~~~~~~~~~~~~~~~~~~~~~~~~~~~~~~~~~~~~~~~~~~~~~~~~~~~~~~~~~~~~~~~~~~~~~~~~~~~~~~~~~~~~~~~~~~~~~~~~~~~~~~~~~~~~~~~~~~~~~~

```
python build_rna_conflict_graphV2.py --num-nt 9
```

~~~~~~~~~~~~~~~~~~~~~~~~~~~~~~~~~~~~~~~~~~~~~~~~~~~~~~~~~~~~~~~~~~~~~~~~~~~~~~~~~~~~~~~~~~~~~~~~~~~~~~~~~~~~~~~~~~~~~~~~~~~~~~~~~~~~~~~

- `--num-nt 9` tells it we are working with 9-mers.
- If you don't give `--input`, it assumes: (the output of Step 1).
- Default guide(s): `DEFAULT_GUIDES = ["CACGAAGUCAAUAC"]` (this is the 15-nt handle used in the bricks scripts).

You can override:

~~~~~~~~~~~~~~~~~~~~~~~~~~~~~~~~~~~~~~~~~~~~~~~~~~~~~~~~~~~~~~~~~~~~~~~~~~~~~~~~~~~~~~~~~~~~~~~~~~~~~~~~~~~~~~~~~~~~~~~~~~~~~~~~~~~~~~~

```
python build_rna_conflict_graphV2.py \
--num-nt 9 \
--input RNAPool_9nt.txt \
--guide CACGAAGUCAAUAC --guide GGGAAAUUU \
```

~~~~~~~~~~~~~~~~~~~~~~~~~~~~~~~~~~~~~~~~~~~~~~~~~~~~~~~~~~~~~~~~~~~~~~~~~~~~~~~~~~~~~~~~~~~~~~~~~~~~~~~~~~~~~~~~~~~~~~~~~~~~~~~~~~~~~~~

The run time of the code could be up to 2 hours.

With defaults you get:

- `FilteredPool_9nt_out.txt`
- `ConflictGraph_9nt_edges_out.txt`

**Step 2a – read and pre-filter**

- **Read all sequences** (must have length `num_nt`).
  **Remove palindromes / self-complementary sequences**
  - A sequence counts as "palindromic" if it is complementary to itself under the same rules used for pairwise complementarity (including GU wobble).

  **Remove sequences that conflict with guide sequences**
  - For each guide:
    - Generate all contiguous 9-mers from the guide.

    - For each pool sequence, check if it is “too complementary” to **any** of those 9-mers (with all reverse-complement orientations).
    - If yes → discard the pool sequence.

This ensures **no KL candidate binds strongly to the constant handle (or any other guide you specify).**

**Step 2b – complementarity rules and the conflict graph**

Now you have a filtered list of sequences `seqs`. The script builds an undirected graph:

- **One node per sequence.**
- An **edge (i, j)** means sequences **i** and **j** are “too complementary” under the custom rules.

Under the hood:

- Base pairs allowed:
    - Watson–Crick: **A-U, U-A, G-C, C-G**
    - GU wobble: **G-U, U-G**
- For each pair `(s1, s2)` it checks **all four combinations**:
    - `s1` vs `s2`
    - `s1` vs reverse-complement(s2)
    - reverse-complement(s1) vs `s2`
    - reverse-complement(s1) vs reverse-complement(s2)

The **complementarity criteria** are exactly summarized in the header of the script:

Two sequences (or their RCs) are considered a **conflicting pair** if **any** of these holds:

- **Global pairing**

    - There exists an alignment with:
    - at least **N–1 paired positions**, and
    - **2 GC pairs**.

    **Very long consecutive pairing**

    - Alignment has a **longest consecutive run ≥ N–1**, and
    - **1 GC pair** in that alignment.

    **Medium consecutive pairing**

    - Longest consecutive run ≥ **N–2**, and
    - **2 GC pairs**.

    **Shorter but GC-rich**

    - Longest consecutive run ≥ **N–3**, and

- **≥ 4 GC pairs**.

**Bulged pairing (N ≥ 9)**

- There exists a **single-bulge alignment** of length **N–1** between the two sequences (one base skipped on one strand) with **≥ 3 GC pairs**.

For **N = 9** these thresholds are:

- N–1 = **8**, N–2 = **7**, N–3 = **6**.
- So two 9-mers conflict if e.g.:
- They can form ~8 bp with several GC pairs, **or**
- They have a contiguous 8-bp run with >1 GC, **or**
- They have a contiguous 7-bp run with >2 GC, **or**
- They have a contiguous 6-bp run with ≥4 GC, **or**
- They can form an **8-bp single-bulged** alignment with ≥3 GC.

This is more nuanced than just "no 5-bp perfect complement"—it balances **length of interaction** and **GC richness**, which controls the expected stability.

**Output of Step 2 (for N=9):**

- **Filtered sequences** (post palindrome + guide filter)

  → `FilteredPool_9nt_out.txt`

- **Conflict graph edge list**

  → `ConflictGraph_9nt_edges_out.txt` (each line: i<TAB>j, with $0 \leq i < j$)

**Step 3 – `select_from_conflict_graph.py`**

**Goal:** choose a large subset of sequences that have **no edges between them** – a big independent set in the conflict graph.

**How to run it (9-nt example)**

If you used the defaults in Step 2:

~~~~~~~~~~~~~~~~~~~~~~~~~~~~~~~~~~~~~~~~~~~~~~~~~~~~~~~~~~~~~~~~~~~~~~~~~~~~~~~~~~~~~~~~~~~~~~~~~~~
python select_from_conflict_graph.py \
--seq-file FilteredPool_9nt_out.txt \
--graph-file ConflictGraph_9nt_edges_out.txt \
--rounds 1000
~~~~~~~~~~~~~~~~~~~~~~~~~~~~~~~~~~~~~~~~~~~~~~~~~~~~~~~~~~~~~~~~~~~~~~~~~~~~~~~~~~~~~~~~~~~~~~~~~~~

It will write: `SelectedPool_from_graph_out.txt`

Each line:

~~~~~~~~~~~~~~~~~~~~~~~~~~~~~~~~~~~~~~~~~~~~~~~~~~~~~~~~~~~~~~~~~~~~~~~~~~~~~~~~~~~~~~~~

```
index<TAB>sequence
```

~~~~~~~~~~~~~~~~~~~~~~~~~~~~~~~~~~~~~~~~~~~~~~~~~~~~~~~~~~~~~~~~~~~~~~~~~~~~~~~~~~~~~~~~

(the index matches the line number in `FilteredPool_9nt_out.txt`).

Key arguments:

- `--seq-file`: filtered sequences from Step 2
- `--graph-file`: edge list from Step 2
- `--rounds`: number of randomized greedy attempts (default 1000)
- `--seed`: random seed (optional but good for reproducibility)
- `--output`: base name for output file (default `SelectedPool_from_graph.txt` → `SelectedPool_from_graph_out.txt`)

**Step 3a – greedy independent set**

Core routine: `greedy_independent_set(adj, rng, seed=None)`

Algorithm:

- Start with:
  - `remaining = [0, 1, ..., n-1]` (all nodes),
  - `selected = []`.

  Optionally add a **seed** set of already-independent nodes.
  While there are nodes remaining:

  - Pick a random node `v` from `remaining`.
  - Add `v` to `selected`.
  - Remove `v` **and all its neighbors** from `remaining`.

This produces a **maximal** independent set (you can't add any more nodes without creating conflicts), but not necessarily maximum.

In `main`, the script runs this many times:

~~~~~~~~~~~~~~~~~~~~~~~~~~~~~~~~~~~~~~~~~~~~~~~~~~~~~~~~~~~~~~~~~~~~~~~~~~~~~~~~~~~~~~~~

```
for round_idx in range(args.rounds):
selected = greedy_independent_set(adj, rng)
if len(selected) > len(best_indices):
best_indices = selected
```
~~~~~~~~~~~~~~~~~~~~~~~~~~~~~~~~~~~~~~~~~~~~~~~~~~~~~~~~~~~~~~~~~~~~~~~~~~~~~~~~~~~~~~~~

~~~~~~~~~~~~~~~~~~~~~~~~~~~~~~~~~~~~~~~~~~~~~~~~~~~~~~~~~~~~~~~~~~~~~~~~~~~~~~~~~~~~~~~~~~~~~~~~~~~~~~~~~~~~~~~

It keeps the largest independent set it has seen so far (`best_indices`).

**Step 3b – optimization pass (`optimize_select_graph`)**

To push the size a bit further, there's a second phase that tries to improve `best_indices`:

- Print "Before optimization: k" where k = `len(best_indices)`.
- If `k == 0`, nothing to do.
- Otherwise:
- Set `count_unchanged = 0`.
- Repeat while `count_unchanged ≤ 120 * current_size`:
- If current size ≤ 1, break.
- Create a **seed** by dropping the first node:
- Run `greedy_independent_set(adj, rng, seed=seed)`.
- If the new set is **larger**, accept it and reset `count_unchanged = 0`.
- Otherwise, accept it but increment `count_unchanged`.
- You can hit **Ctrl-C** to stop earlier.

Intuition: "kick out" one sequence and re-grow greedily: sometimes this lets you escape a local optimum and add more sequences overall.

**Final output:** a large set of **mutually non-conflicting 9-nt KL candidates** in `SelectedPool_from_graph_out.txt`. (Typically, >220 candidates; if you are lucky, >230.)

The result is a **large, algorithmically optimized set of 9-nt KL sequences** that:

- behave nicely on their own,
- don't bind strongly to the constant handle,
- and are predicted not to bind strongly to each other in any orientation.

**Step 4 – `pool_rna_complement.py`**

This final pool is what you feed into tile-design scripts (2D/3D bricks), which then assign one 9-nt bulge (with its reverse-complement loop) to each bKL edge in the structure.

The **tile-design scripts** expect a **KL pool file** where each non-empty line has **at least two** RNA sequences, and they treat the **first column** as the 9-nt bulge sequence.

**How to run it (9-nt example)**

To get a compatible pool file from the graph-selected list, we run:

~~~~~~~~~~~~~~~~~~~~~~~~~~~~~~~~~~~~~~~~~~~~~~~~~~~~~~~~~~~~~~~~~~~~~~~~~~~~~~~~~~~~~~~~~~~~~~~~~~~~~~~~~~~~~~~

```
python pool_rna_complement.py SelectedPool_from_graph_out.txt
```

~~~~~~~~~~~~~~~~~~~~~~~~~~~~~~~~~~~~~~~~~~~~~~~~~~~~~~~~~~~~~~~~~~~~~~~~~~~~~~~~~~~~~~~

The script now produces two outputs:

1. Complement text file (`*_complement.txt`), finalized by the one-command pipeline as:

- `Orthogonal_RNA_Pool_9nt.txt`

2. Orthogonality figure from pairwise NUPACK free energies:

- `orthogonality_9nt_RNA_Pool.png`

Example (first lines of `Orthogonal_RNA_Pool_9nt/Orthogonal_RNA_Pool_9nt.txt`):

~~~~~~~~~~~~~~~~~~~~~~~~~~~~~~~~~~~~~~~~~~~~~~~~~~~~~~~~~~~~~~~~~~~~~~~~~~~~~~~~~~~~~~~

```
CAUGAUUGC     GUACUAACG
CUCUAGUUC     GAGAUCAAG
CUACUGUUC     GAUGACAAG
GUUACCUCA     CAAUGGAGU
UACGAUCUC     AUGCUAGAG
GCUACAUAG     CGAUGUAUC
```

~~~~~~~~~~~~~~~~~~~~~~~~~~~~~~~~~~~~~~~~~~~~~~~~~~~~~~~~~~~~~~~~~~~~~~~~~~~~~~~~~~~~~~~

Example figure (`orthogonality_9nt_RNA_Pool.png`):

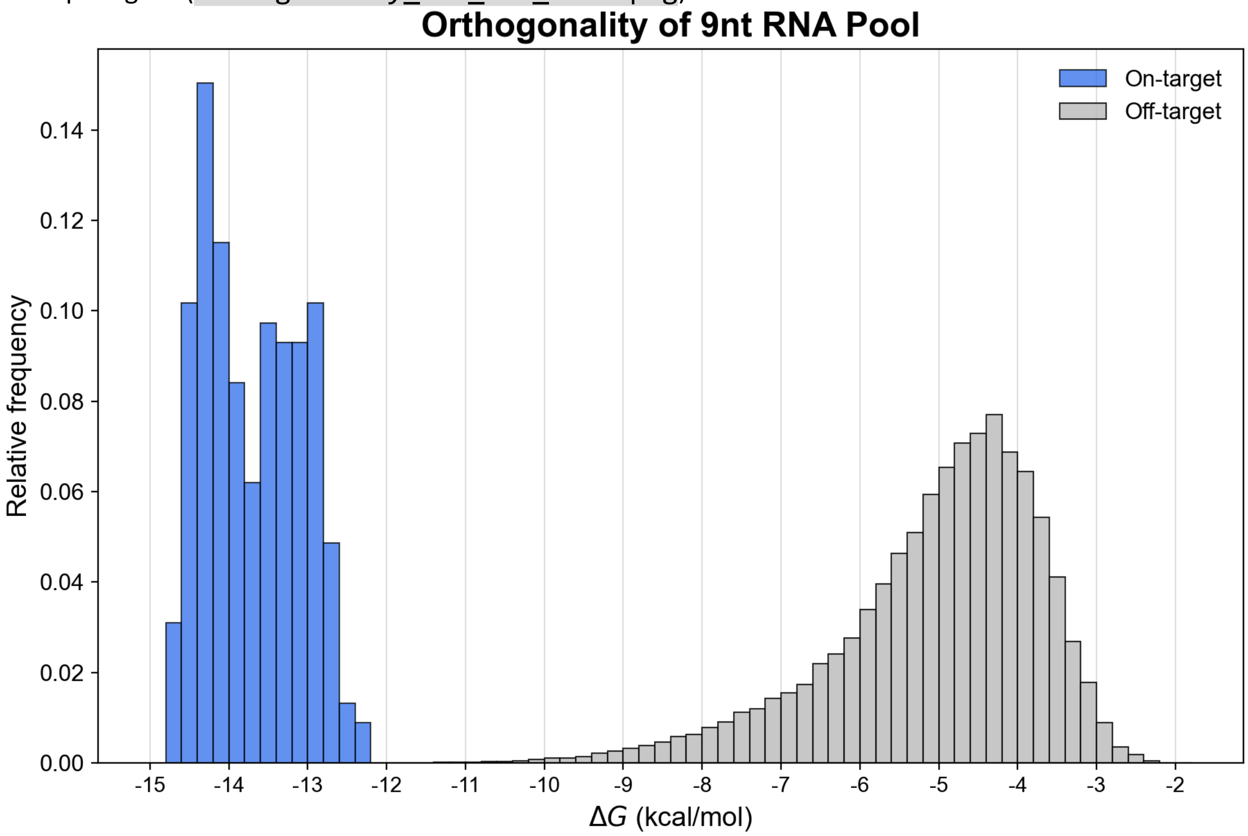
~~~~~~~~~~~~~~~~~~~~~~~~~~~~~~~~~~~~~~~~~~~~~~~~~~~~~~~~~~~~~~~~~~~~~~~~~~~~~~~~~~~~~~~

If NUPACK is unavailable, step 4 still writes the complement text output and prints a warning; the figure is skipped.

NUPACK settings used for the figure:

- `Model(material='rna', celsius=37, sodium=0.1, magnesium=0.010)`
- All pairwise 2-combinations are evaluated.
- Histogram uses bin width `0.2`; On-target and Off-target are plotted separately.

#### II. dsRNA Bricks 2D Builder

##### Outline

This project is a Python GUI tool for designing and exporting 2D dsRNA brick lattices generating tile sequences.

The workflow is:

1. Loading a KL sequence-pool file.
2. Building a 2D lattice from X and Y tile rules.
3. Interactive tile selection from both model view and XY map.
4. Sequence export + NUPACK design in one Generate sequence step.

Current outputs:

- <prefix>_tile_structure.txt
- <prefix>_2D_map.svg
- <prefix>_sequence.txt

##### Requirements

Python:

- Python 3.12 (tested)

Core packages:

- PySide6==6.9.* (recommended)
- numpy
- pyvista
- pyvistaqt
- vtk (tested with 9.5.2)

Sequence design package:

- nupack (install/configure by following the official documentation: https://docs.nupack.org/)

Installation demo (conda + pip):

```
conda create -n dsrna python=3.12 -y

conda activate dsrna

pip install "pyside6==6.9.*" numpy pyvista pyvistaqt vtk
```

Optional quick check:

~~~~~~~~~~~~~~~~~~~~~~~~~~~~~~~~~~~~~~~~~~~~~~~~~~~~~~~~~~~~~~~~~~~~~~~~~~~~~~~~~~~~~~~~~~~~

```
python -c "import PySide6, pyvista, pyvistaqt, vtk, numpy; print('PySide6',
PySide6.__version__); print('PyVista', pyvista.__version__); print('VTK',
vtk.vtkVersion.GetVTKVersion()); print('NumPy', numpy.__version__)"
```

~~~~~~~~~~~~~~~~~~~~~~~~~~~~~~~~~~~~~~~~~~~~~~~~~~~~~~~~~~~~~~~~~~~~~~~~~~~~~~~~~~~~~~~~~~~~

For NUPACK setup and installation, follow: NUPACK Documentation.

**Run**

From this folder:

~~~~~~~~~~~~~~~~~~~~~~~~~~~~~~~~~~~~~~~~~~~~~~~~~~~~~~~~~~~~~~~~~~~~~~~~~~~~~~~~~~~~~~~~~~~~

```
python build_dsRNA_Bricks_2D.py
```

~~~~~~~~~~~~~~~~~~~~~~~~~~~~~~~~~~~~~~~~~~~~~~~~~~~~~~~~~~~~~~~~~~~~~~~~~~~~~~~~~~~~~~~~~~~~

Alternative:

~~~~~~~~~~~~~~~~~~~~~~~~~~~~~~~~~~~~~~~~~~~~~~~~~~~~~~~~~~~~~~~~~~~~~~~~~~~~~~~~~~~~~~~~~~~~

```
python -m function.build_dsRNA_Bricks_2D
```

~~~~~~~~~~~~~~~~~~~~~~~~~~~~~~~~~~~~~~~~~~~~~~~~~~~~~~~~~~~~~~~~~~~~~~~~~~~~~~~~~~~~~~~~~~~~

**Project Layout**

- build_dsRNA_Bricks_2D.py: launcher.
- function/app.py: main GUI window and workflow wiring.
- function/lattice_builder.py: 2D lattice/tile placement rules.
- function/map2d.py: XY map widget and selection sync.
- function/view3d.py: embedded PyVista view.
- function/c_tiles.py: tile geometry definitions.
- function/rna_tile_generator.py: tile RNA generation.
- function/nupack_runner.py: NUPACK execution pipeline.
- KL_231pairs.txt: example KL sequence-pool file.
- demo_9x12_107tile/: demo outputs.

**Workflow Example (9x12, 107 tiles)**

1. Overview
   This example builds a 9x12 lattice (107 tiles total).

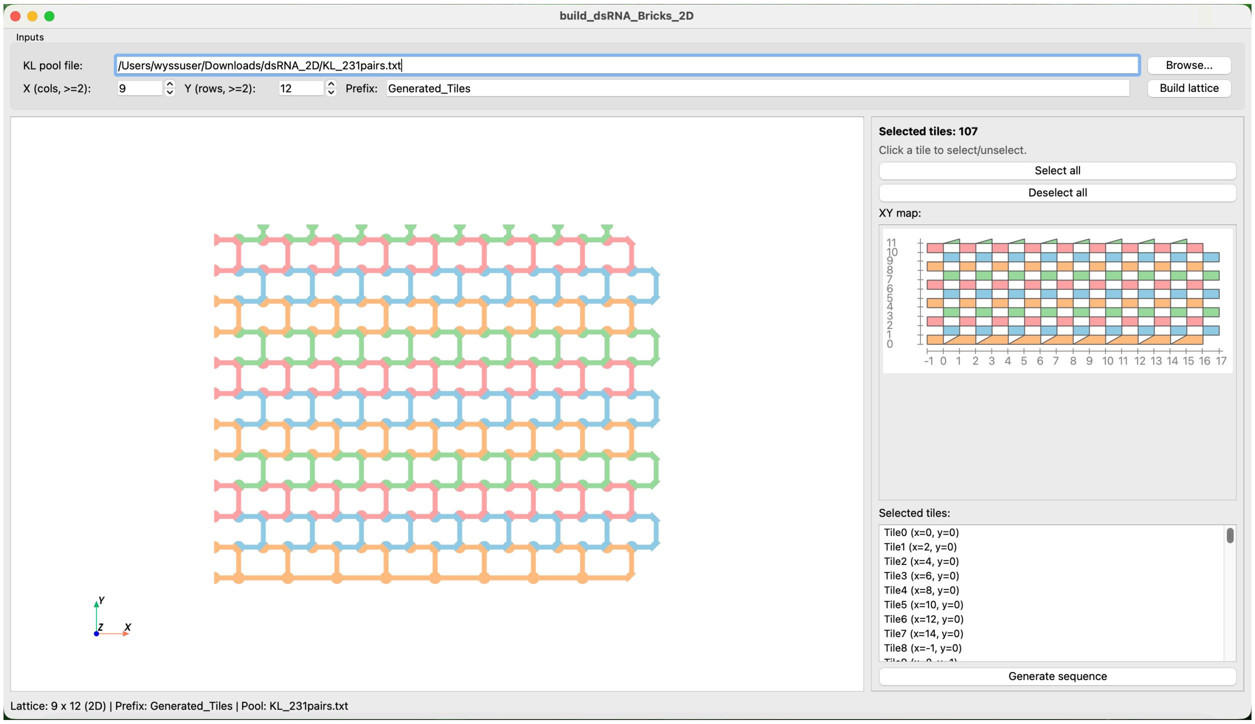

2. Input
   Choose a KL pair pool file. Here we use `KL_231pairs.txt`.

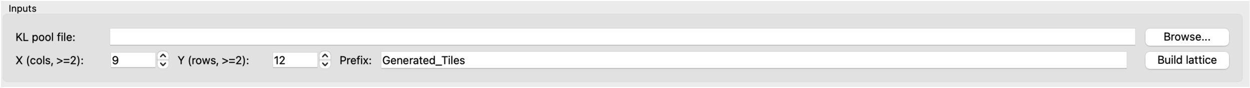

Example lines from `KL_231pairs.txt`:

```
CUAGAUGGA   GAUCUACCU
UGUACCUUC   ACAUGGAAG
UCAGAUUCG   AGUCUAAGC
GCAUGAGUA   CGUACUCAU
GCUCAUCUA   CGAGUAGAU
UAGCCAUUC   AUCGGUAAG
GUUAGAACG   CAAUCUUGC
CCUUAGUUG   GGAAUCAAC
```

Set `X=9`, `Y=12`, and optional prefix.
The required KL pair count for the selected lattice must be smaller than or equal to the pool size.

3. Build the lattice
   Click `Build lattice`. The GUI generates both the model structure and the 2D map.

2D model view:

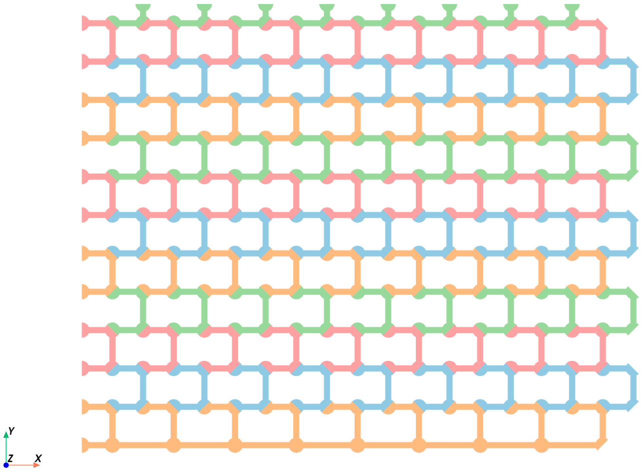

2D XY map:

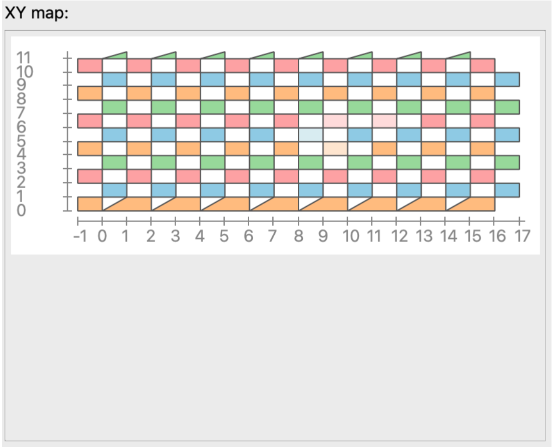

4. Select tiles
   Select/deselect from either the model or XY map. Selection is synchronized and shown in the right-side list. Selection rules before sequence generation:

   - No flexible tiles: each selected tile must have at least 2 interactions with other selected tiles.
   - Single connected group only: selected tiles must form one connected component (no multiple closed groups).

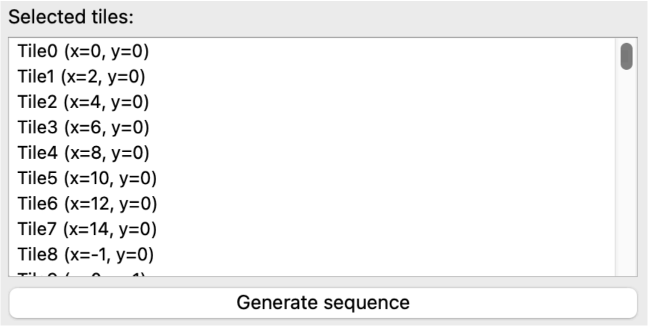

5. Generate sequence
   Click `Generate sequence` to run sequence generation and NUPACK design for selected tiles. Runtime depends on lattice size; for the 107-tile case it is around 5 minutes.

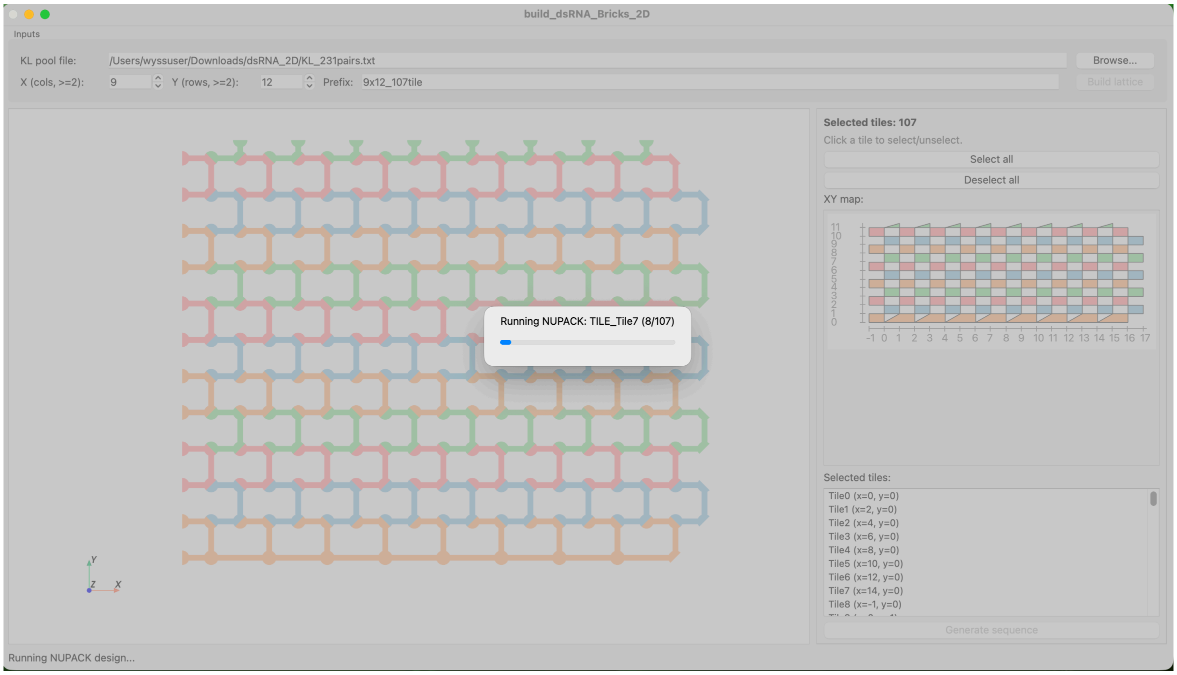

#### Output Example (9x12_107tile)

1. 2D map
   `demo_9x12_107tile/9x12_107tile_2D_map.svg` shows KL pair assignment on all tiles and layers.

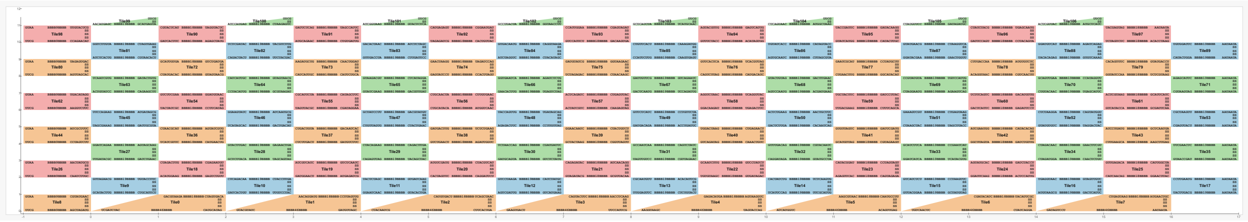

2. Tile structure output
   `demo_9x12_107tile/9x12_107tile_tile_structure.txt` contains tile scaffold sequences (with N placeholders) and designed dot-bracket secondary structures after KL pair assignment.

```
~~~~~~~~~~~~~~~~~~~~~~~~~~~~~~~~~~~~~~~~~~~~~~~~~~~~~~~~~~~~~~~~~~~~~~~~~~~~~~~~

*******TILE_Tile0*******

GNNNNNNUNNNNNNNA CUAGUGACA NNNNNNNUNNNNNNNNGNC AAA GACGUAAGA
GNUNNNNNNNGNNNNNNN ANNNNNNGNNNNNNC GNNNNNNGNNNA CAUGCAUAG
NNNNNNNUNNNNNNNNUNNNNNNNGNNNNNNNUNNNNNNNNGNNNNNG AAA UCGAUCUAC
CNNNNNUNNNNNNNGNNNNNNNUNNNNNNNGNNNNNNNGNNNNNNN ANNNUNNNNNNC UU
CACGAAGUCAAUAC

   ((((((((((((((. .........
(((((((((((((((((( ... ......... )))))))))))))))))) .))))))))))))))
(((((((((((. .........
(((((((((((((((((((((((((((((((((((((((((((((( ... ......... )))))))))
))))))))))))))))))))))))))))))))))))) .))))))))))) .. ..............

~~~~~~~~~~~~~~~~~~~~~~~~~~~~~~~~~~~~~~~~~~~~~~~~~~~~~~~~~~~~~~~~~~~~~~~~~~~~~~~~
```

3. Final sequence output
   `demo_9x12_107tile/9x12_107tile_sequence.txt` contains, for each tile: input scaffold sequence -> input dot-bracket secondary structure -> NUPACK-designed full sequence.

```
~~~~~~~~~~~~~~~~~~~~~~~~~~~~~~~~~~~~~~~~~~~~~~~~~~~~~~~~~~~~~~~~~~~~~~~~~~~~~~~~

*******TILE_Tile0*******

GNNNNNNUNNNNNNNA CUAGUGACA NNNNNNNUNNNNNNNNGNC AAA GACGUAAGA
GNUNNNNNNNGNNNNNNN ANNNNNNGNNNNNNC GNNNNNNGNNNA CAUGCAUAG
NNNNNNNUNNNNNNNNUNNNNNNNGNNNNNNNUNNNNNNNNGNNNNNG AAA UCGAUCUAC
CNNNNNUNNNNNNNGNNNNNNNUNNNNNNNGNNNNNNNGNNNNNNN ANNNUNNNNNNC UU
CACGAAGUCAAUAC

((((((((((((((. .........
(((((((((((((((((( ... ......... )))))))))))))))))) .))))))))))))))
(((((((((((. .........
(((((((((((((((((((((((((((((((((((((((((((((( ... ......... )))))))))
))))))))))))))))))))))))))))))))))))) .))))))))))) .. ..............

GCCGGACUGCCUGCA CUAGUGACA GCCGAGCUGAUCGUGGGC AAA GACGUAAGA
GCUCGCGAUCGGCUCGGC AGCAGGCGGUCUGGC GGUAGGUGCGCA CAUGCAUAG
CCCGUUCUGCCUAGCUGGAGUCGGUGCGAUAUGCGGUGCGGCUCCG AAA UCGAUCUAC
```

CGGAGCUGCACCGCGUGUCGCAUCGACUCCGGCUAGGCGGAGCGGG AGCGUACCUACC UU CACGAAGUCAAUAC

~~~~~~~~~~~~~~~~~~~~~~~~~~~~~~~~~~~~~~~~~~~~~~~~~~~~~~~~~~~~~~~~~~~~~~~~~~~~~~~~

**Sequence Generation Notes**

- Generation uses selected tiles only.
- Validation checks include KL pair pool availability, structure closure, and selected-tile connectivity.
- `Generate sequence` exports the three output files listed above and runs NUPACK with a progress dialog.

**NUPACK Design Principles**

- RNA model with some-nupack3 ensemble, at 37°C and 1.0 M sodium.
- Soft constrains (weight 1.0) for patterns: A4, C4, G4, U4, K6, M6, R6, S6, W6, Y6.
- Sequence optimization stop condition: f_stop = 0.02.
- Each tile design runs 3 rounds (trials), and the best result is selected.

**Reproducibility**

- KL pair assignment is randomized with a fixed seed (42) to make runs reproducible.
- With the same KL pair pool file, lattice size (X, Y), and tile selection, the assignment/order is deterministic.

**File Format Mini-Spec**

1. KL pair pool file (`KL_231pairs.txt` style)
   - One pair per line.
   - Two RNA sequences separated by a tab.
   - Example:

~~~~~~~~~~~~~~~~~~~~~~~~~~~~~~~~~~~~~~~~~~~~~~~~~~~~~~~~~~~~~~~~~~~~~~~~~~~~~~~~

CUAGAUGGA   GAUCUACCU

UGUACCUUC   ACAUGGAAG

~~~~~~~~~~~~~~~~~~~~~~~~~~~~~~~~~~~~~~~~~~~~~~~~~~~~~~~~~~~~~~~~~~~~~~~~~~~~~~~~

2. Tile structure output (`<prefix>_tile_structure.txt`)
   - Repeated tile blocks in this order:
     - tile header line: `*******TILE_NAME*******`
     - tile scaffold sequence (with N placeholders and spacing groups)
~~~~~~~~~~~~~~~~~~~~~~~~~~~~~~~~~~~~~~~~~~~~~~~~~~~~~~~~~~~~~~~~~~~~~~~~~~~~~~~~

- dot-bracket secondary structure (with spacing groups)

3. Final sequence output (`<prefix>_sequence.txt`)
   - Repeated tile blocks in this order:
     - tile header line
     - input scaffold sequence
     - input dot-bracket secondary structure
     - NUPACK-designed final full sequence

**Troubleshooting**

1. Qt window does not show or hangs on startup
   - Confirm PySide6 version:
     - recommended: 6.9.*
     - Check version:

```
bash python -c "import PySide6; print(PySide6.__version__)"
```

2. 3D rendering/picking issues
   - Reinstall GUI/render stack in the active env:

```
bash pip install --upgrade "pyside6==6.9.*" pyvista pyvistaqt vtk
```

3. NUPACK step fails
   - Verify NUPACK installation in the same Python environment used to run the GUI:

```
bash python -c "import nupack; print('nupack ok')"
```

- If import fails, follow the official setup guide: NUPACK Documentation

#### III. dsRNA Bricks 3D Builder

##### Outline

This project is a Python GUI tool for designing and exporting 3D dsRNA brick lattices. It builds RNA nanostructures from predefined tile modules, visualizes them in 3D and 2D, and generates sequence-design outputs for downstream optimization.

The workflow is:

1. Load a KL pair pool file.
2. Set lattice dimensions (X, Y, Z) and a prefix.
3. Build the 3D lattice.
4. Inspect and select tiles in synchronized 3D and 2D views.
5. Generate outputs for structure review and sequence design.

Current outputs:

- <prefix>_tile_structure.txt
- <prefix>_2D_map.svg
- <prefix>_sequence.txt

##### Requirements

Python:

- Python 3.12 (tested)

Core packages:

- PySide6==6.9.* (recommended)
- numpy
- pyvista
- pyvistaqt
- vtk (tested with 9.5.2)

Sequence design package:

- nupack (install/configure by following the official documentation: https://docs.nupack.org/)

Installation demo (conda + pip):

```
conda create -n dsrna python=3.12 -y

conda activate dsrna

pip install "pyside6==6.9.*" numpy pyvista pyvistaqt vtk
```

Optional quick check:

~~~~~~~~~~~~~~~~~~~~~~~~~~~~~~~~~~~~~~~~~~~~~~~~~~~~~~~~~~~~~~~~~~~~~~~~~~~~~~~~~~~~~~~~~~~~~~~~~~~~~

```
python -c "import PySide6, pyvista, pyvistaqt, vtk, numpy; print('PySide6', PySide6.__version__); print('PyVista', pyvista.__version__); print('VTK', vtk.vtkVersion.GetVTKVersion()); print('NumPy', numpy.__version__)"
```

~~~~~~~~~~~~~~~~~~~~~~~~~~~~~~~~~~~~~~~~~~~~~~~~~~~~~~~~~~~~~~~~~~~~~~~~~~~~~~~~~~~~~~~~~~~~~~~~~~~~~

For NUPACK setup and installation, follow: [NUPACK Documentation].

**Run**

From this folder:

~~~~~~~~~~~~~~~~~~~~~~~~~~~~~~~~~~~~~~~~~~~~~~~~~~~~~~~~~~~~~~~~~~~~~~~~~~~~~~~~~~~~~~~~~~~~~~~~~~~~~

```
python build_dsRNA_Bricks_3D.py
```

~~~~~~~~~~~~~~~~~~~~~~~~~~~~~~~~~~~~~~~~~~~~~~~~~~~~~~~~~~~~~~~~~~~~~~~~~~~~~~~~~~~~~~~~~~~~~~~~~~~~~

Alternative:

~~~~~~~~~~~~~~~~~~~~~~~~~~~~~~~~~~~~~~~~~~~~~~~~~~~~~~~~~~~~~~~~~~~~~~~~~~~~~~~~~~~~~~~~~~~~~~~~~~~~~

```
python -m function.build_dsRNA_Bricks_3D
```

~~~~~~~~~~~~~~~~~~~~~~~~~~~~~~~~~~~~~~~~~~~~~~~~~~~~~~~~~~~~~~~~~~~~~~~~~~~~~~~~~~~~~~~~~~~~~~~~~~~~~

**Project Layout**

- `build_dsRNA_Bricks_3D.py`: top-level launcher.
- `function/app.py`: main GUI window and workflow wiring.
- `function/lattice_builder.py`: lattice/tile placement rules.
- `function/map2d.py`: 2D XY layer map widgets and selection.
- `function/view3d.py`: embedded PyVista 3D renderer.
- `function/c_tiles.py`: tile geometry definitions.
- `function/rna_tile_generator.py`: Type I / Type II RNA scaffold generation.
- `function/nupack_runner.py`: NUPACK run pipeline for generated tiles.
- `KL_231pairs.txt`: example KL pair pool file.
- `demo_3x5x4_118tile/`: demo run outputs for a 3x5x4 lattice.

**Workflow Example (3x5x4, 118 tiles)**

6. Overview
   This example builds a 3x5x4 lattice (118 tiles total).

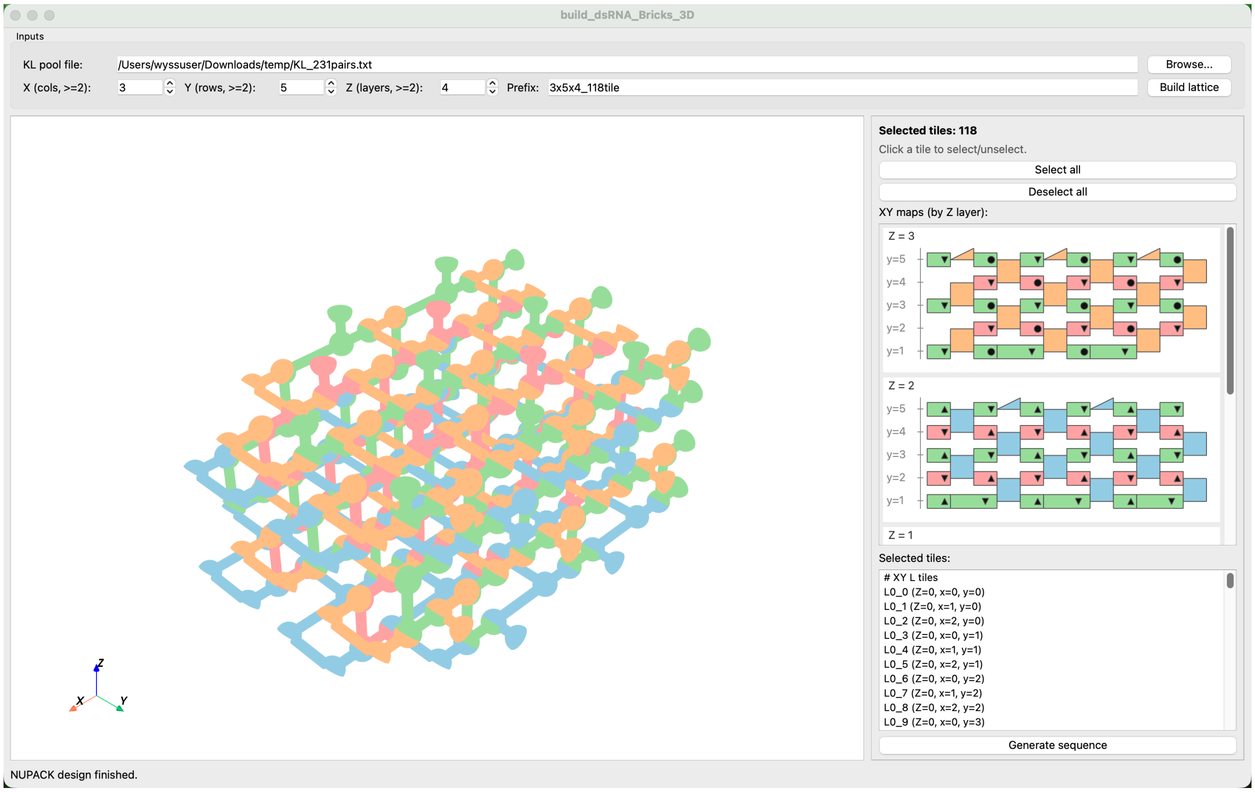

7. Input
   Choose a KL pair pool file. Here we use `KL_231pairs.txt`.

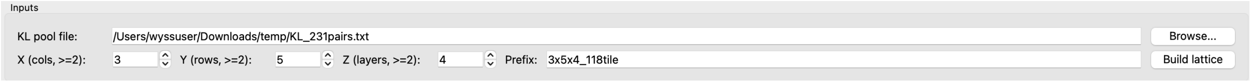

Example lines from `KL_231pairs.txt`:

~~~~~~~~~~~~~~~~~~~~~~~~~~~~~~~~~~~~~~~~~~~~~~~~~~~~~~~~~~~~~~~~~~~~~~~~~~~~~~~~~~~~~~~~~~~~~~

```
CUAGAUGGA   GAUCUACCU
UGUACCUUC   ACAUGGAAG
UCAGAUUCG   AGUCUAAGC
GCAUGAGUA   CGUACUCAU
GCUCAUCUA   CGAGUAGAU
UAGCCAUUC   AUCGGUAAG
GUUAGAACG   CAAUCUUGC
CCUUAGUUG   GGAAUCAAC
```

~~~~~~~~~~~~~~~~~~~~~~~~~~~~~~~~~~~~~~~~~~~~~~~~~~~~~~~~~~~~~~~~~~~~~~~~~~~~~~~~~~~~~~~~~~~~~~

Set X=3, Y=5, Z=4, and optional prefix.
The required KL pair count for the selected lattice must be smaller than or equal to the pool size.

8. Build the lattice
   Click `Build lattice`. The GUI generates both the 3D structure and the layered 2D map.

3D self-assembly view:

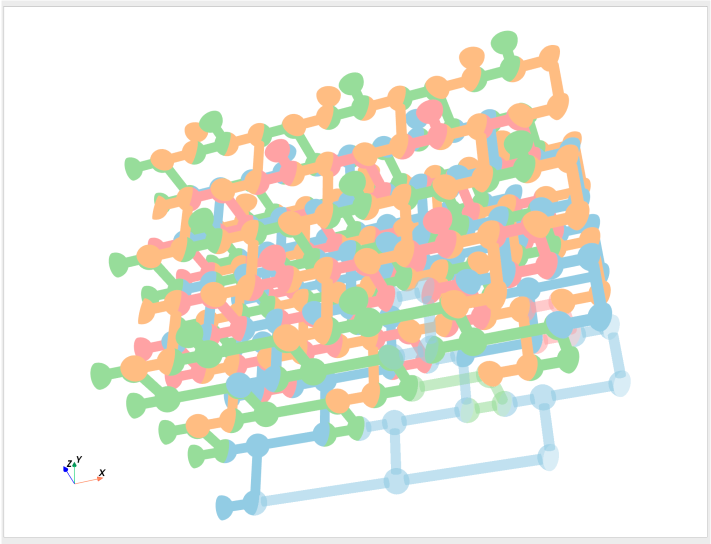

2D layered map:

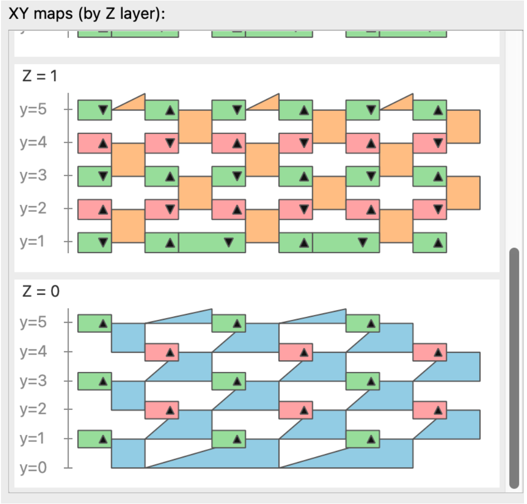

9. Select tiles
   Select/deselect from either the 3D view or 2D map. Selection is synchronized and shown in the right-side list. Selection rules before sequence generation:

- No flexible tiles: each selected tile must have at least 2 interactions with other selected tiles.
- Single connected group only: selected tiles must form one connected component (no multiple closed groups).

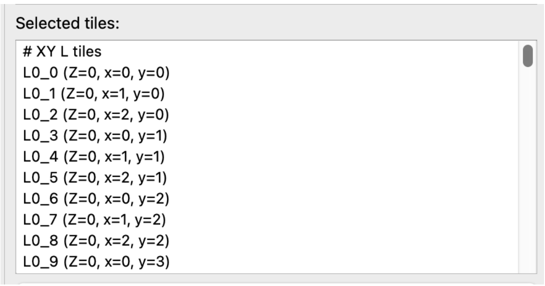

10. Generate sequence
    Click Generate sequence to run sequence generation and NUPACK design for selected tiles. Runtime depends on lattice size; for the 118-tile case it is around 5 minutes.

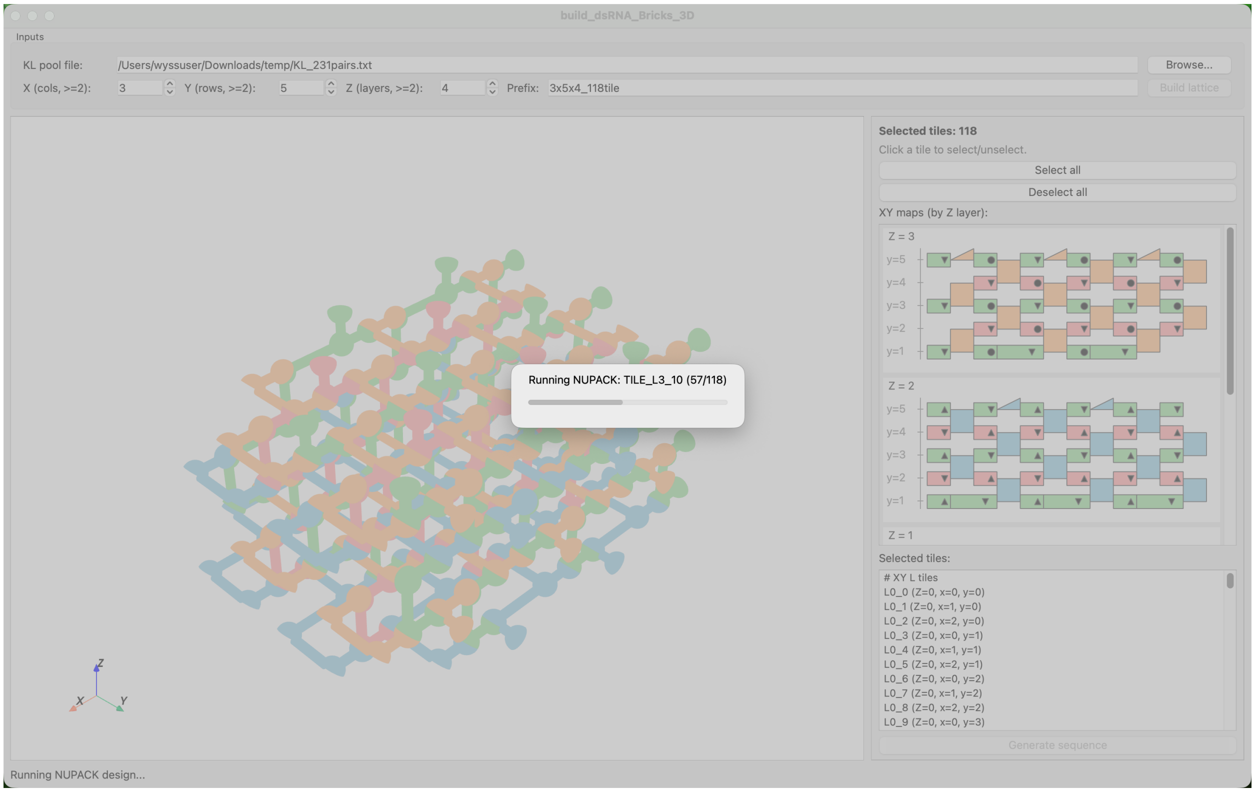

**Output Example (3x5x4_118tile)**

4. 2D map
   `demo_3x5x4_118tile/3x5x4_118tile_2D_map.svg` shows KL pair assignment on all tiles and layers.

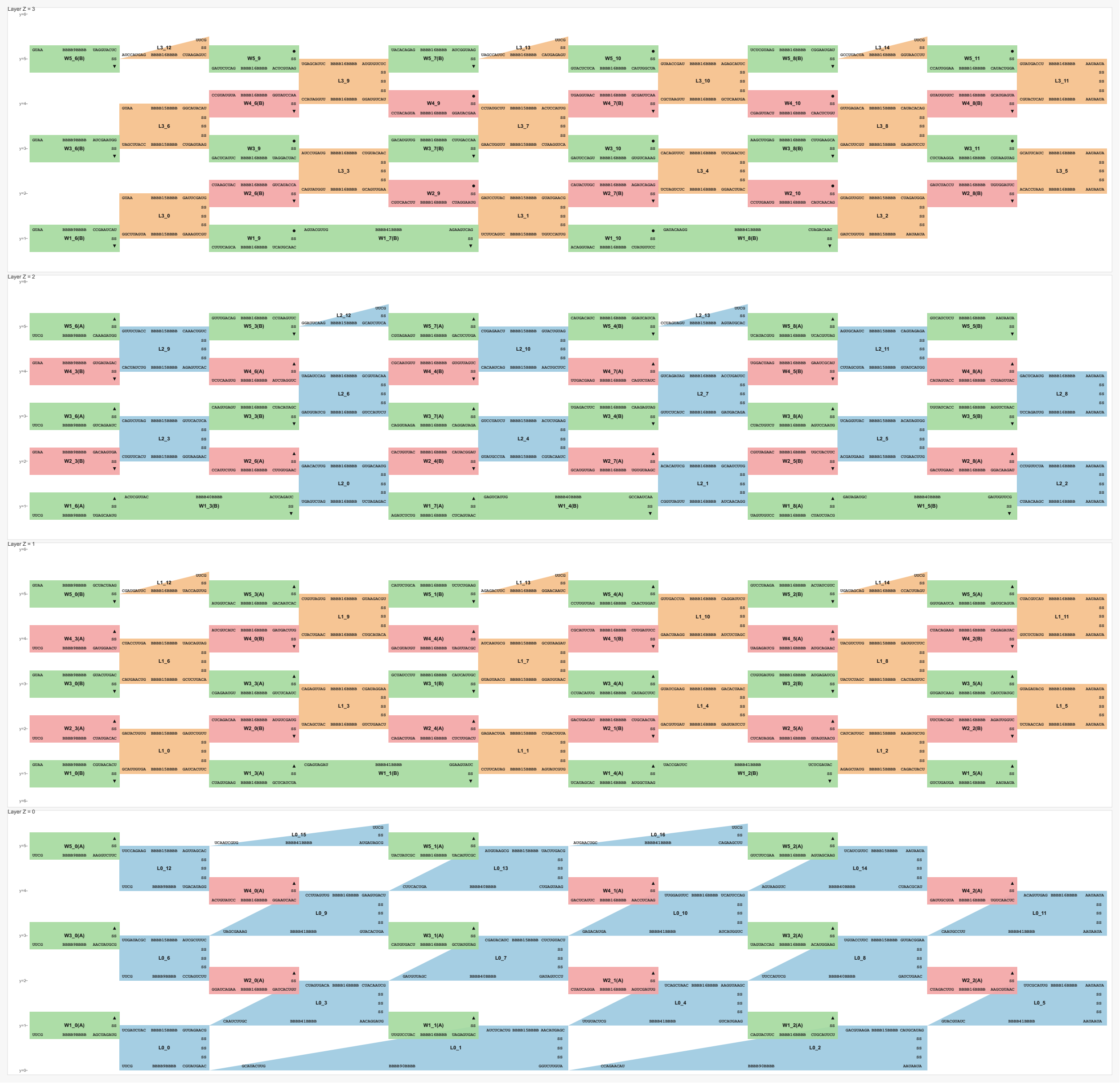

5. Tile structure output `demo_3x5x4_118tile/3x5x4_118tile_tile_structure.txt` contains tile scaffold sequences (with N placeholders) and designed dot-bracket secondary structures after KL pair assignment.

~~~~~~~~~~~~~~~~~~~~~~~~~~~~~~~~~~~~~~~~~~~~~~~~~~~~~~~~~~~~~~~~~~~~~~~~~~~~~~~~~~~~~~~~~~~~~~~~~~~~

```
*******TILE_L0_0*******

GNNNNNNNUNNNNNNNA GUUAGAACG NNNNNNNNUNNNNNNNC AAA UCGAUCUAC
GNNNNNNNGNNNNNNNN ANNNNNNNGNNNNNNNC GNNNNNNNUNNNA CGUAUGAAC NNNNNNNNC UUCG
GNNNNNNNN NNNNGNNNNNNNC UU CACGAAGUCAAUAC
```
~~~~~~~~~~~~~~~~~~~~~~~~~~~~~~~~~~~~~~~~~~~~~~~~~~~~~~~~~~~~~~~~~~~~~~~~~~~~~~~~~~~~~~~~~~~~~~~~~~~~

```
((((((((((((((. .........
((((((((((((((( ... ......... ))))))))))))))) .))))))))))))))
(((((((((((. .........
((((((((( .... ))))))))) .))))))))))) .. ..............
```

~~~~~~~~~~~~~~~~~~~~~~~~~~~~~~~~~~~~~~~~~~~~~~~~~~~~~~~~~~~~~~~~~~~~~~~~~~~~~~~~~~~~~~~~~~~~~~~~~~~~~~~~~~~~~~~~~~~~~~~~~~~~~~~~~~~~~~~

6. Final sequence output
   `demo_3x5x4_118tile/3x5x4_118tile_sequence.txt` contains, for each tile: `input scaffold sequence -> input dot-bracket secondary structure -> NUPACK-designed full sequence`.

~~~~~~~~~~~~~~~~~~~~~~~~~~~~~~~~~~~~~~~~~~~~~~~~~~~~~~~~~~~~~~~~~~~~~~~~~~~~~~~~~~~~~~~~~~~~~~~~~~~~~~~~~~~~~~~~~~~~~~~~~~~~~~~~~~~~~~~

```
*******TILE_L0_0*******

GNNNNNNUNNNNNNA GUUAGAACG NNNNNNNUNNNNNNNC AAA UCGAUCUAC
GNNNNNNGNNNNNNN ANNNNNNGNNNNNNC GNNNNNNUNNNA CGUAUGAAC NNNNNNNNC UUCG
GNNNNNNNN NNNNGNNNNNNC UU CACGAAGUCAAUAC

((((((((((((((. .........
((((((((((((((( ... ......... ))))))))))))))) .))))))))))))))
(((((((((((. .........
((((((((( .... ))))))))) .))))))))))) .. ..............

GGCGCUCUGCUGCCA GUUAGAACG GACCUGCUGCUCCGC AAA UCGAUCUAC
GCGGAGCGGCAGGUC AGGCAGCGGAGUGCC GCUGCGGUAUCA CGUAUGAAC GGCUCAGGC UUCG
GCCUGGGCC AGAUGCCGCAGC UU CACGAAGUCAAUAC
```

~~~~~~~~~~~~~~~~~~~~~~~~~~~~~~~~~~~~~~~~~~~~~~~~~~~~~~~~~~~~~~~~~~~~~~~~~~~~~~~~~~~~~~~~~~~~~~~~~~~~~~~~~~~~~~~~~~~~~~~~~~~~~~~~~~~~~~~

**Sequence Generation Notes**

- Generation uses selected tiles only.
- Validation checks include KL pair pool availability, structure closure, and selected-tile connectivity.
- `Generate sequence` exports the three output files listed above and runs NUPACK with a progress dialog.

**NUPACK Design Principles**

- RNA model with some-nupack3 ensemble, at 37°C and 1.0 M sodium.
- Soft constrains (weight 1.0) for patterns: A4, C4, G4, U4, K6, M6, R6, S6, W6, Y6.
- Sequence optimization stop condition: f_stop = 0.02.
- Each tile design runs 3 rounds (trials), and the best result is selected.

**Reproducibility**
~~~~~~~~~~~~~~~~~~~~~~~~~~~~~~~~~~~~~~~~~~~~~~~~~~~~~~~~~~~~~~~~~~~~~~~~~~~~~~~~~~~~~~~~~~~~~~~~~~~~~~~~~~~~~~~~~~~~~~~~~~~~~~~~~~~~~~~

- KL pair assignment is randomized with a fixed seed (42) to make runs reproducible.
- With the same KL pair pool file, lattice size (X, Y, Z), and tile selection, the assignment/order is deterministic.

**File Format Mini-Spec**

4. KL pair pool file (KL_231pairs.txt style)
   - One pair per line.
   - Two RNA sequences separated by a tab.
   - Example:

~~~
CUAGAUGGA   GAUCUACCU

UGUACCUUC   ACAUGGAAG
~~~

5. Tile structure output (<prefix>_tile_structure.txt)
   - Repeated tile blocks in this order:
     - tile header line: *******TILE_NAME*******
     - tile scaffold sequence (with N placeholders and spacing groups)
     - dot-bracket secondary structure (with spacing groups)

6. Final sequence output (<prefix>_sequence.txt)
   - Repeated tile blocks in this order:
     - tile header line
     - input scaffold sequence
     - input dot-bracket secondary structure
     - NUPACK-designed final full sequence

**Troubleshooting**

4. Qt window does not show or hangs on startup
   - Confirm PySide6 version:
     - recommended: 6.9.*
     - Check version:

~~~
bash python -c "import PySide6; print(PySide6.__version__)"
~~~

5. 3D rendering/picking issues
   - Reinstall GUI/render stack in the active env:

~~~~~~~~~~~~~~~~~~~~~~~~~~~~~~~~~~~~~~~~~~~~~~~~~~~~~~~~~~~~~~~~~~~~~~~~~~~~~~~~~~~~~~~~~~~~~~~~~~~~

```
bash pip install --upgrade "pyside6==6.9.*" pyvista pyvistaqt vtk
```

~~~~~~~~~~~~~~~~~~~~~~~~~~~~~~~~~~~~~~~~~~~~~~~~~~~~~~~~~~~~~~~~~~~~~~~~~~~~~~~~~~~~~~~~~~~~~~~~~~~~

6. NUPACK step fails
   - Verify NUPACK installation in the same Python environment used to run the GUI:

~~~~~~~~~~~~~~~~~~~~~~~~~~~~~~~~~~~~~~~~~~~~~~~~~~~~~~~~~~~~~~~~~~~~~~~~~~~~~~~~~~~~~~~~~~~~~~~~~~~~

```
bash python -c "import nupack; print('nupack ok')"
```

~~~~~~~~~~~~~~~~~~~~~~~~~~~~~~~~~~~~~~~~~~~~~~~~~~~~~~~~~~~~~~~~~~~~~~~~~~~~~~~~~~~~~~~~~~~~~~~~~~~~

- If import fails, follow the official setup guide: NUPACK Documentation

#### Sequences

Color key: red = kissing-loop (KL); purple = boundary tetraloop / bulge; blue = tail; orange = reverse primer / binding site; green = T7 promoter

##### 1) Short DNA strands

| Name (Note) | Sequence | length |
|---|---|---|
| **RH16** | GTATmUmGmAmCmUmUmCmGTGCC | 16 |
| **RH14** | GTATmUmGmAmCmUmUmCmGTG | 14 |
| **SH14** | GTATTGACTTCGTG | 14 |
| **SH12** | GTATTGACTTCG | 12 |
| **SH10** | ATTGACTTCG | 10 |
| **CL35.FWD**<br>Forward primer | **GTTCTAATACGACTCACTATA**GGAT | 25 |
| **CL13RK.REV**<br>Reverse primer | **mCmGTCGTAAGCATGAAGGTTGGAC** | 23 |
| **F15H-REV2**<br>Reverse primer | **GCCTGCGTTGCCCTGGTG** | 18 |

##### 2) dsRNA bricks

| Name (Note) | Sequence | length |
|---|---|---|
| **R4B-Brick1** | GGGCAGGC **CAGCGC** A GGUCUACGG AA **GCUGGC** A CCGUGGACC GCCUGUCC GCUCGGUC **GCGUGC** A CCAGCCUGUUUAGGAGUAUUCCUAG AA **GCACGC** A CUAGGAGUACUCUUAAACGGGCUGG GACCGAGC **UU UUAU CACGAAGUCAAUAC** | 152 |
| **R4B-Brick2** | CCAUCGCU **GACCCG** A UGACGGGUG AA **GGACCG** A CACCUGUCA AGCGAUGG CCUCGGUC **GCCAGC** A UACGGGUG AA **GCGCUG** A CACCUGUA GACCGAGG **UU UUAA CACGAAGUCAAUAC** | 118 |
| **R4B-Brick3** | GCACUGGU **CUGCGC** A GAUGUCAGG AA **CCAGCG** A CCUGGCAUC ACCAGUGC GCUCGCGU **CGGUCC** A UACUCUGG AA **CGGGUC** A CCAGGGUA ACGCGAGC **UU UUAC CACGAAGUCAAUAC** | 118 |
| **R4B-Brick4** | CCUCGACC **UUCG** GGUCGAGG GCCUGCGU **CGCUGG** A UCCGAGUG AA **GCGCAG** A CACUUGGA ACGCAGGC **UU UAUU CACGAAGUCAAUAC** | 88 |
| **R6B-Brick1** | GGGCAGGC **CAGCGC** A GGUCUACGG AA **GCUGGC** A CCGUGGACC GCCUGUCC GCUCGGUC **GCGUGC** A CCAGCCUGUUUAGGAGUAUUCCUAG AA **GCACGC** A CUAGGAGUACUCUUAAACGGGCUGG GACCGAGC **UU UUAU CACGAAGUCAAUAC** | 152 |
| **R6B-Brick2** | CCAUCGCU **GACCCG** A UGACGGGUG AA **GGACCG** A CACCUGUCA AGCGAUGG CCUCGGUC **GCCAGC** A UACGGGUG AA **GCGCUG** A CACCUGUA GACCGAGG **UU UUAA CACGAAGUCAAUAC** | 118 |
| **R6B-Brick3** | GCACUGGU **CUGCGC** A GAUGUCAGG AA **CCAGCG** A CCUGGCAUC ACCAGUGC GCUCGCGU **CGGUCC** A UACUCUGG AA **CGGGUC** A CCAGGGUA ACGCGAGC **UU UUAC CACGAAGUCAAUAC** | 118 |
| **R6B-Brick4** | CCAGCAGU **GAGGCC** A UGACGCGUG AA **GGUGGC** A CACGUGUCA ACUGCUGG CUCCCGCU **CGCUGG** A UCCGGCAG AA **GCGCAG** A CUGCUGGA AGCGGGAG **UU UAUU CACGAAGUCAAUAC** | 118 |
| **R6B-Brick5** | GCAGGCGU **GUCGCG** A CCACUGCAG AA **GCACCC** A CUGCGGUGG ACGCCUGC GCAGGAGC **GCCACC** A UUCUGUCG AA **GGCCUC** A CGACGGAA GCUCCUGC **UU UAAU CACGAAGUCAAUAC** | 118 |
| **R6B-Brick6** | CCGUCGUC **UUCG** GACGACGG GCUCCCGA **GGGUGC** A UCGGUCUG AA **CGCGAC** A CAGAUCGA UCGGGAGC **UU UACU CACGAAGUCAAUAC** | 88 |
| **R8B-Brick1** | GGGCAGGC **CAGCGC** A GGUCUACGG AA **GCUGGC** A CCGUGGACC GCCUGUCC GCUCGGUC **GCGUGC** A CCAGCCUGUUUAGGAGUAUUCCUAG AA **GCACGC** A CUAGGAGUACUCUUAAACGGGCUGG GACCGAGC **UU UUAU CACGAAGUCAAUAC** | 152 |

| Name (Note) | Sequence | length |
|---|---|---|
| R8B-Brick2 | CCAUCGCU **GACCCG** A UGACGGGUG AA **GGACCG** A CACCUGUCA AGCGAUGG CCUCGGUC **GCCAGC** A UACGGGUG AA **GCGCUG** A CACCUGUA GACCGAGG **UU UUAA CACGAAGUCAAUAC** | 118 |
| R8B-Brick3 | GCACUGGU **CUGCGC** A GAUGUCAGG AA **CCAGCG** A CCUGGCAUC ACCAGUGC GCUCGCGU **CGGUCC** A UACUCUGG AA **CGGGUC** A CCAGGGUA ACGCGAGC **UU UUAC CACGAAGUCAAUAC** | 118 |
| R8B-Brick4 | CCAGCAGU **GAGGCC** A UGACGCGUG AA **GGUGGC** A CACGUGUCA ACUGCUGG CUCCCGCU **CGCUGG** A UCCGGCAG AA **GCGCAG** A CUGCUGGA AGCGGGAG **UU UAUU CACGAAGUCAAUAC** | 118 |
| R8B-Brick5 | GCAGGCGU **GUCGCG** A CCACUGCAG AA **GCACCC** A CUGCGGUGG ACGCCUGC GCAGGAGC **GCCACC** A UUCUGUCG AA **GGCCUC** A CGACGGAA GCUCCUGC **UU UAAU CACGAAGUCAAUAC** | 118 |
| R8B-Brick6 | CCAGUGCU **CUCGCC** A GGGUGCCUG AA **GGAGCC** A CAGGUACCC AGCACUGG CCAGCCGA **GGGUGC** A UCCGGUAG AA **CGCGAC** A CUACUGGA UCGGCUGG **UU UACU CACGAAGUCAAUAC** | 118 |
| R8B-Brick7 | GCCAGGCU **GUGCCG** A UCAGUACGG AA **CGUCGC** A CCGUGCUGA AGCCUGGC GCAGGGUC **GGCUCC** A UCAUAGUG AA **GGCGAG** A CACUGUGA GACCCUGC **UU AUUU CACGAAGUCAAUAC** | 118 |
| R8B-Brick8 | CCUCGGUC **UUCG** GACCGAGG GCUCGGGU **GCGACG** A CCCGGUAG AA **CGGCAC** A CUACUGGG ACCCGAGC **UU AAUU CACGAAGUCAAUAC** | 88 |
| R4×3-Brick1 | GGGCAGGC **CAGCGC** A GGACUACGG AA **GCUGGC** A CCGUGGUCC GCCUGUCC GCUCGAGG **GCGUGC** A UGAGUGUCUGUAGAGUACUGAGCGG AA **CGUCCG** A CCGCUCGGUACUUUACAGGCACUCA CCUCGAGC **UU UUAU CACGAAGUCAAUAC** | 152 |
| R4×3-Brick2 | CCAGCAGG **GACCCG** A UCUCGGGUG AA **GGACCG** A CACCUGAGA CCUGCUGG CCCUCGCU **GCCUGG** A CUCGACGG AA **GCGCUG** A CCGUUGAG AGCGAGGG **UU UUAA CACGAAGUCAAUAC** | 118 |
| R4×3-Brick3 | GCCUCUGU **CUGCGC** A GGAGUACGG AA **CCAGCG** A CCGUGCUCC ACAGAGGC GCUCUUGU **CGGUCC** A GCUUCGGG AA **CGGGAG** A CCCGGAGC ACAAGAGC **UU UUAC CACGAAGUCAAUAC** | 118 |
| R4×3-Brick4 | CCAGCAGC **UUCG** GCUGCUGG GCCUCUGC **CGGUGG** A UGGGCCAG AA **GCGCAG** A CUGGUCCA GCAGAGGC **UU UAUU CACGAAGUCAAUAC** | 88 |
| R4×3-Brick5 | GCCUCGGC **GUGCCC** A CUCAUACGG AA **CCAGGC** A CCGUGUGAG GCCGAGGC GCUCCGUC **GGGACC** A UCGUUGUAUGCUGCAACUUGGCUGG AA **GCACGC** A CCAGCCGAGUUGUAGCAUGCAACGA GACGGAGC **UU UAAU CACGAAGUCAAUAC** | 152 |
| R4×3-Brick6 | CCAGCUGC **CUCGGG** A GCGUGCGGG AA **CCUCGG** A CCCGUACGC GCAGCUGG CCCUGCUG **CGCUCG** A UCCGGUCG AA **GGGCAC** A CGACUGGA CAGCAGGG **UU UACU CACGAAGUCAAUAC** | 118 |
| R4×3-Brick7 | GCUCGAGC **CACCGG** A GCUGUAGUG AA **CCACCG** A CACUGCAGC GCUCGAGC GCCUGCGU **CCGAGG** A UGCUCUGG AA **CGGGUC** A CCAGGGCA ACGCAGGC **UU AUUU CACGAAGUCAAUAC** | 118 |
| R4×3-Brick8 | GGGCCUGC **CAGCCG** A GGGUUCCGG AA **CGAGCG** A CCGGGACCC GCAGGUCC GCUCCAGC **CGGACG** A GCUCAAUUGACCGUUGACUGGAGCG AA **GGUCCC** A CGCUCCGGUCAAUGGUCAGUUGAGC GCUGGAGC **UU UUAU CACGAAGUCAAUAC** | 152 |
| R4×3-Brick9 | CCAGUGAG **CUCCCG** A GGCGGACGG AA **CGUGGG** A CCGUUCGCC CUCACUGG CCCUGGUC **GCCAGC** A CGCGGUGG AA **CGGCUG** A CCACUGCG GACCAGGG **UU UUAA CACGAAGUCAAUAC** | 118 |
| R4×3-Brick10 | GGGAGUCC **CUGGCG** A GCUCUGUGG AA **CGACGG** A CCACGGAGC GGACUCCC GCUCAGGU **CCCACG** A GGCUUCUG AA **CCCGAG** A CAGAGGCC ACCUGAGC **UU UUAC CACGAAGUCAAUAC** | 118 |
| R4×3-Brick11 | CCUCACGC **UUCG** GCGUGAGG GCCUGUCUUUAUGGGCAGAACG **CGCUGG** A UCCGCAGG AA **CGCCAG** A CCUGUGGA CGUUCUGUCCAUAAGGACAGGC **UU UAUU CACGAAGUCAAUAC** | 116 |
| R4×3-Brick12 | CCAGAGCC **UUCG** GGCUCUGG GCAGGCGUUCUGUGC **CCGUCG** A GGCGGUGG AA **CCGGUG** A CCACUGCC GCACAGAGCGCCUGC **UU UAAU CACGAAGUCAAUAC** | 102 |
| A4B-Brick1 | GGGCAUCC **CAGCGC** A GGGAUCUAG AA **GCUGGC** A CUAGGUCCC GGAUGCCC GACCGGUC **GCGUGC** A UGGAACUUACCGUG AA **GCACGC** A CACGGUAGGUUCCA GACCGGUC **UU UUAA CACGAAGUCAAUAC** | 130 |

| Name (Note) | Sequence | length |
|---|---|---|
| **A4B-Brick2** | CCUCGCGU **GACCCG** A UGUCUAGUGACGAUGGCGUG AA **GGACCG** A<br>CACGCCGUCGUCAUUAGACA ACGCGAGG CCAGGGUC **GCCAGC** A UACGGGUG AA **GCGCUG** A<br>CACCUGUA GACCCUGG **UU UUAC CACGAAGUCAAUAC** | 140 |
| **A4B-Brick3** | GCAGGCUC **CUGCGC** A GAGGCUUAGCCUAGCCUGGG AA **CCAGCG** A<br>CCCAGGUUAGGCUGAGCCUC GAGCCUGC GCUAGCGU **CGGUCC** A GCGUCUGG AA **CGGGUC** A<br>CCAGGCGC ACGCUAGC **UU UAUU CACGAAGUCAAUAC** | 140 |
| **A4B-Brick4** | CCAGGACC **UUCG** GGUCCUGG GCCUGCCU **CGCUGG** A ACGCGUGGACAGUUGAGUG AA<br>**GCGCAG** A CACUCAGCUGUCUACGCGU AGGCAGGC **UU UAAU CACGAAGUCAAUAC** | 110 |
| **QT5B-Brick1** | GGGCCUAC **GCGCAC** A<br>UGUCCCGUUGCUCAGGAAUUGAUCGGACUAGAUGGCGAUUUGUAUCGGUGUAAUCCUCGGAGGCUUCU<br>G AA **GUGCGC** A<br>CAGAAGCUUCCGAGGGUUACACUGAUACAAGUCGCCAUUUAGUCCGGUCAAUUUCUGAGCAGCGGGAC<br>A GUAGGCUC GCGACCGC **CACGGG** A UGCUGUAUCUG AA **CUCGCC** A CAGAUGCAGCA<br>GCGGUCGC **UU AU CACGAAGUCAAUAC** | 242 |
| **QT5B-Brick2** | CCGUCCUG **CGAGCG** A GCCGUCCUAGCACUGCUCAUAUCGUGAGGACUGCAUGCGUCGGUGACAUG<br>AA **CUGCCG** A CAUGUCAUCGACGCGUGCAGUUCUCACGGUAUGAGUAGUGCUGGGACGGC<br>CAGGACGG CCAGGGUC **GGCGAG** A CCAGCGGCUGG AA **GUCCCG** A CCAGCUGCUGG<br>GACCCUGG **UU AA CACGAAGUCAAUAC** | 204 |
| **QT5B-Brick3** | GCAGGCGU **GCGACC** A UCCGUAUGGAGUGAUGCUUCUGGCAUCAUUCCUACGGAUCUCUGUACAGG<br>AA **CGCCUC** A CCUGUACGGAGAUCUGUAGGAGUGAUGCUAGAAGCGUCACUCUAUACGGA<br>ACGCCUGC GCUCGGAC **CGGCAG** A UCUCGUCCUGG AA **CGCUCG** A CCAGGGCGAGA<br>GUCCGAGC **UU AC CACGAAGUCAAUAC** | 204 |
| **QT5B-Brick4** | CCUAGAGU **CUCCGC** A GCGAUUGUGAUCCGGCUGGUAUCAUAUGGGAUUCGUUCACCGGGAGCCGG<br>AA **CCGUCC** A CCGGCUCUCGGUGAGCGAAUCUCAUAUGGUACCAGUCGGAUCGCAAUCGC<br>ACUCUAGG CCAGUGCU **GAGGCG** A UCCACGGCGUG AA **GGUCGC** A CACGCUGUGGA<br>AGCACUGG **UU UA CACGAAGUCAAUAC** | 204 |
| **QT5B-Brick5** | GCCAGGUG **CGGGAC** A GCUCACGGAGGUGCCUGUAGGUCGGCUCGGG AA **CCCGUG** A<br>CCCGAGCUGACCUACGGGCACCUUCGUGAGC CACCUGGC GCUCUCGC **GGACGG** A<br>CCUGCUAGUAG AA **GCGGAG** A CUACUGGCAGG GCGAGAGC **UU UAAU CACGAAGUCAAUAC** | 168 |
| **HT7B-Brick1** | GGGCUGGC **GGACCC** A UCGAGAAUGCCGUUCGAGUAAGUUCCGACUAGAUGCUCGUCCAGCUG AA<br>**GGGUCC** A CAGCUGGGCGAGCAUUUAGUCGGGACUUACUUGAACGGCGUUCUCGA GCCAGCCC<br>GACGGUGC **CCAGGC** A GGAUUCUAGG AA **CUCGCC** A CCUAGGAUCC GCACCGUC **UU AU**<br>**CACGAAGUCAAUAC** | 196 |
| **HT7B-Brick2** | CCAGAUGG **CGAGCG** A GCCUCGCUGUGCGCGAUCGAUUGCGAGUG AA **CUGCCG** A<br>CACUCGCGAUCGAUUGCGCACGGCGAGGC CCAUCUGG CCCUGGCU **GGCGAG** A UCCAGGCCUG<br>AA **CGCAGC** A CAGGCUUGGA AGCCAGGG **UU AA CACGAAGUCAAUAC** | 160 |
| **HT7B-Brick3** | GCUCCGCU **GCGACC** A UGGCAAUGCGAAGGUAGCUCUGCCACUGG AA **CGCCUC** A<br>CCAGUGGUAGAGCUGCCUUCGUAUUGCCA AGCGGAGC GCUCGGUC **CGGCAG** A UCCGUCUGGG<br>AA **CGCUCG** A CCCAGGCGGA GACCGAGC **UU AC CACGAAGUCAAUAC** | 160 |
| **HT7B-Brick4** | CCAGCCGU **CUCCGC** A UGGCCUAUCUAUACGCCUACGUGGAGCGG AA **CCGUCC** A<br>CCGCUCCGCGUAGGUGUAUAGGUAGGCCA ACGGCUGG CCCUGUGC **GAGGCG** A UCCCGCUCUG<br>AA **GGUCGC** A CAGAGUGGGA GCACAGGG **UU UA CACGAAGUCAAUAC** | 160 |
| **HT7B-Brick5** | GCAGCCGU **CGGGAC** A GCACGCAGCGUCAUUGUCCAUGGCUCGGG AA **CCCGUG** A<br>CCCGAGCUAUGGACGAUGACGUUGCGUGC ACGGCUGC GCUCCUGU **GGACGG** A UCGUUCAAUG<br>AA **GCGGAG** A CAUUGGACGA ACAGGAGC **UU UAAU CACGAAGUCAAUAC** | 162 |
| **HT7B-Brick6** | CCGAGGGU **GCGCAC** A CGGCUCCUGUCUCAGAUCAUGUGCUCGGG AA **CCUCGG** A<br>CCCGAGCGCAUGAUUUGAGACGGGAGCCG ACCCUCGG CCAGACGC **CACGGG** A UCGCGCACGG<br>AA **GUCCCG** A CCGUGUGCGA GCGUCUGG **UU UACU CACGAAGUCAAUAC** | 162 |
| **HT7B-Brick7** | GCCUCUGU **GCUGCG** A CGUCAGUCCUG AA **GCCUGG** A CAGGAUUGACG ACAGAGGC<br>GCUCCCGU **CCGAGG** A UGGCUGAUGG AA **GUGCGC** A CCAUCGGCCA ACGGGAGC **UU AUAU**<br>**CACGAAGUCAAUAC** | 126 |
| **F15S-Brick1**<br>R4×3 B-related; works with R4×3-Brick1–7 or F15H | GGGCCUGC **CAGCCG** A GGGUUCCGG AA **CGAGCG** A CCGGGACCC GCAGGUCC GCUCCGAG<br>**AAUAAU** A GCAGCGUUUAGUGCAGGCUGACGCG AA **GGUCCC** A<br>CGCGUCGGCCUGUACUAAGCGCUGC CUCGGAGC **UU UUAU CACGAAGUCAAUAC** | 152 |

| Name (Note) | Sequence | length |
|---|---|---|
| **F15S-Brick2** | CCGUGAGC **AAAUAA** A GUCGGACGG AA **CGUGGG** A CCGUUCGAC GCUCACGG CCAGGCAG **AAUAAU** A GGCGGUGG AA **CGGCUG** A CCACUGCC CUGCCUGG **UU UUAA CACGAAGUCAAUAC** | 118 |
| **F15S-Brick3** | GCAUCUCG **AAAUAA** A GCACUGUGG AA **CGACGG** A CCACGGUGC CGAGAUGC GCUCGGUG **CCCACG** A UACUGGUG AA **CCCGAG** A CACCGGUA CACCGAGC **UU UUAC CACGAAGUCAAUAC** | 118 |
| **F15S-Brick4** | CCUCGAGC **UUCG** GCUCGAGG GCUGAGUC **CCGUCG** A GGCGGUGG AA **CCGGUG** A CCACUGCC GACUCAGC **UU UAUU CACGAAGUCAAUAC** | 88 |
| **F15S-Brick5** | GCACGGAC **GAGCCC** A UCAAGGC **GUAA** GCCUUGA GUCCGUGC GCACCAGG **CGGACG** A UCUGCCA **UUCG** UGGCAGA CCUGGUGC **UU UAAU CACGAAGUCAAUAC** | 102 |
| **F15S-Brick6** | CCGAGCUG **CUCCCG** A GGUCGCGUG AA **GAGGGC** A CACGUGACC CAGCUCGG CCUCCGGU **GCCAGC** A CUCGGUCG AA **GGGCUC** A CGACUGAG ACCGGAGG **UU UACU CACGAAGUCAAUAC** | 118 |
| **F15S-Brick7** | GCCAGUCC **GUGCGC** A UCGACCU **GUAA** AGGUCGA GGACUGGC GCAGCUAC **GCCCUC** A UGGAGUA **UUCG** UACUCCA GUAGCUGC **UU AAUU CACGAAGUCAAUAC** | 102 |
| **F15S-Brick8** | CCAGGAGC **UUCG** GCUCCUGG GCCUGCGU **CGCUGG** A UCCGGUAG AA **GCGCAC** A CUACUGGA ACGCAGGC **UU ACUU ACGAAGUCAAUAC** | 87 |
| **F15H-Brick1**<br>R4×3 A-related; works with F15S | GGGCUCGCGGUCC **CAGCGC** A GGUCUCAUG AA **GCUGGC** A CAUGGGACC GGACCGUGAGUCC GCAGGUAGCGAACC **GCGUGC** A UCAGGCUGUAUCGGUAACUGUGCGG AA **CGUCCG** A CCGCACGGUUACUGAUACGGCCUGA GGUUCGUUACCUGC **UU UUAU CACGAAGUCAAUAC** | 174 |
| **F15H-Brick2** | CCUCUGUGCCUGU **GACCCG** A UCGGGAGUG AA **GGACCG** A CACUUCCGA ACAGGCGCAGAGG CCAGAUAUCUGAGU **GCCUGG** A CUCGACGG AA **GCGCUG** A CCGUUGAG ACUCAGGUAUCUGG **UU UUAA CACGAAGUCAAUAC** | 140 |
| **F15H-Brick3** | GCUCGUGGCACGU **CUGCGC** A UGGCUGAUG AA **CCAGCG** A CAUCGGCCA ACGUGCUACGAGC GCUAGCGGUGCUAU **CGGUCC** A GGCUGUGG AA **CGGGAG** A CCACGGCC AUAGCAUCGCUAGC **UU UUAC CACGAAGUCAAUAC** | 140 |
| **F15H-Brick4** | CCAGCGUC **UUCG** GACGCUGG GCAGGACC **CGGUGG** A UCCAGCUGCAGCGGAGGUG AA **GCGCAG** A CACCUCUGCUGCGGCUGGA GGUCCUGC **UU UAUU CACGAAGUCAAUAC** | 110 |
| **F15H-Brick5** | GCCUCAGC **GUGCCC** A UCCGUGAUG AA **CCAGGC** A CAUCGCGGA GCUGAGGC GCUCGGUC **GGGACC** A UGGCGUUCUGGUAGGUCGAGGUAGCGGAGACCUCUG AA **GCACGC** A CAGAGGUUUCCGCUGCCUCGAUCUACCAGGACGCCA GACCGAGC **UU UAAU CACGAAGUCAAUAC** | 174 |
| **F15H-Brick6** | CCAGCUGC **CUCGGG** A GCGUGCGGG AA **CCUCGG** A CCCGUACGC GCAGCUGG CCCUGCUG **CGCUCG** A UCCGGUCG AA **GGGCAC** A CGACUGGA CAGCAGGG **UU UACU CACGAAGUCAAUAC** | 118 |
| **F15H-Brick7** | GCUCGAGC **CACCGG** A GCUGUAGUG AA **CCACCG** A CACUGCAGC GCUCGAGC GCCUGCGU **CCGAGG** A UGCUCUGG AA **CGGGUC A CCAGGGCA ACGCAGGC UU AUUU CACGAAGUCAAUAC** | 118 |
| **HT3WJ-Brick1** | GGGCUC GGGUCCAGAGGUGC **GACCCG** A UCCGUACGUG AA **GGACCG** A CACGUGCGGA GCACCUUUGGACCC GCACGACGACGCGU **GCCAGC** A UCCAUGGGAUGGUCUUCAGAUGCUCCCGG AA **GCGCUG** A CCGGGAGUAUCUGAGGACCAUUCCAUGGA ACGCGUUGUCGUGC GAGCCCGACUGCAC **GCGUGC** A UCACCGGGUCUACAUUGAUAUAGGGUGCGAGUCUAUACCGGUUCGUG AA **GCACGC** A CACGAACUGGUAUAGGCUCGCACUCUAUAUCGAUGUAGAUCCGGUGA GUGCAGUC **UU UUAU CACGAAGUCAAUAC** | 324 |
| **HT3WJ-Brick2** | CCGAGC GCACCGCUAUAGGU **CUGCGC** A UCGUGGAUGG AA **CCAGCG** A CCAUCUACGA ACCUAUGGCGGUGC GGACCGGUAUUGCU **CGGUCC** A ACGCGAGUCAGCCUGGCAGUCUGGAGCGG AA **CGGGUC** A CCGCUCCGGACUGCUAGGCUGGCUCGCGU AGCAAUGCCGGUCC GCUCGGGCAGCC **UUCG** GGCUGC **UU UUAA CACGAAGUCAAUAC** | 214 |
| **HT3WJ-Brick3** | GCAUGC GCCAGCCGUCCGAU **CAGCGC** A UGGUUCUCAG AA **GCUGGC** A CUGAGGACCA AUCGGAUGGCUGGC GGUCGGUGCUAAGC **CGCUGG** A AUACCAGGCGAGGUUCCGCAGGAUACUGG AA **GCGCAG** A CCAGUAUUCUGCGGGACCUCGUCUGGUAU GCUUAGUACCGACC GCAUGCGCCUGG **UUCG** CCAGGC **UU UUAC CACGAAGUCAAUAC** | 214 |
| **UR4B-Brick1** | GGGCCUGC **CGCUGG** A UCCACGUCCUCGGG AA **CGGCUG** A CCCGAGGGCGUGGA GCAGGUCC GCUGUAAC **CCGCUC** A GGGACAUGUCCGGG AA **CGCGUG** A CCCGGACGUGUCCC GUUACAGC **UU UUAU CACGAAGUCAAUAC** | 140 |

| Name (Note) | Sequence | length |
|---|---|---|
| **UR4B-Brick2** | CCAGGAUGUCUGG **CCUCGC** A ACUAGGGCUACAGG AA **CCAGCG** A CCUGUAGUCCUAGU CCAGACGUCCUGG CCGUAGGUAUCCGU **GGGACC** A UGUAUCGCGACCUG AA **GAGCGG** A CAGGUCGUGAUACA ACGGAUGCCUACGG **UU UUAA CACGAAGUCAAUAC** | 162 |
| **UR4B-Brick3** | GCCUUCGGCUGUUCCGCUC **CGGGAG** A UGCGUGUAUCCUUG AA **GCGAGG** A CAAGGAUGCACGCA GAGCGGGACAGCUGAAGGC GCUUCUGGAACGUCCAUGC **GACGGG** A UCCUAGUCCUCGCG AA **GGUCCC** A CGCGAGGGCUAGGA GCAUGGGCGUUCUAGAAGC **UU UUAC CACGAAGUCAAUAC** | 184 |
| **UR4B-Brick4** | CCAGACUGACAGC **CAGCCG** A GGCUACGGCCUCAG AA **CUCCCG** A CUGAGGCUGUAGCC GCUGUCGGUCUGG CCUUCGAUCGGUCC **CACGCG** A UGGGUAGGUCCGGG AA **CCCGUC** A CCCGGACUUACCCA GGACCGGUCGAAGG **UU UAUU CACGAAGUCAAUAC** | 162 |
| **UA4B-Brick1** | GGGCUCUG **CGCUGG** A UCCAGUUGAGCGGACCUAUCCUGGG AA **CGGCUG** A CCCAGGGUAGGUUCGCUCGACUGGA CAGAGUCC GCUCGCAC **CCGCUC** A UCUGGCUGACCGGG AA **CGCGUG** A CCCGGUCGGCCAGA GUGCGAGC **UU UUAU CACGAAGUCAAUAC** | 162 |
| **UA4B-Brick2** | CCUCUGUCUGAGU **CCUCGC** A ACCGUGGAGUGUUAGGACGGAUGUG AA **CCAGCG** A CACAUCUGUCCUGACACUUCACGGU ACUCAGGCAGAGG CCAUCGCUUCGGCU **GGGACC** A UCGUGCGCAGCAUG AA **GAGCGG** A CAUGCUGUGCACGA AGCCGAGGCGAUGG **UU UUAA CACGAAGUCAAUAC** | 184 |
| **UA4B-Brick3** | GCAGUAGGGACGUACCCUG **CGGGAG** A UCCGUUUCGUGAGCCUACUGUCGCG AA **GCGAGG** A CGCGACGGUAGGUUCACGGAACGGA CAGGGUGCGUCCUUACUGC GCUCGAGGCAGAUGAGUGC **GACGGG** A CUCGUCUGGGAGCG AA **GGUCCC** A CGCUCCCGGACGAG GCACUCGUCUGCUUCGAGC **UU UUAC CACGAAGUCAAUAC** | 206 |
| **UA4B-Brick4** | CCGAUGUCCAUGC **CAGCCG** A UCUAGAGCUGCGUAGUGUGCCAGUG AA **CUCCCG** A CACUGGUACACUGCGCAGUUCUAGA GCAUGGGCAUCGG CCUACUGUACAGGG **CACGCG** A UCUGACGCCUCGGG AA **CCCGUC** A CCCGAGGUGUCAGA CCCUGUGCAGUAGG **UU UAUU CACGAAGUCAAUAC** | 184 |
| **Z-Brick-7A**<br>Strut 13+8 bp;<br>beam 14 bp + A | GGAUACUUGUCAG **GAGAUCC** CUGCUGGACUCUCG AAA **GGAUCUC** A CGAGAGUUCAGCAG CUGACAGGUAUCC GGUUGGAC **GUCACAG** CUGACUUGAUCACG AAA **CUGUGAC** A CGUGAUCGAGUCAG **GUCCAACC UU CAUGCUUACGACG** | 149 |
| **Z-Brick-7B**<br>Strut 13+8 bp;<br>beam 15 bp | GGAUACUUGUCAG **GAGAUCC** CUUGCUGGACUCUCG AA **GGAUCUC** CGAGAGUUCAGCAAG CUGACAGGUAUCC GGUUGGAC **GUCACAG** CCUGACUUGAUCACG AA **CUGUGAC** CGUGAUCGAGUCAGG **GUCCAACC UU CAUGCUUACGACG** | 149 |
| **Z-Brick-8A**<br>Strut 13+8 bp;<br>beam 13 bp + A | GGAUACUUGUCAG **GAGAUCAC** CUGCUGGACUCUG AAA **GUGAUCUC** A CAGAGUUCAGCAG CUGACAGGUAUCC GGUUGGAC **GUACACAG** CUGACUUGAUCCG AAA **CUGUGUAC** A CGGAUCGAGUCAG **GUCCAACC UU CAUGCUUACGACG** | 149 |
| **Z-Brick-8B**<br>Strut 13+8 bp;<br>beam 14 bp | GGAUACUUGUCAG **GAGAUCAC** CUGCUGGACUCUCG AA **GUGAUCUC** CGAGAGUUCAGCAG CUGACAGGUAUCC GGUUGGAC **GUACACAG** CUGACUUGAUCACG AA **CUGUGUAC** CGUGAUCGAGUCAG **GUCCAACC UU CAUGCUUACGACG** | 149 |
| **Z-Brick-9A**<br>Strut A+12+8+A bp; beam 12 bp + A | GGAUCGUGUUACA **CAUUCAGAG** CCGGAGUAUGCG AAAA **CUCUGAAUG** A CGCAUAUUCCGG AGUAACGCGAUCC GGUUGGACA **GUUACCUAG** GCUCGUGACCCG AAAA **CUAGGUAAC** A CGGGUCGCGAGU A**GUCCAACC UU CAUGCUUACGACG** | 153 |
| **Z-Brick-9B**<br>Strut A+12+8+A bp; beam 13 bp | GGAUAAUGUCGCA **CAUUCAGAG** GCUCAGGGCCUGG AAA **CUCUGAAUG** CCAGGCUCUGAGC AGCGACGUUAUCC GGUUGGACA **GUUACCUAG** GCUCGGUGACCGG AAA **CUAGGUAAC** CCGGUCGCCGAGU A**GUCCAACC UU CAUGCUUACGACG** | 153 |
| **LR3B-Brick1** | GGGCACGGACGAGC A **GCUUCAAUC** GGAUCGCAUCAUCAGGUG AAA **GAAGGUAUC** CACCUGGUGAUGUGAUCC A GCUCGUCUGUGCUC CCAGGUGAGGC A **CUACUCGUA** CCCGUCUUGACGAAAUGAUCACUUAGUUAGUUGAUCUAGGUUGCGG AAA **UACGAGUAG** CCGCAACUUAGAUCAGCUAACUAGGUGAUCGUUUCGUCGAGACGGG A GCCUCGCCUGG **UU AU CACGAAGUCAAUAC** | 242 |
| **LR3B-Brick2** | GCAGUGUCAGCUGC A **CCUUGAGAA** CAGAAGUACGCAGGAGUGG AAA **UUCAUAGCG** CCACUCUUGCGUGCUUCUG A GCAGCUGGCACUGC GCUCCUGGAGC A **GAUACCUUC** GCGAGAUGGCUCUAUACUG AAA **GAUUGAAGC** CAGUAUGGAGCCGUCUCGC A GCUCCGGGAGC **UU AA CACGAAGUCAAUAC** | 190 |
| **LR3B-Brick3** | CCAGCGGA **UUCG** UCCGCUGG GCCUCCGC A **CGCUAUGAA** CCCGCUCCCAAGCGACGG AAA **UUCUCAAGG** CCGUCGCUUGGGAGCGGG A GCGGAGGC **UU AC CACGAAGUCAAUAC** | 113 |

#### 3) Templates for dsRNA bricks

| Name (Note) | Sequence | length |
|---|---|---|
| **R4B-TMP**<br>Template for R4B-Brick1–4 | GTTCTAATACGACTCACTATA GGGCAGGC CAGCGC A GGTCTACGG AA GCTGGC A<br>CCGTGGACC GCCTGTCC GCTCGGTC GCGTGC A CCAGCCTGTTTAGGAGTATTCCTAG AA<br>GCACGC A CTAGGAGTACTCTTAAACGGGCTGG GACCGAGC TT TTAT CACGAAGTCAATAC<br>CCATCGCT GACCCG A TGACGGGTG AA GGACCG A CACCTGTCA AGCGATGG CCTCGGTC<br>GCCAGC A TACGGGTG AA GCGCTG A CACCTGTA GACCGAGG TT TTAA<br>CACGAAGTCAATAC GCACTGGT CTGCGC A GATGTCAGG AA CCAGCG A CCTGGCATC<br>ACCAGTGC GCTCGCGT CGGTCC A TACTCTGG AA CGGGTC A CCAGGGTA ACGCGAGC TT<br>TTAC CACGAAGTCAATAC CCTCGACC TTCG GGTCGAGG GCCTGCGT CGCTGG A<br>TCCGAGTG AA GCGCAG A CACTTGGA ACGCAGGC TT TATT CACGAAGTCAATAC<br>GTCCAACC TT CATGCTTACGACG<br>GGCCGGCATGGTCCCAGCCTCCTCGCTGGCGCCGGCTGGGCAACATGCTTCGGCATGGCGAATGGGAC | 588 |
| **R6B-TMP**<br>Template for R6B-Brick1–6 | GTTCTAATACGACTCACTATA GGGCAGGC CAGCGC A GGTCTACGG AA GCTGGC A<br>CCGTGGACC GCCTGTCC GCTCGGTC GCGTGC A CCAGCCTGTTTAGGAGTATTCCTAG AA<br>GCACGC A CTAGGAGTACTCTTAAACGGGCTGG GACCGAGC TT TTAT CACGAAGTCAATAC<br>CCATCGCT GACCCG A TGACGGGTG AA GGACCG A CACCTGTCA AGCGATGG CCTCGGTC<br>GCCAGC A TACGGGTG AA GCGCTG A CACCTGTA GACCGAGG TT TTAA<br>CACGAAGTCAATAC GCACTGGT CTGCGC A GATGTCAGG AA CCAGCG A CCTGGCATC<br>ACCAGTGC GCTCGCGT CGGTCC A TACTCTGG AA CGGGTC A CCAGGGTA ACGCGAGC TT<br>TTAC CACGAAGTCAATAC CCAGCAGT GAGGCC A TGACGCGTG AA GGTGGC A<br>CACGTGTCA ACTGCTGG CTCCCGCT CGCTGG A TCCGGCAG AA GCGCAG A CTGCTGGA<br>AGCGGGAG TT TATT CACGAAGTCAATAC GCAGGCGT GTCGCG A CCACTGCAG AA<br>GCACCC A CTGCGGTGG ACGCCTGC GCAGGAGC GCCACC A TTCTGTCG AA GGCCTC A<br>CGACGGAA GCTCCTGC TT TAAT CACGAAGTCAATAC CCGTCGTC TTCG GACGACGG<br>GCTCCCGA GGGTGC A TCGGTCTG AA CGCGAC A CAGATCGA TCGGGAGC TT TACT<br>CACGAAGTCAATAC GTCCAACC TT CATGCTTACGACG<br>GGCCGGCATGGTCCCAGCCTCCTCGCTGGCGCCGGCTGGGCAACATGCTTCGGCATGGCGAATGGGAC | 824 |
| **R8B-TMP**<br>Template for R8B-Brick1–8 | GTTCTAATACGACTCACTATA GGGCAGGC CAGCGC A GGTCTACGG AA GCTGGC A<br>CCGTGGACC GCCTGTCC GCTCGGTC GCGTGC A CCAGCCTGTTTAGGAGTATTCCTAG AA<br>GCACGC A CTAGGAGTACTCTTAAACGGGCTGG GACCGAGC TT TTAT CACGAAGTCAATAC<br>CCATCGCT GACCCG A TGACGGGTG AA GGACCG A CACCTGTCA AGCGATGG CCTCGGTC<br>GCCAGC A TACGGGTG AA GCGCTG A CACCTGTA GACCGAGG TT TTAA<br>CACGAAGTCAATAC GCACTGGT CTGCGC A GATGTCAGG AA CCAGCG A CCTGGCATC<br>ACCAGTGC GCTCGCGT CGGTCC A TACTCTGG AA CGGGTC A CCAGGGTA ACGCGAGC TT<br>TTAC CACGAAGTCAATAC CCAGCAGT GAGGCC A TGACGCGTG AA GGTGGC A<br>CACGTGTCA ACTGCTGG CTCCCGCT CGCTGG A TCCGGCAG AA GCGCAG A CTGCTGGA<br>AGCGGGAG TT TATT CACGAAGTCAATAC GCAGGCGT GTCGCG A CCACTGCAG AA<br>GCACCC A CTGCGGTGG ACGCCTGC GCAGGAGC GCCACC A TTCTGTCG AA GGCCTC A<br>CGACGGAA GCTCCTGC TT TAAT CACGAAGTCAATAC CCAGTGCT CTCGCC A GGGTGCCTG<br>AA GGAGCC A CAGGTACCC AGCACTGG CCAGCCGA GGGTGC A TCCGGTAG AA CGCGAC<br>A CTACTGGA TCGGCTGG TT TACT CACGAAGTCAATAC GCCAGGCT GTGCCG A<br>TCAGTACGG AA CGTCGC A CCGTGCTGA AGCCTGGC GCAGGGTC GGCTCC A TCATAGTG<br>AA GGCGAG A CACTGTGA GACCCTGC TT ATTT CACGAAGTCAATAC CCTCGGTC TTCG<br>GACCGAGG GCTCGGGT GCGACG A CCCGGTAG AA CGGCAC A CTACTGGG ACCCGAGC TT<br>AATT CACGAAGTCAATAC GTCCAACC TT CATGCTTACGACG<br>GGCCGGCATGGTCCCAGCCTCCTCGCTGGCGCCGGCTGGGCAACATGCTTCGGCATGGCGAATGGGAC | 1060 |
| **R4×3-A-TMP**<br>Template for R4×3-Brick1–7 (A set) | GTTCTAATACGACTCACTATA GGGCAGGC CAGCGC A GGACTACGG AA GCTGGC A<br>CCGTGGTCC GCCTGTCC GCTCGAGG GCGTGC A TGAGTGTCTGTAGAGTACTGAGCGG AA<br>CGTCCG A CCGCTCGGTACTTTACAGGCACTCA CCTCGAGC TT TTAT CACGAAGTCAATAC<br>CCAGCAGG GACCCG A TCTCGGGTG AA GGACCG A CACCTGAGA CCTGCTGG CCCTCGCT<br>GCCTGG A CTCGACGG AA GCGCTG A CCGTTGAG AGCGAGGG TT TTAA<br>CACGAAGTCAATAC GCCTCTGT CTGCGC A GGAGTACGG AA CCAGCG A CCGTGCTCC<br>ACAGAGGC GCTCTTGT CGGTCC A GCTTCGGG AA CGGGAG A CCCGGAGC ACAAGAGC TT<br>TTAC CACGAAGTCAATAC CCAGCAGC TTCG GCTGCTGG GCCTCTGC CGGTGG A<br>TGGGCCAG AA GCGCAG A CTGGTCCA GCAGAGGC TT TATT CACGAAGTCAATAC<br>GCCTCGGC GTGCCC A CTCATACGG AA CCAGGC A CCGTGTGAG GCCGAGGC GCTCCGTC<br>GGGACC A TCGTTGTATGCTGCAACTTGGCTGG AA GCACGC A<br>CCAGCCGAGTTGTAGCATGCAACGA GACGGAGC TT TAAT CACGAAGTCAATAC CCAGCTGC<br>CTCGGG A GCGTGCGGG AA CCTCGG A CCCGTACGC GCAGCTGG CCCTGCTG CGCTCG A<br>TCCGGTCG AA GGGCAC A CGACTGGA CAGCAGGG TT TACT CACGAAGTCAATAC<br>GCTCGAGC CACCGG A GCTGTAGTG AA CCACCG A CACTGCAGC GCTCGAGC GCCTGCGT<br>CCGAGG A TGCTCTGG AA CGGGTC A CCAGGGCA ACGCAGGC TT ATTT<br>CACGAAGTCAATAC GTCCAACC TT CATGCTTACGACG<br>GGCCGGCATGGTCCCAGCCTCCTCGCTGGCGCCGGCTGGGCAACATGCTTCGGCATGGCGAATGGGAC | 976 |

| Name (Note) | Sequence | length |
|---|---|---|
| **R4×3-B-TMP**<br>Template for R4×3-Brick8–12 (B set) | GTTCTAATACGACTCACTATA GGGCCTGC CAGCCG A GGGTTCCGG AA CGAGCG A<br>CCGGGACCC GCAGGTCC GCTCCAGC CGGACG A GCTCAATTGACCGTTGACTGGAGCG AA<br>GGTCCC A CGCTCCGGTCAATGGTCAGTTGAGC GCTGGAGC TT TTAT CACGAAGTCAATAC<br>CCAGTGAG CTCCCG A GGCGGACGG AA CGTGGG A CCGTTCGCC CTCACTGG CCCTGGTC<br>GCCAGC A CGCGGTGG AA CGGCTG A CCACTGCG GACCAGGG TT TTAA<br>CACGAAGTCAATAC GGGAGTCC CTGGCG A GCTCTGTGG AA CGACGG A CCACGGAGC<br>GGACTCCC GCTCAGGT CCCACG A GGCTTCTG AA CCCGAG A CAGAGGCC ACCTGAGC TT<br>TTAC CACGAAGTCAATAC CCTCACGC TTCG GCGTGAGG GCCTGTCTTTATGGGCAGAACG<br>CGCTGG A TCCGCAGG AA CGCCAG A CCTGTGGA CGTTCTGTCCATAAGGACAGGC TT<br>TATT CACGAAGTCAATAC CCAGAGCC TTCG GGCTCTGG GCAGGCGTTCTGTGC CCGTCG A<br>GGCGGTGG AA CCGGTG A CCACTGCC GCACAGAGCGCCTGC TT TAAT CACGAAGTCAATAC<br>GTCCAACC TT CATGCTTACGACG<br>GGCCGGCATGGTCCCAGCCTCCTCGCTGGCGCCGGCTGGGCAACATGCTTCGGCATGGCGAATGGGAC | 718 |
| **A4B-TMP**<br>Template for A4B-Brick1–4 | GTTCTAATACGACTCACTATA GGGCATCC CAGCGC A GGGATCTAG AA GCTGGC A<br>CTAGGTCCC GGATGCCC GACCGGTC GCGTGC A TGGAACTTACCGTG AA GCACGC A<br>CACGGTAGGTTCCA GACCGGTC TT TTAA CACGAAGTCAATAC CCTCGCGT GACCCG A<br>TGTCTAGTGACGATGGCGTG AA GGACCG A CACGCCGTCGTCATTAGACA ACGCGAGG<br>CCAGGGTC GCCAGC A TACGGGTG AA GCGCTG A CACCTGTA GACCCTGG TT TTAC<br>CACGAAGTCAATAC GCAGGCTC CTGCGC A GAGGCTTAGCCTAGCCTGGG AA CCAGCG A<br>CCCAGGTTAGGCTGAGCCTC GAGCCTGC GCTAGCGT CGGTCC A GCGTCTGG AA CGGGTC A<br>CCAGGCGC ACGCTAGC TT TATT CACGAAGTCAATAC CCAGGACC TTCG GGTCCTGG<br>GCCTGCCT CGCTGG A ACGCGTGGACAGTTGAGTG AA GCGCAG A<br>CACTCAGCTGTCTACGCGT AGGCAGGC TT TAAT CACGAAGTCAATAC GTCCAACC TT<br>CATGCTTACGACG<br>GGCCGGCATGGTCCCAGCCTCCTCGCTGGCGCCGGCTGGGCAACATGCTTCGGCATGGCGAATGGGAC | 632 |
| **QT5B-TMP**<br>Template for QT5B-Brick1–5 | GTTCTAATACGACTCACTATA GGGCCTAC GCGCAC A<br>TGTCCCGTTGCTCAGGAATTGATCGGACTAGATGGCGATTTGTATCGGTGTAATCCTCGGAGGCTTCT<br>G AA GTGCGC A<br>CAGAAGCTTCCGAGGGTTACACTGATACAAGTCGCCATTTAGTCCGGTCAATTTCTGAGCAGCGGGAC<br>A GTAGGCTC GCGACCGC CACGGG A TGCTGTATCTG AA CTCGCC A CAGATGCAGCA<br>GCGGTCGC TT AT CACGAAGTCAATAC CCGTCCTG CGAGCG A<br>GCCGTCCTAGCACTGCTCATATCGTGAGGACTGCATGCGTCGGTGACATG AA CTGCCG A<br>CATGTCATCGACGCGTGCAGTTCTCACGGTATGAGTAGTGCTGGGACGGC CAGGACGG CCAGGGTC<br>GGCGAG A CCAGCGGCTGG AA GTCCCG A CCAGCTGCTGG GACCCTGG TT AA<br>CACGAAGTCAATAC GCAGGCGT GCGACC A<br>TCCGTATGGAGTGATGCTTCTGGCATCATTCCTACGGATCTCTGTACAGG AA CGCCTC A<br>CCTGTACGGAGATCTGTAGGAGTGATGCTAGAAGCGTCACTCTATACGGA ACGCCTGC GCTCGGAC<br>CGGCAG A TCTCGTCCTGG AA CGCTCG A CCAGGGCGAGA GTCCGAGC TT AC<br>CACGAAGTCAATAC CCTAGAGT CTCCGC A<br>GCGATTGTGATCCGGCTGGTATCATATGGGATTCGTTCACCGGGAGCCGG AA CCGTCC A<br>CCGGCTCTCGGTGAGCGAATCTCATATGGTACCAGTCGGATCGCAATCGC ACTCTAGG CCAGTGCT<br>GAGGCG A TCCACGGCGTG AA GGTCGC A CACGCTGTGGA AGCACTGG TT TA<br>CACGAAGTCAATAC GCCAGGTG CGGGAC A GCTCACGGAGGTGCCTGTAGGTCGGCTCGGG AA<br>CCCGTG A CCCGAGCTGACCTACGGGCACCTTCGTGAGC CACCTGGC GCTCTCGC GGACGG A<br>CCTGCTAGTAG AA GCGGAG A CTACTGGCAGG GCGAGAGC TT TAAT CACGAAGTCAATAC<br>GTCCAACC TT CATGCTTACGACG | 1066 |

| Name (Note) | Sequence | length |
|---|---|---|
| **HT7B-TMP**<br>Template for HT7B-Brick1–7 | **GTTCTAATACGACTCACTATA** GGGCTGGC **GGACCC** A<br>TCGAGAATGCCGTTCGAGTAAGTTCCGACTAGATGCTCGTCCAGCTG AA **GGGTCC** A<br>CAGCTGGGCGAGCATTTAGTCGGGACTTACTTGAACGGCGTTCTCGA GCCAGCCC GACGGTGC<br>**CCAGGC** A GGATTCTAGG AA **CTCGCC** A CCTAGGATCC GCACCGTC **TT AT**<br>**CACGAAGTCAATAC** CCAGATGG **CGAGCG** A GCCTCGCTGTGCGCGATCGATTGCGAGTG AA<br>**CTGCCG** A CACTCGCGATCGATTGCGCACGGCGAGGC CCATCTGG CCCTGGCT **GGCGAG** A<br>TCCAGGCCTG AA **CGCAGC** A CAGGCTTGGA AGCCAGGG **TT AA CACGAAGTCAATAC**<br>GCTCCGCT **GCGACC** A TGGCAATGCGAAGGTAGCTCTGCCACTGG AA **CGCCTC** A<br>CCAGTGGTAGAGCTGCCTTCGTATTGCCA AGCGGAGC GCTCGGTC **CGGCAG** A TCCGTCTGGG<br>AA **CGCTCG** A CCCAGGCGGA GACCGAGC **TT AC CACGAAGTCAATAC** CCAGCCGT **CTCCGC**<br>A TGGCCTATCTATACGCCTACGTGGAGCGG AA **CCGTCC** A<br>CCGCTCCGCGTAGGTGTATAGGTAGGCCA ACGGCTGG CCCTGTGC **GAGGCG** A TCCCGCTCTG<br>AA **GGTCGC** A CAGAGTGGGA GCACAGGG **TT TA CACGAAGTCAATAC** GCAGCCGT **CGGGAC**<br>A GCACGCAGCGTCATTGTCCATGGCTCGGG AA **CCCGTG** A<br>CCCGAGCTATGGACGATGACGTTGCGTGC ACGGCTGC GCTCCTGT **GGACGG** A TCGTTCAATG<br>AA **GCGGAG** A CATTGGACGA ACAGGAGC **TT TAAT CACGAAGTCAATAC** CCGAGGGT<br>**GCGCAC** A CGGCTCCTGTCTCAGATCATGTGCTCGGG AA **CCTCGG** A<br>CCCGAGCGCATGATTTGAGACGGGAGCCG ACCCTCGG CCAGACGC **CACGGG** A TCGCGCACGG<br>AA **GTCCCG** A CCGTGTGCGA GCGTCTGG **TT TACT CACGAAGTCAATAC** GCCTCTGT<br>**GCTGCG** A CGTCAGTCCTG AA **GCCTGG** A CAGGATTGACG ACAGAGGC GCTCCCGT<br>**CCGAGG** A TGGCTGATGG AA **GTGCGC** A CCATCGGCCA ACGGGAGC **TT ATAT**<br>**CACGAAGTCAATAC GTCCAACC TT CATGCTTACGACG** | 1170 |
| **F15S-TMP**<br>Template for F15S-Brick1–8 | **GTTCTAATACGACTCACTATA** GGGCCTGC **CAGCCG** A GGGTTCCGG AA **CGAGCG** A<br>CCGGGACCC GCAGGTCC GCTCCGAG **AATAAT** A GCAGCGTTTAGTGCAGGCTGACGCG AA<br>**GGTCCC** A CGCGTCGGCCTGTACTAAGCGCTGC CTCGGAGC **TT TTAT CACGAAGTCAATAC**<br>CCGTGAGC **AAATAA** A GTCGGACGG AA **CGTGGG** A CCGTTCGAC GCTCACGG CCAGGCAG<br>**AATAAT** A GGCGGTGG AA **CGGCTG** A CCACTGCC CTGCCTGG **TT TTAA**<br>**CACGAAGTCAATAC** GCATCTCG **AAATAA** A GCACTGTGG AA **CGACGG** A CCACGGTGC<br>CGAGATGC GCTCGGTG **CCCACG** A TACTGGTG AA **CCCGAG** A CACCGGTA CACCGAGC **TT**<br>**TTAC CACGAAGTCAATAC** CCTCGAGC **TTCG** GCTCGAGG GCTGAGTC **CCGTCG** A<br>GGCGGTGG AA **CCGGTG** A CCACTGCC GACTCAGC **TT TATT CACGAAGTCAATAC**<br>GCACGGAC **GAGCCC** A TCAAGGC **GTAA** GCCTTGA GTCCGTGC GCACCAGG **CGGACG** A<br>TCTGCCA **TTCG** TGGCAGA CCTGGTGC **TT TAAT CACGAAGTCAATAC** CCGAGCTG **CTCCCG**<br>A GGTCGCGTG AA **GAGGGC** A CACGTGACC CAGCTCGG CCTCCGGT **GCCAGC** A<br>CTCGGTCG AA **GGGCTC** A CGACTGAG ACCGGAGG **TT TACT CACGAAGTCAATAC**<br>GCCAGTCC **GTGCGC** A TCGACCT **GTAA** AGGTCGA GGACTGGC GCAGCTAC **GCCCTC** A<br>TGGAGTA **TTCG** TACTCCA GTAGCTGC **TT AATT CACGAAGTCAATAC** CCAGGAGC **TTCG**<br>GCTCCTGG GCCTGCGT **CGCTGG** A TCCGGTAG AA **GCGCAC** A CTACTGGA ACGCAGGC **TT**<br>**ACTT ACGAAGTCAATAC GTCCAACC TT CATGCTTACGACG** | 929 |
| **F15H-TMP**<br>Template for F15H-Brick1–7 | **GTTCTAATACGACTCACTATA** GGGCTCGCGGTCC **CAGCGC** A GGTCTCATG AA **GCTGGC** A<br>CATGGGACC GGACCGTGAGTCC GCAGGTAGCGAACC **GCGTGC** A<br>TCAGGCTGTATCGGTAACTGTGCGG AA **CGTCCG** A CCGCACGGTTACTGATACGGCCTGA<br>GGTTCGTTACCTGC **TT TTAT CACGAAGTCAATAC** CCTCTGTGCCTGT **GACCCG** A<br>TCGGGAGTG AA **GGACCG** A CACTTCCGA ACAGGCGCAGAGG CCAGATATCTGAGT **GCCTGG**<br>A CTCGACGG AA **GCGCTG** A CCGTTGAG ACTCAGGTATCTGG **TT TTAA**<br>**CACGAAGTCAATAC** GCTCGTGGCACGT **CTGCGC** A TGGCTGATG AA **CCAGCG** A<br>CATCGGCCA ACGTGCTACGAGC GCTAGCGGTGCTAT **CGGTCC** A GGCTGTGG AA **CGGGAG** A<br>CCACGGCC ATAGCATCGCTAGC **TT TTAC CACGAAGTCAATAC** CCAGCGTC **TTCG**<br>GACGCTGG GCAGGACC **CGGTGG** A TCCAGCTGCAGCGGAGGTG AA **GCGCAG** A<br>CACCTCTGCTGCGGCTGGA GGTCCTGC **TT TATT CACGAAGTCAATAC** GCCTCAGC **GTGCCC**<br>A TCCGTGATG AA **CCAGGC** A CATCGCGGA GCTGAGGC GCTCGGTC **GGGACC** A<br>TGGCGTTCTGGTAGGTCGAGGTAGCGGAGACCTCTG AA **GCACGC** A<br>CAGAGGTTTCCGCTGCCTCGATCTACCAGGACGCCA GACCGAGC **TT TAAT CACGAAGTCAATAC**<br>CCAGCTGC **CTCGGG** A GCGTGCGGG AA **CCTCGG** A CCCGTACGC GCAGCTGG CCCTGCTG<br>**CGCTCG** A TCCGGTCG AA **GGGCAC** A CGACTGGA CAGCAGGG **TT TACT**<br>**CACGAAGTCAATAC** GCTCGAGC **CACCGG** A GCTGTAGTG AA **CCACCG** A CACTGCAGC<br>GCTCGAGC GCCTGCGT **CCGAGG** A TGCTCTGG AA **CGGGTC A CCAGGGCA ACGCAGGC TT**<br>**ATTT CACGAAGTCAATAC GTCCAACC TT CATGCTTACGACG** | 1018 |

| Name (Note) | Sequence | length |
|---|---|---|
| **HT3WJ-Gene**<br>Template for HT3WJ-Brick1–3 | **GTTCTAATACGACTCACTATA** GGGCTC GGGTCCAGAGGTGC **GACCCG** A TCCGTACGTG AA<br>**GGACCG** A CACGTGCGGA GCACCTTTGGACCC GCACGACGACGCGT **GCCAGC** A<br>TCCATGGGATGGTCTTCAGATGCTCCCGG AA **GCGCTG** A<br>CCGGGAGTATCTGAGGACCATTCCATGGA ACGCGTTGTCGTGC GAGCCCGACTGCAC **GCGTGC** A<br>TCACCGGGTCTACATTGATATAGGGTGCGAGTCTATACCGGTTCGTG AA **GCACGC** A<br>CACGAACTGGTATAGGCTCGCACTCTATATCGATGTAGATCCGGTGA GTGCAGTC **TT** **TTAT**<br>**CACGAAGTCAATAC** CCGAGC GCACCGCTATAGGT **CTGCGC** A TCGTGGATGG AA **CCAGCG** A<br>CCATCTACGA ACCTATGGCGGTGC GGACCGGTATTGCT **CGGTCC** A<br>ACGCGAGTCAGCCTGGCAGTCTGGAGCGG AA **CGGGTC** A<br>CCGCTCCGGACTGCTAGGCTGGCTCGCGT AGCAATGCCGGTCC GCTCGGGCAGCC **TTCG**<br>GGCTGC **TT** **TTAA** **CACGAAGTCAATAC** GCATGC GCCAGCCGTCCGAT **CAGCGC** A<br>TGGTTCTCAG AA **GCTGGC** A CTGAGGACCA ATCGGATGGCTGGC GGTCGGTGCTAAGC<br>**CGCTGG** A ATACCAGGCGAGGTTCCGCAGGATACTGG AA **GCGCAG** A<br>CCAGTATTCTGCGGGACCTCGTCTGGTAT GCTTAGTACCGACC GCATGCGCCTGG **TTCG**<br>CCAGGC **TT** **TTAC** **CACGAAGTCAATAC** **GTCCAACC** **TT** **CATGCTTACGACG** | 796 |
| **UR4B-Gene**<br>Template for UR4B-Brick1–4 | **GTTCTAATACGACTCACTATA** GGGCCTGC **CGCTGG** A TCCACGTCCTCGGG AA **CGGCTG** A<br>CCCGAGGGCGTGGA GCAGGTCC GCTGTAAC **CCGCTC** A GGGACATGTCCGGG AA **CGCGTG** A<br>CCCGGACGTGTCCC GTTACAGC **TT** **TTAT** **CACGAAGTCAATAC** CCAGGATGTCTGG **CCTCGC**<br>A ACTAGGGCTACAGG AA **CCAGCG** A CCTGTAGTCCTAGT CCAGACGTCCTGG<br>CCGTAGGTATCCGT **GGGACC** A TGTATCGCGACCTG AA **GAGCGG** A CAGGTCGTGATACA<br>ACGGATGCCTACGG **TT** **TTAA** **CACGAAGTCAATAC** GCCTTCGGCTGTTCCGCTC **CGGGAG** A<br>TGCGTGTATCCTTG AA **GCGAGG** A CAAGGATGCACGCA GAGCGGGACAGCTGAAGGC<br>GCTTCTGGAACGTCCATGC **GACGGG** A TCCTAGTCCTCGCG AA **GGTCCC** A<br>CGCGAGGGCTAGGA GCATGGGCGTTCTAGAAGC **TT** **TTAC** **CACGAAGTCAATAC**<br>CCAGACTGACAGC **CAGCCG** A GGCTACGGCCTCAG AA **CTCCCG** A CTGAGGCTGTAGCC<br>GCTGTCGGTCTGG CCTTCGATCGGTCC **CACGCG** A TGGGTAGGTCCGGG AA **CCCGTC** A<br>CCCGGACTTACCCA GGACCGGTCGAAGG **TT** **TATT** **CACGAAGTCAATAC** **GTCCAACC** **TT**<br>**CATGCTTACGACG** | 692 |
| **UA4B-Gene**<br>Template for UA4B-Brick1–4 | **GTTCTAATACGACTCACTATA** GGGCTCTG **CGCTGG** A TCCAGTTGAGCGGACCTATCCTGGG AA<br>**CGGCTG** A CCCAGGGTAGGTTCGCTCGACTGGA CAGAGTCC GCTCGCAC **CCGCTC** A<br>TCTGGCTGACCGGG AA **CGCGTG** A CCCGGTCGGCCAGA GTGCGAGC **TT** **TTAT**<br>**CACGAAGTCAATAC** CCTCTGTCTGAGT **CCTCGC** A ACCGTGGAGTGTTAGGACGGATGTG AA<br>**CCAGCG** A CACATCTGTCCTGACACTTCACGGT ACTCAGGCAGAGG CCATCGCTTCGGCT<br>**GGGACC** A TCGTGCGCAGCATG AA **GAGCGG** A CATGCTGTGCACGA AGCCGAGGCGATGG **TT**<br>**TTAA** **CACGAAGTCAATAC** GCAGTAGGGACGTACCCTG **CGGGAG** A<br>TCCGTTTCGTGAGCCTACTGTCGCG AA **GCGAGG** A CGCGACGGTAGGTTCACGGAACGGA<br>CAGGGTGCGTCCTTACTGC GCTCGAGGCAGATGAGTGC **GACGGG** A CTCGTCTGGGAGCG AA<br>**GGTCCC** A CGCTCCCGGACGAG GCACTCGTCTGCTTCGAGC **TT** **TTAC** **CACGAAGTCAATAC**<br>CCGATGTCCATGC **CAGCCG** A TCTAGAGCTGCGTAGTGTGCCAGTG AA **CTCCCG** A<br>CACTGGTACACTGCGCAGTTCTAGA GCATGGGCATCGG CCTACTGTACAGGG **CACGCG** A<br>TCTGACGCCTCGGG AA **CCCGTC** A CCCGAGGTGTCAGA CCCTGTGCAGTAGG **TT** **TATT**<br>**CACGAAGTCAATAC** **GTCCAACC** **TT** **CATGCTTACGACG** | 780 |
| **Z-Brick-7A-GENE**<br>Strut 13+8 bp; beam 14 bp + A | **GTTCTAATACGACTCACTATA** GGATACTTGTCAG **GAGATCC** CTGCTGGACTCTCG AAA<br>**GGATCTC** A CGAGAGTTCAGCAG CTGACAGGTATCC GGTTGGAC **GTCACAG**<br>CTGACTTGATCACG AAA **CTGTGAC** A CGTGATCGAGTCAG **GTCCAACC** **TT**<br>**CATGCTTACGACG** | 170 |
| **Z-Brick-7B-GENE**<br>Strut 13+8 bp; beam 15 bp | **GTTCTAATACGACTCACTATA** GGATACTTGTCAG **GAGATCC** CTTGCTGGACTCTCG AA<br>**GGATCTC** CGAGAGTTCAGCAAG CTGACAGGTATCC GGTTGGAC **GTCACAG**<br>CCTGACTTGATCACG AA **CTGTGAC** CGTGATCGAGTCAGG **GTCCAACC** **TT** **CATGCTTACGACG** | 170 |
| **Z-Brick-8A-GENE**<br>Strut 13+8 bp; beam 13 bp + A | **GTTCTAATACGACTCACTATA** GGATACTTGTCAG **GAGATCAC** CTGCTGGACTCTG AAA<br>**GTGATCTC** A CAGAGTTCAGCAG CTGACAGGTATCC GGTTGGAC **GTACACAG**<br>CTGACTTGATCCG AAA **CTGTGTAC** A CGGATCGAGTCAG **GTCCAACC** **TT** **CATGCTTACGACG** | 170 |
| **Z-Brick-8B-GENE**<br>Strut 13+8 bp; beam 14 bp | **GTTCTAATACGACTCACTATA** GGATACTTGTCAG **GAGATCAC** CTGCTGGACTCTCG AA<br>**GTGATCTC** CGAGAGTTCAGCAG CTGACAGGTATCC GGTTGGAC **GTACACAG**<br>CTGACTTGATCACG AA **CTGTGTAC** CGTGATCGAGTCAG **GTCCAACC** **TT** **CATGCTTACGACG** | 170 |
| **Z-Brick-9A-GENE**<br>Strut A+12+8+A bp; beam 12 bp + A | **GTTCTAATACGACTCACTATA** GGATCGTGTTACA **CATTCAGAG** CCGGAGTATGCG AAAA<br>**CTCTGAATG** A CGCATATTCCGG AGTAACGCGATCC GGTTGGACA **GTTACCTAG**<br>GCTCGTGACCCG AAAA **CTAGGTAAC** A CGGGTCGCGAGT A**GTCCAACC** **TT**<br>**CATGCTTACGACG** | 174 |

| Name (Note) | Sequence | length |
|---|---|---|
| **Z-Brick-9B-GENE** Strut A+12+8+A bp; beam 13 bp | GTTCTAATACGACTCACTATA GGATAATGTCGCA CATTCAGAG GCTCAGGGCCTGG AAA<br>CTCTGAATG CCAGGCTCTGAGC AGCGACGTTATCC GGTTGGACA GTTACCTAG<br>GCTCGGTGACCGG AAA CTAGGTAAC CCGGTCGCCGAGT AGTCCAACC TT CATGCTTACGACG | 174 |
| **LR3B-TMP** Template for LR3B-Brick1–3 | GTTCTAATACGACTCACTATA GGGCACGGACGAGC A GCTTCAATC GGATCGCATCATCAGGTG<br>AAA GAAGGTATC CACCTGGTGATGTGATCC A GCTCGTCTGTGCTC CCAGGTGAGGC A<br>CTACTCGTA CCCGTCTTGACGAAATGATCACTTAGTTAGTTGATCTAGGTTGCGG AAA<br>TACGAGTAG CCGCAACTTAGATCAGCTAACTAGGTGATCGTTTCGTCGAGACGGG A<br>GCCTCGCCTGG TT AT CACGAAGTCAATAC GCAGTGTCAGCTGC A CCTTGAGAA<br>CAGAAGTACGCAGGAGTGG AAA TTCATAGCG CCACTCTTGCGTGCTTCTG A<br>GCAGCTGGCACTGC GCTCCTGGAGC A GATACCTTC GCGAGATGGCTCTATACTG AAA<br>GATTGAAGC CAGTATGGAGCCGTCTCGC A GCTCCGGGAGC TT AA CACGAAGTCAATAC<br>CCAGCGGA TTCG TCCGCTGG GCCTCCGC A CGCTATGAA CCCGCTCCCAAGCGACGG AAA<br>TTCTCAAGG CCGTCGCTTGGGAGCGGG A GCGGAGGC TT AC CACGAAGTCAATAC<br>GTCCAACC TT CATGCTTACGACG | 589 |

#### 4) Bricks for 2D assembly

**2D_14** (14 bricks): 2D-a + 2D-z.

**2D_34** (34 bricks): 2D-a + 2D-b + 2D-c + 2D-y.

**2D_62** (62 bricks): 2D-a + 2D-b + 2D-c + 2D-d + 2D-e + 2D-f + 2D-x.

**2D_107** (107 bricks): 2D-a + 2D-b + 2D-c + 2D-d + 2D-e + 2D-f + 2D-g + 2D-h + 2D-i + 2D-j + 2D-k + 2D-l.

| Name (Note) | Sequence | length |
|---|---|---|
| **2D-a-Tile0** | GGGCUAUGACACGC A CUUCAAGAC AGCGGUAUG GUAA CAUACCGCU A GCGUGUCGUAGCUC<br>GCUCGUACCGG A CUGCUAACC UCCGCUGCU UUCG AGCAGCGGA A CCGGUGCGAGC UU AU<br>CACGAAGUCAAUAC | 134 |
| **2D-a-Tile1** | CCUGAGGGCUAGGC A GCUUCAAUC GGGUCGCUCGCUUCACAG AAA UAGAGCAAC<br>CUGUGAGGCGAGUGACCC A GCCUAGCUCUCAGG GCUCGUGGACC A CUACUCGUA<br>CACGGCAGGGACCUCGCCUACAUCUAGUCGUUGACCUUUACUGGUG AAA GGUUAGCAG<br>CACCAGUGAAGGUCAGCGACUAGGUGUAGGUGAGGUCCUUGCCGUG A GGUCCGCGAGC UU AA<br>CACGAAGUCAAUAC | 242 |
| **2D-a-Tile9** | CCUACAGUGGACCC A UAGUUGAGC GGUCCCGUCUGCGGACGUG AAA GCUUGUUAC<br>CACGUCUGCAGAUGGGACC A GGGUCCAUUGUAGG GCUGGUGAGCC A GUUGCUCUA<br>CCGCUUGCACCGUGAGGUG AAA GUCUUGAAG CACCUCGCGGUGUAAGCGG A GGCUCGCCAGC<br>UU AC CACGAAGUCAAUAC | 190 |
| **2D-a-Tile10** | GCAGUGUCAGCUGC A CCUUGAGAA CAGAAGUACGCAGGAGUGG AAA UUCAUAGCG<br>CCACUCUUGCGUGCUUCUG A GCAGCUGGCACUGC GCUCCUGGAGC A GAUACCUUC<br>GCGAGAUGGCUCUAUACUG AAA GAUUGAAGC CAGUAUGGAGCCGUCUCGC A GCUCCGGGAGC<br>UU UA CACGAAGUCAAUAC | 190 |
| **2D-a-Tile18** | GCCUACUUCUGCGC A GACAGUUGA CCUAAGUCA GUAA UGACUUAGG A GCGCAGAGGUAGGC<br>GCUCCUGCAGG A GUAACAAGC CCAGCCGGU UUCG ACCGGCUGG A CCUGCGGGAGC UU<br>UAAU CACGAAGUCAAUAC | 136 |
| **2D-a-Tile19** | CCAUGUGCACCUGG A CUUUGCUCA CGGCCUAGGUAGGACCGG AAA CAAUCGACA<br>CCGGUCCUACCUAGGCCG A CCAGGUGUACAUGG GCACUUAUCGG A CGCUAUGAA<br>CCGCCUGUCACUAUCGUG AAA GCUCAACUA CACGAUAGUGACAGGCGG A CCGAUGAGUGC UU<br>UACU CACGAAGUCAAUAC | 188 |
| **2D-a-Tile20** | GCUACCUAGAGUGC A UGUAUCCUC GCCAGGGCGUCUGCGGUG AAA GAAUGGCUA<br>CACCGCAGACGCCUUGGC A GCACUCUGGGUAGC GCAUGUGUCGC A UCACUACAG<br>CCCGAGGGCUCUGAGUGG AAA UUCUCAAGG CCACUCAGAGCCCUCGGG A GCGACGCAUGC UU<br>AUAU CACGAAGUCAAUAC | 188 |
| **2D-a-Tile28** | GCCUCGUGUUACGG A CUAACUAGC CCGUCUUGUCUGGGUCGUG AAA GAAGUAUGC<br>CACGACUCAGACGAGACGG A CCGUAACGCGAGGC GCUGUUGCAGC A UAGCCAUUC<br>GAGUGAUUCGAAUGGUCGG AAA UGAGCAAAG CCGACCGUUCGAGUCACUC A GCUGCGACAGC<br>UU AAUU CACGAAGUCAAUAC | 192 |

| Name (Note) | Sequence | length |
|---|---|---|
| **2D-z-Tile2** | GGGCCUGGGCGACU A **GUCUCAUGA** ACCGGUACCGCUUGCGUG AAA **GAAGGUAUC**<br>CACGCAGGCGGUGCCGGU A AGUCGCCUAGGCUC GCUACCUGCAGG **AAUAAUA**<br>CGUCGUCGGUUCUGAGAUCGAGUGAUGUUAUCUACAUCGGUGACGG AAA **UACGAGUAG**<br>CCGUCACUGAUGUAGGUAACAUCGCUCGAUUUCAGAACUGACGACG CCUGCGGGUAGC **UU AU**<br>**CACGAAGUCAAUAC** | 240 |
| **2D-z-Tile11** | CCUAGUGGGCCUAGC **AAAUAAA** GCUACGGGACACUGCUCGG AAA **CUGUAGUGA**<br>CCGAGCGGUGUCUCGUAGC GCUAGGCUCACUAGG GCCUCUGCGACC **AAUAAUA**<br>GCCAGUUCACGAGCUGCGG AAA **UCAUGAGAC** CCGCAGUUCGUGGACUGGC GGUCGUAGAGGC<br>**UU AA CACGAAGUCAAUAC** | 186 |
| **2D-z-Tile27** | CCAGCUUCGAGGGC A **UCGAUAGAG** GCCGACACA **GUAA** UGUGUCGGC A GCCCUCGGAGCUGG<br>GCCUGGUCGGC A **UGUCGAUUG** CCAGGUGCAGCAGCCGGUG AAA **UCAACUGUC**<br>CACCGGUUGCUGUACCUGG A GCCGAUCAGGC **UU AC CACGAAGUCAAUAC** | 162 |
| **2D-z-Tile29** | CCAGUAGUGCCACGG **AAAUAAA** GCAGAAUCUUGGGCCUCGC AAA **GCAAUCAUG**<br>GCGAGGUCCAAGGUUCUGC CCGUGGCGCUACUGG GCGUUAUGUAGG **AAUAAUA**<br>GCAGCGUACAUGGUUAGCG AAA **GAGGAUACA** CGCUAAUCAUGUGCGCUGC CCUACGUAACGC<br>**UU UA CACGAAGUCAAUAC** | 186 |
| **2D-z-Tile37** | CCUCGAGC **UUCG** GCUCGAGG GCUCCGGC A **GCAUACUUC** ACCGAGGCAGAGUACCGG AAA<br>**CUCUAUCGA** CCGGUACUCUGCCUCGGU A GCCGGAGC **UU UAAU CACGAAGUCAAUAC** | 115 |
| **2D-z-Tile38** | GCAGGGUC **UUCG** GACCCUGC GCCUCCGG A **CAUGAUUGC** CCGUAGGGUCCUGAGGGC AAA<br>**GCUAGUUAG** GCCCUCAGGACCCUACGG A CCGGAGGC **UU UACU CACGAAGUCAAUAC** | 115 |
| **2D-b-Tile2** | GGGCGAGUCUGACU A **GUCUCAUGA** ACAGCUAGCCUGGAGGUG AAA **GAAGGUAUC**<br>CACCUCUAGGCUGGCUGU A AGUCAGAUUCGCUC GCCUCUGAGCC A **UUGCAUUCG**<br>GGCUCCCGUAGACCUGUCCACUGUGAGCAUGACAGCCAUCUACCGG AAA **UACGAGUAG**<br>CCGGUAGGUGGCUGUUAUGCUCAUAGUGGAUAGGUCUAUGGGAGCC A GGCUCGGAGGC **UU AU**<br>**CACGAAGUCAAUAC** | 242 |
| **2D-b-Tile3** | CCACUCUUACAUCC A **UCAGGUUAC** GCAGCGCAGCUUGGGCUG AAA **GUGUACCUA**<br>CAGCCCGAGCUGUGCUGC A GGAUGUAGGAGUGG GCUCCGGACGG A **UUCGACAUG**<br>CCUGCGGUACGUGUCUGUCGUGGUACAUACUUCAAGGUGUCACCGG AAA **CGAAUGCAA**<br>CCGGUGAUACCUUGAGGUAUGUAUCACGACGGACACGUGCCGCAGG A CCGUCUGGAGC **UU AA**<br>**CACGAAGUCAAUAC** | 242 |
| **2D-b-Tile11** | CCAUCGUGAGUAGC A **GCAAUCUUC** GUCACGUCUCCGUGACCGG AAA **CUGUAGUGA**<br>CCGGUCGCGGAGGCGUGAC A GCUACUCGCGAUGG GCCUCGCGUCC A **UAGGUACAC**<br>GGAGCAUCGGAGUCCAGUG AAA **UCAUGAGAC** CACUGGGCUCCGGUGCUCC A GGACGUGAGGC<br>**UU AC CACGAAGUCAAUAC** | 190 |
| **2D-b-Tile12** | GCCAAGUGGACUGC A **GACUACCUA** CCGAGCGGUACUGGCGGUG AAA **GCUGAUUGA**<br>CACCGCUAGUACUGCUCGG A GCAGUCCGCUUGGC GCUCCGCAGCC A **GAGAACAUG**<br>GCAUUAUGCUUGUCCUGUG AAA **GUAACCUGA** CACAGGGCAAGCGUAAUGC A GGCUGUGGAGC<br>**UU UA CACGAAGUCAAUAC** | 190 |
| **2D-b-Tile21** | CCAUCCUACUUGGC A **UGCAGAUUC** GACCCUAUCACUCCGGUG AAA **GGUUACAUG**<br>CACCGGAGUGAUAGGGUC A GCCAAGUGGGAUGG GCAUAUGGUCC A **UCAAUCAGC**<br>ACCGAGGUGAAGGUCGUG AAA **GAAGAUUGC** CACGACCUUCACCUCGGU A GGACCGUAUGC **UU**<br>**UAAU CACGAAGUCAAUAC** | 188 |
| **2D-b-Tile22** | GGUCCAGGCUCGGG A **GUGCAUUGA** UCGACUACGCCUGCCUGG AAA **UACAUGUGG**<br>CCAGGCAGGCGUAGUCGA A CCCGAGCUUGGACC GCUACUAGUGG A **CUUAGCUUC**<br>AGGUCUCAAACUCAGGUG AAA **UAGGUAGUC** CACCUGAGUUUGAGACCU A CCACUGGUAGC **UU**<br>**UACU CACGAAGUCAAUAC** | 188 |
| **2D-b-Tile29** | CCUAUGGAGCAAGC A **CAUACAUGG** CCGAUUGAGGCGGUCUCGC AAA **GCAAUCAUG**<br>GCGAGAUCGCCUUAAUCGG A GCUUGCUUCAUAGG GCUCCGCUGAG A **CAUGUAACC**<br>GCGUGGGCCUAGGGCUCUG AAA **GAGGAUACA** CAGAGCUCUAGGUCCACGC A CUCAGUGGAGC<br>**UU AUAU CACGAAGUCAAUAC** | 192 |
| **2D-b-Tile30** | GCAGUCGAGAGCCC A **CUUCUGACA** GGCCAAGGCGUGGGCUCGG AAA **UACAGUCAC**<br>CCGAGCUCACGCUUUGGCC A GGGCUCUUGACUGC GCUGGUACCGG A **CCACAUGUA**<br>CGUACGGAAUACUGGUCUG AAA **GAAUCUGCA** CAGACCGGUAUUUCGUACG A CCGGUGCCAGC<br>**UU AAUU CACGAAGUCAAUAC** | 192 |
| **2D-c-Tile27** | GGGCAAUGAGUACC A **UCGAUAGAG** GCAGCCGGAUCCGGACGGC AAA **CAUUCGUAC**<br>GCCGUCUGGAUCUGGCUGC A GGUACUCGUUGUCC GCUAGUCAUGC A **UGUCGAUUG**<br>CCAAUGUCAAUUGCCGGUG AAA **UCAACUGUC** CACCGGUAAUUGGCAUUGG A GCAUGGCUAGC<br>**UU AU CACGAAGUCAAUAC** | 190 |

| Name (Note) | Sequence | length |
|---|---|---|
| **2D-c-Tile36** | GCCUAAUGUCCUGG A **GGCUAUGUA** CCGCUUCGG **GUAA** CCGAAGCGG A CCAGGACGUUAGGC GCAGCUGCUCC A **GUACGAAUG** GAAGCUGCG **UUCG** CGCAGCUUC A GGAGCGGCUGC **UU AA CACGAAGUCAAUAC** | 134 |
| **2D-c-Tile37** | CAGAUGUCGCUAGG A **UAGCAGUAG** UCCGAGCCGAUUGGGACG AAA **GAGUCAUGA** CGUCCCAGUCGGCUCGGA A CCUAGCGGCAUCUG GCGACUACGGC A **GCAUACUUC** ACCGCUCCACUUUACGUG AAA **CUCUAUCGA** CACGUAAGGUGGAGCGGU A GCCGUGGUCGC **UU AC CACGAAGUCAAUAC** | 186 |
| **2D-c-Tile38** | GGUGUAUUCCAAGC A **UUCCGAAAG** CGCGAGUCAUCUGGAGCG AAA **GAAGUCUCA** CGCUCCAGAUGACUCGCG A GCUUGGAGUACACC GCGACGCCUGG A **CAUGAUUGC** CUCACUCGGGAGCGUCCG AAA **GCUAGUUAG** CGGACGCUCCCGAGUGAG A CCAGGUGUCGC **UU UA CACGAAGUCAAUAC** | 186 |
| **2D-c-Tile39** | CCAUCAGGUCGAGG A **UAGGACUAG** ACGUUUCUAUCUGCGUGG AAA **CUCAAUGGA** CCACGCAGAUAGAGACGU A CCUCGACUUGAUGG GCCUAUCGACC A **GUGACUGUA** GCCGAGCAUGAGGCCUGG AAA **CCAUGUAUG** CCAGGCCUCAUGCUCGGU A GGUCGGUAGGC **UU UAAU CACGAAGUCAAUAC** | 188 |
| **2D-c-Tile40** | GCAUGAGAGUCGGC A **GGAUCUUGA** CCGUAGGGUCUUGGUCGG AAA **CAGUCUUAG** CCGACCAGGACCCUACGG A GCCGACUUUCAUGC GCUUGGAUCGC A **GAUGACUUG** ACCGAGCAUGCUACGUGG AAA **UGUCAGAAG** CCACGUAGCAUGCUCGGU A GCGAUUCAAGC **UU UACU CACGAAGUCAAUAC** | 188 |
| **2D-c-Tile46** | GCCUGCGGACUGGC A **CACUGUUCA** CCCGACGCUCUCGGACGUG AAA **CAUCUAUGC** CACGUCUGAGAGUGUCGGG A GCCAGUCUGCAGGC GCUCGGCCAGG A **UGAGACUUC** GCCUAAGCGUGCUGGUCGG AAA **CUACUGCUA** CCGACCGGCACGUUUAGGC A CCUGGUCGAGC **UU AUAU CACGAAGUCAAUAC** | 192 |
| **2D-c-Tile47** | CAGCUCUCGAUCGG A **UACUUAGCG** CGAUACGGUCGGUGACGCG AAA **CAAUGUCUC** CGCGUCGCCGACUGUAUCG A CCGAUCGGGAGCUG GCGGAGCGUAG A **UCCAUUGAG** ACGCCUUGUAAUGCCGAGC AAA **CUUUCGGAA** GCUCGGUAUUACGAGGCGU A CUACGUUCCGC **UU AAUU CACGAAGUCAAUAC** | 192 |
| **2D-c-Tile48** | GCCUAGUACGUGGC A **UAGUACAGG** GCCAUCUAGCCAGGGCGUG AAA **GAUCAUGUG** CACGCCUUGGCUGGAUGGC A GCCACGUGCUAGGC GCUCGGCACGG A **CUAAGACUG** GCUCCGGUCUUAGCCGGUG AAA **CUAGUCCUA** CACCGGUUAAGAUCGGAGC A CCGUGUCGAGC **UU ACUU CACGAAGUCAAUAC** | 192 |
| **2D-y-Tile4** | GGGCCAUGACCUGC A **CUUGGAUCA** CCCGUGCGUCGGGACGGC AAA **CAUGUUCUC** GCCGUCUCGACGUACGGG A GCAGGUCGUGGCUC CCAGUCGCGGUG **AAUAAUA** CCCAGUAUGUUUCUGGACGAUCUCGAAUGCUGUACAUAUGCACCGG AAA **CAUGUCGAA** CCGGUGCGUAUGUACGGCAUUCGGGAUCGUUCAGAAACGUACUGGG CACCGUGACUGG **UU AU CACGAAGUCAAUAC** | 240 |
| **2D-y-Tile13** | CCUUAGGGCACGCUG **AAAUAAA** GCCCUUGCCGGAUCCGGUG AAA **GAAGCUAAG** CACCGGGUCCGGUAAGGGC CAGCGUGUCCUAAGG GCGUAUUUCAGG **AAUAAUA** GCUAGAUCAAGCGGUCGUG AAA **UGAUCCAAG** CACGACUGCUUGGUCUAGC CCUGAGAUACGC **UU AA CACGAAGUCAAUAC** | 186 |
| **2D-y-Tile31** | CCGACGGUCAAGUCC **AAAUAAA** GGUGGAUCGCCUGGCGUCG AAA **CAAGUCAUC** CGACGCUAGGCGGUCCACC GGACUUGGCCGUCGG CCUUGCUGUACG **AAUAAUA** GUCCGUUUAUGCGUUCCGG AAA **UCAAUGCAC** CCGGAAUGCAUAGACGGAC CGUACGGCAAGG **UU AC CACGAAGUCAAUAC** | 186 |
| **2D-y-Tile45** | CCAGGCGGACAGGG A **CACUUCAGA** GCCGAUACG **GUAA** CGUAUCGGC A CCCUGUCUGCCUGG GCUACUGAGGC A **UCAUGACUC** GCCAGUUUGAAGUGCAGUG AAA **UACAUAGCC** CACUGCGCUUCAGACUGGC A GCCUCGGUAGC **UU UA CACGAAGUCAAUAC** | 162 |
| **2D-y-Tile49** | CCAGCAUGGAUUGGC **AAAUAAA** GCUCAAGUUCACGGACACG AAA **UAGGUCAAG** CGUGUCUGUGAAUUUGAGC GCCAAUCUAUGCUGG CCUUGGGCACCG **AAUAAUA** GGAUUCUCAAGGUCAUCUG AAA **UCAAGAUCC** CAGAUGGCCUUGGGAAUCC CGGUGUCCAAGG **UU UAAU CACGAAGUCAAUAC** | 188 |
| **2D-y-Tile55** | CCUCGGUC **UUCG** GACCGAGG GCUCCCGG A **GCAUAGAUG** ACCGUUCGUCAGCCGGUG AAA **UCUGAAGUG** CACCGGCUGACGAGCGGU A CCGGGAGC **UU UACU CACGAAGUCAAUAC** | 115 |
| **2D-y-Tile56** | GCCUCACC **UUCG** GGUGAGGC CCUCGGCC A **GAGACAUUG** CCACCUCGUGAGAUGCGG AAA **UGAACAGUG** CCGCAUCUCACGAGGUGG A GGCCGAGG **UU AUAU CACGAAGUCAAUAC** | 115 |
| **2D-y-Tile57** | CCGAUCGC **UUCG** GCGAUCGG GCACCGGG A **CACAUGAUC** UCCACUGGAUGGUCGGUG AAA **CGCUAAGUA** CACCGACUAUCCGGUGGA A CCCGGUGC **UU AAUU CACGAAGUCAAUAC** | 115 |

| Name (Note) | Sequence | length |
|---|---|---|
| **2D-y-Tile58** | GCCAGGUC UUCG GACCUGGC GCACCGCG A **CUUGACCUA** CCCGAGGUAGGGUCGCGG AAA **CCUGUACUA** CCGCGACUCUACCUCGGG A CGCGGUGC **UU ACUU CACGAAGUCAAUAC** | 115 |
| **2D-d-Tile4** | GGGCUCUGCACUGC A **CUUGGAUCA** CUGCCUACGACGGACCGC AAA **CAUGUUCUC** GCGGUCUGUCGUGGGCAG A GCAGUGCGGAGUCC GCUCCUGAGCC A **GUAUCUCCA** UCGCGGAUGUUGCCGGUCACAGUCUAAUACUUCAGAUUUGCGCUGG AAA **CAUGUCGAA** CCAGCGCGAAUCUGAGGUAUUAGGCUGUGAUCGGCAACGUCCGCGA A GGCUCGGGAGC **UU AU CACGAAGUCAAUAC** | 242 |
| **2D-d-Tile5** | CCGUCCUGACUCGC A **CGAUCUUAC** CCGUCGGCUGGUCUGCUG AAA **GUAAGGACA** CAGCAGGCCAGCUGACGG A GCGAGUCGGGACGG CCUACGGCUGG A **CAGAACUCA** CCCUGGCGAGCAUCGUCUCGUAUGACGAUUGACCAGGUUAUGCUGG AAA **UGGAGAUAC** CCAGCAUGACCUGGUUAAUCGUCGUACGAGGCGAUGCUUGCCAGGG A CCAGCUGUAGG **UU AA CACGAAGUCAAUAC** | 242 |
| **2D-d-Tile13** | CCGAGCGAUCGGGC A **CCUUGAUUC** GCGUUGGAUGUAGGCGGUG AAA **GAAGCUAAG** CACCGCUUACAUUCAACGC A GCCCGAUUGCUCGG CCAGCUGUAGG A **UGUCCUUAC** CGCGAGUUAACGGGACCGG AAA **UGAUCCAAG** CCGGUCUCGUUAGCUCGCG A CCUACGGCUGG **UU AC CACGAAGUCAAUAC** | 190 |
| **2D-d-Tile14** | GCCUAGUAGACAGG A **GUUGCUAGA** GCCUACUGAGAUGGCGGUG AAA **GAAGACGUA** CACCGCUAUCUCGGUAGGC A CCUGUCUGCUAGGC GCUCCGGGAGC A **GGUACUACA** GCGUCUGGAUUUGCCCGUG AAA **GUAAGAUCG** CACGGGUAAAUCUAGACGC A GCUCCUGGAGC **UU UA CACGAAGUCAAUAC** | 190 |
| **2D-d-Tile23** | CCAUGUUUCAGGCC A **CAUUGCAUC** UCGGUUGAAACUGCGGUG AAA **UCAGUACUC** CACCGCAGUUUCAGCCGA A GGCCUGAGACAUGG CCUUGUCCAGC A **UACGUCUUC** GGGUCUCCACUUAGAGUG AAA **GAAUCAAGG** CACUCUAGGUGGAGACCC A GCUGGGCAAGG **UU UAAU CACGAAGUCAAUAC** | 188 |
| **2D-d-Tile24** | GAGCUAUAUCUGCC A **UUCCACUUG** GCUCGGCCAUAGGUCCGG AAA **CUAACUCUG** CCGGACCUAUGGCUGAGC A GGCAGAUGUAGCUC CCAUGGUAGGG A **UAGUAACGC** CUGUCUGGACUUAGACAG AAA **UCUAGCAAC** CUGUCUAGGUCCAGACAG A CCCUAUCAUGG **UU UACU CACGAAGUCAAUAC** | 188 |
| **2D-d-Tile31** | CCGUCGUGGAUGGC A **CCUAUGUUC** GCGAGCGGCUCGGGCAGCG AAA **CAAGUCAUC** CGCUGCUCGAGCUGCUCGC A GCCAUCCGCGACGG CCAGGUGAGGC A **GAGUACUGA** GCUCCAUGGUCGGCACCGG AAA **UCAAUGCAC** CCGGUGUCGACCGUGGAGC A GCCUCGCCUGG **UU AUAU CACGAAGUCAAUAC** | 192 |
| **2D-d-Tile32** | GCCUGUGAGGUAGG A **UCAUGCUAG** CCCUAGGUAGGCUGACGUG AAA **GGCAAUGAA** CACGUCGGCCUAUCUAGGG A CCUACCUUACAGGC GCUCCGACGGG A **CAGAGUUAG** GCUCAUGCUGCCGGAGGUG AAA **GAUGCAAUG** CACCUCUGGCAGUAUGAGC A CCCGUUGGAGC **UU AAUU CACGAAGUCAAUAC** | 192 |
| **2D-e-Tile41** | GGGCACGCGACCGC A **GCUACUAUC** GUCCGGCUAUAGCGAGCG AAA **UCGAAUCUC** CGCUCGCUAUAGCUGGAC A GCGGUCGUGUGCUC CCGUGGGACCC A **UUCAUUGCC** GACCAGCUCACUCCGGUG AAA **GAACAUAGG** CACCGGAGUGAGCUGGUC A GGGUCUCACGG **UU AU CACGAAGUCAAUAC** | 186 |
| **2D-e-Tile42** | GCAAGCGGACUAGC A **UGAAUGUCC** GUGCAGAAGCCUGCGGUG AAA **GAUAGAUGC** CACCGCAGGCUUCUGCAC A GCUAGUCUGCUUGC GCGUCUGUUGC A **UGUUCCUAG** UCACAGGGUACUCUGCGG AAA **CUAGCAUGA** CCGCAGAGUACCCUGUGA A GCAACGGACGC **UU AA CACGAAGUCAAUAC** | 186 |
| **2D-e-Tile49** | CCAGGGUCUAAAGC A **UGGAACAAG** GCCUGUGCAACUGCGCGUG AAA **UAGGUCAAG** CACGCGUAGUUGUACAGGC A GCUUUAGGCCCUGG CCAGGUCCAGC A **GAGAUUCGA** GGACACGGGUGCUGGAGUG AAA **UCAAGAUCC** CACUCCGGCACCUGUGUCC A GCUGGGCCUGG **UU AC CACGAAGUCAAUAC** | 190 |
| **2D-e-Tile50** | GCAGUUGAGACCCG A **UGAGCAUUC** CCAGCGGUAAAGGCCUCGG AAA **CAGAUGUAG** CCGAGGUCUUUAUCGCUGG A CGGGUCUUAACUGC GCUCCGGAGCC A **GCAUCUAUC** GGCGUCUAUCCUUGCGGUG AAA **GAUAGUAGC** CACCGCGAGGAUGGACGCC A GGCUCUGGAGC **UU UA CACGAAGUCAAUAC** | 190 |
| **2D-e-Tile57** | CCAGGCGAUUCCGG A **UCAGUGAAG** CCGGUUAUGACUGCGGUG AAA **GAUGUCUGA** CACCGCAGUCAUAGCCGG A CCGGAAUUGCCUGG CCUGGGUCUCG A **CACAUGAUC** UCAGCUCGAAAGCUCCGG AAA **CGCUAAGUA** CCGGAGCUUUCGAGCUGA A CGAGAUCCAGG **UU UAAU CACGAAGUCAAUAC** | 188 |

| Name (Note) | Sequence | length |
|---|---|---|
| **2D-e-Tile58** | GCCAGCGACAGGCC A **GGAUACCAA** GCCGAGGCACUUGGUCGG AAA **UCCUUGUAG** CCGACCAGGUGCCUCGGC A GGCCUGUUGCUGGC GCAGAGACGCG A **CUUGACCUA** CAGCCUGAAAUUCGGCUG AAA **CCUGUACUA** CAGCCGAGUUUCAGGCUG A CGCGUUUCUGC **UU UACU CACGAAGUCAAUAC** | 188 |
| **2D-e-Tile59** | CCAGCAUAUUGAGG A **UAGACAUGC** GGAGCUCCAGGGACCCGC AAA **CUUUCGUCA** GCGGGUCUCUGGAGCUCC A CCUCAAUGUGCUGG CCCUGUACAGC A **CUACAUCUG** CCGCAGGAUCUUGGACGC AAA **CUUGUUCCA** GCGUCCAGGAUCCUGCGG A GCUGUGCAGGG **UU AUAU CACGAAGUCAAUAC** | 188 |
| **2D-e-Tile60** | GGGAACUUGGGCUC A **GGUAAGUAC** GUCAAGCGAGCUGGAGGC AAA **GUAAGCGUA** GCCUCCAGCUCGCUUGAC A GAGCCCAGGUUCCC GCUUGUCCUGC A **CUACGAUCA** CCGGCUCCUACUUGCCCG AAA **GAAUGCUCA** CGGGCAAGUAGGAGCCGG A GCAGGGCAAGC **UU AAUU CACGAAGUCAAUAC** | 188 |
| **2D-f-Tile45** | GGGCAUGACCGGUC A **CACUUCAGA** GGCCCUGUAUGGUGCUGUG AAA **GUUGAUGUC** CACAGCGCCAUAUAGGGCC A GACCGGUUAUGCUC CCGGUUCAGGC A **UCAUGACUC** GGUCUAUCGCGUGCCCGUG AAA **UACAUAGCC** CACGGGUACGCGGUAGACC A GCCUGGACCGG **UU AU CACGAAGUCAAUAC** | 190 |
| **2D-f-Tile54** | GCAGCUGCCAAGCC A **CGAUGAAGA** CCGUGAGUA **GUAA** UACUCACGG A GGCUUGGUAGCUGC GCUCCGUGCCC A **GACAUCAAC** GGCCAAAGG **UUCG** CCUUUGGCC A GGGCAUGGAGC **UU AA CACGAAGUCAAUAC** | 134 |
| **2D-f-Tile55** | CCAUCUUGCCGUGC A **GUACCUUAG** CCCGAGGAUCCUGGACGG AAA **CAUCUUGAC** CCGUCCAGGAUCCUCGGG A GCACGGCGAGAUGG CCUCAGAUCCG A **GCAUAGAUG** ACCCAGGCCUCUGACCGG AAA **UCUGAAGUG** CCGGUCAGAGGCCUGGGU A CGGAUUUGAGG **UU AC CACGAAGUCAAUAC** | 186 |
| **2D-f-Tile56** | GCAGUGUGCAGCCC A **CGAUCUGAA** GCGUCUUUAGAGGUCAUC AAA **UAGAUUCCG** GAUGACCUCUAAAGACGC A GGGCUGCGCACUGC GCACCUUGGCC A **GAGACAUUG** CACGCUGUUCGGCCACGG AAA **UGAACAGUG** CCGUGGCUGAACAGCGUG A GGCCAGGGUGC **UU UA CACGAAGUCAAUAC** | 186 |
| **2D-f-Tile64** | GCCUACGCCUCACC A **UACCGUAAC** CGAGCAUCAGUGGGCUCCG AAA **UCUCGUAUC** CGGAGCUCACUGGUGCUCG A GGUGAGGUGUAGGC GCUGGUCGAGG A **CGGAAUCUA** GGCGACGCGAUCGCCUCUG AAA **CUAAGGUAC** CAGAGGUGAUCGUGUCGCC A CCUCGGCCAGC **UU UAAU CACGAAGUCAAUAC** | 192 |
| **2D-f-Tile65** | CCGUCCUUGCGUGC A **CAGUCAAUC** AGGCCAUAGUUCGUAUCGC AAA **CGUUCAAAG** GCGAUAUGAACUGUGGCCU A GCACGCAGGGACGG CCUCCGGGAGC A **UCAGACAUC** GCUGUCGGAGGCGGAGUGG AAA **UUCAGAUCG** CCACUCUGCCUCUGACAGC A GCUCCUGGAGG **UU UACU CACGAAGUCAAUAC** | 192 |
| **2D-f-Tile66** | GCACCGUCUGGGCC A **CUAGUUACG** CCGAGUGCAGUCGGGACGG AAA **CAGCUAUCA** CCGUCCUGACUGUACUCGG A GGCCCAGGCGGUGC GCUGUUGCGAC A **CUACAAGGA** CCGAGUGGCAAUGCCCUGG AAA **CUUCACUGA** CCAGGGUAUUGCUACUCGG A GUCGCGACAGC **UU AUAU CACGAAGUCAAUAC** | 192 |
| **2D-f-Tile67** | CCAAGUGCUCCGGC A **CCAGUUGUA** CGCGGAGUACAGUGAGCAG AAA **CAAGUCCAA** CUGCUCGCUGUAUUCCGCG A GCCGGAGUACUUGG CCCUCGCCUGC A **UGACGAAAG** CCGUCCGACUCGGCUCGUG AAA **UUGGUAUCC** CACGAGUCGAGUUGGACGG A GCAGGUGAGGG **UU AAUU CACGAAGUCAAUAC** | 192 |
| **2D-f-Tile68** | GCCACUGCGGCAGG A **CUCAGAAUG** CCGCAAGCGUCCGGACCGC AAA **GCAUUCUCA** GCGGUCUGGACGUUUGCGG A CCUGCCGUAGUGGC GCAGGGUCCGG A **UACGCUUAC** CCUUGCUUGCUAUCCGGUG AAA **GCAUGUCUA** CACCGGGUAGCAGGCAAGG A CCGGAUCCUGC **UU ACUU CACGAAGUCAAUAC** | 192 |
| **2D-x-Tile6** | GGGCUUGAGGUACC A **GAUCACUUC** GCAGCUGGUGCUCGUCUG AAA **UGUAGUACC** CAGACGGGCACCGGCUGC A GGUACCUUAAGCUC GCCUCGUGCAGC **AAUAAUA** GCCAGGAUGUGAACGGCACUGCGGAGAUUAUCUGUGCAUAUCACGC AAA **UGAGUUCUG** GCGUGAUGUGCACAGGUAAUCUCUGCAGUGUCGUUCACGUCCUGGC GCUGCGCGAGGC **UU AU CACGAAGUCAAUAC** | 240 |
| **2D-x-Tile15** | CCAGCUUUAUCAUGC **AAAUAAA** GACGUGUAGCUGGGCUGUG AAA **GCGUUACUA** CACAGCUCAGCUGCACGUC GCAUGAUGAAGCUGG CCCUCGUUGCGG **AAUAAUA** GUGACUUCGUAGGCGGACG AAA **GAAGUGAUC** CGUCCGUCUACGGAGUCAC CCGCAGCGAGGG **UU AA CACGAAGUCAAUAC** | 186 |

| Name (Note) | Sequence | length |
|---|---|---|
| **2D-x-Tile33** | CCAGCAUUUAGGUCC **AAAUAAA** GCUGCUGUCGACUGACCCG AAA **CUAGGAACA** CGGGUCGGUCGAUAGCAGC GGACCUAGAUGCUGG CCUAGUUCGCUC **AAUAAUA** GGGUCCGAGGCUUCGACGG AAA **CAAGUGGAA** CCGUCGGAGCCUUGGACCC GAGCGGACUAGG **UU AC CACGAAGUCAAUAC** | 186 |
| **2D-x-Tile51** | CCUCCAGGCAGUCGG **AAAUAAA** CGAGCCUAUAUCUGAGGUG AAA **UGAUCGUAG** CACCUCGGAUAUGGGCUCG CCGACUGUCUGGAGG CCGUUCUUAGCC **AAAUAAA** GGUCGUUAACCGUACAGGC AAA **GGACAUUCA** GCCUGUGCGGUUGACGACC GGCUAGGAACGG **UU UA CACGAAGUCAAUAC** | 186 |
| **2D-x-Tile63** | CACCCUGCGUGAGG A **UCGACUAAC** GCCACUACU **GUAA** AGUAGUGGC A CCUCACGUAGGGUG CCCGAGUCGCC A **GUCAAGAUG** CCGCUGGCUCCGUGAGAGC AAA UCUUCAUCG GCUCUCGCGGAGUCAGCGG A GGCGAUUCGGG **UU UAAU CACGAAGUCAAUAC** | 164 |
| **2D-x-Tile69** | CCUGGACUUCAAUCG **AAAUAAA** GCUUGAUCUACAUGCGGAC AAA **GUUUCACAG** GUCCGCGUGUAGGUCAAGC CGAUUGAGGUCCAGG CCUGCUGGCAGC **AAUAAUA** GGAUCGGGUUCAUAUCCUG AAA **GUACUUACC** CAGGAUGUGAACUCGAUCC GCUGCUAGCAGG **UU UACU CACGAAGUCAAUAC** | 188 |
| **2D-x-Tile73** | CCUCGCGA **UUCG** UCGCGAGG CCCUGGCC A **GAUACGAGA** CCGCAGGUAGCUCCAGGC AAA **GUUAGUCGA** GCCUGGAGCUACCUGCGG A GGCCAGGG **UU AUAU CACGAAGUCAAUAC** | 115 |
| **2D-x-Tile74** | GCUCGGUC **UUCG** GACCGAGC GCUCCGGG A **CUUUGAACG** CCCUAGGGCACUACCCUG AAA **GUUACGGUA** CAGGGUAGUGCCCUAGGG A CCCGGAGC **UU AAUU CACGAAGUCAAUAC** | 115 |
| **2D-x-Tile75** | CCAGGAGC **UUCG** GCUCCUGG CCUCCGGG A **UGAUAGCUG** CCGCAGGGCGAGAGCGUG AAA **GAUUGACUG** CACGCUCUCGCCCUGCGG A CCCGGAGG **UU ACUU CACGAAGUCAAUAC** | 115 |
| **2D-x-Tile76** | GCCAGCAC **UUCG** GUGCUGGC GCUCCCGC A **UUGGACUUG** CCCGAGGUACAGUCUCGG AAA **CGUAACUAG** CCGAGACUGUACCUCGGG A GCGGGAGC **UU UAUA CACGAAGUCAAUAC** | 115 |
| **2D-x-Tile77** | CCAGGGUC **UUCG** GACCCUGG CCUCGCCC A **UGAGAAUGC** GGUGAGCCUACUCAGGUG AAA **UACAACUGG** CACCUGAGUAGGCUCACC A GGGCGAGG **UU AUUA CACGAAGUCAAUAC** | 115 |
| **2D-x-Tile78** | GCAGGAGC **UUCG** GCUCCUGC GCCUCCGC A **CUGUGAAAC** GCUCCUGCCGCUCUGGGC AAA **CAUUCUGAG** GCCCAGAGCGGCAGGAGC A GCGGAGGC **UU CUUA CACGAAGUCAAUAC** | 115 |
| **2D-g-Tile6** | GGGCAUGAUGCUCC A **GAUCACUUC** GCACCUGUCUCUAUCCUG AAA **UGUAGUACC** CAGGAUGGAGACGGGUGC A GGAGCAUUAUGCCC GACCGUACGGG A **CCGAUUCUA** UCCAGGGUUCGGCUGGUAGAUGGACUGAGUUGCUUUGAUCGAGGGC AAA **UGAGUUCUG** GCCCUCGGUCAAAGCGACUCAGUUCAUCUAUCAGCCGAGCCCUGGA A CCCGUGCGGUC **UU AU CACGAAGUCAAUAC** | 242 |
| **2D-g-Tile7** | CCGUACGCGUGAGG A **CGUGAAUGA** GCCGAGGGUCGUUCCGUG AAA **GUAGAUACG** CACGGAGCGACCUUCGGU A CCUCACGUGUACGG CCAGUUCGCCG A **UCCUAUGAG** CACAGCGGUCGUGCUGAUAGCGGUAGAGUCUGAGGUGCGGCAGUGG AAA **UAGAAUCGG** CCACUGCUGCACCUCGGACUCUAUCGCUAUUAGCACGAUCGCUGUG A UGGCGGACUGG **UU AA CACGAAGUCAAUAC** | 242 |
| **2D-g-Tile8** | GCCAAGGGUCACGC A **UCUCGUUAG** CCGGUGCGCUUUGCGCUG AAA **UAGUCUACC** CAGCGCGAAGCGUACCGG A GCGUGACUCUUGGC GCUCUGUGCAGG **AAUAAUA** GUCCUACUGGUUCGGGCAUAUCUCAUAGUAGGACCGUGUCUUGCGG AAA **CUCAUAGGA** CCGCAAGGCACGGUCUUACUAUGGGAUAUGUCCGAACCGGUAGGAC CCUGCGCAGAGC **UU AC CACGAAGUCAAUAC** | 240 |
| **2D-g-Tile15** | CCAGUGUAGCCUCG A **UACAAGUGC** CUGGAUUGACUUGGCGGUG AAA **GCGUUACUA** CACCGCUAAGUCGAUCCAG A CGAGGCUGCACUGG CGUGCUGCAGG A **CGUAUCUAC** CCUGCCUGAGGUGCCGGUG AAA **GAAGUGAUC** CACCGGUACCUCGGGCAGG A CCUGCGGCACG **UU UA CACGAAGUCAAUAC** | 190 |
| **2D-g-Tile16** | GCACGUUCGGAGCC A **GUUCCAAUG** UCGACGGAGUAAGGCGGUG AAA **CAGAGAUGA** CACCGCUUUACUUCGUCGA A GGCUCCGGACGUGC GCUGGUGCUGC A **GGUAGACUA** GCCUACUUGUGCUAGCGUC AAA **UCAUUCACG** GACGCUGGCACAGGUAGGC A GCAGCGCCAGC **UU UAAU CACGAAGUCAAUAC** | 192 |
| **2D-g-Tile17** | CCGUCUCGUGAACGG **AAAUAAA** GUCCACUUCGUUUCCGGUG AAA **GAACUUACG** CACCGGGAACGAGGUGGAC CCGUUCAUGAGACGG CCUGAGGUAGCG **AAUAAUA** GGUUAGUUCCGGGAUCCGG AAA **CUAACGAGA** CCGGAUUCCGGAGCUAACC CGCUAUCUCAGG **UU UACU CACGAAGUCAAUAC** | 188 |
| **2D-g-Tile25** | CCGUCUGAGUUCGC A **CAUCGUUAC** CCUGCUUGGCUUGCGGUG AAA **CUUCUAAGC** CACCGCAGGCCAAGCAGG A GCGAACUUAGACGG CCUCUGCCUGC A **UCAUCUCUG** CCCGAGCGACGGUAGGUG AAA **GCACUUGUA** CACCUACUGUCGCUCGGG A GCAGGUAGAGG **UU AUAU CACGAAGUCAAUAC** | 188 |

| Name (Note) | Sequence | length |
|---|---|---|
| **2D-g-Tile26** | GCAGCUUCACAGGC A **GACUUACUC** GCCAUUUCCGAGGCGGUG AAA **GAACAGAUC** CACCGCCUCGGAAGUGGC A GCCUGUGGAGCUGC GCUGAGCCGAC A **CGUAAGUUC** GCGCAGGGUGCUACACGG AAA **CAUUGGAAC** CCGUGUAGCACCCUGCGC A GUCGGUUCAGC **UU AAUU CACGAAGUCAAUAC** | 188 |
| **2D-h-Tile33** | GGGCAAUUAGUAGC A **UGUUACCUC** CGCUAAGUUGCUUUCGCAG AAA **CUAGGAACA** CUGCGAGAGCAAUUUAGCG A GCUACUAGUUGCUC CCAGGUGUGCA A **GCUUAGAAG** GCAUGCGUCGUCUGGUCGG AAA **CAAGUGGAA** CCGACCGGACGAUGCAUGC A UGCACGCCUGG **UU AU CACGAAGUCAAUAC** | 190 |
| **2D-h-Tile34** | GCCAGUUAGGACGG A **UGACACUAG** CCCUAAUCUGUAGGCGGUG AAA **GUUAGAACG** CACCGCUUACAGGUUAGGG A CCGUCCUGACUGGC GCUCCGGCAGU A **GAUCUGUUC** GCUUAUUGAGCGUGAGGUG AAA **GUAACGAUG** CACCUCGCGCUCGAUAAGC A ACUGCUGGAGC **UU AA CACGAAGUCAAUAC** | 190 |
| **2D-h-Tile35** | CCUAGGCGACCUAGC **AAAUAAA** GGCUACGCAUGCUGCAGCG AAA **GUCUAAGGA** CGCUGCGGCAUGUGUAGCC GCUAGGUUGCCUAGG CCAUACGGAGCG **AAUAAUA** GGAUAGGUGCCUUGUGCCG AAA **GAGUAAGUC** CGGCACGAGGCAUCUAUCC CGCUCUGUAUGG **UU AC CACGAAGUCAAUAC** | 186 |
| **2D-h-Tile43** | CCAGAGGUAUCGCC A **GAUUCAUCC** GGGUAGUAGAGGCUCCCG AAA **GAUGAUCAG** CGGGAGCUUCUACUACCC A GGCGAUAUCUCUGG CCAGGUAGUGC A **CGUUCUAAC** CCGCAGCGGUAGGUGCCG AAA **GAGGUAACA** CGGCACCUACCGCUGCGG A GCACUGCCUGG **UU UA CACGAAGUCAAUAC** | 186 |
| **2D-h-Tile44** | GCAGUGGCGUGUCC A **GUCUGUAAG** CCCGAGCCCACUGAGGCG AAA **GUAGAGUCA** CGCCUCAGUGGGCUCGGG A GGACACGUCACUGC GCACAUACAGG A **UCCUUAGAC** ACCGAGCAGAGGCGUCGG AAA **CUAGUGUCA** CCGACGCUUCUGCUCGGU A CCUGUGUGUGC **UU UAAU CACGAAGUCAAUAC** | 188 |
| **2D-h-Tile51** | CCUAGCGAGUAUCC A **GUGUCAAAC** GCCUGCUGUGAGUGCGUGG AAA **UGAUCGUAG** CCACGCGCUCACGGCAGGC A GGAUACUUGCUAGG CCCUGUGCUGU A **CUGAUCAUC** GCUAUCUACAGCUGGAGGC AAA **GGACAUUCA** GCCUCCGGCUGUGGAUAGC A ACAGCGCAGGG **UU UACU CACGAAGUCAAUAC** | 192 |
| **2D-h-Tile52** | GCCAGUGCCAAGCC A **CUCGAUACA** GUGCCGUUGUAGUAAGUGG AAA **CAACAUACG** CCACUUGCUACAGCGGCAC A GGCUUGGUACUGGC GCUGGUGCAGG A **UGACUCUAC** CCUAAGUUUCGAGCGAGGC AAA **GGAUGAAUC** GCCUCGUUCGAAGCUUAGG A CCUGCGCCAGC **UU AUAU CACGAAGUCAAUAC** | 192 |
| **2D-h-Tile53** | CCAGCGUUCGAACGG **AAAUAAA** GUCCCUGAGCUCUGACGUG AAA **UGAAUGAGC** CACGUCGGAGCUUAGGGAC CCGUUCGGACGCUGG CCAGCAUCCGAG **AAUAAUA** GCGAAGGUUCUGUAGACGG AAA **CUUACAGAC** CCGUCUGCAGAAUCUUCGC CUCGGGUGCUGG **UU AAUU CACGAAGUCAAUAC** | 188 |
| **2D-h-Tile61** | CCAGUAUGCAACCG A **UGACGAUUC** GGCUAGGUUCAGGUGGCG AAA **GGAACUAUC** CGCCACCUGAACCUAGCC A CGGUUGCGUACUGG CGCUGUAAUCC A **CGUAUGUUG** CCAGCUCUUGGGCGUCCG AAA **GUUUGACAC** CGGACGCUCAAGAGCUGG A GGAUUGCAGCG **UU ACUU CACGAAGUCAAUAC** | 188 |
| **2D-i-Tile62** | GGGCAGUCCGACCC A **UACCAGUUC** AGCGCUUACGAGGGCUCG AAA **CAAUGAUGG** CGAGCCCUCGUAAGCGCU A GGGUCGGGCUGUCC GGUGUGCCGCU A **GCUCAUUCA** GGGACUGAUCUUGACCGG AAA **UGUAUCGAG** CCGGUCAGGAUCAGUCCC A AGCGGUACACC **UU AU CACGAAGUCAAUAC** | 186 |
| **2D-i-Tile69** | CCAGGGUCAGCGGC A **CAUACCUUG** GCCUCAUUCGCAUGCUCGC AAA **GUUUCACAG** GCGAGCGUGCGAGUGAGGC A GCCGCUGGCCCUGG CCAGCGGGUGC A **GAUAGUUCC** GCAUGGGUAGACUGGUCUG AAA **GUACUUACC** CAGACCGGUCUAUCCAUGC A GCACCUGCUGG **UU AA CACGAAGUCAAUAC** | 190 |
| **2D-i-Tile70** | GCUCCUGUAGUCCC A **UCGAGAUAG** GCCACUGCGGCUUGCGGUG AAA **GCUUGAUGA** CACCGCGAGCCGUAGUGGC A GGGACUAUAGGAGC GCUGCUAGCCU A **CCAUCAUUG** CGUCAAUGCUUGUCCUGCG AAA **GAAUCGUCA** CGCAGGGCAAGCGUUGACG A AGGCUGGCAGC **UU AC CACGAAGUCAAUAC** | 190 |
| **2D-i-Tile71** | CCACCGUGACUGAGC **AAAUAAA** GUCACUUGCCACGGACCUG AAA **CUUACUUGC** CAGGUCUGUGGCGAGUGAC GCUCAGUUACGGUGG CCUGCAUCGGUC **AAUAAUA** GCUCUGUUCCGCGUGGCUG AAA **GAACUGGUA** CAGCCAUGCGGAGCAGAGC GACCGGUGCAGG **UU UA CACGAAGUCAAUAC** | 186 |

| Name (Note) | Sequence | length |
|---|---|---|
| **2D-i-Tile80** | GGGUGAGCUAACGC A GACAUGGAA CCGCAUCUAACUCGAGUG AAA GAUCUAACG CACUCGGGUUAGGUGCGG A GCGUUAGUUCACCC GCUCCUACCGC A GCAAGUAAG UCCAGUCGUCAUUCGCGG AAA CUAUCUCGA CCGCGAGUGACGGCUGGA A GCGGUGGGAGC UU UAAU CACGAAGUCAAUAC | 188 |
| **2D-i-Tile86** | GCCACAUCGACGCC A GUAUGCAGA GCCUUCUGGACAUGCCUCG AAA CUAUGCAAG CGAGGCGUGUCCGGAAGGC A GGCGUCGGUGUGGC GCUGCUCAGGU A GGUACAAAG GGAAUGUAAUACUGACCGG AAA UCCUAACUG CCGGUCGGUAUUGCAUUCC A ACCUGGGCAGC UU UACU CACGAAGUCAAUAC | 192 |
| **2D-i-Tile87** | CCUACCGUGUGCUC A CUAUGCCUA GGACUCUCGAUGUGACCGG AAA CUGUCUACA CCGGUCGCAUCGGGAGUCC A GAGCACAUGGUAGG CCCUGUCGCUC A GACUCAUUG CGUCCUGCGCCAUCGGGAC AAA GAUUCUCAG GUCCCGGUGGCGUAGGACG A GAGCGGCAGGG UU AUAU CACGAAGUCAAUAC | 192 |
| **2D-i-Tile88** | GCAGGUUCAUUACC A UAGUUCGUC GGUAAGGGCGUCGCACGUG AAA UUCUGAUCC CACGUGUGACGCUCUUACC A GGUAAUGGACCUGC GCAGGGCCAGG A CGUUAGAUC GCUACAGUCACAUGGCGUG AAA GACUAAGUG CACGCCGUGUGAUUGUAGC A CCUGGUCCUGC UU AAUU CACGAAGUCAAUAC | 192 |
| **2D-i-Tile89** | CCUCGGUGGUACGAC AAAUAAA CCUCGUUCAUCCGGCUCGC AAA UGUUCACUC GCGAGCUGGAUGGACGAGG GUCGUACUACCGAGG CCCUGUGCCUGG AAUAAUA GCUGCGUACUGGGCAGCCG AAA UUCCAUGUC CGGCUGUCCAGUGCGCAGC CCAGGUACAGGG UU ACUU CACGAAGUCAAUAC | 188 |
| **2D-j-Tile63** | GGGCGUGCUCAGGC A UCGACUAAC GCUACGGUAAGAGCCUCGC AAA GUACAUUCC GCGAGGUUCUUAUCGUAGC A GCCUGAGUACGCUC CCAGCGUACGG A GUCAAGAUG CCUCGAGUACUAUCCGAGC AAA UCUUCAUCG GCUCGGGUAGUAUUCGAGG A CCGUAUGCUGG UU AU CACGAAGUCAAUAC | 190 |
| **2D-j-Tile72** | GCAGUGUGACUCGG A CACAGUAUC GGAGCAGAG GUAA CUCUGCUCC A CCGAGUCGCACUGC GCUGCGGAUCC A GGAAUGUAC GCGUCCAAU UUCG AUUGGACGC A GGAUCUGCAGC UU AA CACGAAGUCAAUAC | 134 |
| **2D-j-Tile73** | CCAGCCUUGUGAGC A GGUCUUCAA GCGGAUACUUAGGCAGCG AAA GUCACAUCA CGCUGCUUAAGUGUCCGC A GCUCACAGGGCUGG CCUGUUCCGCA A GAUACGAGA CUCGGUACAGUUUCGGGC AAA GUUAGUCGA GCCCGAGACUGUGCCGAG A UGCGGGACAGG UU AC CACGAAGUCAAUAC | 186 |
| **2D-j-Tile74** | GCACAUUUCCAGCC A CGUACAAUC GGCAAGCGUACGGCACGG AAA UGAGUUGAC CCGUGCUGUACGUUUGCC A GGCUGGAGAUGUGC GGUCUUAAGGC A CUUUGAACG CCCAGUACCUGGCAGGUG AAA GUUACGGUA CACCUGUCAGGUGCUGGG A GCCUUGAGACC UU UA CACGAAGUCAAUAC | 186 |
| **2D-j-Tile75** | CCUCGCGAGAGUCC A CAUCUAGAG GUCGGGUCUGAGGCGGUG AAA CUGAUACUC CACCGCUUCAGAUCCGAC A GGACUCUUGCGAGG CCAACGUACGG A UGAUAGCUG CGCCUGCGUGAGCCGGUG AAA GAUUGACUG CACCGGUUCACGUAGGCG A CCGUAUGUUGG UU UAAU CACGAAGUCAAUAC | 188 |
| **2D-j-Tile76** | GCCUGAGGUACGGC A CGUGUAAGA UCCGAGUUUGAGGCGUGG AAA UUGCAAGAG CCACGCUUCAAAUUCGGA A GCCGUACUUCAGGC GCAUGUUAGGG A UUGGACUUG CCUGCGGAUGAGCCCUGG AAA CGUAACUAG CCAGGGUUCAUCUGCAGG A CCCUAGCAUGC UU UACU CACGAAGUCAAUAC | 188 |
| **2D-j-Tile77** | CCGAGCGCUCUGCC A CAGUUAGGA GCGAAUGUCGUGGCGGUG AAA GCGAUAGUA CACCGCUACGACGUUCGC A GGCAGAGUGCUCGG CCUCCGAGACG A UGAGAAUGC GGAGCGGGUGAUGUUCGG AAA UACAACUGG CCGAACGUCACCUGCUCC A CGUCUUGGAGG UU AUAU CACGAAGUCAAUAC | 188 |
| **2D-j-Tile78** | GCAUCAGCAGUCCC A CUGAGAAUC GCCAGGUACAUUGCGCUG AAA CUUUGUACC CAGCGCGAUGUAUCUGGC A GGGACUGUUGAUGC GCGAGUGGGAC A CUGUGAAAC GCUGCGGUCUCGUUCCGC AAA CAUUCUGAG GCGGAAUGAGACUGCAGC A GUCCCGCUCGC UU AAUU CACGAAGUCAAUAC | 188 |
| **2D-j-Tile79** | CCAAGGGUCAUAGC A CACUUAGUC GCGACGCUGGCUGAGGUG AAA CAAUGAGUC CACCUCGGCCAGUGUCGC A GCUAUGAUCCUUGG CCCUGGACGAC A UCAUCAAGC CCUGCUGUCUAUACUCGG AAA CAAGGUAUG CCGAGUGUAGACGGCAGG A GUCGUUCAGGG UU ACUU CACGAAGUCAAUAC | 188 |
| **2D-k-Tile81** | GGGCGAUGGUAGCC A GUAUCCAUG UCGCAGUAGCUAGGCGAGC AAA GGAUUCACA GCUCGCUUAGCUGCUGCGA A GGCUACCGUCGCUC CCAGGUCUGCU A UGAUGUGAC GCUAAGGUCACUGCCGGUG AAA GAUACUGUG CACCGGUAGUGAUCUUAGC A AGCAGGCCUGG UU AU CACGAAGUCAAUAC | 190 |

| Name (Note) | Sequence | length |
|---|---|---|
| **2D-k-Tile82** | GCAGCGUAUCGGGC A **CUAGAGAUC** GCAACGGGUUAGUGCAGUG AAA **GUUCAGGAA** CACUGCGCUAACUCGUUGC A GCCCGAUGCGCUGC GCUGGUCGAGC A **GUCAACUCA** GCUAUCUAGCUCGCCUCUG AAA **UUGAAGACC** CAGAGGUGAGCUGGAUAGC A GCUCGGCCAGC **UU AA CACGAAGUCAAUAC** | 190 |
| **2D-k-Tile83** | CCGAGGGUAGUGAG A **UAGCCAAUG** CCUUACUUAUUGGCCUCGG AAA **CAGUAGAGA** CCGAGGUCAAUAGGUAAGG A CUCACUAUCCUCGG CCAGCGCGUGC A **GAGUAUCAG** GCGUCAGUACCCGGUCCUG AAA **GAUUGUACG** CAGGACUGGGUAUUGACGC A GCACGUGCUGG **UU AC CACGAAGUCAAUAC** | 190 |
| **2D-k-Tile84** | GAGCAUUAUGGGCC A **GCAGUUCUA** CCGUACGUGGAAGCCUCGG AAA **UAGCUCUAG** CCGAGGUUUCCAUGUACGG A GGCCCAUGAUGCUC GCCCUGUCCGA A **CUCUUGCAA** GGACAUUGCGAGGCCGUGG AAA **CUCUAGAUG** CCACGGUCUCGCGAUGUCC A UCGGAUAGGGC **UU UA CACGAAGUCAAUAC** | 190 |
| **2D-k-Tile85** | CCUCCGUUGUAGGC A **UCACUCUUG** CGCCUGUUGCUUGGCGGUG AAA **GGUAUCGUA** CACCGCUAAGCAGCAGGCG A GCCUACAGCGGAGG CAUCCGGGAGU A **UACUAUCGC** AGGGUCUCGGUCUAUCGUG AAA **UCUUACACG** CACGAUGGACCGGGACCCU A ACUCCUGGAUG **UU UAAU CACGAAGUCAAUAC** | 192 |
| **2D-k-Tile90** | GCCUCGUGAUCUAC A **UCCUAAGUC** UGUAACUGC **GUAA** GCAGUUACA A GUAGAUCGCGAGGC GCUACUAGUAC A **UGUGAAUCC** GGGAGUCUA **UUCG** UAGACUCCC A GUACUGGUAGC **UU UACU CACGAAGUCAAUAC** | 136 |
| **2D-k-Tile91** | CCACAUGCAGAUCC A **GACCAAUCA** GGCUGUCAGUAGGCGCAG AAA **UACGGAAAC** CUGCGCUUACUGGCAGCC A GGAUCUGUAUGUGG GUACUGGUGCC A **UUCCUGAAC** GGCAGUCUUGUGCCUCGG AAA **CAUGGAUAC** CCGAGGUACAAGGCUGCU A GGCACUAGUAC **UU AUAU CACGAAGUCAAUAC** | 188 |
| **2D-k-Tile99** | CCAGGCUG **UUCG** CAGCCUGG CGCUGCAC A **GUUUCCGUA** GGGCUGUGAAUCGCAGCUG AAA **GACUUAGGA** CAGCUGUGAUUCGCAGCCC A GUGCAGCG **UU AAUU CACGAAGUCAAUAC** | 117 |
| **2D-k-Tile100** | GCUCGAGG **UUCG** CCUCGAGC GGCUCCGC A **GUCUGAAUC** UGCACGUGUAUCGUCGGUG AAA **UGAUUGGUC** CACCGAUGAUACGCGUGCA A GCGGAGCC **UU ACUU CACGAAGUCAAUAC** | 117 |
| **2D-k-Tile101** | CCAGCGUC **UUCG** GACGCUGG CCCUCGCC A **GUAUGACGA** GCCACGUGGAUAUCUCGUG AAA **UUGCCUAUC** CACGAGGUAUCCGCGUGGC A GGCGAGGG **UU UAUA CACGAAGUCAAUAC** | 117 |
| **2D-k-Tile102** | GCCAGCAC **UUCG** GUGCUGGC GCUCCGGGU A **CUUGUGAUG** CGGGAGUCAACGGCUAUCG AAA **GUAACUCUC** CGAUAGUCGUUGGCUCCCG A ACCCGGAGC **UU AUUA CACGAAGUCAAUAC** | 119 |
| **2D-l-Tile92** | GGGCAUGUAGAUUC A **GAUAGGCAA** GUCCGUAUAAGGAUAGUC AAA **GAUUCAGAC** GACUAUUCUUAUGCGGAC A GAAUCUAUAUGUCC GCAAGUGCAGC A **UCUCUACUG** UCGUCUGUGGCGAGGGUG AAA **GAUCUCUAG** CACCCUUGCCACGGACGA A GCUGCGCUUGC **UU AU CACGAAGUCAAUAC** | 186 |
| **2D-l-Tile93** | CCUCGAGCAGCGCC A **GAGAGUUAC** GGGUAUAAGGGUGAGCGG AAA **UCGUCAUAC** CCGCUCGCCCUUGUACCC A GGCGCUGUUCGAGG CCCUGUAAUCG A **CUAGAGCUA** CGCGUGCAGCAGUUCCGG AAA **CAUUGGCUA** CCGGAAUUGCUGUACGCG A CGAUUGCAGGG **UU AA CACGAAGUCAAUAC** | 186 |
| **2D-l-Tile94** | GCCUUAUUGUGCCC A **CUUUCAUGC** AGGGUGCCGCUGAAGUCG AAA **CAUCACAAG** CGACUUUAGCGGUACCCU A GGGCACAGUAAGGC GCUCCGUGCCG A **UACGAUACC** CUCGCUGACCUGUAGCUG AAA **UAGAACUGC** CAGCUAUAGGUCGGCGAG A CGGCAUGGAGC **UU AC CACGAAGUCAAUAC** | 186 |
| **2D-l-Tile95** | CAGAUCUCUGACGG A **CUGGAUAGA** CGCGUUGCACAUGCCUGG AAA **UCUGCUUAG** CCAGGCGUGUGCGACGCG A CCGUCAGGGAUCUG CCUCUGAGCGG A **CUUGCAUAG** CCUGGUGAAGCGCAGCGG AAA **CAAGAGUGA** CCGCUGUGCUUCGCCAGG A CCGCUUAGAGG **UU UA CACGAAGUCAAUAC** | 186 |
| **2D-l-Tile96** | GCAGGAGCUAAUCC A **CUCAAGUUG** CAGGGUUCUUGUCCAGUG AAA **GUGACAGAA** CACUGGGCAAGAGCCCUG A GGAUUAGUUCCUGC GCAGGUGUGGG A **UGUAGACAG** CCCUGGCACAGUCCUCGG AAA **UCUGCAUAC** CCGAGGGCUGUGUCAGGG A CCCACGCCUGC **UU UAAU CACGAAGUCAAUAC** | 188 |
| **2D-l-Tile97** | CCAGCGUUCGAGUC A **UCCUAGUAC** UGGCGGAACCGGGACGUG AAA **GUUAGCUAC** CACGUCUCGGUUUCGCCA A GACUCGAGCGCUGG GCUUAUGUACC A **GGAUCAGAA** GCUCCUGGUAUGCCGGUG AAA **UAGGCAUAG** CACCGGUAUACCGGGAGC A GGUACGUAAGC **UU UACU CACGAAGUCAAUAC** | 188 |

| Name (Note) | Sequence | length |
|---|---|---|
| **2D-l-Tile98** | GCCAUACUGUUACGC **AAAUAAA** CCUGCGAAUACGGCCUGG AAA **CUACUGGAA** CCAGGCUGUAUUUGCAGG GCGUAACGGUAUGGC GCUACGGUACC A **GAGUGAACA** GCCUCUGUCAUGGCGGUG AAA **GACGAACUA** CACCGCUAUGACGGAGGC A GGUACUGUAGC **UU AUAU CACGAAGUCAAUAC** | 186 |
| **2D-l-Tile103** | GCAGGCAG **UUCG** CUGCCUGC CCCUCCGG A **CUAAGCAGA** CCCGUCUACUCGGCAUCCC AAA **GCAUGAAAG** GGGAUGUCGAGUGGACGGG A CCGGAGGG **UU AAUU CACGAAGUCAAUAC** | 117 |
| **2D-l-Tile104** | GCCUCACC **UUCG** GGUGAGGC GCAGCGCC A **UUCUGUCAC** CCGGAGUGCGAUUAACGUC AAA **UCUAUCCAG** GACGUUGAUCGCGCUCCGG A GGCGCUGC **UU ACUU CACGAAGUCAAUAC** | 117 |
| **2D-l-Tile105** | CCAGGCUG **UUCG** CAGCCUGG UCUCCGGG A **GUAGCUAAC** ACGGACGGUUGAUUCCUCG AAA **CAACUUGAG** CGAGGAGUCAACUGUCCGU A CCCGGAGA **UU UAUA CACGAAGUCAAUAC** | 117 |
| **2D-l-Tile106** | GCAGGGUC **UUCG** GACCCUGC GCUCGCCC A **UUCCAGUAG** GGCGUCGGGAUAUCUCGUC AAA **GUACUAGGA** GACGAGGUAUCCUGACGCC A GGGCGAGC **UU AUUA CACGAAGUCAAUAC** | 117 |

#### 5) Plasmid for 2D assembly

**2D_14** (2 plasmids): 2D-a-Plasmid, 2D-z-Plasmid.

**2D_34** (4 plasmids): 2D-a-Plasmid, 2D-b-Plasmid, 2D-c-Plasmid, 2D-y-Plasmid.

**2D_62** (7 plasmids): 2D-a-Plasmid, 2D-b-Plasmid, 2D-c-Plasmid, 2D-d-Plasmid, 2D-e-Plasmid, 2D-f-Plasmid, 2D-x-Plasmid.

**2D_107** (12 plasmids): 2D-a-Plasmid, 2D-b-Plasmid, 2D-c-Plasmid, 2D-d-Plasmid, 2D-e-Plasmid, 2D-f-Plasmid, 2D-g-Plasmid, 2D-h-Plasmid, 2D-i-Plasmid, 2D-j-Plasmid, 2D-k-Plasmid, 2D-l-Plasmid.

| Name (Note) | Sequence | length |
|---|---|---|
| **2D-a-Plasmid** Template for 2D-a tiles | **GTTCTAATACGACTCACTATA** GGGCTATGACACGC A **CTTCAAGAC** AGCGGTATG **GTAA** CATACCGCT A GCGTGTCGTAGCTC GCTCGTACCGG A **CTGCTAACC** TCCGCTGCT **TTCG** AGCAGCGGA A CCGGTGCGAGC **TT AT CACGAAGTCAATAC** CCTGAGGGCTAGGC A **GCTTCAATC** GGGTCGCTCGCTTCACAG AAA **TAGAGCAAC** CTGTGAGGCGAGTGACCC A GCCTAGCTCTCAGG GCTCGTGGACC A **CTACTCGTA** CACGGCAGGGACCTCGCCTACATCTAGTCGTTGACCTTTACTGGTG AAA **GGTTAGCAG** CACCAGTGAAGGTCAGCGACTAGGTGTAGGTGAGGTCCTTGCCGTG A GGTCCGCGAGC **TT AA CACGAAGTCAATAC** CCTACAGTGGACCC A **TAGTTGAGC** GGTCCCGTCTGCGGACGTG AAA **GCTTGTTAC** CACGTCTGCAGATGGGACC A GGGTCCATTGTAGG GCTGGTGAGCC A **GTTGCTCTA** CCGCTTGCACCGTGAGGTG AAA **GTCTTGAAG** CACCTCGCGGTGTAAGCGG A GGCTCGCCAGC **TT AC CACGAAGTCAATAC** GCAGTGTCAGCTGC A **CCTTGAGAA** CAGAAGTACGCAGGAGTGG AAA **TTCATAGCG** CCACTCTTGCGTGCTTCTG A GCAGCTGGCACTGC GCTCCTGGAGC A **GATACCTTC** GCGAGATGGCTCTATACTG AAA **GATTGAAGC** CAGTATGGAGCCGTCTCGC A GCTCCGGGAGC **TT TA CACGAAGTCAATAC** GCCTACTTCTGCGC A **GACAGTTGA** CCTAAGTCA **GTAA** TGACTTAGG A GCGCAGAGGTAGGC GCTCCTGCAGG A **GTAACAAGC** CCAGCCGGT **TTCG** ACCGGCTGG A CCTGCGGGAGC **TT TAAT CACGAAGTCAATAC** CCATGTGCACCTGG A **CTTTGCTCA** CGGCCTAGGTAGGACCGG AAA **CAATCGACA** CCGGTCCTACCTAGGCCG A CCAGGTGTACATGG GCACTTATCGG A **CGCTATGAA** CCGCCTGTCACTATCGTG AAA **GCTCAACTA** CACGATAGTGACAGGCGG A CCGATGAGTGC **TT TACT CACGAAGTCAATAC** GCTACCTAGAGTGC A **TGTATCCTC** GCCAGGGCGTCTGCGGTG AAA **GAATGGCTA** CACCGCAGACGCCTTGGC A GCACTCTGGGTAGC GCATGTGTCGC A **TCACTACAG** CCCGAGGGCTCTGAGTGG AAA **TTCTCAAGG** CCACTCAGAGCCCTCGGG A GCGACGCATGC **TT ATAT CACGAAGTCAATAC** GCCTCGTGTTACGG A **CTAACTAGC** CCGTCTTGTCTGGGTCGTG AAA **GAAGTATGC** CACGACTCAGACGAGACGG A CCGTAACGCGAGGC GCTGTTGCAGC A **TAGCCATTC** GAGTGATTCGAATGGTCGG AAA **TGAGCAAAG** CCGACCGTTCGAGTCACTC A GCTGCGACAGC **TT AATT CACGAAGTCAATAC** **GTCCAACC TT CATGCTTACGACG** | 1504 |

| Name (Note) | Sequence | length |
|---|---|---|
| **2D-z-Plasmid**<br>Template for 2D-z tiles | **GTTCTAATACGACTCACTATA** GGGCCTGGGCGACT A **GTCTCATGA** ACCGGTACCGCTTGCGTG<br>AAA **GAAGGTATC** CACGCAGGCGGTGCCGGT A AGTCGCCTAGGCTC GCTACCTGCAGG<br>**AATAATA** CGTCGTCGGTTCTGAGATCGAGTGATGTTATCTACATCGGTGACGG AAA **TACGAGTAG**<br>CCGTCACTGATGTAGGTAACATCGCTCGATTTCAGAACTGACGACG CCTGCGGGTAGC **TT AT**<br>**CACGAAGTCAATAC** CCTAGTGGGCCTAGC **AAATAAA** GCTACGGGACACTGCTCGG AAA<br>**CTGTAGTGA** CCGAGCGGTGTCTCGTAGC GCTAGGCTCACTAGG GCCTCTGCGACC **AATAATA**<br>GCCAGTTCACGAGCTGCGG AAA **TCATGAGAC** CCGCAGTTCGTGGACTGGC GGTCGTAGAGGC<br>**TT AA CACGAAGTCAATAC** CCAGCTTCGAGGGC A **TCGATAGAG** GCCGACACA **GTAA**<br>TGTGTCGGC A GCCCTCGGAGCTGG GCCTGGTCGGC A **TGTCGATTG**<br>CCAGGTGCAGCAGCCGGTG AAA **TCAACTGTC** CACCGGTTGCTGTACCTGG A GCCGATCAGGC<br>**TT AC CACGAAGTCAATAC** CCAGTAGTGCCACGG **AAATAAA** GCAGAATCTTGGGCCTCGC AAA<br>**GCAATCATG** GCGAGGTCCAAGGTTCTGC CCGTGGCGCTACTGG GCGTTATGTAGG **AATAATA**<br>GCAGCGTACATGGTTAGCG AAA **GAGGATACA** CGCTAATCATGTGCGCTGC CCTACGTAACGC<br>**TT TA CACGAAGTCAATAC** CCTCGAGC **TTCG** GCTCGAGG GCTCCGGC A **GCATACTTC**<br>ACCGAGGCAGAGTACCGG AAA **CTCTATCGA** CCGGTACTCTGCCTCGGT A GCCGGAGC **TT**<br>**TAAT CACGAAGTCAATAC** GCAGGGTC **TTCG** GACCCTGC GCCTCCGG A **CATGATTGC**<br>CCGTAGGGTCCTGAGGGC AAA **GCTAGTTAG** GCCCTCAGGACCCTACGG A CCGGAGGC **TT**<br>**TACT CACGAAGTCAATAC GTCCAACC TT CATGCTTACGACG** | 1048 |
| **2D-b-Plasmid**<br>Template for 2D-b tiles | **GTTCTAATACGACTCACTATA** GGGCGAGTCTGACT A **GTCTCATGA** ACAGCTAGCCTGGAGGTG<br>AAA **GAAGGTATC** CACCTCTAGGCTGGCTGT A AGTCAGATTCGCTC GCCTCTGAGCC A<br>**TTGCATTCG** GGCTCCCGTAGACCTGTCCACTGTGAGCATGACAGCCATCTACCGG AAA<br>**TACGAGTAG** CCGGTAGGTGGCTGTTATGCTCATAGTGGATAGGTCTATGGGAGCC A<br>GGCTCGGAGGC **TT AT CACGAAGTCAATAC** CCACTCTTACATCC A **TCAGGTTAC**<br>GCAGCGCAGCTTGGGCTG AAA **GTGTACCTA** CAGCCCGAGCTGTGCTGC A GGATGTAGGAGTGG<br>GCTCCGGACGG A **TTCGACATG**<br>CCTGCGGTACGTGTCTGTCGTGGTACATACTTCAAGGTGTCACCGG AAA **CGAATGCAA**<br>CCGGTGATACCTTGAGGTATGTATCACGACGGACACGTGCCGCAGG A CCGTCTGGAGC **TT AA**<br>**CACGAAGTCAATAC** CCATCGTGAGTAGC A **GCAATCTTC** GTCACGTCTCCGTGACCGG AAA<br>**CTGTAGTGA** CCGGTCGCGGAGGCGTGAC A GCTACTCGCGATGG GCCTCGCGTCC A<br>**TAGGTACAC** GGAGCATCGGAGTCCAGTG AAA **TCATGAGAC** CACTGGGCTCCGGTGCTCC A<br>GGACGTGAGGC **TT AC CACGAAGTCAATAC** GCCAAGTGGACTGC A **GACTACCTA**<br>CCGAGCGGTACTGGCGGTG AAA **GCTGATTGA** CACCGCTAGTACTGCTCGG A<br>GCAGTCCGCTTGGC GCTCCGCAGCC A **GAGAACATG** GCATTATGCTTGTCCTGTG AAA<br>**GTAACCTGA** CACAGGGCAAGCGTAATGC A GGCTGTGGAGC **TT TA CACGAAGTCAATAC**<br>CCATCCTACTTGGC A **TGCAGATTC** GACCCTATCACTCCGGTG AAA **GGTTACATG**<br>CACCGGAGTGATAGGGTC A GCCAAGTGGGATGG GCATATGGTCC A **TCAATCAGC**<br>ACCGAGGTGAAGGTCGTG AAA **GAAGATTGC** CACGACCTTCACCTCGGT A GGACCGTATGC **TT**<br>**TAAT CACGAAGTCAATAC** GGTCCAGGCTCGGG A **GTGCATTGA** TCGACTACGCCTGCCTGG<br>AAA **TACATGTGG** CCAGGCAGGCGTAGTCGA A CCCGAGCTTGGACC GCTACTAGTGG A<br>**CTTAGCTTC** AGGTCTCAAACTCAGGTG AAA **TAGGTAGTC** CACCTGAGTTTGAGACCT A<br>CCACTGGTAGC **TT TACT CACGAAGTCAATAC** CCTATGGAGCAAGC A **CATACATGG**<br>CCGATTGAGGCGGTCTCGC AAA **GCAATCATG** GCGAGATCGCCTTAATCGG A<br>GCTTGCTTCATAGG GCTCCGCTGAG A **CATGTAACC** GCGTGGGCCTAGGGCTCTG AAA<br>**GAGGATACA** CAGAGCTCTAGGTCCACGC A CTCAGTGGAGC **TT ATAT CACGAAGTCAATAC**<br>GCAGTCGAGAGCCC A **CTTCTGACA** GGCCAAGGCGTGGGCTCGG AAA **TACAGTCAC**<br>CCGAGCTCACGCTTTGGCC A GGGCTCTTGACTGC GCTGGTACCGG A **CCACATGTA**<br>CGTACGGAATACTGGTCTG AAA **GAATCTGCA** CAGACCGGTATTTCGTACG A CCGGTGCCAGC<br>**TT AATT CACGAAGTCAATAC GTCCAACC TT CATGCTTACGACG** | 1668 |

| Name (Note) | Sequence | length |
|---|---|---|
| **2D-c-Plasmid**<br>Template for 2D-c tiles | **GTTCTAATACGACTCACTATA** GGGCAATGAGTACC A **TCGATAGAG** GCAGCCGGATCCGGACGGC<br>AAA **CATTCGTAC** GCCGTCTGGATCTGGCTGC A GGTACTCGTTGTCC GCTAGTCATGC A<br>**TGTCGATTG** CCAATGTCAATTGCCGGTG AAA **TCAACTGTC** CACCGGTAATTGGCATTGG A<br>GCATGGCTAGC **TT AT CACGAAGTCAATAC** GCCTAATGTCCTGG A **GGCTATGTA**<br>CCGCTTCGG **GTAA** CCGAAGCGG A CCAGGACGTTAGGC GCAGCTGCTCC A **GTACGAATG**<br>GAAGCTGCG **TTCG** CGCAGCTTC A GGAGCGGCTGC **TT AA CACGAAGTCAATAC**<br>CAGATGTCGCTAGG A **TAGCAGTAG** TCCGAGCCGATTGGGACG AAA **GAGTCATGA**<br>CGTCCCAGTCGGCTCGGA A CCTAGCGGCATCTG GCGACTACGGC A **GCATACTTC**<br>ACCGCTCCACTTTACGTG AAA **CTCTATCGA** CACGTAAGGTGGAGCGGT A GCCGTGGTCGC **TT**<br>**AC CACGAAGTCAATAC** GGTGTATTCCAAGC A **TTCCGAAAG** CGCGAGTCATCTGGAGCG AAA<br>**GAAGTCTCA** CGCTCCAGATGACTCGCG A GCTTGGAGTACACC GCGACGCCTGG A<br>**CATGATTGC** CTCACTCGGGAGCGTCCG AAA **GCTAGTTAG** CGGACGCTCCCGAGTGAG A<br>CCAGGTGTCGC **TT TA CACGAAGTCAATAC** CCATCAGGTCGAGG A **TAGGACTAG**<br>ACGTTTCTATCTGCGTGG AAA **CTCAATGGA** CCACGCAGATAGAGACGT A CCTCGACTTGATGG<br>GCCTATCGACC A **GTGACTGTA** GCCGAGCATGAGGCCTGG AAA **CCATGTATG**<br>CCAGGCCTCATGCTCGGT A GGTCGGTAGGC **TT TAAT CACGAAGTCAATAC**<br>GCATGAGAGTCGGC A **GGATCTTGA** CCGTAGGGTCTTGGTCGG AAA **CAGTCTTAG**<br>CCGACCAGGACCCTACGG A GCCGACTTTCATGC GCTTGGATCGC A **GATGACTTG**<br>ACCGAGCATGCTACGTGG AAA **TGTCAGAAG** CCACGTAGCATGCTCGGT A GCGATTCAAGC **TT**<br>**TACT CACGAAGTCAATAC** GCCTGCGGACTGGC A **CACTGTTCA** CCCGACGCTCTCGGACGTG<br>AAA **CATCTATGC** CACGTCTGAGAGTGTCGGG A GCCAGTCTGCAGGC GCTCGGCCAGG A<br>**TGAGACTTC** GCCTAAGCGTGCTGGTCGG AAA **CTACTGCTA** CCGACCGGCACGTTTAGGC A<br>CCTGGTCGAGC **TT ATAT CACGAAGTCAATAC** CAGCTCTCGATCGG A **TACTTAGCG**<br>CGATACGGTCGGTGACGCG AAA **CAATGTCTC** CGCGTCGCCGACTGTATCG A<br>CCGATCGGGAGCTG GCGGAGCGTAG A **TCCATTGAG** ACGCCTTGTAATGCCGAGC AAA<br>**CTTTCGGAA** GCTCGGTATTACGAGGCGT A CTACGTTCCGC **TT AATT CACGAAGTCAATAC**<br>GCCTAGTACGTGGC A **TAGTACAGG** GCCATCTAGCCAGGGCGTG AAA **GATCATGTG**<br>CACGCCTTGGCTGGATGGC A GCCACGTGCTAGGC GCTCGGCACGG A **CTAAGACTG**<br>GCTCCGGTCTTAGCCGGTG AAA **CTAGTCCTA** CACCGGTTAAGATCGGAGC A CCGTGTCGAGC<br>**TT ACTT CACGAAGTCAATAC** GTCCAACC TT CATGCTTACGACG | 1692 |
| **2D-y-Plasmid**<br>Template for 2D-y tiles | **GTTCTAATACGACTCACTATA** GGGCCATGACCTGC A **CTTGGATCA** CCCGTGCGTCGGGACGGC<br>AAA **CATGTTCTC** GCCGTCTCGACGTACGGG A GCAGGTCGTGGCTC CCAGTCGCGGTG<br>**AATAATA** CCCAGTATGTTTCTGGACGATCTCGAATGCTGTACATATGCACCGG AAA **CATGTCGAA**<br>CCGGTGCGTATGTACGGCATTCGGGATCGTTCAGAAACGTACTGGG CACCGTGACTGG **TT AT**<br>**CACGAAGTCAATAC** CCTTAGGGCACGCTG **AAATAAA** GCCCTTGCCGGATCCGGTG AAA<br>**GAAGCTAAG** CACCGGGTCCGGTAAGGGC CAGCGTGTCCTAAGG GCGTATTTCAGG **AATAATA**<br>GCTAGATCAAGCGGTCGTG AAA **TGATCCAAG** CACGACTGCTTGGTCTAGC CCTGAGATACGC<br>**TT AA CACGAAGTCAATAC** CCGACGGTCAAGTCC **AAATAAA** GGTGGATCGCCTGGCGTCG AAA<br>**CAAGTCATC** CGACGCTAGGCGGTCCACC GGACTTGGCCGTCGG CCTTGCTGTACG **AATAATA**<br>GTCCGTTTATGCGTTCCGG AAA **TCAATGCAC** CCGGAATGCATAGACGGAC CGTACGGCAAGG<br>**TT AC CACGAAGTCAATAC** CCAGGCGGACAGGG A **CACTTCAGA** GCCGATACG **GTAA**<br>CGTATCGGC A CCCTGTCTGCCTGG GCTACTGAGGC A **TCATGACTC**<br>GCCAGTTTGAAGTGCAGTG AAA **TACATAGCC** CACTGCGCTTCAGACTGGC A GCCTCGGTAGC<br>**TT TA CACGAAGTCAATAC** CCAGCATGGATTGGC **AAATAAA** GCTCAAGTTCACGGACACG AAA<br>**TAGGTCAAG** CGTGTCTGTGAATTTGAGC GCCAATCTATGCTGG CCTTGGGCACCG **AATAATA**<br>GGATTCTCAAGGTCATCTG AAA **TCAAGATCC** CAGATGGCCTTGGGAATCC CGGTGTCCAAGG<br>**TT TAAT CACGAAGTCAATAC** CCTCGGTC **TTCG** GACCGAGG GCTCCCGG A **GCATAGATG**<br>ACCGTTCGTCAGCCGGTG AAA **TCTGAAGTG** CACCGGCTGACGAGCGGT A CCGGGAGC **TT**<br>**TACT CACGAAGTCAATAC** GCCTCACC **TTCG** GGTGAGGC CCTCGGCC A **GAGACATTG**<br>CCACCTCGTGAGATGCGG AAA **TGAACAGTG** CCGCATCTCACGAGGTGG A GGCCGAGG **TT**<br>**ATAT CACGAAGTCAATAC** CCGATCGC **TTCG** GCGATCGG GCACCGGG A **CACATGATC**<br>TCCACTGGATGGTCGGTG AAA **CGCTAAGTA** CACCGACTATCCGGTGGA A CCCGGTGC **TT**<br>**AATT CACGAAGTCAATAC** GCCAGGTC **TTCG** GACCTGGC GCACCGCG A **CTTGACCTA**<br>CCCGAGGTAGGGTCGCGG AAA **CCTGTACTA** CCGCGACTCTACCTCGGG A CGCGGTGC **TT**<br>**ACTT CACGAAGTCAATAC** GTCCAACC TT CATGCTTACGACG | 1466 |

| Name (Note) | Sequence | length |
|---|---|---|
| **2D-d-Plasmid** Template for 2D-d tiles | GTTCTAATACGACTCACTATA GGGCTCTGCACTGC A CTTGGATCA CTGCCTACGACGGACCGC AAA CATGTTCTC GCGGTCTGTCGTGGGCAG A GCAGTGCGGAGTCC GCTCCTGAGCC A GTATCTCCA TCGCGGATGTTGCCGGTCACAGTCTAATACTTCAGATTTGCGCTGG AAA CATGTCGAA CCAGCGCGAATCTGAGGTATTAGGCTGTGATCGGCAACGTCCGCGA A GGCTCGGGAGC TT AT CACGAAGTCAATAC CCGTCCTGACTCGC A CGATCTTAC CCGTCGGCTGGTCTGCTG AAA GTAAGGACA CAGCAGGCCAGCTGACGG A GCGAGTCGGGACGG CCTACGGCTGG A CAGAACTCA CCCTGGCGAGCATCGTCTCGTATGACGATTGACCAGGTTATGCTGG AAA TGGAGATAC CCAGCATGACCTGGTTAATCGTCGTACGAGGCGATGCTTGCCAGGG A CCAGCTGTAGG TT AA CACGAAGTCAATAC CCGAGCGATCGGGC A CCTTGATTC GCGTTGGATGTAGGCGGTG AAA GAAGCTAAG CACCGCTTACATTCAACGC A GCCCGATTGCTCGG CCAGCTGTAGG A TGTCCTTAC CGCGAGTTAACGGGACCGG AAA TGATCCAAG CCGGTCTCGTTAGCTCGCG A CCTACGGCTGG TT AC CACGAAGTCAATAC GCCTAGTAGACAGG A GTTGCTAGA GCCTACTGAGATGGCGGTG AAA GAAGACGTA CACCGCTATCTCGGTAGGC A CCTGTCTGCTAGGC GCTCCGGGAGC A GGTACTACA GCGTCTGGATTTGCCCGTG AAA GTAAGATCG CACGGGTAAATCTAGACGC A GCTCCTGGAGC TT TA CACGAAGTCAATAC CCATGTTTCAGGCC A CATTGCATC TCGGTTGAAACTGCGGTG AAA TCAGTACTC CACCGCAGTTTCAGCCGA A GGCCTGAGACATGG CCTTGTCCAGC A TACGTCTTC GGGTCTCCACTTAGAGTG AAA GAATCAAGG CACTCTAGGTGGAGACCC A GCTGGGCAAGG TT TAAT CACGAAGTCAATAC GAGCTATATCTGCC A TTCCACTTG GCTCGGCCATAGGTCCGG AAA CTAACTCTG CCGGACCTATGGCTGAGC A GGCAGATGTAGCTC CCATGGTAGGG A TAGTAACGC CTGTCTGGACTTAGACAG AAA TCTAGCAAC CTGTCTAGGTCCAGACAG A CCCTATCATGG TT TACT CACGAAGTCAATAC CCGTCGTGGATGGC A CCTATGTTC GCGAGCGGCTCGGGCAGCG AAA CAAGTCATC CGCTGCTCGAGCTGCTCGC A GCCATCCGCGACGG CCAGGTGAGGC A GAGTACTGA GCTCCATGGTCGGCACCGG AAA TCAATGCAC CCGGTGTCGACCGTGGAGC A GCCTCGCCTGG TT ATAT CACGAAGTCAATAC GCCTGTGAGGTAGG A TCATGCTAG CCCTAGGTAGGCTGACGTG AAA GGCAATGAA CACGTCGGCCTATCTAGGG A CCTACCTTACAGGC GCTCCGACGGG A CAGAGTTAG GCTCATGCTGCCGGAGGTG AAA GATGCAATG CACCTCTGGCAGTATGAGC A CCCGTTGGAGC TT AATT CACGAAGTCAATAC GTCCAACC TT CATGCTTACGACG | 1668 |
| **2D-e-Plasmid** Template for 2D-e tiles | GTTCTAATACGACTCACTATA GGGCACGCGACCGC A GCTACTATC GTCCGGCTATAGCGAGCG AAA TCGAATCTC CGCTCGCTATAGCTGGAC A GCGGTCGTGTGCTC CCGTGGGACCC A TTCATTGCC GACCAGCTCACTCCGGTG AAA GAACATAGG CACCGGAGTGAGCTGGTC A GGGTCTCACGG TT AT CACGAAGTCAATAC GCAAGCGGACTAGC A TGAATGTCC GTGCAGAAGCCTGCGGTG AAA GATAGATGC CACCGCAGGCTTCTGCAC A GCTAGTCTGCTTGC GCGTCTGTTGC A TGTTCCTAG TCACAGGGTACTCTGCGG AAA CTAGCATGA CCGCAGAGTACCCTGTGA A GCAACGGACGC TT AA CACGAAGTCAATAC CCAGGGTCTAAAGC A TGGAACAAG GCCTGTGCAACTGCGCGTG AAA TAGGTCAAG CACGCGTAGTTGTACAGGC A GCTTTAGGCCCTGG CCAGGTCCAGC A GAGATTCGA GGACACGGGTGCTGGAGTG AAA TCAAGATCC CACTCCGGCACCTGTGTCC A GCTGGGCCTGG TT AC CACGAAGTCAATAC GCAGTTGAGACCCG A TGAGCATTC CCAGCGGTAAAGGCCTCGG AAA CAGATGTAG CCGAGGTCTTTATCGCTGG A CGGGTCTTAACTGC GCTCCGGAGCC A GCATCTATC GGCGTCTATCCTTGCGGTG AAA GATAGTAGC CACCGCGAGGATGGACGCC A GGCTCTGGAGC TT TA CACGAAGTCAATAC CCAGGCGATTCCGG A TCAGTGAAG CCGGTTATGACTGCGGTG AAA GATGTCTGA CACCGCAGTCATAGCCGG A CCGGAATTGCCTGG CCTGGGTCTCG A CACATGATC TCAGCTCGAAAGCTCCGG AAA CGCTAAGTA CCGGAGCTTTCGAGCTGA A CGAGATCCAGG TT TAAT CACGAAGTCAATAC GCCAGCGACAGGCC A GGATACCAA GCCGAGGCACTTGGTCGG AAA TCCTTGTAG CCGACCAGGTGCCTCGGC A GGCCTGTTGCTGGC GCAGAGACGCG A CTTGACCTA CAGCCTGAAATTCGGCTG AAA CCTGTACTA CAGCCGAGTTTCAGGCTG A CGCGTTTCTGC TT TACT CACGAAGTCAATAC CCAGCATATTGAGG A TAGACATGC GGAGCTCCAGGGACCCGC AAA CTTTCGTCA GCGGGTCTCTGGAGCTCC A CCTCAATGTGCTGG CCCTGTACAGC A CTACATCTG CCGCAGGATCTTGGACGC AAA CTTGTTCCA GCGTCCAGGATCCTGCGG A GCTGTGCAGGG TT ATAT CACGAAGTCAATAC GGGAACTTGGGCTC A GGTAAGTAC GTCAAGCGAGCTGGAGGC AAA GTAAGCGTA GCCTCCAGCTCGCTTGAC A GAGCCCAGGTTCCC GCTTGTCCTGC A CTACGATCA CCGGCTCCTACTTGCCCG AAA GAATGCTCA CGGGCAAGTAGGAGCCGG A GCAGGGCAAGC TT AATT CACGAAGTCAATAC GTCCAACC TT CATGCTTACGACG | 1548 |

| Name (Note) | Sequence | length |
|---|---|---|
| **2D-f-Plasmid**<br>Template for 2D-f tiles | GTTCTAATACGACTCACTATA GGGCATGACCGGTC A CACTTCAGA GGCCCTGTATGGTGCTGTG<br>AAA GTTGATGTC CACAGCGCCATATAGGGCC A GACCGGTTATGCTC CCGGTTCAGGC A<br>TCATGACTC GGTCTATCGCGTGCCCGTG AAA TACATAGCC CACGGGTACGCGGTAGACC A<br>GCCTGGACCGG TT AT CACGAAGTCAATAC GCAGCTGCCAAGCC A CGATGAAGA<br>CCGTGAGTA GTAA TACTCACGG A GGCTTGGTAGCTGC GCTCCGTGCCC A GACATCAAC<br>GGCCAAAGG TTCG CCTTTGGCC A GGGCATGGAGC TT AA CACGAAGTCAATAC<br>CCATCTTGCCGTGC A GTACCTTAG CCCGAGGATCCTGGACGG AAA CATCTTGAC<br>CCGTCCAGGATCCTCGGG A GCACGGCGAGATGG CCTCAGATCCG A GCATAGATG<br>ACCCAGGCCTCTGACCGG AAA TCTGAAGTG CCGGTCAGAGGCCTGGGT A CGGATTTGAGG TT<br>AC CACGAAGTCAATAC GCAGTGTGCAGCCC A CGATCTGAA GCGTCTTTAGAGGTCATC AAA<br>TAGATTCCG GATGACCTCTAAAGACGC A GGGCTGCGCACTGC GCACCTTGGCC A<br>GAGACATTG CACGCTGTTCGGCCACGG AAA TGAACAGTG CCGTGGCTGAACAGCGTG A<br>GGCCAGGGTGC TT TA CACGAAGTCAATAC GCCTACGCCTCACC A TACCGTAAC<br>CGAGCATCAGTGGGCTCCG AAA TCTCGTATC CGGAGCTCACTGGTGCTCG A<br>GGTGAGGTGTAGGC GCTGGTCGAGG A CGGAATCTA GGCGACGCGATCGCCTCTG AAA<br>CTAAGGTAC CAGAGGTGATCGTGTCGCC A CCTCGGCCAGC TT TAAT CACGAAGTCAATAC<br>CCGTCCTTGCGTGC A CAGTCAATC AGGCCATAGTTCGTATCGC AAA CGTTCAAAG<br>GCGATATGAACTGTGGCCT A GCACGCAGGGACGG CCTCCGGGAGC A TCAGACATC<br>GCTGTCGGAGGCGGAGTGG AAA TTCAGATCG CCACTCTGCCTCTGACAGC A GCTCCTGGAGG<br>TT TACT CACGAAGTCAATAC GCACCGTCTGGGCC A CTAGTTACG<br>CCGAGTGCAGTCGGGACGG AAA CAGCTATCA CCGTCCTGACTGTACTCGG A<br>GGCCCAGGCGGTGC GCTGTTGCGAC A CTACAAGGA CCGAGTGGCAATGCCCTGG AAA<br>CTTCACTGA CCAGGGTATTGCTACTCGG A GTCGCGACAGC TT ATAT CACGAAGTCAATAC<br>CCAAGTGCTCCGGC A CCAGTTGTA CGCGGAGTACAGTGAGCAG AAA CAAGTCCAA<br>CTGCTCGCTGTATTCCGCG A GCCGGAGTACTTGG CCCTCGCCTGC A TGACGAAAG<br>CCGTCCGACTCGGCTCGTG AAA TTGGTATCC CACGAGTCGAGTTGGACGG A GCAGGTGAGGG<br>TT AATT CACGAAGTCAATAC GCCACTGCGGCAGG A CTCAGAATG<br>CCGCAAGCGTCCGGACCGC AAA GCATTCTCA GCGGTCTGGACGTTTGCGG A<br>CCTGCCGTAGTGGC GCAGGGTCCGG A TACGCTTAC CCTTGCTTGCTATCCGGTG AAA<br>GCATGTCTA CACCGGGTAGCAGGCAAGG A CCGGATCCTGC TT ACTT CACGAAGTCAATAC<br>GTCCAACC TT CATGCTTACGACG | 1700 |
| **2D-x-Plasmid**<br>Template for 2D-x tiles | GTTCTAATACGACTCACTATA GGGCTTGAGGTACC A GATCACTTC GCAGCTGGTGCTCGTCTG<br>AAA TGTAGTACC CAGACGGGCACCGGCTGC A GGTACCTTAAGCTC GCCTCGTGCAGC<br>AATAATA GCCAGGATGTGAACGGCACTGCGGAGATTATCTGTGCATATCACGC AAA TGAGTTCTG<br>GCGTGATGTGCACAGGTAATCTCTGCAGTGTCGTTCACGTCCTGGC GCTGCGCGAGGC TT AT<br>CACGAAGTCAATAC CCAGCTTTATCATGC AAATAAA GACGTGTAGCTGGGCTGTG AAA<br>GCGTTACTA CACAGCTCAGCTGCACGTC GCATGATGAAGCTGG CCCTCGTTGCGG AATAATA<br>GTGACTTCGTAGGCGGACG AAA GAAGTGATC CGTCCGTCTACGGAGTCAC CCGCAGCGAGGG<br>TT AA CACGAAGTCAATAC CCAGCATTTAGGTCC AAATAAA GCTGCTGTCGACTGACCCG AAA<br>CTAGGAACA CGGGTCGGTCGATAGCAGC GGACCTAGATGCTGG CCTAGTTCGCTC AATAATA<br>GGGTCCGAGGCTTCGACGG AAA CAAGTGGAA CCGTCGGAGCCTTGGACCC GAGCGGACTAGG<br>TT AC CACGAAGTCAATAC CCTCCAGGCAGTCGG AAATAAA CGAGCCTATATCTGAGGTG AAA<br>TGATCGTAG CACCTCGGATATGGGCTCG CCGACTGTCTGGAGG CCGTTCTTAGCC AAATAAA<br>GGTCGTTAACCGTACAGGC AAA GGACATTCA GCCTGTGCGGTTGACGACC GGCTAGGAACGG<br>TT TA CACGAAGTCAATAC CACCCTGCGTGAGG A TCGACTAAC GCCACTACT GTAA<br>AGTAGTGGC A CCTCACGTAGGGTG CCCGAGTCGCC A GTCAAGATG<br>CCGCTGGCTCCGTGAGAGC AAA TCTTCATCG GCTCTCGCGGAGTCAGCGG A GGCGATTCGGG<br>TT TAAT CACGAAGTCAATAC CCTGGACTTCAATCG AAATAAA GCTTGATCTACATGCGGAC<br>AAA GTTTCACAG GTCCGCGTGTAGGTCAAGC CGATTGAGGTCCAGG CCTGCTGGCAGC<br>AATAATA GGATCGGGTTCATATCCTG AAA GTACTTACC CAGGATGTGAACTCGATCC<br>GCTGCTAGCAGG TT TACT CACGAAGTCAATAC CCTCGCGA TTCG TCGCGAGG CCCTGGCC<br>A GATACGAGA CCGCAGGTAGCTCCAGGC AAA GTTAGTCGA GCCTGGAGCTACCTGCGG A<br>GGCCAGGG TT ATAT CACGAAGTCAATAC GCTCGGTC TTCG GACCGAGC GCTCCGGG A<br>CTTTGAACG CCCTAGGGCACTACCCTG AAA GTTACGGTA CAGGGTAGTGCCCTAGGG A<br>CCCGGAGC TT AATT CACGAAGTCAATAC CCAGGAGC TTCG GCTCCTGG CCTCCGGG A<br>TGATAGCTG CCGCAGGGCGAGAGCGTG AAA GATTGACTG CACGCTCTCGCCCTGCGG A<br>CCCGGAGG TT ACTT CACGAAGTCAATAC GCCAGCAC TTCG GTGCTGGC GCTCCCGC A<br>TTGGACTTG CCCGAGGTACAGTCTCGG AAA CGTAACTAG CCGAGACTGTACCTCGGG A<br>GCGGGAGC TT TATA CACGAAGTCAATAC CCAGGGTC TTCG GACCCTGG CCTCGCCC A<br>TGAGAATGC GGTGAGCCTACTCAGGTG AAA TACAACTGG CACCTGAGTAGGCTCACC A<br>GGGCGAGG TT ATTA CACGAAGTCAATAC GCAGGAGC TTCG GCTCCTGC GCCTCCGC A<br>CTGTGAAAC GCTCCTGCCGCTCTGGGC AAA CATTCTGAG GCCCAGAGCGGCAGGAGC A<br>GCGGAGGC TT CTTA CACGAAGTCAATAC GTCCAACC TT CATGCTTACGACG | 1884 |

| Name (Note) | Sequence | length |
|---|---|---|
| **2D-g-Plasmid**<br>Template for 2D-g tiles | GTTCTAATACGACTCACTATA GGGCATGATGCTCC A GATCACTTC GCACCTGTCTCTATCCTG AAA TGTAGTACC CAGGATGGAGACGGGTGC A GGAGCATTATGCCC GACCGTACGGG A CCGATTCTA TCCAGGGTTCGGCTGGTAGATGGACTGAGTTGCTTTGATCGAGGGC AAA TGAGTTCTG GCCCTCGGTCAAAGCGACTCAGTTCATCTATCAGCCGAGCCCTGGA A CCCGTGCGGTC TT AT CACGAAGTCAATAC CCGTACGCGTGAGG A CGTGAATGA GCCGAGGGTCGTTCCGTG AAA GTAGATACG CACGGAGCGACCTTCGGT A CCTCACGTGTACGG CCAGTTCGCCG A TCCTATGAG CACAGCGGTCGTGCTGATAGCGGTAGAGTCTGAGGTGCGGCAGTGG AAA TAGAATCGG CCACTGCTGCACCTCGGACTCTATCGCTATTAGCACGATCGCTGTG A TGGCGGACTGG TT AA CACGAAGTCAATAC GCCAAGGGTCACGC A TCTCGTTAG CCGGTGCGCTTTGCGCTG AAA TAGTCTACC CAGCGCGAAGCGTACCGG A GCGTGACTCTTGGC GCTCTGTGCAGG AATAATA GTCCTACTGGTTCGGGCATATCTCATAGTAGGACCGTGTCTTGCGG AAA CTCATAGGA CCGCAAGGCACGGTCTTACTATGGGATATGTCCGAACCGGTAGGAC CCTGCGCAGAGC TT AC CACGAAGTCAATAC CCAGTGTAGCCTCG A TACAAGTGC CTGGATTGACTTGGCGGTG AAA GCGTTACTA CACCGCTAAGTCGATCCAG A CGAGGCTGCACTGG CGTGCTGCAGG A CGTATCTAC CCTGCCTGAGGTGCCGGTG AAA GAAGTGATC CACCGGTACCTCGGGCAGG A CCTGCGGCACG TT TA CACGAAGTCAATAC GCACGTTCGGAGCC A GTTCCAATG TCGACGGAGTAAGGCGGTG AAA CAGAGATGA CACCGCTTTACTTCGTCGA A GGCTCCGGACGTGC GCTGGTGCTGC A GGTAGACTA GCCTACTTGTGCTAGCGTC AAA TCATTCACG GACGCTGGCACAGGTAGGC A GCAGCGCCAGC TT TAAT CACGAAGTCAATAC CCGTCTCGTGAACGG AAATAAA GTCCACTTCGTTTCCGGTG AAA GAACTTACG CACCGGGAACGAGGTGGAC CCGTTCATGAGACGG CCTGAGGTAGCG AATAATA GGTTAGTTCCGGGATCCGG AAA CTAACGAGA CCGGATTCCGGAGCTAACC CGCTATCTCAGG TT TACT CACGAAGTCAATAC CCGTCTGAGTTCGC A CATCGTTAC CCTGCTTGGCTTGCGGTG AAA CTTCTAAGC CACCGCAGGCCAAGCAGG A GCGAACTTAGACGG CCTCTGCCTGC A TCATCTCTG CCCGAGCGACGGTAGGTG AAA GCACTTGTA CACCTACTGTCGCTCGGG A GCAGGTAGAGG TT ATAT CACGAAGTCAATAC GCAGCTTCACAGGC A GACTTACTC GCCATTTCCGAGGCGGTG AAA GAACAGATC CACCGCCTCGGAAGTGGC A GCCTGTGGAGCTGC GCTGAGCCGAC A CGTAAGTTC GCGCAGGGTGCTACACGG AAA CATTGGAAC CCGTGTAGCACCCTGCGC A GTCGGTTCAGC TT AATT CACGAAGTCAATAC GTCCAACC TT CATGCTTACGACG | 1714 |
| **2D-h-Plasmid**<br>Template for 2D-h tiles | GTTCTAATACGACTCACTATA GGGCAATTAGTAGC A TGTTACCTC CGCTAAGTTGCTTTCGCAG AAA CTAGGAACA CTGCGAGAGCAATTTAGCG A GCTACTAGTTGCTC CCAGGTGTGCA A GCTTAGAAG GCATGCGTCGTCTGGTCGG AAA CAAGTGGAA CCGACCGGACGATGCATGC A TGCACGCCTGG TT AT CACGAAGTCAATAC GCCAGTTAGGACGG A TGACACTAG CCCTAATCTGTAGGCGGTG AAA GTTAGAACG CACCGCTTACAGGTTAGGG A CCGTCCTGACTGGC GCTCCGGCAGT A GATCTGTTC GCTTATTGAGCGTGAGGTG AAA GTAACGATG CACCTCGCGCTCGATAAGC A ACTGCTGGAGC TT AA CACGAAGTCAATAC CCTAGGCGACCTAGC AAATAAA GGCTACGCATGCTGCAGCG AAA GTCTAAGGA CGCTGCGGCATGTGTAGCC GCTAGGTTGCCTAGG CCATACGGAGCG AATAATA GGATAGGTGCCTTGTGCCG AAA GAGTAAGTC CGGCACGAGGCATCTATCC CGCTCTGTATGG TT AC CACGAAGTCAATAC CCAGAGGTATCGCC A GATTCATCC GGGTAGTAGAGGCTCCCG AAA GATGATCAG CGGGAGCTTCTACTACCC A GGCGATATCTCTGG CCAGGTAGTGC A CGTTCTAAC CCGCAGCGGTAGGTGCCG AAA GAGGTAACA CGGCACCTACCGCTGCGG A GCACTGCCTGG TT TA CACGAAGTCAATAC GCAGTGGCGTGTCC A GTCTGTAAG CCCGAGCCCACTGAGGCG AAA GTAGAGTCA CGCCTCAGTGGGCTCGGG A GGACACGTCACTGC GCACATACAGG A TCCTTAGAC ACCGAGCAGAGGCGTCGG AAA CTAGTGTCA CCGACGCTTCTGCTCGGT A CCTGTGTGTGC TT TAAT CACGAAGTCAATAC CCTAGCGAGTATCC A GTGTCAAAC GCCTGCTGTGAGTGCGTGG AAA TGATCGTAG CCACGCGCTCACGGCAGGC A GGATACTTGCTAGG CCCTGTGCTGT A CTGATCATC GCTATCTACAGCTGGAGGC AAA GGACATTCA GCCTCCGGCTGTGGATAGC A ACAGCGCAGGG TT TACT CACGAAGTCAATAC GCCAGTGCCAAGCC A CTCGATACA GTGCCGTTGTAGTAAGTGG AAA CAACATACG CCACTTGCTACAGCGGCAC A GGCTTGGTACTGGC GCTGGTGCAGG A TGACTCTAC CCTAAGTTTCGAGCGAGGC AAA GGATGAATC GCCTCGTTCGAAGCTTAGG A CCTGCGCCAGC TT ATAT CACGAAGTCAATAC CCAGCGTTCGAACGG AAATAAA GTCCCTGAGCTCTGACGTG AAA TGAATGAGC CACGTCGGAGCTTAGGGAC CCGTTCGGACGCTGG CCAGCATCCGAG AATAATA GCGAAGGTTCTGTAGACGG AAA CTTACAGAC CCGTCTGCAGAATCTTCGC CTCGGGTGCTGG TT AATT CACGAAGTCAATAC CCAGTATGCAACCG A TGACGATTC GGCTAGGTTCAGGTGGCG AAA GGAACTATC CGCCACCTGAACCTAGCC A CGGTTGCGTACTGG CGCTGTAATCC A CGTATGTTG CCAGCTCTTGGGCGTCCG AAA GTTTGACAC CGGACGCTCAAGAGCTGG A GGATTGCAGCG TT ACTT CACGAAGTCAATAC GTCCAACC TT CATGCTTACGACG | 1744 |

| Name (Note) | Sequence | length |
|---|---|---|
| **2D-i-Plasmid**<br>Template for 2D-i tiles | GTTCTAATACGACTCACTATA GGGCAGTCCGACCC A TACCAGTTC AGCGCTTACGAGGGCTCG<br>AAA CAATGATGG CGAGCCCTCGTAAGCGCT A GGGTCGGGCTGTCC GGTGTGCCGCT A<br>GCTCATTCA GGGACTGATCTTGACCGG AAA TGTATCGAG CCGGTCAGGATCAGTCCC A<br>AGCGGTACACC TT AT CACGAAGTCAATAC CCAGGGTCAGCGGC A CATACCTTG<br>GCCTCATTCGCATGCTCGC AAA GTTTCACAG GCGAGCGTGCGAGTGAGGC A<br>GCCGCTGGCCCTGG CCAGCGGGTGC A GATAGTTCC GCATGGGTAGACTGGTCTG AAA<br>GTACTTACC CAGACCGGTCTATCCATGC A GCACCTGCTGG TT AA CACGAAGTCAATAC<br>GCTCCTGTAGTCCC A TCGAGATAG GCCACTGCGGCTTGCGGTG AAA GCTTGATGA<br>CACCGCGAGCCGTAGTGGC A GGGACTATAGGAGC GCTGCTAGCCT A CCATCATTG<br>CGTCAATGCTTGTCCTGCG AAA GAATCGTCA CGCAGGGCAAGCGTTGACG A AGGCTGGCAGC<br>TT AC CACGAAGTCAATAC CCACCGTGACTGAGC AAATAAA GTCACTTGCCACGGACCTG AAA<br>CTTACTTGC CAGGTCTGTGGCGAGTGAC GCTCAGTTACGGTGG CCTGCATCGGTC AATAATA<br>GCTCTGTTCCGCGTGGCTG AAA GAACTGGTA CAGCCATGCGGAGCAGAGC GACCGGTGCAGG<br>TT TA CACGAAGTCAATAC GGGTGAGCTAACGC A GACATGGAA CCGCATCTAACTCGAGTG<br>AAA GATCTAACG CACTCGGGTTAGGTGCGG A GCGTTAGTTCACCC GCTCCTACCGC A<br>GCAAGTAAG TCCAGTCGTCATTCGCGG AAA CTATCTCGA CCGCGAGTGACGGCTGGA A<br>GCGGTGGGAGC TT TAAT CACGAAGTCAATAC GCCACATCGACGCC A GTATGCAGA<br>GCCTTCTGGACATGCCTCG AAA CTATGCAAG CGAGGCGTGTCCGGAAGGC A<br>GGCGTCGGTGTGGC GCTGCTCAGGT A GGTACAAAG GGAATGTAATACTGACCGG AAA<br>TCCTAACTG CCGGTCGGTATTGCATTCC A ACCTGGGCAGC TT TACT CACGAAGTCAATAC<br>CCTACCGTGTGCTC A CTATGCCTA GGACTCTCGATGTGACCGG AAA CTGTCTACA<br>CCGGTCGCATCGGGAGTCC A GAGCACATGGTAGG CCCTGTCGCTC A GACTCATTG<br>CGTCCTGCGCCATCGGGAC AAA GATTCTCAG GTCCCGGTGGCGTAGGACG A GAGCGGCAGGG<br>TT ATAT CACGAAGTCAATAC GCAGGTTCATTACC A TAGTTCGTC<br>GGTAAGGGCGTCGCACGTG AAA TTCTGATCC CACGTGTGACGCTCTTACC A<br>GGTAATGGACCTGC GCAGGGCCAGG A CGTTAGATC GCTACAGTCACATGGCGTG AAA<br>GACTAAGTG CACGCCGTGTGATTGTAGC A CCTGGTCCTGC TT AATT CACGAAGTCAATAC<br>CCTCGGTGGTACGAC AAATAAA CCTCGTTCATCCGGCTCGC AAA TGTTCACTC<br>GCGAGCTGGATGGACGAGG GTCGTACTACCGAGG CCCTGTGCCTGG AATAATA<br>GCTGCGTACTGGGCAGCCG AAA TTCCATGTC CGGCTGTCCAGTGCGCAGC CCAGGTACAGGG<br>TT ACTT CACGAAGTCAATAC GTCCAACC TT CATGCTTACGACG | 1748 |
| **2D-j-Plasmid**<br>Template for 2D-j tiles | GTTCTAATACGACTCACTATA GGGCGTGCTCAGGC A TCGACTAAC GCTACGGTAAGAGCCTCGC<br>AAA GTACATTCC GCGAGGTTCTTATCGTAGC A GCCTGAGTACGCTC CCAGCGTACGG A<br>GTCAAGATG CCTCGAGTACTATCCGAGC AAA TCTTCATCG GCTCGGGTAGTATTCGAGG A<br>CCGTATGCTGG TT AT CACGAAGTCAATAC GCAGTGTGACTCGG A CACAGTATC<br>GGAGCAGAG GTAA CTCTGCTCC A CCGAGTCGCACTGC GCTGCGGATCC A GGAATGTAC<br>GCGTCCAAT TTCG ATTGGACGC A GGATCTGCAGC TT AA CACGAAGTCAATAC<br>CCAGCCTTGTGAGC A GGTCTTCAA GCGGATACTTAGGCAGCG AAA GTCACATCA<br>CGCTGCTTAAGTGTCCGC A GCTCACAGGGCTGG CCTGTTCCGCA A GATACGAGA<br>CTCGGTACAGTTTCGGGC AAA GTTAGTCGA GCCCGAGACTGTGCCGAG A TGCGGACAGG TT<br>AC CACGAAGTCAATAC GCACATTTCCAGCC A CGTACAATC GGCAAGCGTACGGCACGG AAA<br>TGAGTTGAC CCGTGCTGTACGTTTGCC A GGCTGGAGATGTGC GGTCTTAAGGC A<br>CTTTGAACG CCCAGTACCTGGCAGGTG AAA GTTACGGTA CACCTGTCAGGTGCTGGG A<br>GCCTTGAGACC TT TA CACGAAGTCAATAC CCTCGCGAGAGTCC A CATCTAGAG<br>GTCGGGTCTGAGGCGGTG AAA CTGATACTC CACCGCTTCAGATCCGAC A GGACTCTTGCGAGG<br>CCAACGTACGG A TGATAGCTG CGCCTGCGTGAGCCGGTG AAA GATTGACTG<br>CACCGGTTCACGTAGGCG A CCGTATGTTGG TT TAAT CACGAAGTCAATAC<br>GCCTGAGGTACGGC A CGTGTAAGA TCCGAGTTTGAGGCGTGG AAA TTGCAAGAG<br>CCACGCTTCAAATTCGGA A GCCGTACTTCAGGC GCATGTTAGGG A TTGGACTTG<br>CCTGCGGATGAGCCCTGG AAA CGTAACTAG CCAGGGTTCATCTGCAGG A CCCTAGCATGC TT<br>TACT CACGAAGTCAATAC CCGAGCGCTCTGCC A CAGTTAGGA GCGAATGTCGTGGCGGTG<br>AAA GCGATAGTA CACCGCTACGACGTTCGC A GGCAGAGTGCTCGG CCTCCGAGACG A<br>TGAGAATGC GGAGCGGGTGATGTTCGG AAA TACAACTGG CCGAACGTCACCTGCTCC A<br>CGTCTTGGAGG TT ATAT CACGAAGTCAATAC GCATCAGCAGTCCC A CTGAGAATC<br>GCCAGGTACATTGCGCTG AAA CTTTGTACC CAGCGCGATGTATCTGGC A GGGACTGTTGATGC<br>GCGAGTGGGAC A CTGTGAAAC GCTGCGGTCTCGTTCCGC AAA CATTCTGAG<br>GCGGAATGAGACTGCAGC A GTCCCGCTCGC TT AATT CACGAAGTCAATAC<br>CCAAGGGTCATAGC A CACTTAGTC GCGACGCTGGCTGAGGTG AAA CAATGAGTC<br>CACCTCGGCCAGTGTCGC A GCTATGATCCTTGG CCCTGGACGAC A TCATCAAGC<br>CCTGCTGTCTATACTCGG AAA CAAGGTATG CCGAGTGTAGACGGCAGG A GTCGTTCAGGG TT<br>ACTT CACGAAGTCAATAC GTCCAACC TT CATGCTTACGACG | 1680 |

| Name (Note) | Sequence | length |
|---|---|---|
| **2D-k-Plasmid**<br>Template for 2D-k tiles | GTTCTAATACGACTCACTATA GGGCGATGGTAGCC A GTATCCATG TCGCAGTAGCTAGGCGAGC<br>AAA GGATTCACA GCTCGCTTAGCTGCTGCGA A GGCTACCGTCGCTC CCAGGTCTGCT A<br>TGATGTGAC GCTAAGGTCACTGCCGGTG AAA GATACTGTG CACCGGTAGTGATCTTAGC A<br>AGCAGGCCTGG TT AT CACGAAGTCAATAC GCAGCGTATCGGGC A CTAGAGATC<br>GCAACGGGTTAGTGCAGTG AAA GTTCAGGAA CACTGCGCTAACTCGTTGC A<br>GCCCGATGCGCTGC GCTGGTCGAGC A GTCAACTCA GCTATCTAGCTCGCCTCTG AAA<br>TTGAAGACC CAGAGGTGAGCTGGATAGC A GCTCGGCCAGC TT AA CACGAAGTCAATAC<br>CCGAGGGTAGTGAG A TAGCCAATG CCTTACTTATTGGCCTCGG AAA CAGTAGAGA<br>CCGAGGTCAATAGGTAAGG A CTCACTATCCTCGG CCAGCGCGTGC A GAGTATCAG<br>GCGTCAGTACCCGGTCCTG AAA GATTGTACG CAGGACTGGGTATTGACGC A GCACGTGCTGG<br>TT AC CACGAAGTCAATAC GAGCATTATGGGCC A GCAGTTCTA CCGTACGTGGAAGCCTCGG<br>AAA TAGCTCTAG CCGAGGTTTCCATGTACGG A GGCCCATGATGCTC GCCCTGTCCGA A<br>CTCTTGCAA GGACATTGCGAGGCCGTGG AAA CTCTAGATG CCACGGTCTCGCGATGTCC A<br>TCGGATAGGGC TT TA CACGAAGTCAATAC CCTCCGTTGTAGGC A TCACTCTTG<br>CGCCTGTTGCTTGGCGGTG AAA GGTATCGTA CACCGCTAAGCAGCAGGCG A<br>GCCTACAGCGGAGG CATCCGGGAGT A TACTATCGC AGGGTCTCGGTCTATCGTG AAA<br>TCTTACACG CACGATGGACCGGGACCCT A ACTCCTGGATG TT TAAT CACGAAGTCAATAC<br>GCCTCGTGATCTAC A TCCTAAGTC TGTAACTGC GTAA GCAGTTACA A GTAGATCGCGAGGC<br>GCTACTAGTAC A TGTGAATCC GGGAGTCTA TTCG TAGACTCCC A GTACTGGTAGC TT<br>TACT CACGAAGTCAATAC CCACATGCAGATCC A GACCAATCA GGCTGTCAGTAGGCGCAG<br>AAA TACGGAAAC CTGCGCTTACTGGCAGCC A GGATCTGTATGTGG GTACTGGTGCC A<br>TTCCTGAAC GGCAGTCTTGTGCCTCGG AAA CATGGATAC CCGAGGTACAAGGCTGCT A<br>GGCACTAGTAC TT ATAT CACGAAGTCAATAC CCAGGCTG TTCG CAGCCTGG CGCTGCAC A<br>GTTTCCGTA GGGCTGTGAATCGCAGCTG AAA GACTTAGGA CAGCTGTGATTCGCAGCCC A<br>GTGCAGCG TT AATT CACGAAGTCAATAC GCTCGAGG TTCG CCTCGAGC GGCTCCGC A<br>GTCTGAATC TGCACGTGTATCGTCGGTG AAA TGATTGGTC CACCGATGATACGCGTGCA A<br>GCGGAGCC TT ACTT CACGAAGTCAATAC CCAGCGTC TTCG GACGCTGG CCCTCGCC A<br>GTATGACGA GCCACGTGGATATCTCGTG AAA TTGCCTATC CACGAGGTATCCGCGTGGC A<br>GGCGAGGG TT TATA CACGAAGTCAATAC GCCAGCAC TTCG GTGCTGGC GCTCCGGGT A<br>CTTGTGATG CGGGAGTCAACGGCTATCG AAA GTAACTCTC CGATAGTCGTTGGCTCCCG A<br>ACCCGGAGC TT ATTA CACGAAGTCAATAC GTCCAACC TT CATGCTTACGACG | 1790 |
| **2D-l-Plasmid**<br>Template for 2D-l tiles | GTTCTAATACGACTCACTATA GGGCATGTAGATTC A GATAGGCAA GTCCGTATAAGGATAGTC<br>AAA GATTCAGAC GACTATTCTTATGCGGAC A GAATCTATATGTCC GCAAGTGCAGC A<br>TCTCTACTG TCGTCTGTGGCGAGGGTG AAA GATCTCTAG CACCCTTGCCACGGACGA A<br>GCTGCGCTTGC TT AT CACGAAGTCAATAC CCTCGAGCAGCGCC A GAGAGTTAC<br>GGGTATAAGGGTGAGCGG AAA TCGTCATAC CCGCTCGCCCTTGTACCC A GGCGCTGTTCGAGG<br>CCCTGTAATCG A CTAGAGCTA CGCGTGCAGCAGTTCCGG AAA CATTGGCTA<br>CCGGAATTGCTGTACGCG A CGATTGCAGGG TT AA CACGAAGTCAATAC GCCTTATTGTGCCC<br>A CTTTCATGC AGGGTGCCGCTGAAGTCG AAA CATCACAAG CGACTTTAGCGGTACCCT A<br>GGGCACAGTAAGGC GCTCCGTGCCG A TACGATACC CTCGCTGACCTGTAGCTG AAA<br>TAGAACTGC CAGCTATAGGTCGGCGAG A CGGCATGGAGC TT AC CACGAAGTCAATAC<br>CAGATCTCTGACGG A CTGGATAGA CGCGTTGCACATGCCTGG AAA TCTGCTTAG<br>CCAGGCGTGTGCGACGCG A CCGTCAGGGATCTG CCTCTGAGCGG A CTTGCATAG<br>CCTGGTGAAGCGCAGCGG AAA CAAGAGTGA CCGCTGTGCTTCGCCAGG A CCGCTTAGAGG TT<br>TA CACGAAGTCAATAC GCAGGAGCTAATCC A CTCAAGTTG CAGGGTTCTTGTCCAGTG AAA<br>GTGACAGAA CACTGGGCAAGAGCCCTG A GGATTAGTTCCTGC GCAGGTGTGGG A<br>TGTAGACAG CCCTGGCACAGTCCTCGG AAA TCTGCATAC CCGAGGGCTGTGTCAGGG A<br>CCCACGCCTGC TT TAAT CACGAAGTCAATAC CCAGCGTTCGAGTC A TCCTAGTAC<br>TGGCGGAACCGGGACGTG AAA GTTAGCTAC CACGTCTCGGTTTCGCCA A GACTCGAGCGCTGG<br>GCTTATGTACC A GGATCAGAA GCTCCTGGTATGCCGGTG AAA TAGGCATAG<br>CACCGGTATACCGGGAGC A GGTACGTAAGC TT TACT CACGAAGTCAATAC<br>GCCATACTGTTACGC AAATAAA CCTGCGAATACGGCCTGG AAA CTACTGGAA<br>CCAGGCTGTATTTGCAGG GCGTAACGGTATGGC GCTACGGTACC A GAGTGAACA<br>GCCTCTGTCATGGCGGTG AAA GACGAACTA CACCGCTATGACGGAGGC A GGTACTGTAGC TT<br>ATAT CACGAAGTCAATAC GCAGGCAG TTCG CTGCCTGC CCCTCCGG A CTAAGCAGA<br>CCCGTCTACTCGGCATCCC AAA GCATGAAAG GGGATGTCGAGTGGACGGG A CCGGAGGG TT<br>AATT CACGAAGTCAATAC GCCTCACC TTCG GGTGAGGC GCAGCGCC A TTCTGTCAC<br>CCGGAGTGCGATTAACGTC AAA TCTATCCAG GACGTTGATCGCGCTCCGG A GGCGCTGC TT<br>ACTT CACGAAGTCAATAC CCAGGCTG TTCG CAGCCTGG TCTCCGGG A GTAGCTAAC<br>ACGGACGGTTGATTCCTCG AAA CAACTTGAG CGAGGAGTCAACTGTCCGT A CCCGGAGA TT<br>TATA CACGAAGTCAATAC GCAGGGTC TTCG GACCCTGC GCTCGCCC A TTCCAGTAG<br>GGCGTCGGGATATCTCGTC AAA GTACTAGGA GACGAGGTATCCTGACGCC A GGGCGAGC TT<br>ATTA CACGAAGTCAATAC GTCCAACC TT CATGCTTACGACG | 1818 |

#### 6) Bricks for 3D assembly

**3D_23** (23 bricks): 3D-a + 3D-b + 3D-za + 3D-yz.

**3D_71** (71 bricks): 3D-a + 3D-b + 3D-c + 3D-d + 3D-e + 3D-f + 3D-g + 3D-xy + 3D-xz.

**3D_118** (118 bricks): 3D-a + 3D-b + 3D-c + 3D-d + 3D-e + 3D-f + 3D-g + 3D-h + 3D-i + 3D-j + 3D-k + 3D-l + 3D-m + 3D-n.

| Name (Note) | Sequence | length |
|---|---|---|
| **3D-a-L0-0** | GGGCAAGUCGCUGCA **UUCGACAUG** UAUCCAGGGACCCGG AAA **CGAAUGCAA**<br>CCGGGUCUCUGGAUA AGCAGCGAUUUGCCC GCUCCUGAGACA **CUGCUAACC** CGAUCGUGC<br>**UUCG** GCACGAUCG AGUCUCGGGAGC **UU AU CACGAAGUCAAUAC** | 154 |
| **3D-a-L0-3** | CCAUAUUGGAGCCCA **GUCUCAUGA** GCCACCGUCACCGACG AAA **GAAGGUAUC**<br>CGUCGGUGGCGGUGGC AGGGCUCCGAUAUGG GCUCCUGCUGCA **GUAUCUCCA**<br>GCCCAGGCGUGUUCAUGCUUGUACUAAGAUCUUAUCAGCGG AAA **CAUGUCGAA**<br>CCGCUGAUGAGAUCUUGGUACAAGUAUGAACAUGCCUGGGU AGCAGCGGGAGC **UU AA**<br>**CACGAAGUCAAUAC** | 228 |
| **3D-a-W0-0** | CCUGUAGCCUCGCCA **CGUAAGUUC** GGAUACUGA **GUAA** UCAGUAUCC AGGCGAGGUUACAGG<br>GCUCGUCCUGCA **UUGCAUUCG** GGUAUGAGA **UUCG** UCUCAUACC AGCAGGGCGAGC **UU AC**<br>**CACGAAGUCAAUAC** | 134 |
| **3D-a-W0-1** | CAGAGGUAACGGCCA **CUUUGCUCA** CCGACAGCGCUCAGCUGUCGGAUGUCCAGUAGGACGACCGG<br>AAA **CAAUCGACA** CCGGUCGUUCUACUGGGCAUCCGAUAGCUGAGUGCUGUCGG<br>AGGCCGUUGCCUCUG GCCGUGCGAGCA **CAGAACUCA** CCUACGAUCGUAAGCG AAA<br>**UGGAGAUAC** CGCUUACGGUCGUAGG AGCUCGUACGGC **UU UA CACGAAGUCAAUAC** | 228 |
| **3D-za-L0-1** | GGGCAGAUCCUUGCA **CCGAUUCUA** AUGUACGGGAUCCGC AAA **UGAGUUCUG**<br>GCGGAUCUCGUACAU AGCAAGGGUCUGUCC GCAUGGGCAUGG **AAUAAUA**<br>CACUCAAGCUAUGGGAUGAGGCGUAGUUAGCAGAUAGUCAGUCCUUGCUUUCGGACACGUAGGAUCAG<br>CGAAGGUCCAAUGUUACCGGUG AAA **GGUUAGCAG**<br>CACCGGUGACAUUGGGCCUUCGUUGAUCCUGCGUGUCUGAAAGCAGGGACUGGCUAUCUGUUAACUAU<br>GCCUCAUUCCAUAGUUUGAGUG CCAUGCUCAUGC **UU AU CACGAAGUCAAUAC** | 322 |
| **3D-za-L0-4** | GCCUACGUCAGGCUG **AAAUAAA** GACUAACUCGACCGUG AAA **GUAACCUGA**<br>CACGGUCGGGUUAGUC CAGCCUGAUGUAGGC CCAGGGCAGGGU **AAUAAUA**<br>CGACUUCAUCAGAAUGGGCCUAGGGCUUCGCAGGGACCUGG AAA **UAGAAUCGG**<br>CCAGGUCCUUGCGAAGUCCUAGGCUCAUUCUGGUGAAGUCG ACCCUGUCCUGG **UU AA**<br>**CACGAAGUCAAUAC** | 224 |
| **3D-za-L1-0** | CCAGUGGGACACGGA **GUGCAUUGA** CCUGACAGC **GUAA** GCUGUCAGG ACCGUGUCUCACUGG<br>GAGUCGCCUGCA **GACAGUUGA** GGGCAAUGCCGAGUG AAA **GAACUUACG** CACUCGGUAUUGCCC<br>AGCAGGUGACUC **UU AC CACGAAGUCAAUAC** | 154 |
| **3D-za-L1-1** | CCAGACGGAAGCCCA **UUCCACUUG** CAUAGUGUCUACCGG AAA **CUAACUCUG**<br>CCGGUAGGCACUAUG AGGGCUUCUGUCUGG GCAGAGUCAUGC **AAUAAUA** CCAGAUCUCGGACGG<br>AAA **UGAGCAAAG** CCGUCCGGGAUCUGG GCAUGAUUCUGC **UU UA CACGAAGUCAAUAC** | 172 |
| **3D-za-W0-3** | GCAGCGUC **UUCG** GACGCUGC CCAGGCGCA **UGUCGAUUG** CCGGUGAGGUCAGGUG AAA<br>**UCAACUGUC** CACCUGACUUCACCGG AGCGCCUGG **UU UAAU CACGAAGUCAAUAC** | 111 |
| **3D-za-W1-4** | CCAGAGCC **UUCG** GGCUCUGG GGCUCCGCA **CAGAGUUAG** CCACGGUUACCGAGUG AAA<br>**GAUGCAAUG** CACUCGGUGACCGUGG AGCGGAGCC **UU UACU CACGAAGUCAAUAC** | 111 |
| **3D-b-L0-6** | GGGCACUGUUAGCCA **GGUACUACA** GGCGAGGGUGCGGUG AAA **GUAAGAUCG**<br>CACCGCAUCCUCGCC AGGCUAACGGUGUCC GACACGGUCGCA **GCUUCAAUC** GUCGGUCAC<br>**UUCG** GUGACCGAC AGCGACUGUGUC **UU AU CACGAAGUCAAUAC** | 154 |
| **3D-b-L1-3** | CCAUGUGGCUGCCGA **CAUGAUUGC** UGCCUUGGAGUUCCCG AAA **GCUAGUUAG**<br>CGGGAACUUCAAGGCA ACGGCAGCUACAUGG CCAGUUCUAGGA **CAUUGCAUC**<br>UCCGGGAGUGCCGCUG AAA **UCAGUACUC** CAGCGGCAUUCCCGGA ACCUAGGACUGG **UU AA**<br>**CACGAAGUCAAUAC** | 178 |
| **3D-b-W1-0** | GCCUGAUUCGUAGCA **GAGUACUGA** UAACUACGAGCGGAUG AAA **UCAAUGCAC**<br>CAUCCGCUUGUAGUUA AGCUACGAGUCAGGC CCAGAUCAUGCA **GAUACCUUC**<br>GAGCGGAGUUUCGGUG AAA **GAUUGAAGC** CACCGAAAUUCCGCUC AGCAUGGUCUGG **UU AC**<br>**CACGAAGUCAAUAC** | 178 |
| **3D-b-W1-1** | CCGUCAUUACUGCAA **GCUUAGAAG** GUCGGUCUGCGACCGG AAA **CAAGUGGAA**<br>CCGGUCGCGGACCGAC AUGCAGUAGUGACGG CCUACUUCGCCA **UCAGGUUAC**<br>ACGGGCAUCGCAGCUG AAA **GUGUACCUA** CAGCUGCGGUGCCCGU AGGCGAGGUAGG **UU UA**<br>**CACGAAGUCAAUAC** | 178 |

| Name (Note) | Sequence | length |
|---|---|---|
| **3D-b-W2-0** | GCCAGAUAUGACCCA **UCGAUAGAG** GCGUCAGAU **GUAA** AUCUGACGC AGGGUCAUGUCUGGC<br>GGAGGUGCUGGA **CGAUCUUAC** GCCGUCCUG **UUCG** CAGGACGGC ACCAGCGCCUCC **UU UAAU**<br>**CACGAAGUCAAUAC** | 136 |
| **3D-b-W2-1** | GGGAACGGUAUGCCA **CAUACAUGG** GCGUGUCUACAUCCGC AAA **GCAAUCAUG**<br>GCGGAUGUGGACACGC AGGCAUACUGUUCCC GCUAGUUAUGGA **CGUAUCUAC**<br>CCUGGUCGUCUCACUG AAA **GAAGUGAUC** CAGUGAGAUGACCAGG ACCAUAGCUAGC **UU UACU**<br>**CACGAAGUCAAUAC** | 180 |
| **3D-yz-L0-7** | GGGCAUCUAUGUCGC **AAAUAAA** CCUUGGCUGAGGGUG AAA **GUAGAUACG**<br>CACCCUCGGCCAAGG GCGACAUGGAUGCCC CGAGAGCAGCCA **UAGGUACAC**<br>GCGCUACGGAGUAUGUCGGAUUGUAUUUCUGGUUCAUCGG AAA **UCAUGAGAC**<br>CCGAUGAAUCAGAAAUGCAAUCCGGCAUACUCUGUAGCGC AGGCUGUUCUCG **UU AU**<br>**CACGAAGUCAAUAC** | 222 |
| **3D-yz-L0-9** | CCUGGAGG **UUCG** CCUCCAGG CCCUGCCCA **GAUCACUUC**<br>GCCUCCGCUUCACGACUUAGUCGCUUGCGGUUUAGGCGCUG AAA **UGUAGUACC**<br>CAGCGCCUGAACCGCAGGCGACUAGGUCGUGAGGCGGAGGC AGGGCAGGG **UU AA**<br>**CACGAAGUCAAUAC** | 159 |
| **3D-yz-L1-4** | GCCAAGGUCCGCCUG **AAAUAAA** GGCAGGAGCGCUCCGG AAA **UGUCAGAAG**<br>CCGGAGCGUUCCUGCC CAGGCGGAUCUUGGC GCUAGUUAGGGC **AAUAAUA**<br>GUGGUCCUGACCGGUG AAA **CUUCUAAGC** CACCGGUCGGGACCAC GCCCUAGCUAGC **UU AC**<br>**CACGAAGUCAAUAC** | 174 |
| **3D-yz-L1-6** | CCGUCUGAG **UUCG** CUCAGACGG CCUCCGGCA **GCAUACUUC** ACGGACCGGGAGCGG AAA<br>**CUCUAUCGA** CCGCUCCUGGUCCGU AGCCGGAGG **UU UA CACGAAGUCAAUAC** | 109 |
| **3D-yz-L1-7** | CCAGGGUC **UUCG** GACCCUGG CCAGCCGAA **GUGACUGUA** UCCAGACGGCGAGUG AAA<br>**CCAUGUAUG** CACUCGCUGUCUGGA AUCGGCUGG **UU UAAU CACGAAGUCAAUAC** | 109 |
| **3D-yz-W2-3** | CCUCGGGA **UUCG** UCCCGAGG GCUCGCGCA **CUAACUAGC** CAGGUCCGCUUGGCUG AAA<br>**GAAGUAUGC** CAGCCAAGUGGACCUG AGCGCGAGC **UU UACU CACGAAGUCAAUAC** | 111 |
| **3D-yz-W2-4** | CCACCUGC **UUCG** GCAGGUGG CCCUCGCCA **CUUCUGACA** GCGCGUAGAGAUUCGG AAA<br>**UACAGUCAC** CCGAAUCUUUACGCGC AGGCGAGGG **UU AUAU CACGAAGUCAAUAC** | 111 |
| **3D-zz-L0-7** | GGGCUGUC **UUCG** GACAGCCC GACCGUACGGGA **UAGGUACAC**<br>GCUCGGAUGACGUUCGGAGAUUCUGAAGUUCUAAGCCGUG AAA **UCAUGAGAC**<br>CACGGCUUGGAACUUCGGAAUCUCUGAACGUCGUCCGAGC ACCCGUGCGGUC **UU AU**<br>**CACGAAGUCAAUAC** | 163 |
| **3D-zz-L1-3** | CCAGGACC **UUCG** GGUCCUGG GACAGUCGGCCA **CAUUGCAUC** ACCUGCCGUAGCUGUG AAA<br>**UCAGUACUC** CACAGCUAUGGCAGGU AGGCCGGCUGUC **UU AA CACGAAGUCAAUAC** | 115 |
| **3D-zz-W1-0** | CCGUAGGCCUAAGCA **GAGUACUGA** UAGAGAAUACUCGUCG AAA **UCAAUGCAC**<br>CGACGAGUGUUCUCUA AGCUUAGGUCUACGG CUCCGUCGUGCA **GAUACCUUC** GCGCAUAGU<br>**UUCG** ACUAUGCGC AGCACGGCGGAG **UU AC CACGAAGUCAAUAC** | 156 |
| **3D-zz-W1-1** | GCCUGUUAAGUAGCG **AAAUAAA** GGUAUGAGAGCGUCGG AAA **CAAGUGGAA**<br>CCGACGCUUUCAUACC CGCUACUUGACAGGC CCUCAGUCCGCA **UCAGGUUAC**<br>GGCACCGGAAGUCCUG AAA **GUGUACCUA** CAGGACUUUCGGUGCC AGCGGAUUGAGG **UU UA**<br>**CACGAAGUCAAUAC** | 176 |
| **3D-c-L0-1** | GGGCAUUGGGCUGCA **CCGAUUCUA** GUUAUCAUCCGAGGC AAA **UGAGUUCUG**<br>GCCUCGGGUGAUAAC AGCAGCCCGAUGCCC GACCUGGGACCA **CUACUCGUA**<br>CCGGCUAUGUUUAGGUGAUGGUGAACAUUUCGUUCAAUAGGCGUGGUAUAUGUUAGAUCGUAAUCGAG<br>CUGUAGUCGAUGAGGGCUCGUG AAA **GGUUAGCAG**<br>CACGAGCUCUCAUCGGCUACAGUUCGAUUAUGAUCUAGCAUAUACUACGCCUGUUGAACGGAAUGUUU<br>ACCAUCAUCUAAACGUAGCCGG AGGUCCUAGGUC **UU AU CACGAAGUCAAUAC** | 324 |
| **3D-c-L0-2** | CCGACCGUGGGACCA **GUUGCUCUA** UAACUUGUGCUCGUG AAA **GUCUUGAAG**<br>CACGAGCGCAAGUUA AGGUCCCAUGGUCGG GCAUCAGGAGCC **AAUAAUA**<br>GCUCCCAUUCACAGUGCUGAGUGGUACAUUGAUGGCAGUAGAUUGGGCAGGAGUAACUAUGAUACAAU<br>GUUACUUAAUCACGGAGGUCGG AAA **UACGAGUAG**<br>CCGACCUUCGUGAUUGAGUAACGUUGUAUCGUAGUUAUUCCUGCCUAAUCUAUUGCCAUCGAUGUACU<br>ACUCAGCGCUGUGAGUGGGAGC GGCUCUUGAUGC **UU AA CACGAAGUCAAUAC** | 322 |
| **3D-c-L0-4** | GCCUCCGGUGGCGCA **GAGAACAUG** CCACGAGUGCGUUGCG AAA **GUAACCUGA**<br>CGCAACGCGCUCGUGG AGCGCCACUGGAGGC CCAGUGCGCCUA **UCCUAUGAG**<br>CAGUAGCGUACGAAGUGUGAAACGGAUGAUCAGGGACGUGG AAA **UAGAAUCGG**<br>CCACGUCCUUGAUCAUUCGUUUCAUACUUCGUGCGCUACUG AAGGCGUACUGG **UU AC**<br>**CACGAAGUCAAUAC** | 228 |

| Name (Note) | Sequence | length |
|---|---|---|
| **3D-c-L0-5** | GCAUUCCGAAGACCG **AAAUAAA** GCAGGGUGCGCCUCUG AAA **GUAAGGACA** CAGAGGCGUACCCUGC CGGUCUUUGGAAUGC CCGUGGGACUCG **AAUAAUA** GGAUGUACGGAUUGCGGAGUACAUUGAUCUCUGGAAGCCAG AAA **UAGAGCAAC** CUGGCUUCUAGAGAUCGAUGUACUUCGCAAUCUGUACAUCC CGAGUUCCACGG **UU UA CACGAAGUCAAUAC** | 224 |
| **3D-c-L0-7** | GCACGGGCAUCCGGA **CGUGAAUGA** CCAGUGGUCCAGGUG AAA **GUAGAUACG** CACCUGGGCCACUGG ACCGGAUGUCCGUGC GCUCGUAGCGGA **UAGGUACAC** GGCGACUUGAUAUUGGCUUUAGCUGAGUCGCUAGGGCCUG AAA **UCAUGAGAC** CAGGCCCUGGCGACUCGGCUAAAGUCAAUAUCGAGUCGCC ACCGCUGCGAGC **UU UAAU CACGAAGUCAAUAC** | 226 |
| **3D-c-L0-8** | GCCAGAGCAUGACCA **GUAACAAGC** CCGAGGCGGACGGUG AAA **CUAACGAGA** CACCGUCUGCCUCGG AGGUCAUGUUCUGGC CCAUCGGAGGCA **CUUGGAUCA** GCGCAGGUUGAAGAUGAUCCAUAGGCUGGAUUCCAAUCGC AAA **CAUGUUCUC** GCGAUUGGGAUCCAGCUUAUGGAUUAUCUUCAGCCUGCGU AGCCUCUGAUGG **UU UACU CACGAAGUCAAUAC** | 226 |
| **3D-d-L0-9** | GGGCUCUGCCAGGCA **CCUUGAGAA** CCGUGCGUGAUCCUGG AAA **UUCAUAGCG** CCAGGAUCGCGCACGG AGCCUGGCGGAGCUC GCCCUUGCGGUA **GAUCACUUC** GCUCGAAUUCAGCUCGUAGGCAAGGCAGACAAUUGCAACUG AAA **UGUAGUACC** CAGUUGCAGUUGUCUGUCUUGCCUGCGAGCUGGAUUCGAGC AACCGCGAGGGC **UU AU CACGAAGUCAAUAC** | 228 |
| **3D-d-L1-0** | CAGUGUGGACGAGCA **GUGCAUUGA** UCUGACAUAGCGGUG AAA **UACAUGUGG** CACCGCUGUGUCAGA AGCUCGUCUACACUG GCUCGGUGCGCA **GACAGUUGA** CCGUACGUGGUCGUG AAA **GAACUUACG** CACGACCGCGUACGG AGCGCAUCGAGC **UU AA CACGAAGUCAAUAC** | 174 |
| **3D-d-L1-1** | GCCAUGGUAGCAGCA **UUCCACUUG** GCCGUUCGACGACGG AAA **CUAACUCUG** CCGUCGUUGAACGGC AGCUGCUAUCAUGGC GUGCGUUCUGGA **UAGCCAUUC** AGGGAUGUCCAGCGG AAA **UGAGCAAAG** CCGCUGGGCAUCCCU ACCAGAGCGCAC **UU AC CACGAAGUCAAUAC** | 174 |
| **3D-d-L1-2** | GAGCGGUAGGUAGCA **GACUUACUC** CAUCCGUGAUGCGUG AAA **GAACAGAUC** CACGCAUUACGGAUG AGCUACCUGCCGCUC CCAGGUCCUGCA **UGCAGAUUC** GACCAGGUGCUCGUG AAA **GGUUACAUG** CACGAGCGCCUGGUC AGCAGGGCCUGG **UU UA CACGAAGUCAAUAC** | 174 |
| **3D-d-L1-4** | CCAUGGGUGAUCGCA **GAUGACUUG** GCCAGACGCCGUCCGG AAA **UGUCAGAAG** CCGGACGGUGUCUGGC AGCGAUCAUCCAUGG GAGUCUCUGCCA **CAUCGUUAC** CCGCUGAGACAGUCGG AAA **CUUCUAAGC** CCGACUGUUUCAGCGG AGGCAGGGACUC **UU UAAU CACGAAGUCAAUAC** | 180 |
| **3D-d-L1-5** | CCUCAACGGUCACGG **AAAUAAA** GCUCGAGUUCCGACGG AAA **CUAGCAUGA** CCGUCGGAGCUCGAGC CCGUGACUGUUGAGG GCACCGUAUAGG **AAUAAUA** CCGACUCGCCUCGGUC AAA **CAUUCGUAC** GACCGAGGUGAGUCGG CCUAUGCGGUGC **UU UACU CACGAAGUCAAUAC** | 176 |
| **3D-d-L1-6** | GCAGUCUGUAGCGCA **UGUUACCUC** GCCGUCCGAGACCAG AAA **CUAGGAACA** CUGGUCUUGGACGGC AGCGCUACGGACUGC CCAUCUCAGGCA **GCAUACUUC** UACCGGUUCACGCGG AAA **CUCUAUCGA** CCGCGUGGACCGGUA AGCCUGGGAUGG **UU AUAU CACGAAGUCAAUAC** | 176 |
| **3D-d-L1-7** | GCAGGAGCAGAUCCA **GGCUAUGUA** UUGUAACGGUGAGCG AAA **GUCUAAGGA** CGCUCACUGUUACAA AGGAUCUGUUCCUGC CGUGCGAAGGAA **GUGACUGUA** UCUACCGUGAGGGUG AAA **CCAUGUAUG** CACCCUCGCGGUAGA AUCCUUUGCACG **UU AAUU CACGAAGUCAAUAC** | 176 |
| **3D-d-L1-8** | GCCUCGUACAGAGCA **UUCCGAAAG** CGCAGGUUAUCCGCG AAA **GAAGUCUCA** CGCGGAUGACCUGCG AGCUCUGUGCGAGGC GCUCGGACGGGA **UUCAUUGCC** ACCGGCUGCGACGUG AAA **GAACAUAGG** CACGUCGUAGCCGGU ACCCGUUCGAGC **UU ACUU CACGAAGUCAAUAC** | 176 |
| **3D-e-L2-0** | GGGCCAUAGCUAGGA **UGAGCAUUC** GCCGAUCUGGACGGUG AAA **CAGAUGUAG** CACCGUCCGGAUCGGC ACCUAGCUGUGGCCC CCGGAUAGGCCA **GCAUAGAUG** ACCAGGCGAUAUCCGG AAA **UCUGAAGUG** CCGGAUAUUGCCUGGU AGGCCUGUCCGG **UU AU CACGAAGUCAAUAC** | 178 |
| **3D-e-L2-1** | CGACCUUACCGAGCA **CUCGAUACA** GCACAUCUGCUUCUGG AAA **CAACAUACG** CCAGAAGCGGAUGUGC AGCUCGGUGAGGUCG CCCAUGUCGACA **UACUUAGCG** CCAGUGAUGCGUCUCG AAA **CAAUGUCUC** CGAGACGCGUCACUGG AGUCGAUAUGGG **UU AA CACGAAGUCAAUAC** | 178 |

| Name (Note) | Sequence | length |
|---|---|---|
| **3D-e-L2-2** | CCUGAUCUUAGGGUG **AAAUAAA** CCGGACAGAUCCCGUG AAA **CAUCUUGAC** CACGGGAUUUGUCCGG CACCCUAGGAUCAGG GCACCGUUCGGG **AAUAAUA** CAGUCCGGCAGGACGG AAA **CCUGUACUA** CCGUCCUGUCGGACUG CCCGAGCGGUGC **UU AC CACGAAGUCAAUAC** | 174 |
| **3D-e-L2-4** | CCUCACUAGCUUCGA **CUACAAGGA** CCAUUACGCGACCGG AAA **CUUCACUGA** CCGGUCGUGUAAUGG ACGAAGCUGGUGAGG CCUCUGACCGCA **GUGUCAAAC** GCUCGAGUGCGACGG AAA **UGAUCGUAG** CCGUCGCGCUCGAGC AGCGGUUAGAGG **UU UA CACGAAGUCAAUAC** | 174 |
| **3D-e-L2-5** | GCCAGCUUUCAGGCA **UACGCUUAC** CCUGACGUUCGGUCG AAA **GCAUGUCUA** CGACCGAGCGUCAGG AGCCUGAAGGCUGGC CCUGCUGCAGCA **CGAUGAAGA** CCCGCUCGCUCGGUG AAA **UGAAUGAGC** CACCGAGUGAGCGGG AGCUGCGGCAGG **UU UAAU CACGAAGUCAAUAC** | 176 |
| **3D-e-L2-6** | GCAGUGGAUAUGGCA **UACCAGUUC** AGUAUAGUUGCGGUCG AAA **CAAUGAUGG** CGACCGCAGCUAUACU AGCCAUAUUCACUGC CCAGGUUACGCA **UCAGACAUC** AGGGUAAUAGUCACGG AAA **UUCAGAUCG** CCGUGACUGUUACCCU AGCGUAGCCUGG **UU UACU CACGAAGUCAAUAC** | 180 |
| **3D-e-L2-7** | GCCAGCUUGAGGCCA **UACCGUAAC** GUAGUGGGACAUCCCG AAA **UCUCGUAUC** CGGGAUGUUCCACUAC AGGCCUCAGGCUGGC GCUACUCAUGCA **UGACGAAAG** CCUGGCAGCUCACGUG AAA **UUGGUAUCC** CACGUGAGUUGCCAGG AGCAUGGGUAGC **UU AUAU CACGAAGUCAAUAC** | 180 |
| **3D-e-L2-8** | GCCAUCUUUGACAGC **AAAUAAA** GCUCCAUGAACGUCGG AAA **CAGCUAUCA** CCGACGUUUAUGGAGC GCUGUCAGAGAUGGC CCAUGGUAACGG **AAUAAUA** GUCUGUCUGAGCCCUG AAA **GUACUUACC** CAGGGCUCGGACAGAC CCGUUGCCAUGG **UU AAUU CACGAAGUCAAUAC** | 176 |
| **3D-f-L0-10** | GGGCAAGCUGUUCCA **UCAAUCAGC** ACCUGGCUAACUCGUG AAA **GAAGAUUGC** CACGAGUUGGCCAGGU AGGAACAGUUUGCCC GACCGGACCGCA **GGUAGACUA** CACGCCGAGUACUCGAUGGUACAGGAUGUACCGCUGUAUAC AAA **UCAUUCACG** GUAUACAGUGGUACAUUCUGUACCGUCGAGUAUUCGGCGUG AGCGGUUCGGUC **UU AU CACGAAGUCAAUAC** | 228 |
| **3D-f-W0-2** | GCAUGAUAGACGGGA **CAUGUAACC** UCCGUUGCUACAUUGGUCAUGCGUUGACAAGAGUACGACUG AAA **GAGGAUACA** CAGUCGUAUUCUUGUCGACGCAUGGCCAAUGUGGCAACGGA ACCCGUCUGUCAUGC GCCAGGUGCCCA **CUUCAAGAC** GGUGGCAUGUUAUCGG AAA **CUCAUAGGA** CCGAUAACGUGCCACU AGGGCAUCUGGC **UU AA CACGAAGUCAAUAC** | 228 |
| **3D-f-W0-4** | GCACAGUGGCUGACA **GAGACAUUG** GCGAACUGAGGUGACUCAGACCUUAAGUUUCUGCUCAGUG AAA **UGAACAGUG** CACUGAGCGGAAACUUGAGGUCUGGGUCACCUUAGUUCGU AGUCAGCCGCUGUGC CCGAGUAAUGCA **UGUAUCCUC** GGGCUCGGCCAGACUG AAA **GAAUGGCUA** CAGUCUGGUCGAGCCC AGCAUUGCUCGG **UU AC CACGAAGUCAAUAC** | 226 |
| **3D-f-W0-5** | GCCAAGGUGCGAGCA **UAGUACAGG** UCCGGAUGGUGACAAGUCAUGAGGAUGUUCAGUUGGACUG AAA **GAUCAUGUG** CAGUCCAAUUGAACAUUCUCAUGAUUUGUCACUAUCCGGA AGCUCGCAUCUUGGC CCAGGUGGCAGC **AAUAAUA** GCUCCCAGGGUCCGUG AAA **GAAUCUGCA** CACGGACCUUGGGAGC GCUGCUACCUGG **UU UA CACGAAGUCAAUAC** | 224 |
| **3D-f-W1-2** | GCAGUGGUCCUUGCA **GUACGAAUG** GGUCCGAGGAUCGGUG AAA **GAGUAAGUC** CACCGAUCUUCGGACC AGCAAGGAUCACUGC CCGUAUGUGUCA **UGUCCUUAC** CUACGCGGUGCGCUCG AAA **UGAUCCAAG** CGAGCGCAUCGCGUAG AGACACGUACGG **UU UAAU CACGAAGUCAAUAC** | 180 |
| **3D-f-W1-3** | GGAGUGGCUGCUUAA **CUUGACCUA** GCUGCCCAC **GUAA** GUGGGCAGC AUAAGCAGUCACUCC CCUCAGGGCGGA **CCACAUGUA** CGCGAAGUC **UUCG** GACUUCGCG ACCGCCUUGAGG **UU UACU CACGAAGUCAAUAC** | 136 |
| **3D-f-W1-4** | GCACCGUUCACCCUA **CUACGAUCA** GCAGACUUACGGACCG AAA **GAAUGCUCA** CGGUCCGUGAGUCUGC AAGGGUGAGCGGUGC CCUAGUCCCGCA **CAGAGUUAG** GCGCUAGGAACGCCUG AAA **GAUGCAAUG** CAGGCGUUUCUAGCGC AGCGGGGCUAGG **UU AUAU CACGAAGUCAAUAC** | 180 |
| **3D-f-W1-5** | CAUGUAGGCCCUGCA **GCUCAUUCA** UGAUCUGUCCUAGUGG AAA **UGUAUCGAG** CCACUAGGGCAGAUCA AGCAGGGCUUACAUG CCUGUGAGGCCA **GAUCUGUUC** GCCUCCGUGACGCGUG AAA **GUAACGAUG** CACGCGUCGCGGAGGC AGGCCUUACAGG **UU AAUU CACGAAGUCAAUAC** | 180 |

| Name (Note) | Sequence | length |
|---|---|---|
| **3D-g-L0-11** | GGGCUGUUCGAGUGC **AAAUAAA** CGAUCGCUGACCGCUG AAA **GAAGCUAAG** CAGCGGUCGGCGAUCG GCACUCGGACAGCCC GACCAGUGACCG **AAUAAUA** CCAUCGUGGACGGCUAUGUGAGCGGUAUUCAUUACCGGCUG AAA **GCUUGUUAC** CAGCCGGUGAUGAAUAUCGCUCACGUAGCCGUUCACGAUGG CGGUCGCUGGUC **UU AU CACGAAGUCAAUAC** | 224 |
| **3D-g-W2-2** | CAGCUAUCGACUCGA **CCUAUGUUC** GCGUGAUGUGGACCCG AAA **CAAGUCAUC** CGGGUCCAUAUCACGC ACGAGUCGGUAGCUG GCAGAUCUGGAA **UCUCGUUAG** CCGACUAUAGGGUGCG AAA **UAGUCUACC** CGCACCCUGUAGUCGG AUCCAGGUCUGC **UU AA CACGAAGUCAAUAC** | 178 |
| **3D-g-W2-3** | GCAUUGUAGAACCGA **CGAUCUGAA** GGCGCUGUGGCUCGAC AAA **UAGAUUCCG** GUCGAGCCGCAGCGCC ACGGUUCUGCAAUGC CCGUAGGAGGCA **CUAACUAGC** GCUGGGAGUUCCACCG AAA **GAAGUAUGC** CGGUGGAAUUCCCAGU AGCCUCUUACGG **UU AC CACGAAGUCAAUAC** | 178 |
| **3D-g-W2-4** | GCAUGAUAGACGCCA **GGAUACCAA** GCGAGUCGAGUGACGG AAA **UCCUUGUAG** CCGUCACUUGACUCGC AGGCGUCUGUCAUGC GCAUCUCAUCCA **CUUCUGACA** GCGUGGCGACUAACGG AAA **UACAGUCAC** CCGUUAGUUGCCACGU AGGAUGGGAUGC **UU UA CACGAAGUCAAUAC** | 178 |
| **3D-g-W2-5** | CACCUAGUACGAGCA **GGUAAGUAC** GUCGCUUGUGACGGGC AAA **GUAAGCGUA** GCCCGUCAUAAGCGAC AGCUCGUAUUAGGUG CCGGUUCAGGGA **UCAUGCUAG** CCCGUCAUGGCACGUG AAA **GGCAAUGAA** CACGUGCCGUGACGGG ACCCUGGACCGG **UU UAAU CACGAAGUCAAUAC** | 180 |
| **3D-g-W3-1** | GCACUCGUUACGCCA **UCAUGACUC** GGCAGACGACUGCGUG AAA **UACAUAGCC** CACGCAGUUGUCUGCC AGGCGUAAUGAGUGC CCAGGUCCUGCA **GCAAUCUUC** GCUGUGAUUCGUUCGG AAA **CUGUAGUGA** CCGAACGAGUCACAGC AGCAGGGCCUGG **UU UACU CACGAAGUCAAUAC** | 180 |
| **3D-g-W3-3** | CCAUGUUCGUCCGGA **GAUAGUUCC** GUGGCAUCU **GUAA** AGAUGCCAC ACCGGACGGACAUGG GCUGCGGACUCA **UGUUCCUAG** GGUCGCGGA **UUCG** UCCGCGACC AGAGUCUGCAGC **UU AUAU CACGAAGUCAAUAC** | 136 |
| **3D-g-W3-4** | CCAUGCGUCCGACCA **GGAAUGUAC** GUCGGACUUCACGGUG AAA **GAACUGGUA** CACCGUGAGGUCCGAC AGGUCGGAUGCAUGG GCAAUGGUACGA **UCCUUAGAC** GAGGUCUUAUCCACGG AAA **CUAGUGUCA** CCGUGGAUGAGACCUC ACGUACUAUUGC **UU AAUU CACGAAGUCAAUAC** | 180 |
| **3D-g-W3-5** | CCAGCAGUGUCCGGA **CUUUGAACG** CGAGGUCUUUGCGGUG AAA **GUUACGGUA** CACCGCAAGGACCUCG ACCGGACAUUGCUGG GGUAUGCUGGCA **UGAGACUUC** CACUGGCUGAAAUCUG AAA **CUACUGCUA** CAGAUUUCGGCCAGUG AGCCAGUAUACC **UU ACUU CACGAAGUCAAUAC** | 180 |
| **3D-xy-W0-3** | GGGCUCGCGCAGGGA **CACUUCAGA** GCCGUUAUC **GUAA** GAUAACGGC ACCCUGCGUGAGCUC GCCCAUGGAGGA **UGUCGAUUG** CCGUGAUUGCUCGGUG AAA **UCAACUGUC** CACCGAGCGAUCACGG ACCUCCGUGGGC **UU AU CACGAAGUCAAUAC** | 156 |
| **3D-xy-W0-7** | GCAGGCUG **UUCG** CAGCCUGC CCAGGCGCA **CACUGUUCA** AGCCCGUGGCGAAGUG AAA **CAUCUAUGC** CACUUCGCUACGGGCU AGCGCCUGG **UU AA CACGAAGUCAAUAC** | 109 |
| **3D-xy-W0-8** | GCCACCUG **UUCG** CAGGUGGC CCAGCCGCA **CACAUGAUC** GCCAGGCUGAUACCGG AAA **CGCUAAGUA** CCGGUAUCGGCCUGGC AGCGGCUGG **UU AC CACGAAGUCAAUAC** | 109 |
| **3D-xy-W1-6** | GCUCCUGG **UUCG** CCAGGAGC GCUCCCGCA **CUACAUCUG** GGGACUCGGGUAUCUC AAA **CUUGUUCCA** GAGAUACCUGAGUCCC AGCGGGAGC **UU UA CACGAAGUCAAUAC** | 109 |
| **3D-xy-W1-7** | CGGAGCUG **UUCG** CAGCUCCG CCAGCCGGA **CGUAUGUUG** CCCGUGAGCUUAUCCG AAA **GUUUGACAC** CGGAUAAGUUCACGGG ACCGGCUGG **UU UAAU CACGAAGUCAAUAC** | 111 |
| **3D-xy-W1-8** | GCAGACGG **UUCG** CCGUCUGC GCUCCCGGA **GUCAAGAUG** GCUGACAUGACGAGGC AAA **UCUUCAUCG** GCCUCGUCGUGUCAGC ACCGGGAGC **UU UACU CACGAAGUCAAUAC** | 111 |
| **3D-xy-W2-7** | CCAGCCUG **UUCG** CAGGCUGG CGCUCGGGA **UCAGUGAAG** GCAGCCUGACGACGUG AAA **GAUGUCUGA** CACGUCGUUAGGCUGC ACCCGAGCG **UU AUAU CACGAAGUCAAUAC** | 111 |
| **3D-xy-W2-8** | CCAGCUGG **UUCG** CCAGCUGG GCCUCCGGA **UAGACAUGC** CACGAGGUACUUGGGC AAA **CUUUCGUCA** GCCCAAGUGCCUCGUG ACCGGAGGC **UU AAUU CACGAAGUCAAUAC** | 111 |
| **3D-xy-W3-6** | CAUCCCUG **UUCG** CAGGGAUG CCAGCCGCA **CCAUCAUUG** CCUAGGCUACAUCCCG AAA **GAAUCGUCA** CGGGAUGUGGCCUAGG AGCGGCUGG **UU ACUU CACGAAGUCAAUAC** | 111 |
| **3D-xy-W3-7** | GCCUCGUG **UUCG** CACGAGGC GCUGCCGGA **GAUACGAGA** CAGCCUGGCUCAGGGC AAA **GUUAGUCGA** GCCCUGAGUCAGGCUG ACCGGCAGC **UU UAUA CACGAAGUCAAUAC** | 111 |

| Name (Note) | Sequence | length |
| --- | --- | --- |
| **3D-xy-W3-8** | GCCUGAGG **UUCG** CCUCAGGC GGUCCGGGA **UGAUAGCUG** GGGGUGCGAGUACGUG AAA **GAUUGACUG** CACGUACUUGCACCCC ACCCGGACC **UU AUUA CACGAAGUCAAUAC** | 111 |
| **3D-xz-L0-13** | GGGCUAGG **UUCG** CCUAGCCC GCCCUGGGA **UCACUACAG** CCCGAGCGCCAGCAUUCUCACAUGCGAUCGUUCCGGACGG AAA **UUCUCAAGG** CCGUCCGGGACGAUCGUAUGUGAGGAUGCUGGUGCUCGGG ACCCAGGGC **UU AU CACGAAGUCAAUAC** | 157 |
| **3D-xz-L0-14** | CAACCCGG **UUCG** CCGGGUUG CGAGCGGCA **GACUACCUA** CCCGUGGUCCUAGUGGACAGGCCGACGACUCGUCGCGGUG AAA **GCUGAUUGA** CACCGCGAUGAGUCGUUGGCCUGUUCACUAGGGCCACGGG AGCCGCUCG **UU AA CACGAAGUCAAUAC** | 157 |
| **3D-xz-L1-9** | GCCUGGUG **UUCG** CACCAGGC GCUCCCGGA **UGACACUAG** CCCUGCUUGACAGGUG AAA **GUUAGAACG** CACCUGUCGAGCAGGG ACCGGGAGC **UU AC CACGAAGUCAAUAC** | 109 |
| **3D-xz-L1-10** | GCUCCGUG **UUCG** CACGGAGC CGAGCCCGA **UAGCAGUAG** UCACCCUGCGGCUGCG AAA **GAGUCAUGA** CGCAGCCGUAGGGUGA ACGGGCUCG **UU UA CACGAAGUCAAUAC** | 109 |
| **3D-xz-L2-3** | GCACAGGCGAGCACA **CGGAAUCUA** GCUAACGAC **GUAA** GUCGUUAGC AGUGCUCGUCUGUGC GCUGCUGAUGCA **UGGAACAAG** CAUCUUUGUCCUACG AAA **UAGGUCAAG** CGUAGGAUAAAGAUG AGCAUCGGCAGC **UU UAAU CACGAAGUCAAUAC** | 156 |
| **3D-xz-L2-9** | CCAGCUGG **UUCG** CCAGCUGG CCAGCCCGA **UGACGAUUC** CUGGAGCGGGAGCGC AAA **GGAACUAUC** GCGCUCCUGCUCCAG ACGGGCUGG **UU UACU CACGAAGUCAAUAC** | 109 |
| **3D-xz-L2-10** | CCUCGCUG **UUCG** CAGCGAGG GGAGCGCGA **UCGACUAAC** ACGGAUCGCGAGGUC AAA **GUACAUUCC** GACCUCGUGAUCCGU ACGCGCUCC **UU AUAU CACGAAGUCAAUAC** | 109 |
| **3D-xz-L2-11** | CCAGCUGG **UUCG** CCAGCUGG CCUACCCCA **CAGUCAAUC** GGCGUGGGACGAAGC AAA **CGUUCAAAG** GCUUCGUUCCACGCC AGGGGUAGG **UU AAUU CACGAAGUCAAUAC** | 109 |
| **3D-xz-W3-0** | CCGUCAGUAGCAGCA **CGUUCUAAC** CCGGUCUUAAUGGGCG AAA **GAGGUAACA** CGCCCAUUGAGACCGG AGCUGCUAUUGACGG GCAGCGCAGGCA **CGCUAUGAA** GGUCCGACG **UUCG** CGUCGGACC AGCCUGUGCUGC **UU ACUU CACGAAGUCAAUAC** | 158 |
| **3D-xz-W3-2** | CCACCGUAGUGCGUG **AAAUAAA** GCGUUCGGAUCUCCGC AAA **CUUUCGGAA** GCGGAGAUUCGAACGC CACGCACUGCGGUGG CCUACGCGCACA **CUUAGCUUC** GACCGCAGGGCUAGCG AAA **UAGGUAGUC** CGCUAGCCUUGCGGUC AGUGCGUGUAGG **UU UAUA CACGAAGUCAAUAC** | 178 |
| **3D-h-L0-12** | GGGCCUUCGAGCGCA **UACGUCUUC** GGUAGUAUGCUCUCG AAA **GAAUCAAGG** CGAGAGCGUACUACC AGCGCUCGGAGGCCC GCUCCGGUACCA **UAGUUGAGC** CUGGUGCUG **UUCG** CAGCACCAG AGGUACUGGAGC **UU AU CACGAAGUCAAUAC** | 154 |
| **3D-h-L0-13** | CCUCAUGCGCUGGGA **UACAAGUGC** CUCGGACUGGUCGUG AAA **GCGUUACUA** CACGACCGGUCCGAG ACCCAGCGUAUGAGG GCAGGUGCUGGA **UCACUACAG** GCCUUGUGAUCGCACUAGUGGCCGUCGUCUAUCGCGACGG AAA **UUCUCAAGG** CCGUCGCGGUAGACGAUGGCCACUGGUGCGAUUACAAGGC ACCAGCGCCUGC **UU AA CACGAAGUCAAUAC** | 224 |
| **3D-h-L0-14** | CCAGCGUUUGCGACG **AAAUAAA** CGAAUGCGGAGACGG AAA **CAUUGGAAC** CCGUCUCUGCAUUCG CGUCGCAGACGCUGG CCUCCGAGCGCA **GACUACCUA** CCGGUGCUUGUAAUGGCGUUGAUGCCGUGUCGGCGAGGUG AAA **GCUGAUUGA** CACCUCGCUGACACGGUAUCAACGUCAUUACAGGCACCGG AGCGCUUGGAGG **UU AC CACGAAGUCAAUAC** | 222 |
| **3D-h-L0-15** | CCAGGAGG **UUCG** CCUCCUGG GCACCGCGA **GUUGCUAGA** ACCCGGUCUGGUCGUUUACCUAUAGCAUACUAUCAGCGGUG AAA **GAAGACGUA** CACCGCUGGUAGUAUGUUAUAGGUGAACGACCGGACCGGGU ACGCGGUGC **UU UA CACGAAGUCAAUAC** | 159 |
| **3D-h-L0-16** | GCACCGUG **UUCG** CACGGUGC GCUGCGGUA **UCAUCUCUG** CCCAGAGGUUCGGAACGAGCGGUCUGAACGAUGAUGCGGUG AAA **GCACUUGUA** CACCGCAUUAUCGUUCGGACCGCUUGUUCCGAGCCUCUGGG AACCGCAGC **UU UAAU CACGAAGUCAAUAC** | 161 |
| **3D-h-L1-9** | GCAGCGUACAUUCCA **GAGAUUCGA** GCGGCAGGGAUGCGUG AAA **UCAAGAUCC** CACGCAUCUCUGCCGC AGGAAUGUGCGCUGC GCUGCUAGCGGA **UGACACUAG** CCGGAGAUGAUAUGUG AAA **GUUAGAACG** CACAUAUCGUCUCCGG ACCGCUGGCAGC **UU UACU CACGAAGUCAAUAC** | 180 |

| Name (Note) | Sequence | length |
|---|---|---|
| **3D-h-L1-10** | GCAGCCGUGGCAGCA **CUGAUCAUC** GCAUCUUGGCCGAGGC AAA **GGACAUUCA** GCCUCGGCUAAGAUGC AGCUGCCAUGGCUGC GCGUGGGACGGA **UAGCAGUAG** CCUGACCGUGGAUGCG AAA **GAGUCAUGA** CGCAUCCAUGGUCAGG ACCGUCUCACGC **UU** **AUAU** **CACGAAGUCAAUAC** | 180 |
| **3D-h-L1-11** | CCUGGCAUAGCGCUG **AAAUAAA** GCCAGAGUCUCACCGG AAA **CUUACAGAC** CCGGUGAGGCUCUGGC CAGCGCUGUGCCAGG CCAUCUGGAGGC **AAUAAUA** GCUCCGAUCCUCGCGG AAA **CUCAAUGGA** CCGCGAGGGUCGGAGC GCCUCUAGAUGG **UU** **AAUU** **CACGAAGUCAAUAC** | 176 |
| **3D-h-L1-12** | GGCACCUG **UUCG** CAGGUGCC GCUCCGGGA **CUAAGACUG** CCGGACGUGACGGUG AAA **CUAGUCCUA** CACCGUCGCGUCCGG ACCCGGAGC **UU** **ACUU** **CACGAAGUCAAUAC** | 109 |
| **3D-h-L1-13** | GCCUCGAG **UUCG** CUCGAGGC GGUCCCGCA **GCAUCUAUC** UCGGACGUCCUGCUG AAA **GAUAGUAGC** CAGCAGGGCGUCCGA AGCGGGACC **UU** **UAUA** **CACGAAGUCAAUAC** | 109 |
| **3D-h-L1-14** | GCCUCGUG **UUCG** CACGAGGC GGUCCCGCA **UGACUCUAC** CCGUCCUGCCGAGGC AAA **GGAUGAAUC** GCCUCGGUAGGACGG AGCGGGACC **UU** **AUUA** **CACGAAGUCAAUAC** | 109 |
| **3D-i-L2-3** | GGGCCAGUACAGGCA **CGGAAUCUA** GCGCCUCGAGCCGUG AAA **CUAAGGUAC** CACGGCUUGAGGCGC AGCCUGUAUUGGCUC GAUCGUAGCGCA **UGGAACAAG** GCACGCAGAUCCUCG AAA **UAGGUCAAG** CGAGGAUUUGCGUGU AGCGCUGCGAUC **UU** **AU** **CACGAAGUCAAUAC** | 174 |
| **3D-i-L2-9** | CAGUAGUUGCCUGCA **UUGGACUUG** CGAUCCGGCACCUGG AAA **CGUAACUAG** CCAGGUGUCGGAUCG AGCAGGCAGCUACUG GCGCUGGCAGCA **UGACGAUUC** CACGGUCGCCGAGCG AAA **GGAACUAUC** CGCUCGGUGACCGUG AGCUGCUAGCGC **UU** **AA** **CACGAAGUCAAUAC** | 174 |
| **3D-i-L2-10** | CCUAUUGGGAUAGCA **CUGUGAAAC** GCGGAAUGAGACCGC AAA **CAUUCUGAG** GCGGUCUUAUUCCGC AGCUAUCCUAAUAGG GCGUCUUCGGCA **UCGACUAAC** CCGCUGCGCAGGAGC AAA **GUACAUUCC** GCUCCUGUGCAGCGG AGCCGAGGACGC **UU** **AC** **CACGAAGUCAAUAC** | 174 |
| **3D-i-L2-11** | CCUCCAGUAGAGCCA **GCAAGUAAG** GUGCGAGUAUGCCUG AAA **CUAUCUCGA** CAGGCAUGCUCGCAC AGGCUCUAUUGGAGG CCAUAUGCUGCA **CAGUCAAUC** ACCGACCGUCGGUGC AAA **CGUUCAAAG** GCACCGAUGGUCGGU AGCAGCGUAUGG **UU** **UA** **CACGAAGUCAAUAC** | 174 |
| **3D-i-L2-12** | GCAGACGG **UUCG** CCGUCUGC CCUGGGCCA **UGAGAAUGC** GGGUCCGGCAGCACCG AAA **UACAACUGG** CGGUGCUGUCGGACCC AGGCCCAGG **UU** **UAAU** **CACGAAGUCAAUAC** | 111 |
| **3D-i-L2-13** | CCUCGCUG **UUCG** CAGCGAGG GCUCGCCCA **UCAUCAAGC** ACCCGCUGCACAUGCG AAA **CAAGGUAUG** CGCAUGUGUAGCGGGU AGGGCGAGC **UU** **UACU** **CACGAAGUCAAUAC** | 111 |
| **3D-i-L3-9** | CCAGCCGGACGCCGA **CUUGUGAUG** GCGGAAUUGGCUCGCG AAA **GUAACUCUC** CGCGAGCCGAUUCCGC ACGGCGUCUGGCUGG GCUCCGCUCGGA **CUAUGCCUA** CCCAGAGUUGGUAGCG AAA **CUGUCUACA** CGCUACCAGCUCUGGG ACCGAGUGGAGC **UU** **AUAU** **CACGAAGUCAAUAC** | 180 |
| **3D-i-L3-10** | GCAGUUUAUCGACCA **UUCUGUCAC** GGCACUAUUCAUGUAG AAA **UCUAUCCAG** CUACAUGAGUAGUGCC AGGUCGAUGAACUGC GGUUGUGGCACA **UCCUAAGUC** UCGGUCGGCUCUAGGC AAA **UGUUCACUC** GCCUAGAGUCGACCGA AGUGCCGCAACC **UU** **AAUU** **CACGAAGUCAAUAC** | 180 |
| **3D-i-L3-11** | GGGUCGAGCAUGCUG **AAAUAAA** CCGUGUAUUCACGGUG AAA **GUACUAGGA** CACCGUGAGUACACGG CAGCAUGUUCGACCC GCUCUGUAGUGG **AAUAAUA** GCUACCUGGGACCUCC AAA **GAUUCAGAC** GGAGGUCCUAGGUAGC CCACUGCAGAGC **UU** **ACUU** **CACGAAGUCAAUAC** | 176 |
| **3D-i-L3-12** | GGUCCGGA **UUCG** UCCGGACC CCAGGCCCA **GUAUGACGA** GCCGUGCGCGAGGUG AAA **UUGCCUAUC** CACCUCGUGCACGGC AGGGCCUGG **UU** **UAUA** **CACGAAGUCAAUAC** | 109 |
| **3D-i-L3-13** | CGAGCGUG **UUCG** CACGCUCG CCAGGCCCA **CUAAGCAGA** GAGAUCUUACUGCCC AAA **GCAUGAAAG** GGGCAGUGAGAUCUC AGGGCCUGG **UU** **AUUA** **CACGAAGUCAAUAC** | 109 |
| **3D-i-L3-14** | CCUCGCUG **UUCG** CAGCGAGG GCCUCCCGA **GUAGCUAAC** CGGGUCGUCCAGCCG AAA **CAACUUGAG** CGGCUGGGCGACCCG ACGGGAGGC **UU** **CUUA** **CACGAAGUCAAUAC** | 109 |
| **3D-j-L3-0** | GGGCCUUAGCACAGA **CUCUUGCAA** GCCAGUCCG **GUAA** CGGACUGGC ACUGUGCUGAGGCCC CGUACGAGGCCA **UGAUGUGAC** GCCUGCAGCUCGGUG AAA **GAUACUGUG** CACCGAGUUGCAGGC AGGCCUUGUACG **UU** **AU** **CACGAAGUCAAUAC** | 154 |

| Name (Note) | Sequence | length |
|---|---|---|
| **3D-j-L3-1** | GCAGCAGGUGCCGCA **GGUACAAAG** GCAGCAAUGGGUCGG AAA **UCCUAACUG** CCGACCCGUUGCUGC AGCGGCACUUGCUGC GCUGAGUCCGGA **CGUACAAUC** UCGACUCGGGACCGG AAA **UGAGUUGAC** CCGGUCCUGAGUCGA ACCGGAUUCAGC **UU AA CACGAAGUCAAUAC** | 174 |
| **3D-j-L3-2** | GCAGUUGCAUAGGCA **CGUUAGAUC** GGAGUAAGCAGGGUG AAA **GACUAAGUG** CACCCUGUUUACUCC AGCCUAUGUAACUGC CCAGACUCUGAG **AAUAAUA** CCAGAGGUGGACUGG AAA **CUCUAGAUG** CCAGUCCGCCUCUGG CUCAGGGUCUGG **UU AC CACGAAGUCAAUAC** | 172 |
| **3D-j-L3-3** | CCAGCGUCUUACCGA **CUAGAGAUC** GCAGCGAUGAGCCGUG AAA **GUUCAGGAA** CACGGCUCGUCGCUGC ACGGUAAGGCGCUGG GGCUGGAUCUCA **UACUAUCGC** GGAGCCGUAAUGUCAG AAA **UCUUACACG** CUGACAUUGCGGCUCC AGAGAUUCAGCC **UU UA CACGAAGUCAAUAC** | 178 |
| **3D-j-L3-4** | GCACCGGUUGCGGCA **GCAGUUCUA** GGAUGUAGUCGGACGG AAA **UAGCUCUAG** CCGUCCGAUUACAUCC AGCCGCAAUCGGUGC CCUACGAUCGCA **GACUCAUUG** ACCUCCGUCACAUGGC AAA **GAUUCUCAG** GCCAUGUGGCGGAGGU AGCGAUUGUAGG **UU UAAU CACGAAGUCAAUAC** | 180 |
| **3D-j-L3-5** | CCAGCAGGAACGUCG **AAAUAAA** GCUGGCUGCCAGGUGG AAA **CUAUGCAAG** CCACCUGGUAGCCAGC CGACGUUUCUGCUGG GCUUACUAGUGG **AAUAAUA** CCUGCCGUACCGCGUG AAA **UUCCAUGUC** CACGCGGUGCGGCAGG CCACUGGUAAGC **UU UACU CACGAAGUCAAUAC** | 176 |
| **3D-j-L3-6** | GCCUCCGGUUCCCGA **GUAUGCAGA** GGCCUUGGC **GUAA** GCCAAGGCC ACGGGAACUGGAGGC GCUUAUGGCGCA **GUAUCCAUG** GCAGGCCUUAGCGGC AAA **GGAUUCACA** GCCGCUAGGGCCUGC AGCGCCGUAAGC **UU AUAU CACGAAGUCAAUAC** | 156 |
| **3D-j-L3-7** | GUGCGUGCACGACCA **UAGUUCGUC** GGACCAUGGGUCGUG AAA **UUCUGAUCC** CACGACCUAUGGUCC AGGUCGUGUACGCAC CGUACGAGUGCA **UAGCCAAUG** GUCGAUCGCCCUCGG AAA **CAGUAGAGA** CCGAGGGUGAUCGAC AGCACUUGUACG **UU AAUU CACGAAGUCAAUAC** | 176 |
| **3D-j-L3-8** | GCCUACUAGCGCGAA **GACCAAUCA** GGAGCCAGUACGCAG AAA **UACGGAAAC** CUGCGUAUUGGCUCC AUCGCGCUGGUAGGC CCUAGUCUAGCA **UCACUCUUG** CCAGGCGGACCGGUG AAA **GGUAUCGUA** CACCGGUUCGCCUGG AGCUAGGCUAGG **UU ACUU CACGAAGUCAAUAC** | 176 |
| **3D-k-W3-0** | GGGCAUGUGAGCGCA **CGUUCUAAC** GCAGCGGUAUCGAGCG AAA **GAGGUAACA** CGCUCGAUGCCGCUGC AGCGCUCAUAUGCUC GCACCUUGCGCA **CGCUAUGAA** GCCCUGCUUGAUACCG AAA **GCUCAACUA** CGGUAUCAGGCAGGGC AGCGCAGGGUGC **UU AU CACGAAGUCAAUAC** | 178 |
| **3D-k-W3-2** | CCAGGGUUAGAUGCA **UCCAUUGAG** CGGCUGAGCUUGCGGC AAA **CUUUCGGAA** GCCGCAAGUUCAGCCG AGCAUCUAGCCCUGG CGCCGGCAUGGA **CUUAGCUUC** AGGGCCAGCGCUCGCG AAA **UAGGUAGUC** CGCGAGCGUUGGCCCU ACCAUGUCGGCG **UU AA CACGAAGUCAAUAC** | 178 |
| **3D-k-W3-6** | CCAGAGGCUACUGGA **UGUAGACAG** CCAUGCGGAGCCUCGG AAA **UCUGCAUAC** CCGAGGCUUCGCAUGG ACCAGUAGUCUCUGG CCUCCGACGGCA **CCAUCAUUG** CCCGUGAUUACACGCG AAA **GAAUCGUCA** CGCGUGUAGUCACGGG AGCCGUUGGAGG **UU AC CACGAAGUCAAUAC** | 178 |
| **3D-k-W3-7** | GCCAAUGCUGUCCCA **GAGUGAACA** GGAUCCCGCUACGGUG AAA **GACGAACUA** CACCGUAGUGGGAUCC AGGGACAGUAUUGGC GCUCAGGACCCA **GAUACGAGA** CAGUCUGGCUUAGGCC AAA **GUUAGUCGA** GGCCUAAGUCAGACUG AGGGUCUUGAGC **UU UA CACGAAGUCAAUAC** | 178 |
| **3D-k-W3-8** | GAGUGCGGUAGCCCA **GUCUGAAUC** UAAUGAGUAACGCGUG AAA **UGAUUGGUC** CACGCGUUGCUCAUUA AGGGCUACUGCACUC GCGGUGCUCGGA **UGAUAGCUG** CCGGAGUGGCGAGGUG AAA **GAUUGACUG** CACCUCGCUACUCCGG ACCGAGUACCGC **UU UAAU CACGAAGUCAAUAC** | 180 |
| **3D-k-W3-9** | GCCUCAGG **UUCG** CCUGAGGC CGUCGCGCA **GGAUCAGAA** GUGGCGUUUCAGGGUG AAA **UAGGCAUAG** CACCCUGAGACGCCAC AGCGCGACG **UU UACU CACGAAGUCAAUAC** | 111 |
| **3D-k-W3-10** | GCACCUGG **UUCG** CCAGGUGC CGGUCCAGCA **GUUUCCGUA** GCAGGGAUUCAGUCCG AAA **GACUUAGGA** CGGACUGAGUCCCUGC AGCUGGACCG **UU AUAU CACGAAGUCAAUAC** | 113 |
| **3D-k-W4-9** | GCUCAGGG **UUCG** CCCUGAGC GGUCCGGCA **GAGAGUUAC** CGUGAGCUCUAUCCGG AAA **UCGUCAUAC** CCGGAUAGGGCUCACG AGCCGGACC **UU AAUU CACGAAGUCAAUAC** | 111 |
| **3D-k-W4-10** | CGAGUCGG **UUCG** CCGACUCG GCCUCGGGA **CUGGAUAGA** CAGUGGAGCGUAGGUG AAA **UCUGCUUAG** CACCUACGUUCCACUG ACCCGAGGC **UU ACUU CACGAAGUCAAUAC** | 111 |

| Name (Note) | Sequence | length |
|---|---|---|
| **3D-k-W4-11** | GGCUCAGG **UUCG** CCUGAGCC GGUCGCGCA **UCCUAGUAC** GUGCCGUGCCGACCUG AAA **GUUAGCUAC** CAGGUCGGUACGGCAC AGCGCGACC **UU UAUA CACGAAGUCAAUAC** | 111 |
| **3D-l-W4-0** | GGGCAAGUGGUCGCA **UAGGACUAG** GCGCUAUAG **GUAA** CUAUAGCGC AGCGACCAUUUGCCC GCUCGUCUGCCA **CCUUGAUUC** GGCAGCUGU **UUCG** ACAGCUGCC AGGCAGGCGAGC **UU AU CACGAAGUCAAUAC** | 134 |
| **3D-l-W4-1** | GCCUCCGCUGGUCCA **GCUACUAUC** GGAGCGAGAUAGGUCG AAA **UCGAAUCUC** CGACCUAUUUCGCUCC AGGACCAGUGGAGGC GCUGCUGUCGGA **UAGUAACGC** CACCGGUUUAGGCACG AAA **UCUAGCAAC** CGUGCCUAGACCGGUG ACCGACGGCAGC **UU AA CACGAAGUCAAUAC** | 178 |
| **3D-l-W4-2** | CCUACAGUCUGACCA **GAUUCAUCC** ACGGAGUGCGUCGGUG AAA **GAUGAUCAG** CACCGACGUACUCCGU AGGUCAGAUUGUAGG GCUACUGCACCA **GUUCCAAUG** UGACCGUGCGGUUCGG AAA **CAGAGAUGA** CCGAACCGUACGGUCA AGGUGCGGUAGC **UU AC CACGAAGUCAAUAC** | 178 |
| **3D-l-W4-3** | GCACCGUUGGAUGCA **CCAGUUGUA** GGCACCGGUCCCGCAG AAA **CAAGUCCAA** CUGCGGGAUCGGUGCC AGCAUCCAGCGGUGC CCCAUGCGUGCA **GGAUCUUGA** CCGCCUAUUGCCGUGG AAA **CAGUCUUAG** CCACGGCAGUAGGCGG AGCACGUAUGGG **UU UA CACGAAGUCAAUAC** | 178 |
| **3D-l-W4-4** | CAGCAUUGCCUGGCA **CAUACCUUG** GCAACAGGCAGGUCGC AAA **GUUUCACAG** GCGACCUGUCUGUUGC AGCCAGGCGAUGCUG GCUAGUACCCUA **UGAAUGUCC** GCUGCGUUAUGGACUG AAA **GAUAGAUGC** CAGUCCAUGACGCAGU AAGGGUGCUAGC **UU UAAU CACGAAGUCAAUAC** | 180 |
| **3D-l-W4-5** | GCAACCUUUGACCUG **AAAUAAA** GCAUAGUGCGGACGUG AAA **CUUACUUGC** CACGUCCGUACUAUGC CAGGUCAGAGGUUGC CCAGGUGACGCA **GUCUGUAAG** CCACGUCUCGUGACCG AAA **GUAGAGUCA** CGGUCACGGGACGUGG AGCGUCGCCUGG **UU UACU CACGAAGUCAAUAC** | 178 |
| **3D-l-W4-6** | GGCAUAGUCCGCUCA **GAUAGGCAA** GCGCUCUCG **GUAA** CGAGAGCGC AGAGCGGAUUAUGCC GCUCCUGUGGCA **CUAGUUACG** CCGUUGAAU **UUCG** AUUCAACGG AGCCACGGGAGC **UU AUAU CACGAAGUCAAUAC** | 136 |
| **3D-l-W4-7** | CACCUCGUAGUUCCA **CUUUCAUGC** AUCUACAGGGACAGCG AAA **CAUCACAAG** CGCUGUCCUUGUAGAU AGGAACUAUGAGGUG GCUACUUACGGA **CUCAGAAUG** CCCUGGUUACGUCGGC AAA **GCAUUCUCA** GCCGACGUGACCAGGG ACCGUAGGUAGC **UU AAUU CACGAAGUCAAUAC** | 180 |
| **3D-l-W4-8** | GCACCUUAUCGAGGA **CUCAAGUUG** GCCCUCGGGUCCCGUG AAA **GUGACAGAA** CACGGGACUCGAGGGC ACCUCGAUGAGGUGC GCACGUCCUGCA **UCGAGAUAG** GCGGACGGCAUGCGUG AAA **GCUUGAUGA** CACGCAUGUCGUCCGC AGCAGGGCGUGC **UU ACUU CACGAAGUCAAUAC** | 180 |
| **3D-m-W0-3** | GGGCUCGGAGCGGUA **CACUUCAGA** CCGUAGUGUACGCAUGCUCCGCAGGACGUUCUGCAGACCG AAA **GUUGAUGUC** CGGUCUGCGGAACGUCUUGCGGAGUAUGCGUAUACUACGG AGCCGCUCUGAGUCC GCACCUGUUCCA **UGUCGAUUG** GCCUGCAUCGUCGUCG AAA **UCAACUGUC** CGACGACGGUGCAGGU AGGAACGGGUGC **UU AU CACGAAGUCAAUAC** | 226 |
| **3D-m-W0-6** | GCCAUCGGAUGCGGA **CACAGUAUC** AGAGCCCAC **GUAA** GUGGGCUCU ACCGCAUCUGAUGGC CCAGGUGCGGCA **GACAUCAAC** UUAGCAGCG **UUCG** CGCUGCUAA AGCCGCGCCUGG **UU AA CACGAAGUCAAUAC** | 134 |
| **3D-m-W0-7** | GCCUCGGAGCGUGCA **GUCAACUCA** GCGCAAUGGCCUAAUCUCAUCUAUGCUCCCGAUCGGAACUG AAA **UUGAAGACC** CAGUUCCGGUCGGGAGUAUAGAUGGGAUUAGGUCAUUGCGU AGCACGCUUCGAGGC CCAGGGUCCGCA **CACUGUUCA** CCUGCGGGAGCCAGUG AAA **CAUCUAUGC** CACUGGCUUCCGCAGG AGCGGAUCCUGG **UU AC CACGAAGUCAAUAC** | 228 |
| **3D-m-W0-8** | GCGUCUUGACCGGGA **CAUCUAGAG** CCGCAACUGUUGCUUGGUACGGUAUGCGAUGCGACUAUGUG AAA **CUGAUACUC** CACAUAGUUGCAUCGCGUACCGUAUCAAGCAAUAGUUGCGG ACCCGGUCGAGACGC CCUAGGGACCGA **CACAUGAUC** UCCGGCUUGUAAACGG AAA **CGCUAAGUA** CCGUUUACGAGCCGGA ACGGUCUCUAGG **UU UA CACGAAGUCAAUAC** | 228 |
| **3D-m-W0-9** | GCCACCUG **UUCG** CAGGUGGC CCUCGCGCA **GGUCUUCAA** GGUCGCAUGAGAUCCG AAA **GUCACAUCA** CGGAUCUCGUGCGACC AGCGCGAGG **UU UAAU CACGAAGUCAAUAC** | 111 |
| **3D-m-W0-10** | CCAGCGUG **UUCG** CACGCUGG CCCUCGCCA **GAGUAUCAG** CCGGAGCUCGUACCAG AAA **GAUUGUACG** CUGGUACGGGCUCCGG AGGCGAGGG **UU UACU CACGAAGUCAAUAC** | 111 |

| Name (Note) | Sequence | length |
|---|---|---|
| **3D-m-W1-6** | GCUACAGAGUGGGCA **CGUGUAAGA** GCAGCGAGAGUCGUGG AAA **UUGCAAGAG** CCACGACUUUCGCUGC AGCCCACUUUGUAGC GCAGCGCCAGCA **CUACAUCUG** CCGGACGGGAGUACGC AAA **CUUGUUCCA** GCGUACUCUCGUCCGG AGCUGGUGCUGC **UU AUAU CACGAAGUCAAUAC** | 180 |
| **3D-m-W1-7** | CAGAUGUCUGGACCA **CUGAGAAUC** GCUAGGAUUAUGGCUG AAA **CUUUGUACC** CAGCCAUAGUCCUAGC AGGUCCAGGCAUCUG CCGCUGCACUCA **CGUAUGUUG** ACCCUCAGGGCUUGCG AAA **GUUUGACAC** CGCAAGCCUUGAGGGU AGAGUGUAGCGG **UU AAUU CACGAAGUCAAUAC** | 180 |
| **3D-m-W1-8** | CGCAGUGCACCUCGA **GACAUGGAA** CUGGUAGGACGAGGUG AAA **GAUCUAACG** CACCUCGUUCUACCAG ACGAGGUGUACUGCG GCUCAGCGGCAA **GUCAAGAUG** UCCCAGGUGAAUGGAC AAA **UCUUCAUCG** GUCCAUUCGCCUGGGA AUGCCGUUGAGC **UU ACUU CACGAAGUCAAUAC** | 180 |
| **3D-n-W2-6** | GGGCAUGAUCGAGCA **UGUGAAUCC** GGCAACGCC **GUAA** GGCGUUGCC AGCUCGAUUAUGCCC CCAGGUCGAGCA **GUACCUUAG** CCGCUUACC **UUCG** GGUAAGCGG AGCUCGGCCUGG **UU AU CACGAAGUCAAUAC** | 134 |
| **3D-n-W2-7** | CCAGACGGGCUCGCA **UCUCUACUG** UGCUCCGGGCCUCGUG AAA **GAUCUCUAG** CACGAGGCUCGGAGCA AGCGAGCCUGUCUGG CCGUGUCGACCA **UCAGUGAAG** GCUCGAUGAUGUGCAG AAA **GAUGUCUGA** CUGCACAUUAUCGAGC AGGUCGGCACGG **UU AA CACGAAGUCAAUAC** | 178 |
| **3D-n-W2-8** | CCUAUUUGAACCGCA **UACGAUACC** CAUCAGGUGCACCGUG AAA **UAGAACUGC** CACGGUGCGCCUGAUG AGCGGUUCGAAUAGG GCUGCGAACGGA **UAGACAUGC** CUCUGGCUAUGCGGUC AAA **CUUUCGUCA** GACCGCAUGGCCAGAG ACCGUUUGCAGC **UU AC CACGAAGUCAAUAC** | 178 |
| **3D-n-W2-9** | GCCUCAGG **UUCG** CCUGAGGC GGUCCGCGA **UUCCUGAAC** GCGCUCCGGUAGCCAG AAA **CAUGGAUAC** CUGGCUACUGGAGCGC ACGCGGACC **UU UA CACGAAGUCAAUAC** | 109 |
| **3D-n-W2-10** | GCCCUCGG **UUCG** CCGAGGGC CGGCGGUGA **CUAGAGCUA** CCGACGUGCUCUCGCG AAA **CAUUGGCUA** CGCGAGAGUACGUCGG ACACCGCCG **UU UAAU CACGAAGUCAAUAC** | 111 |
| **3D-n-W2-11** | GGUCCGGA **UUCG** UCCGGACC GCCUCCGGA **CUUGCAUAG** CGUGGGUUCAAGCCGG AAA **CAAGAGUGA** CCGGCUUGGACCCACG ACCGGAGGC **UU UACU CACGAAGUCAAUAC** | 111 |
| **3D-n-W1-9** | GCCUCAGG **UUCG** CCUGAGGC CGUCGCCCA **CAGUUAGGA** GGGCAGCGGAUCCCUG AAA **GCGAUAGUA** CAGGGAUCUGCUGCCC AGGGCGACG **UU AUAU CACGAAGUCAAUAC** | 111 |
| **3D-n-W1-10** | GCCUCAGG **UUCG** CCUGAGGC GGUCCCGCA **CACUUAGUC** GCGACCAUGCGCUGUG AAA **CAAUGAGUC** CACAGCGCGUGGUCGU AGCGGGACC **UU AAUU CACGAAGUCAAUAC** | 111 |

#### 7) Plasmid for 3D assembly

**3D_23** (4 plasmids): 3D-a-Plasmid, 3D-b-Plasmid, 3D-za-Plasmid, 3D-yz-Plasmid.

**3D_71** (9 plasmids): 3D-a-Plasmid, 3D-b-Plasmid, 3D-c-Plasmid, 3D-d-Plasmid, 3D-e-Plasmid, 3D-f-Plasmid, 3D-g-Plasmid, 3D-xy-Plasmid, 3D-xz-Plasmid.

**3D_118** (14 plasmids): 3D-a-Plasmid, 3D-b-Plasmid, 3D-c-Plasmid, 3D-d-Plasmid, 3D-e-Plasmid, 3D-f-Plasmid, 3D-g-Plasmid, 3D-h-Plasmid, 3D-i-Plasmid, 3D-j-Plasmid, 3D-k-Plasmid, 3D-l-Plasmid, 3D-m-Plasmid, 3D-n-Plasmid.

| Name (Note) | Sequence | length |
|---|---|---|
| **3D-a-Plasmid**<br>Template for 3D-a tiles | **GTTCTAATACGACTCACTATA** GGGCAAGTCGCTGCA **TTCGACATG** TATCCAGGGACCCGG AAA<br>**CGAATGCAA** CCGGGTCTCTGGATA AGCAGCGATTTGCCC GCTCCTGAGACA **CTGCTAACC**<br>CGATCGTGC **TTCG** GCACGATCG AGTCTCGGGAGC **TT AT CACGAAGTCAATAC**<br>CCATATTGGAGCCCA **GTCTCATGA** GCCACCGTCACCGACG AAA **GAAGGTATC**<br>CGTCGGTGGCGGTGGC AGGGCTCCGATATGG GCTCCTGCTGCA **GTATCTCCA**<br>GCCCAGGCGTGTTCATGCTTGTACTAAGATCTTATCAGCGG AAA **CATGTCGAA**<br>CCGCTGATGAGATCTTGGTACAAGTATGAACATGCCTGGGT AGCAGCGGGAGC **TT AA**<br>**CACGAAGTCAATAC** CCTGTAGCCTCGCCA **CGTAAGTTC** GGATACTGA **GTAA** TCAGTATCC<br>AGGCGAGGTTACAGG GCTCGTCCTGCA **TTGCATTCG** GGTATGAGA **TTCG** TCTCATACC<br>AGCAGGGCGAGC **TT AC CACGAAGTCAATAC** CAGAGGTAACGGCCA **CTTTGCTCA**<br>CCGACAGCGCTCAGCTGTCGGATGTCCAGTAGGACGACCGG AAA **CAATCGACA**<br>CCGGTCGTTCTACTGGGCATCCGATAGCTGAGTGCTGTCGG AGGCCGTTGCCTCTG<br>GCCGTGCGAGCA **CAGAACTCA** CCTACGATCGTAAGCG AAA **TGGAGATAC**<br>CGCTTACGGTCGTAGG AGCTCGTACGGC **TT TA CACGAAGTCAATAC GTCCAACC TT**<br>**CATGCTTACGACG** | 788 |
| **3D-za-Plasmid**<br>Template for 3D-za tiles | **GTTCTAATACGACTCACTATA** GGGCAGATCCTTGCA **CCGATTCTA** ATGTACGGGATCCGC AAA<br>**TGAGTTCTG** GCGGATCTCGTACAT AGCAAGGGTCTGTCC GCATGGGCATGG **AATAATA**<br>CACTCAAGCTATGGGATGAGGCGTAGTTAGCAGATAGTCAGTCCTTGCTTTCGGACACGTAGGATCAG<br>CGAAGGTCCAATGTTACCGGTG AAA **GGTTAGCAG**<br>CACCGGTGACATTGGGCCTTCGTTGATCCTGCGTGTCTGAAAGCAGGGACTGGCTATCTGTTAACTAT<br>GCCTCATTCCATAGTTTGAGTG CCATGCTCATGC **TT AT CACGAAGTCAATAC**<br>GCCTACGTCAGGCTG **AAATAAA** GACTAACTCGACCGTG AAA **GTAACCTGA**<br>CACGGTCGGGTTAGTC CAGCCTGATGTAGGC CCAGGGCAGGGT **AATAATA**<br>CGACTTCATCAGAATGGGCCTAGGGCTTCGCAGGGACCTGG AAA **TAGAATCGG**<br>CCAGGTCCTTGCGAAGTCCTAGGCTCATTCTGGTGAAGTCG ACCCTGTCCTGG **TT AA**<br>**CACGAAGTCAATAC** CCAGTGGGACACGGA **GTGCATTGA** CCTGACAGC **GTAA** GCTGTCAGG<br>ACCGTGTCTCACTGG GAGTCGCCTGCA **GACAGTTGA** GGGCAATGCCGAGTG AAA **GAACTTACG**<br>CACTCGGTATTGCCC AGCAGGTGACTC **TT AC CACGAAGTCAATAC** CCAGACGGAAGCCCA<br>**TTCCACTTG** CATAGTGTCTACCGG AAA **CTAACTCTG** CCGGTAGGCACTATG<br>AGGGCTTCTGTCTGG GCAGAGTCATGC **AATAATA** CCAGATCTCGGACGG AAA **TGAGCAAAG**<br>CCGTCCGGGATCTGG GCATGATTCTGC **TT TA CACGAAGTCAATAC** GCAGCGTC **TTCG**<br>GACGCTGC CCAGGCGCA **TGTCGATTG** CCGGTGAGGTCAGGTG AAA **TCAACTGTC**<br>CACCTGACTTCACCGG AGCGCCTGG **TT TAAT CACGAAGTCAATAC** CCAGAGCC **TTCG**<br>GGCTCTGG GGCTCCGCA **CAGAGTTAG** CCACGGTTACCGAGTG AAA **GATGCAATG**<br>CACTCGGTGACCGTGG AGCGGAGCC **TT TACT CACGAAGTCAATAC GTCCAACC TT**<br>**CATGCTTACGACG** | 1138 |
| **3D-b-Plasmid**<br>Template for 3D-b tiles | **GTTCTAATACGACTCACTATA** GGGCACTGTTAGCCA **GGTACTACA** GGCGAGGGTGCGGTG AAA<br>**GTAAGATCG** CACCGCATCCTCGCC AGGCTAACGGTGTCC GACACGGTCGCA **GCTTCAATC**<br>GTCGGTCAC **TTCG** GTGACCGAC AGCGACTGTGTC **TT AT CACGAAGTCAATAC**<br>CCATGTGGCTGCCGA **CATGATTGC** TGCCTTGGAGTTCCCG AAA **GCTAGTTAG**<br>CGGGAACTTCAAGGCA ACGGCAGCTACATGG CCAGTTCTAGGA **CATTGCATC**<br>TCCGGGAGTGCCGCTG AAA **TCAGTACTC** CAGCGGCATTCCCGGA ACCTAGGACTGG **TT AA**<br>**CACGAAGTCAATAC** GCCTGATTCGTAGCA **GAGTACTGA** TAACTACGAGCGGATG AAA<br>**TCAATGCAC** CATCCGCTTGTAGTTA AGCTACGAGTCAGGC CCAGATCATGCA **GATACCTTC**<br>GAGCGGAGTTTCGGTG AAA **GATTGAAGC** CACCGAAATTCCGCTC AGCATGGTCTGG **TT AC**<br>**CACGAAGTCAATAC** CCGTCATTACTGCAA **GCTTAGAAG** GTCGGTCTGCGACCGG AAA<br>**CAAGTGGAA** CCGGTCGCGGACCGAC ATGCAGTAGTGACGG CCTACTTCGCCA **TCAGGTTAC**<br>ACGGGCATCGCAGCTG AAA **GTGTACCTA** CAGCTGCGGTGCCCGT AGGCGAGGTAGG **TT TA**<br>**CACGAAGTCAATAC** GCCAGATATGACCCA **TCGATAGAG** GCGTCAGAT **GTAA** ATCTGACGC<br>AGGGTCATGTCTGGC GGAGGTGCTGGA **CGATCTTAC** GCCGTCCTG **TTCG** CAGGACGGC<br>ACCAGCGCCTCC **TT TAAT CACGAAGTCAATAC** GGGAACGGTATGCCA **CATACATGG**<br>GCGTGTCTACATCCGC AAA **GCAATCATG** GCGGATGTGGACACGC AGGCATACTGTTCCC<br>GCTAGTTATGGA **CGTATCTAC** CCTGGTCGTCTCACTG AAA **GAAGTGATC**<br>CAGTGAGATGACCAGG ACCATAGCTAGC **TT TACT CACGAAGTCAATAC GTCCAACC TT**<br>**CATGCTTACGACG** | 1048 |

| Name (Note) | Sequence | length |
|---|---|---|
| **3D-yz-Plasmid**<br>Template for 3D-yz tiles | **GTTCTAATACGACTCACTATA** GGGCATCTATGTCGC **AAATAAA** CCTTGGCTGAGGGTG AAA<br>**GTAGATACG** CACCCTCGGCCAAGG GCGACATGGATGCCC CGAGAGCAGCCA **TAGGTACAC**<br>GCGCTACGGAGTATGTCGGATTGTATTTCTGGTTCATCGG AAA **TCATGAGAC**<br>CCGATGAATCAGAAATGCAATCCGGCATACTCTGTAGCGC AGGCTGTTCTCG **TT AT**<br>**CACGAAGTCAATAC** CCTGGAGG **TTCG** CCTCCAGG CCCTGCCCA **GATCACTTC**<br>GCCTCCGCTTCACGACTTAGTCGCTTGCGGTTTAGGCGCTG AAA **TGTAGTACC**<br>CAGCGCCTGAACCGCAGGCGACTAGGTCGTGAGGCGGAGGC AGGGCAGGG **TT AA**<br>**CACGAAGTCAATAC** GCCAAGGTCCGCCTG **AAATAAA** GGCAGGAGCGCTCCGG AAA<br>**TGTCAGAAG** CCGGAGCGTTCCTGCC CAGGCGGATCTTGGC GCTAGTTAGGGC **AATAATA**<br>GTGGTCCTGACCGGTG AAA **CTTCTAAGC** CACCGGTCGGGACCAC GCCCTAGCTAGC **TT AC**<br>**CACGAAGTCAATAC** CCGTCTGAG **TTCG** CTCAGACGG CCTCCGGCA **GCATACTTC**<br>ACGGACCGGGAGCGG AAA **CTCTATCGA** CCGCTCCTGGTCCGT AGCCGGAGG **TT TA**<br>**CACGAAGTCAATAC** CCAGGGTC **TTCG** GACCCTGG CCAGCCGAA **GTGACTGTA**<br>TCCAGACGGCGAGTG AAA **CCATGTATG** CACTCGCTGTCTGGA ATCGGCTGG **TT TAAT**<br>**CACGAAGTCAATAC** CCTCGGGA **TTCG** TCCCGAGG GCTCGCGCA **CTAACTAGC**<br>CAGGTCCGCTTGGCTG AAA **GAAGTATGC** CAGCCAAGTGGACCTG AGCGCGAGC **TT TACT**<br>**CACGAAGTCAATAC** CCACCTGC **TTCG** GCAGGTGG CCCTCGCCA **CTTCTGACA**<br>GCGCGTAGAGATTCGG AAA **TACAGTCAC** CCGAATCTTTACGCGC AGGCGAGGG **TT ATAT**<br>**CACGAAGTCAATAC** **GTCCAACC TT CATGCTTACGACG** | 1039 |
| **3D-zz-Plasmid**<br>Template for 3D-zz tiles | **GTTCTAATACGACTCACTATA** GGGCTGTC **TTCG** GACAGCCC GACCGTACGGGA **TAGGTACAC**<br>GCTCGGATGACGTTCGGAGATTCTGAAGTTCTAAGCCGTG AAA **TCATGAGAC**<br>CACGGCTTGGAACTTCGGAATCTCTGAACGTCGTCCGAGC ACCCGTGCGGTC **TT AT**<br>**CACGAAGTCAATAC** CCAGGACC **TTCG** GGTCCTGG GACAGTCGGCCA **CATTGCATC**<br>ACCTGCCGTAGCTGTG AAA **TCAGTACTC** CACAGCTATGGCAGGT AGGCCGGCTGTC **TT AA**<br>**CACGAAGTCAATAC** CCGTAGGCCTAAGCA **GAGTACTGA** TAGAGAATACTCGTCG AAA<br>**TCAATGCAC** CGACGAGTGTTCTCTA AGCTTAGGTCTACGG CTCCGTCGTGCA **GATACCTTC**<br>GCGCATAGT **TTCG** ACTATGCGC AGCACGGCGGAG **TT AC CACGAAGTCAATAC**<br>GCCTGTTAAGTAGCG **AAATAAA** GGTATGAGAGCGTCGG AAA **CAAGTGGAA**<br>CCGACGCTTTCATACC CGCTACTTGACAGGC CCTCAGTCCGCA **TCAGGTTAC**<br>GGCACCGGAAGTCCTG AAA **GTGTACCTA** CAGGACTTTCGGTGCC AGCGGATTGAGG **TT TA**<br>**CACGAAGTCAATAC** **GTCCAACC TT CATGCTTACGACG** | 654 |
| **3D-c-Plasmid**<br>Template for 3D-c tiles | **GTTCTAATACGACTCACTATA** GGGCATTGGGCTGCA **CCGATTCTA** GTTATCATCCGAGGC AAA<br>**TGAGTTCTG** GCCTCGGGTGATAAC AGCAGCCCGATGCCC GACCTGGGACCA **CTACTCGTA**<br>CCGGCTATGTTTAGGTGATGGTGAACATTTCGTTCAATAGGCGTGGTATATGTTAGATCGTAATCGAG<br>CTGTAGTCGATGAGGGCTCGTG AAA **GGTTAGCAG**<br>CACGAGCTCTCATCGGCTACAGTTCGATTATGATCTAGCATATACTACGCCTGTTGAACGGAATGTTT<br>ACCATCATCTAAACGTAGCCGG AGGTCCTAGGTC **TT AT CACGAAGTCAATAC**<br>CCGACCGTGGGACCA **GTTGCTCTA** TAACTTGTGCTCGTG AAA **GTCTTGAAG**<br>CACGAGCGCAAGTTA AGGTCCCATGGTCGG GCATCAGGAGCC **AATAATA**<br>GCTCCCATTCACAGTGCTGAGTGGTACATTGATGGCAGTAGATTGGGCAGGAGTAACTATGATACAAT<br>GTTACTTAATCACGGAGGTCGG AAA **TACGAGTAG**<br>CCGACCTTCGTGATTGAGTAACGTTGTATCGTAGTTATTCCTGCCTAATCTATTGCCATCGATGTACT<br>ACTCAGCGCTGTGAGTGGGAGC GGCTCTTGATGC **TT AA CACGAAGTCAATAC**<br>GCCTCCGGTGGCGCA **GAGAACATG** CCACGAGTGCGTTGCG AAA **GTAACCTGA**<br>CGCAACGCGCTCGTGG AGCGCCACTGGAGGC CCAGTGCGCCTA **TCCTATGAG**<br>CAGTAGCGTACGAAGTGTGAAACGGATGATCAGGGACGTGG AAA **TAGAATCGG**<br>CCACGTCCTTGATCATTCGTTTCATACTTCGTGCGCTACTG AAGGCGTACTGG **TT AC**<br>**CACGAAGTCAATAC** GCATTCCGAAGACCG **AAATAAA** GCAGGGTGCGCCTCTG AAA<br>**GTAAGGACA** CAGAGGCGTACCCTGC CGGTCTTTGGAATGC CCGTGGGACTCG **AATAATA**<br>GGATGTACGGATTGCGGAGTACATTGATCTCTGGAAGCCAG AAA **TAGAGCAAC**<br>CTGGCTTCTAGAGATCGATGTACTTCGCAATCTGTACATCC CGAGTTCCACGG **TT TA**<br>**CACGAAGTCAATAC** GCACGGGCATCCGGA **CGTGAATGA** CCAGTGGTCCAGGTG AAA<br>**GTAGATACG** CACCTGGGCCACTGG ACCGGATGTCCGTGC GCTCGTAGCGGA **TAGGTACAC**<br>GGCGACTTGATATTGGCTTTAGCTGAGTCGCTAGGGCCTG AAA **TCATGAGAC**<br>CAGGCCCTGGCGACTCGGCTAAAGTCAATATCGAGTCGCC ACCGCTGCGAGC **TT TAAT**<br>**CACGAAGTCAATAC** GCCAGAGCATGACCA **GTAACAAGC** CCGAGGCGGACGGTG AAA<br>**CTAACGAGA** CACCGTCTGCCTCGG AGGTCATGTTCTGGC CCATCGGAGGCA **CTTGGATCA**<br>GCGCAGGTTGAAGATGATCCATAGGCTGGATTCCAATCGC AAA **CATGTTCTC**<br>GCGATTGGGATCCAGCTTATGGATTATCTTCAGCCTGCGT AGCCTCTGATGG **TT TACT**<br>**CACGAAGTCAATAC** **GTCCAACC TT CATGCTTACGACG** | 1594 |

| Name (Note) | Sequence | length |
|---|---|---|
| **3D-d-Plasmid**<br>Template for 3D-d tiles | **GTTCTAATACGACTCACTATA** GGGCTCTGCCAGGCA **CCTTGAGAA** CCGTGCGTGATCCTGG AAA<br>**TTCATAGCG** CCAGGATCGCGCACGG AGCCTGGCGGAGCTC GCCCTTGCGGTA **GATCACTTC**<br>GCTCGAATTCAGCTCGTAGGCAAGGCAGACAATTGCAACTG AAA **TGTAGTACC**<br>CAGTTGCAGTTGTCTGTCTTGCCTGCGAGCTGGATTCGAGC AACCGCGAGGGC **TT AT**<br>**CACGAAGTCAATAC** CAGTGTGGACGAGCA **GTGCATTGA** TCTGACATAGCGGTG AAA<br>**TACATGTGG** CACCGCTGTGTCAGA AGCTCGTCTACACTG GCTCGGTGCGCA **GACAGTTGA**<br>CCGTACGTGGTCGTG AAA **GAACTTACG** CACGACCGCGTACGG AGCGCATCGAGC **TT AA**<br>**CACGAAGTCAATAC** GCCATGGTAGCAGCA **TTCCACTTG** GCCGTTCGACGACGG AAA<br>**CTAACTCTG** CCGTCGTTGAACGGC AGCTGCTATCATGGC GTGCGTTCTGGA **TAGCCATTC**<br>AGGGATGTCCAGCGG AAA **TGAGCAAAG** CCGCTGGGCATCCCT ACCAGAGCGCAC **TT AC**<br>**CACGAAGTCAATAC** GAGCGGTAGGTAGCA **GACTTACTC** CATCCGTGATGCGTG AAA<br>**GAACAGATC** CACGCATTACGGATG AGCTACCTGCCGCTC CCAGGTCCTGCA **TGCAGATTC**<br>GACCAGGTGCTCGTG AAA **GGTTACATG** CACGAGCGCCTGGTC AGCAGGGCCTGG **TT TA**<br>**CACGAAGTCAATAC** CCATGGGTGATCGCA **GATGACTTG** GCCAGACGCCGTCCGG AAA<br>**TGTCAGAAG** CCGGACGGTGTCTGGC AGCGATCATCCATGG GAGTCTCTGCCA **CATCGTTAC**<br>CCGCTGAGACAGTCGG AAA **CTTCTAAGC** CCGACTGTTTCAGCGG AGGCAGGGACTC **TT TAAT**<br>**CACGAAGTCAATAC** CCTCAACGGTCACGG **AAATAAA** GCTCGAGTTCCGACGG AAA<br>**CTAGCATGA** CCGTCGGAGCTCGAGC CCGTGACTGTTGAGG GCACCGTATAGG **AATAATA**<br>CCGACTCGCCTCGGTC AAA **CATTCGTAC** GACCGAGGTGAGTCGG CCTATGCGGTGC **TT TACT**<br>**CACGAAGTCAATAC** GCAGTCTGTAGCGCA **TGTTACCTC** GCCGTCCGAGACCAG AAA<br>**CTAGGAACA** CTGGTCTTGGACGGC AGCGCTACGGACTGC CCATCTCAGGCA **GCATACTTC**<br>TACCGGTTCACGCGG AAA **CTCTATCGA** CCGCGTGGACCGGTA AGCCTGGGATGG **TT ATAT**<br>**CACGAAGTCAATAC** GCAGGAGCAGATCCA **GGCTATGTA** TTGTAACGGTGAGCG AAA<br>**GTCTAAGGA** CGCTCACTGTTACAA AGGATCTGTTCCTGC CGTGCGAAGGAA **GTGACTGTA**<br>TCTACCGTGAGGGTG AAA **CCATGTATG** CACCCTCGCGGTAGA ATCCTTTGCACG **TT AATT**<br>**CACGAAGTCAATAC** GCCTCGTACAGAGCA **TTCCGAAAG** CGCAGGTTATCCGCG AAA<br>**GAAGTCTCA** CGCGGATGACCTGCG AGCTCTGTGCGAGGC GCTCGGACGGGA **TTCATTGCC**<br>ACCGGCTGCGACGTG AAA **GAACATAGG** CACGTCGTAGCCGGT ACCCGTTCGAGC **TT ACTT**<br>**CACGAAGTCAATAC** **GTCCAACC TT CATGCTTACGACG** | 1678 |
| **3D-e-Plasmid**<br>Template for 3D-e tiles | **GTTCTAATACGACTCACTATA** GGGCCATAGCTAGGA **TGAGCATTC** GCCGATCTGGACGGTG AAA<br>**CAGATGTAG** CACCGTCCGGATCGGC ACCTAGCTGTGGCCC CCGGATAGGCCA **GCATAGATG**<br>ACCAGGCGATATCCGG AAA **TCTGAAGTG** CCGGATATTGCCTGGT AGGCCTGTCCGG **TT AT**<br>**CACGAAGTCAATAC** CGACCTTACCGAGCA **CTCGATACA** GCACATCTGCTTCTGG AAA<br>**CAACATACG** CCAGAAGCGGATGTGC AGCTCGGTGAGGTCG CCCATGTCGACA **TACTTAGCG**<br>CCAGTGATGCGTCTCG AAA **CAATGTCTC** CGAGACGCGTCACTGG AGTCGATATGGG **TT AA**<br>**CACGAAGTCAATAC** CCTGATCTTAGGGTG **AAATAAA** CCGGACAGATCCCGTG AAA<br>**CATCTTGAC** CACGGGATTTGTCCGG CACCCTAGGATCAGG GCACCGTTCGGG **AATAATA**<br>CAGTCCGGCAGGACGG AAA **CCTGTACTA** CCGTCCTGTCGGACTG CCCGAGCGGTGC **TT AC**<br>**CACGAAGTCAATAC** CCTCACTAGCTTCGA **CTACAAGGA** CCATTACGCGACCGG AAA<br>**CTTCACTGA** CCGGTCGTGTAATGG ACGAAGCTGGTGAGG CCTCTGACCGCA **GTGTCAAAC**<br>GCTCGAGTGCGACGG AAA **TGATCGTAG** CCGTCGCGCTCGAGC AGCGGTTAGAGG **TT TA**<br>**CACGAAGTCAATAC** GCCAGCTTTCAGGCA **TACGCTTAC** CCTGACGTTCGGTCG AAA<br>**GCATGTCTA** CGACCGAGCGTCAGG AGCCTGAAGGCTGGC CCTGCTGCAGCA **CGATGAAGA**<br>CCCGCTCGCTCGGTG AAA **TGAATGAGC** CACCGAGTGAGCGGG AGCTGCGGCAGG **TT TAAT**<br>**CACGAAGTCAATAC** GCAGTGGATATGGCA **TACCAGTTC** AGTATAGTTGCGGTCG AAA<br>**CAATGATGG** CGACCGCAGCTATACT AGCCATATTCACTGC CCAGGTTACGCA **TCAGACATC**<br>AGGGTAATAGTCACGG AAA **TTCAGATCG** CCGTGACTGTTACCCT AGCGTAGCCTGG **TT TACT**<br>**CACGAAGTCAATAC** GCCAGCTTGAGGCCA **TACCGTAAC** GTAGTGGGACATCCCG AAA<br>**TCTCGTATC** CGGGATGTTCCACTAC AGGCCTCAGGCTGGC GCTACTCATGCA **TGACGAAAG**<br>CCTGGCAGCTCACGTG AAA **TTGGTATCC** CACGTGAGTTGCCAGG AGCATGGGTAGC **TT ATAT**<br>**CACGAAGTCAATAC** GCCATCTTTGACAGC **AAATAAA** GCTCCATGAACGTCGG AAA<br>**CAGCTATCA** CCGACGTTTATGGAGC GCTGTCAGAGATGGC CCATGGTAACGG **AATAATA**<br>GTCTGTCTGAGCCCTG AAA **GTACTTACC** CAGGGCTCGGACAGAC CCGTTGCCATGG **TT AATT**<br>**CACGAAGTCAATAC** **GTCCAACC TT CATGCTTACGACG** | 1460 |

| Name (Note) | Sequence | length |
|---|---|---|
| **3D-f-Plasmid**<br>Template for 3D-f tiles | GTTCTAATACGACTCACTATA GGGCAAGCTGTTCCA TCAATCAGC ACCTGGCTAACTCGTG AAA<br>GAAGATTGC CACGAGTTGGCCAGGT AGGAACAGTTTGCCC GACCGGACCGCA GGTAGACTA<br>CACGCCGAGTACTCGATGGTACAGGATGTACCGCTGTATAC AAA TCATTCACG<br>GTATACAGTGGTACATTCTGTACCGTCGAGTATTCGGCGTG AGCGGTTCGGTC TT AT<br>CACGAAGTCAATAC GCATGATAGACGGGA CATGTAACC<br>TCCGTTGCTACATTGGTCATGCGTTGACAAGAGTACGACTG AAA GAGGATACA<br>CAGTCGTATTCTTGTCGACGCATGGCCAATGTGGCAACGGA ACCCGTCTGTCATGC<br>GCCAGGTGCCCA CTTCAAGAC GGTGGCATGTTATCGG AAA CTCATAGGA<br>CCGATAACGTGCCACT AGGGCATCTGGC TT AA CACGAAGTCAATAC GCACAGTGGCTGACA<br>GAGACATTG GCGAACTGAGGTGACTCAGACCTTAAGTTTCTGCTCAGTG AAA TGAACAGTG<br>CACTGAGCGGAAACTTGAGGTCTGGGTCACCTTAGTTCGT AGTCAGCCGCTGTGC<br>CCGAGTAATGCA TGTATCCTC GGGCTCGGCCAGACTG AAA GAATGGCTA<br>CAGTCTGGTCGAGCCC AGCATTGCTCGG TT AC CACGAAGTCAATAC GCCAAGGTGCGAGCA<br>TAGTACAGG TCCGGATGGTGACAAGTCATGAGGATGTTCAGTTGGACTG AAA GATCATGTG<br>CAGTCCAATTGAACATTCTCATGATTTGTCACTATCCGGA AGCTCGCATCTTGGC<br>CCAGGTGGCAGC AATAATA GCTCCCAGGGTCCGTG AAA GAATCTGCA CACGGACCTTGGGAGC<br>GCTGCTACCTGG TT TA CACGAAGTCAATAC GCAGTGGTCCTTGCA GTACGAATG<br>GGTCCGAGGATCGGTG AAA GAGTAAGTC CACCGATCTTCGGACC AGCAAGGATCACTGC<br>CCGTATGTGTCA TGTCCTTAC CTACGCGGTGCGCTCG AAA TGATCCAAG<br>CGAGCGCATCGCGTAG AGACACGTACGG TT TAAT CACGAAGTCAATAC GGAGTGGCTGCTTAA<br>CTTGACCTA GCTGCCCAC GTAA GTGGGCAGC ATAAGCAGTCACTCC CCTCAGGGCGGA<br>CCACATGTA CGCGAAGTC TTCG GACTTCGCG ACCGCCTTGAGG TT TACT<br>CACGAAGTCAATAC GCACCGTTCACCCTA CTACGATCA GCAGACTTACGGACCG AAA<br>GAATGCTCA CGGTCCGTGAGTCTGC AAGGGTGAGCGGTGC CCTAGTCCCGCA CAGAGTTAG<br>GCGCTAGGAACGCCTG AAA GATGCAATG CAGGCGTTTCTAGCGC AGCGGGGCTAGG TT ATAT<br>CACGAAGTCAATAC CATGTAGGCCCTGCA GCTCATTCA TGATCTGTCCTAGTGG AAA<br>TGTATCGAG CCACTAGGGCAGATCA AGCAGGGCTTACATG CCTGTGAGGCCA GATCTGTTC<br>GCCTCCGTGACGCGTG AAA GTAACGATG CACGCGTCGCGGAGGC AGGCCTTACAGG TT AATT<br>CACGAAGTCAATAC GTCCAACC TT CATGCTTACGACG | 1626 |
| **3D-g-Plasmid**<br>Template for 3D-g tiles | GTTCTAATACGACTCACTATA GGGCTGTTCGAGTGC AAATAAA CGATCGCTGACCGCTG AAA<br>GAAGCTAAG CAGCGGTCGGCGATCG GCACTCGGACAGCCC GACCAGTGACCG AATAATA<br>CCATCGTGGACGGCTATGTGAGCGGTATTCATTACCGGCTG AAA GCTTGTTAC<br>CAGCCGGTGATGAATATCGCTCACGTAGCCGTTCACGATGG CGGTCGCTGGTC TT AT<br>CACGAAGTCAATAC CAGCTATCGACTCGA CCTATGTTC GCGTGATGTGGACCCG AAA<br>CAAGTCATC CGGGTCCATATCACGC ACGAGTCGGTAGCTG GCAGATCTGGAA TCTCGTTAG<br>CCGACTATAGGGTGCG AAA TAGTCTACC CGCACCCTGTAGTGCGG ATCCAGGTCTGC TT AA<br>CACGAAGTCAATAC GCATTGTAGAACCGA CGATCTGAA GGCGCTGTGGCTCGAC AAA<br>TAGATTCCG GTCGAGCCGCAGCGCC ACGGTTCTGCAATGC CCGTAGGAGGCA CTAACTAGC<br>GCTGGGAGTTCCACCG AAA GAAGTATGC CGGTGGAATTCCCAGT AGCCTCTTACGG TT AC<br>CACGAAGTCAATAC GCATGATAGACGCCA GGATACCAA GCGAGTCGAGTGACGG AAA<br>TCCTTGTAG CCGTCACTTGACTCGC AGGCGTCTGTCATGC GCATCTCATCCA CTTCTGACA<br>GCGTGGCGACTAACGG AAA TACAGTCAC CCGTTAGTTGCCACGT AGGATGGGATGC TT TA<br>CACGAAGTCAATAC CACCTAGTACGAGCA GGTAAGTAC GTCGCTTGTGACGGGC AAA<br>GTAAGCGTA GCCCGTCATAAGCGAC AGCTCGTATTAGGTG CCGGTTCAGGGA TCATGCTAG<br>CCCGTCATGGCACGTG AAA GGCAATGAA CACGTGCCGTGACGGG ACCCTGGACCGG TT TAAT<br>CACGAAGTCAATAC GCACTCGTTACGCCA TCATGACTC GGCAGACGACTGCGTG AAA<br>TACATAGCC CACGCAGTTGTCTGCC AGGCGTAATGAGTGC CCAGGTCCTGCA GCAATCTTC<br>GCTGTGATTCGTTCGG AAA CTGTAGTGA CCGAACGAGTCACAGC AGCAGGGCCTGG TT TACT<br>CACGAAGTCAATAC CCATGTTCGTCCGGA GATAGTTCC GTGGCATCT GTAA AGATGCCAC<br>ACCGGACGGACATGG GCTGCGGACTCA TGTTCCTAG GGTCGCGGA TTCG TCCGCGACC<br>AGAGTCTGCAGC TT ATAT CACGAAGTCAATAC CCATGCGTCCGACCA GGAATGTAC<br>GTCGGACTTCACGGTG AAA GAACTGGTA CACCGTGAGGTCCGAC AGGTCGGATGCATGG<br>GCAATGGTACGA TCCTTAGAC GAGGTCTTATCCACGG AAA CTAGTGTCA<br>CCGTGGATGAGACCTC ACGTACTATTGC TT AATT CACGAAGTCAATAC CCAGCAGTGTCCGGA<br>CTTTGAACG CGAGGTCTTTGCGGTG AAA GTTACGGTA CACCGCAAGGACCTCG<br>ACCGGACATTGCTGG GGTATGCTGGCA TGAGACTTC CACTGGCTGAAATCTG AAA<br>CTACTGCTA CAGATTTCGGCCAGTG AGCCAGTATACC TT ACTT CACGAAGTCAATAC<br>GTCCAACC TT CATGCTTACGACG | 1658 |

| Name (Note) | Sequence | length |
|---|---|---|
| **3D-xy-Plasmid**<br>Template for 3D-xy tiles | **GTTCTAATACGACTCACTATA** GGGCTCGCGCAGGGA **CACTTCAGA** GCCGTTATC **GTAA**<br>GATAACGGC ACCCTGCGTGAGCTC GCCCATGGAGGA **TGTCGATTG** CCGTGATTGCTCGGTG<br>AAA **TCAACTGTC** CACCGAGCGATCACGG ACCTCCGTGGGC **TT AT CACGAAGTCAATAC**<br>GCAGGCTG **TTCG** CAGCCTGC CCAGGCGCA **CACTGTTCA** AGCCCGTGGCGAAGTG AAA<br>**CATCTATGC** CACTTCGCTACGGGCT AGCGCCTGG **TT AA CACGAAGTCAATAC** GCCACCTG<br>**TTCG** CAGGTGGC CCAGCCGCA **CACATGATC** GCCAGGCTGATACCGG AAA **CGCTAAGTA**<br>CCGGTATCGGCCTGGC AGCGGCTGG **TT AC CACGAAGTCAATAC** GCTCCTGG **TTCG**<br>CCAGGAGC GCTCCCGCA **CTACATCTG** GGGACTCGGGTATCTC AAA **CTTGTTCCA**<br>GAGATACCTGAGTCCC AGCGGGAGC **TT TA CACGAAGTCAATAC** CGGAGCTG **TTCG**<br>CAGCTCCG CCAGCCGGA **CGTATGTTG** CCCGTGAGCTTATCCG AAA **GTTTGACAC**<br>CGGATAAGTTCACGGG ACCGGCTGG **TT TAAT CACGAAGTCAATAC** GCAGACGG **TTCG**<br>CCGTCTGC GCTCCCGGA **GTCAAGATG** GCTGACATGACGAGGC AAA **TCTTCATCG**<br>GCCTCGTCGTGTCAGC ACCGGGAGC **TT TACT CACGAAGTCAATAC** CCAGCCTG **TTCG**<br>CAGGCTGG CGCTCGGGA **TCAGTGAAG** GCAGCCTGACGACGTG AAA **GATGTCTGA**<br>CACGTCGTTAGGCTGC ACCCGAGCG **TT ATAT CACGAAGTCAATAC** CCAGCTGG **TTCG**<br>CCAGCTGG GCCTCCGGA **TAGACATGC** CACGAGGTACTTGGGC AAA **CTTTCGTCA**<br>GCCCAAGTGCCTCGTG ACCGGAGGC **TT AATT CACGAAGTCAATAC** CATCCCTG **TTCG**<br>CAGGGATG CCAGCCGCA **CCATCATTG** CCTAGGCTACATCCCG AAA **GAATCGTCA**<br>CGGGATGTGGCCTAGG AGCGGCTGG **TT ACTT CACGAAGTCAATAC** GCCTCGTG **TTCG**<br>CACGAGGC GCTGCCGGA **GATACGAGA** CAGCCTGGCTCAGGGC AAA **GTTAGTCGA**<br>GCCCTGAGTCAGGCTG ACCGGCAGC **TT TATA CACGAAGTCAATAC** GCCTGAGG **TTCG**<br>CCTCAGGC GGTCCGGGA **TGATAGCTG** GGGGTGCGAGTACGTG AAA **GATTGACTG**<br>CACGTACTTGCACCCC ACCCGGACC **TT ATTA CACGAAGTCAATAC** **GTCCAACC TT**<br>**CATGCTTACGACG** | 1304 |
| **3D-xz-Plasmid**<br>Template for 3D-xz tiles | **GTTCTAATACGACTCACTATA** GGGCTAGG **TTCG** CCTAGCCC GCCCTGGGA **TCACTACAG**<br>CCCGAGCGCCAGCATTCTCACATGCGATCGTTCCGGACGG AAA **TTCTCAAGG**<br>CCGTCCGGGACGATCGTATGTGAGGATGCTGGTGCTCGGG ACCCAGGGC **TT AT**<br>**CACGAAGTCAATAC** CAACCCGG **TTCG** CCGGGTTG CGAGCGGCA **GACTACCTA**<br>CCCGTGGTCCTAGTGGACAGGCCGACGACTCGTCGCGGTG AAA **GCTGATTGA**<br>CACCGCGATGAGTCGTTGGCCTGTTCACTAGGGCCACGGG AGCCGCTCG **TT AA**<br>**CACGAAGTCAATAC** GCCTGGTG **TTCG** CACCAGGC GCTCCCGGA **TGACACTAG**<br>CCCTGCTTGACAGGTG AAA **GTTAGAACG** CACCTGTCGAGCAGGG ACCGGGAGC **TT AC**<br>**CACGAAGTCAATAC** GCTCCGTG **TTCG** CACGGAGC CGAGCCCGA **TAGCAGTAG**<br>TCACCCTGCGGCTGCG AAA **GAGTCATGA** CGCAGCCGTAGGGTGA ACGGGCTCG **TT TA**<br>**CACGAAGTCAATAC** GCACAGGCGAGCACA **CGGAATCTA** GCTAACGAC **GTAA** GTCGTTAGC<br>AGTGCTCGTCTGTGC GCTGCTGATGCA **TGGAACAAG** CATCTTTGTCCTACG AAA **TAGGTCAAG**<br>CGTAGGATAAAGATG AGCATCGGCAGC **TT TAAT CACGAAGTCAATAC** CCAGCTGG **TTCG**<br>CCAGCTGG CCAGCCCGA **TGACGATTC** CTGGAGCGGGAGCGC AAA **GGAACTATC**<br>GCGCTCCTGCTCCAG ACGGGCTGG **TT TACT CACGAAGTCAATAC** CCTCGCTG **TTCG**<br>CAGCGAGG GGAGCGCGA **TCGACTAAC** ACGGATCGCGAGGTC AAA **GTACATTCC**<br>GACCTCGTGATCCGT ACGCGCTCC **TT ATAT CACGAAGTCAATAC** CCAGCTGG **TTCG**<br>CCAGCTGG CCTACCCCA **CAGTCAATC** GGCGTGGGACGAAGC AAA **CGTTCAAAG**<br>GCTTCGTTCCACGCC AGGGGTAGG **TT AATT CACGAAGTCAATAC** CCGTCAGTAGCAGCA<br>**CGTTCTAAC** CCGGTCTTAATGGGCG AAA **GAGGTAACA** CGCCCATTGAGACCGG<br>AGCTGCTATTGACGG GCAGCGCAGGCA **CGCTATGAA** GGTCCGACG **TTCG** CGTCGGACC<br>AGCCTGTGCTGC **TT ACTT CACGAAGTCAATAC** CCACCGTAGTGCGTG **AAATAAA**<br>GCGTTCGGATCTCCGC AAA **CTTTCGGAA** GCGGAGATTCGAACGC CACGCACTGCGGTGG<br>CCTACGCGCACA **CTTAGCTTC** GACCGCAGGGCTAGCG AAA **TAGGTAGTC**<br>CGCTAGCCTTGCGGTC AGTGCGTGTAGG **TT TATA CACGAAGTCAATAC** **GTCCAACC TT**<br>**CATGCTTACGACG** | 1395 |

| Name (Note) | Sequence | length |
|---|---|---|
| **3D-h-Plasmid**<br>Template for 3D-h tiles | GTTCTAATACGACTCACTATA GGGCCTTCGAGCGCA TACGTCTTC GGTAGTATGCTCTCG AAA<br>GAATCAAGG CGAGAGCGTACTACC AGCGCTCGGAGGCCC GCTCCGGTACCA TAGTTGAGC<br>CTGGTGCTG TTCG CAGCACCAG AGGTACTGGAGC TT AT CACGAAGTCAATAC<br>CCTCATGCGCTGGGA TACAAGTGC CTCGGACTGGTCGTG AAA GCGTTACTA<br>CACGACCGGTCCGAG ACCCAGCGTATGAGG GCAGGTGCTGGA TCACTACAG<br>GCCTTGTGATCGCACTAGTGGCCGTCGTCTATCGCGACGG AAA TTCTCAAGG<br>CCGTCGCGGTAGACGATGGCCACTGGTGCGATTACAAGGC ACCAGCGCCTGC TT AA<br>CACGAAGTCAATAC CCAGCGTTTGCGACG AAATAAA CGAATGCGGAGACGG AAA CATTGGAAC<br>CCGTCTCTGCATTCG CGTCGCAGACGCTGG CCTCCGAGCGCA GACTACCTA<br>CCGGTGCTTGTAATGGCGTTGATGCCGTGTCGGCGAGGTG AAA GCTGATTGA<br>CACCTCGCTGACACGGTATCAACGTCATTACAGGCACCGG AGCGCTTGGAGG TT AC<br>CACGAAGTCAATAC CCAGGAGG TTCG CCTCCTGG GCACCGCGA GTTGCTAGA<br>ACCCGGTCTGGTCGTTTACCTATAGCATACTATCAGCGGTG AAA GAAGACGTA<br>CACCGCTGGTAGTATGTTATAGGTGAACGACCGGACCGGGT ACGCGGTGC TT TA<br>CACGAAGTCAATAC GCACCGTG TTCG CACGGTGC GCTGCGGTA TCATCTCTG<br>CCCAGAGGTTCGGAACGAGCGGTCTGAACGATGATGCGGTG AAA GCACTTGTA<br>CACCGCATTATCGTTCGGACCGCTTGTTCCGAGCCTCTGGG AACCGCAGC TT TAAT<br>CACGAAGTCAATAC GCAGCGTACATTCCA GAGATTCGA GCGGCAGGGATGCGTG AAA<br>TCAAGATCC CACGCATCTCTGCCGC AGGAATGTGCGCTGC GCTGCTAGCGGA TGACACTAG<br>CCGGAGATGATATGTG AAA GTTAGAACG CACATATCGTCTCCGG ACCGCTGGCAGC TT TACT<br>CACGAAGTCAATAC GCAGCCGTGGCAGCA CTGATCATC GCATCTTGGCCGAGGC AAA<br>GGACATTCA GCCTCGGCTAAGATGC AGCTGCCATGGCTGC GCGTGGGACGGA TAGCAGTAG<br>CCTGACCGTGGATGCG AAA GAGTCATGA CGCATCCATGGTCAGG ACCGTCTCACGC TT ATAT<br>CACGAAGTCAATAC CCTGGCATAGCGCTG AAATAAA GCCAGAGTCTCACCGG AAA<br>CTTACAGAC CCGGTGAGGCTCTGGC CAGCGCTGTGCCAGG CCATCTGGAGGC AATAATA<br>GCTCCGATCCTCGCGG AAA CTCAATGGA CCGCGAGGGTCGGAGC GCCTCTAGATGG TT AATT<br>CACGAAGTCAATAC GGCACCTG TTCG CAGGTGCC GCTCCGGGA CTAAGACTG<br>CCGGACGTGACGGTG AAA CTAGTCCTA CACCGTCGCGTCCGG ACCCGGAGC TT ACTT<br>CACGAAGTCAATAC GCCTCGAG TTCG CTCGAGGC GGTCCCGCA GCATCTATC<br>TCGGACGTCCTGCTG AAA GATAGTAGC CAGCAGGGCGTCCGA AGCGGGACC TT TATA<br>CACGAAGTCAATAC GCCTCGTG TTCG CACGAGGC GGTCCCGCA TGACTCTAC<br>CCGTCCTGCCGAGGC AAA GGATGAATC GCCTCGGTAGGACGG AGCGGGACC TT ATTA<br>CACGAAGTCAATAC GTCCAACC TT CATGCTTACGACG | 1827 |
| **3D-i-Plasmid**<br>Template for 3D-i tiles | GTTCTAATACGACTCACTATA GGGCCAGTACAGGCA CGGAATCTA GCGCCTCGAGCCGTG AAA<br>CTAAGGTAC CACGGCTTGAGGCGC AGCCTGTATTGGCTC GATCGTAGCGCA TGGAACAAG<br>GCACGCAGATCCTCG AAA TAGGTCAAG CGAGGATTTGCGTGT AGCGCTGCGATC TT AT<br>CACGAAGTCAATAC CAGTAGTTGCCTGCA TTGGACTTG CGATCCGGCACCTGG AAA<br>CGTAACTAG CCAGGTGTCGGATCG AGCAGGCAGCTACTG GCGCTGGCAGCA TGACGATTC<br>CACGGTCGCCGAGCG AAA GGAACTATC CGCTCGGTGACCGTG AGCTGCTAGCGC TT AA<br>CACGAAGTCAATAC CCTATTGGGATAGCA CTGTGAAAC GCGGAATGAGACCGC AAA<br>CATTCTGAG GCGGTCTTATTCCGC AGCTATCCTAATAGG GCGTCTTCGGCA TCGACTAAC<br>CCGCTGCGCAGGAGC AAA GTACATTCC GCTCCTGTGCAGCGG AGCCGAGGACGC TT AC<br>CACGAAGTCAATAC CCTCCAGTAGAGCCA GCAAGTAAG GTGCGAGTATGCCTG AAA<br>CTATCTCGA CAGGCATGCTCGCAC AGGCTCTATTGGAGG CCATATGCTGCA CAGTCAATC<br>ACCGACCGTCGGTGC AAA CGTTCAAAG GCACCGATGGTCGGT AGCAGCGTATGG TT TA<br>CACGAAGTCAATAC GCAGACGG TTCG CCGTCTGC CCTGGGCCA TGAGAATGC<br>GGGTCCGGCAGCACCG AAA TACAACTGG CGGTGCTGTCGGACCC AGGCCCAGG TT TAAT<br>CACGAAGTCAATAC CCTCGCTG TTCG CAGCGAGG GCTCGCCCA TCATCAAGC<br>ACCCGCTGCACATGCG AAA CAAGGTATG CGCATGTGTAGCGGGT AGGGCGAGC TT TACT<br>CACGAAGTCAATAC CCAGCCGGACGCCGA CTTGTGATG GCGGAATTGGCTCGCG AAA<br>GTAACTCTC CGCGAGCCGATTCCGC ACGGCGTCTGGCTGG GCTCCGCTCGGA CTATGCCTA<br>CCCAGAGTTGGTAGCG AAA CTGTCTACA CGCTACCAGCTCTGGG ACCGAGTGGAGC TT ATAT<br>CACGAAGTCAATAC GCAGTTTATCGACCA TTCTGTCAC GGCACTATTCATGTAG AAA<br>TCTATCCAG CTACATGAGTAGTGCC AGGTCGATGAACTGC GGTTGTGGCACA TCCTAAGTC<br>TCGGTCGGCTCTAGGC AAA TGTTCACTC GCCTAGAGTCGACCGA AGTGCCGCAACC TT AATT<br>CACGAAGTCAATAC GGGTCGAGCATGCTG AAATAAA CCGTGTATTCACGGTG AAA<br>GTACTAGGA CACCGTGAGTACACGG CAGCATGTTCGACCC GCTCTGTAGTGG AATAATA<br>GCTACCTGGGACCTCC AAA GATTCAGAC GGAGGTCCTAGGTAGC CCACTGCAGAGC TT ACTT<br>CACGAAGTCAATAC GGTCCGGA TTCG TCCGGACC CCAGGCCCA GTATGACGA<br>GCCGTGCGCGAGGTG AAA TTGCCTATC CACCTCGTGCACGGC AGGGCCTGG TT TATA<br>CACGAAGTCAATAC CGAGCGTG TTCG CACGCTCG CCAGGCCCA CTAAGCAGA<br>GAGATCTTACTGCCC AAA GCATGAAAG GGGCAGTGAGATCTC AGGGCCTGG TT ATTA<br>CACGAAGTCAATAC CCTCGCTG TTCG CAGCGAGG GCCTCCCGA GTAGCTAAC<br>CGGGTCGTCCAGCCG AAA CAACTTGAG CGGCTGGGCGACCCG ACGGGAGGC TT CTTA<br>CACGAAGTCAATAC GTCCAACC TT CATGCTTACGACG | 1825 |

| Name (Note) | Sequence | length |
|---|---|---|
| **3D-j-Plasmid**<br>Template for 3D-j tiles | GTTCTAATACGACTCACTATA GGGCCTTAGCACAGA CTCTTGCAA GCCAGTCCG GTAA<br>CGGACTGGC ACTGTGCTGAGGCCC CGTACGAGGCCA TGATGTGAC GCCTGCAGCTCGGTG AAA<br>GATACTGTG CACCGAGTTGCAGGC AGGCCTTGTACG TT AT CACGAAGTCAATAC<br>GCAGCAGGTGCCGCA GGTACAAAG GCAGCAATGGGTCGG AAA TCCTAACTG<br>CCGACCCGTTGCTGC AGCGGCACTTGCTGC GCTGAGTCCGGA CGTACAATC<br>TCGACTCGGGACCGG AAA TGAGTTGAC CCGGTCCTGAGTCGA ACCGGATTCAGC TT AA<br>CACGAAGTCAATAC GCAGTTGCATAGGCA CGTTAGATC GGAGTAAGCAGGGTG AAA<br>GACTAAGTG CACCCTGTTTACTCC AGCCTATGTAACTGC CCAGACTCTGAG AATAATA<br>CCAGAGGTGGACTGG AAA CTCTAGATG CCAGTCCGCCTCTGG CTCAGGGTCTGG TT AC<br>CACGAAGTCAATAC CCAGCGTCTTACCGA CTAGAGATC GCAGCGATGAGCCGTG AAA<br>GTTCAGGAA CACGGCTCGTCGCTGC ACGGTAAGGCGCTGG GGCTGGATCTCA TACTATCGC<br>GGAGCCGTAATGTCAG AAA TCTTACACG CTGACATTGCGGCTCC AGAGATTCAGCC TT TA<br>CACGAAGTCAATAC GCACCGGTTGCGGCA GCAGTTCTA GGATGTAGTCGGACGG AAA<br>TAGCTCTAG CCGTCCGATTACATCC AGCCGCAATCGGTGC CCTACGATCGCA GACTCATTG<br>ACCTCCGTCACATGGC AAA GATTCTCAG GCCATGTGGCGGAGGT AGCGATTGTAGG TT TAAT<br>CACGAAGTCAATAC CCAGCAGGAACGTCG AAATAAA GCTGGCTGCCAGGTGG AAA<br>CTATGCAAG CCACCTGGTAGCCAGC CGACGTTTCTGCTGG GCTTACTAGTGG AATAATA<br>CCTGCCGTACCGCGTG AAA TTCCATGTC CACGCGGTGCGGCAGG CCACTGGTAAGC TT TACT<br>CACGAAGTCAATAC GCCTCCGGTTCCCGA GTATGCAGA GGCCTTGGC GTAA GCCAAGGCC<br>ACGGGAACTGGAGGC GCTTATGGCGCA GTATCCATG GCAGGCCTTAGCGGC AAA GGATTCACA<br>GCCGCTAGGGCCTGC AGCGCCGTAAGC TT ATAT CACGAAGTCAATAC GTGCGTGCACGACCA<br>TAGTTCGTC GGACCATGGGTCGTG AAA TTCTGATCC CACGACCTATGGTCC<br>AGGTCGTGTACGCAC CGTACGAGTGCA TAGCCAATG GTCGATCGCCCTCGG AAA CAGTAGAGA<br>CCGAGGGTGATCGAC AGCACTTGTACG TT AATT CACGAAGTCAATAC GCCTACTAGCGCGAA<br>GACCAATCA GGAGCCAGTACGCAG AAA TACGGAAAC CTGCGTATTGGCTCC<br>ATCGCGCTGGTAGGC CCTAGTCTAGCA TCACTCTTG CCAGGCGGACCGGTG AAA GGTATCGTA<br>CACCGGTTCGCCTGG AGCTAGGCTAGG TT ACTT CACGAAGTCAATAC GTCCAACC TT<br>CATGCTTACGACG | 1586 |
| **3D-k-Plasmid**<br>Template for 3D-k tiles | GTTCTAATACGACTCACTATA GGGCATGTGAGCGCA CGTTCTAAC GCAGCGGTATCGAGCG AAA<br>GAGGTAACA CGCTCGATGCCGCTGC AGCGCTCATATGCTC GCACCTTGCGCA CGCTATGAA<br>GCCCTGCTTGATACCG AAA GCTCAACTA CGGTATCAGGCAGGGC AGCGCAGGGTGC TT AT<br>CACGAAGTCAATAC CCAGGGTTAGATGCA TCCATTGAG CGGCTGAGCTTGCGGC AAA<br>CTTTCGGAA GCCGCAAGTTCAGCCG AGCATCTAGCCCTGG CGCCGGCATGGA CTTAGCTTC<br>AGGGCCAGCGCTCGCG AAA TAGGTAGTC CGCGAGCGTTGGCCCT ACCATGTCGGCG TT AA<br>CACGAAGTCAATAC CCAGAGGCTACTGGA TGTAGACAG CCATGCGGAGCCTCGG AAA<br>TCTGCATAC CCGAGGCTTCGCATGG ACCAGTAGTCTCTGG CCTCCGACGGCA CCATCATTG<br>CCCGTGATTACACGCG AAA GAATCGTCA CGCGTGTAGTCACGGG AGCCGTTGGAGG TT AC<br>CACGAAGTCAATAC GCCAATGCTGTCCCA GAGTGAACA GGATCCCGCTACGGTG AAA<br>GACGAACTA CACCGTAGTGGGATCC AGGGACAGTATTGGC GCTCAGGACCCA GATACGAGA<br>CAGTCTGGCTTAGGCC AAA GTTAGTCGA GGCCTAAGTCAGACTG AGGGTCTTGAGC TT TA<br>CACGAAGTCAATAC GAGTGCGGTAGCCCA GTCTGAATC TAATGAGTAACGCGTG AAA<br>TGATTGGTC CACGCGTTGCTCATTA AGGGCTACTGCACTC GCGGTGCTCGGA TGATAGCTG<br>CCGGAGTGGCGAGGTG AAA GATTGACTG CACCTCGCTACTCCGG ACCGAGTACCGC TT TAAT<br>CACGAAGTCAATAC GCCTCAGG TTCG CCTGAGGC CGTCGCGCA GGATCAGAA<br>GTGGCGTTTCAGGGTG AAA TAGGCATAG CACCCTGAGACGCCAC AGCGCGACG TT TACT<br>CACGAAGTCAATAC GCACCTGG TTCG CCAGGTGC CGGTCCAGCA GTTTCCGTA<br>GCAGGGATTCAGTCCG AAA GACTTAGGA CGGACTGAGTCCCTGC AGCTGGACCG TT ATAT<br>CACGAAGTCAATAC GCTCAGGG TTCG CCCTGAGC GGTCCGGCA GAGAGTTAC<br>CGTGAGCTCTATCCGG AAA TCGTCATAC CCGGATAGGGCTCACG AGCCGGACC TT AATT<br>CACGAAGTCAATAC CGAGTCGG TTCG CCGACTCG GCCTCGGGA CTGGATAGA<br>CAGTGGAGCGTAGGTG AAA TCTGCTTAG CACCTACGTTCCACTG ACCCGAGGC TT ACTT<br>CACGAAGTCAATAC GGCTCAGG TTCG CCTGAGCC GGTCGCGCA TCCTAGTAC<br>GTGCCGTGCCGACCTG AAA GTTAGCTAC CAGGTCGGTACGGCAC AGCGCGACC TT TATA<br>CACGAAGTCAATAC GTCCAACC TT CATGCTTACGACG | 1493 |

| Name (Note) | Sequence | length |
|---|---|---|
| **3D-l-Plasmid**<br>Template for 3D-l tiles | **GTTCTAATACGACTCACTATA** GGGCAAGTGGTCGCA **TAGGACTAG** GCGCTATAG **GTAA**<br>CTATAGCGC AGCGACCATTTGCCC GCTCGTCTGCCA **CCTTGATTC** GGCAGCTGT **TTCG**<br>ACAGCTGCC AGGCAGGCGAGC **TT AT CACGAAGTCAATAC** GCCTCCGCTGGTCCA<br>**GCTACTATC** GGAGCGAGATAGGTCG AAA **TCGAATCTC** CGACCTATTTCGCTCC<br>AGGACCAGTGGAGGC GCTGCTGTCGGA **TAGTAACGC** CACCGGTTTAGGCACG AAA<br>**TCTAGCAAC** CGTGCCTAGACCGGTG ACCGACGGCAGC **TT AA CACGAAGTCAATAC**<br>CCTACAGTCTGACCA **GATTCATCC** ACGGAGTGCGTCGGTG AAA **GATGATCAG**<br>CACCGACGTACTCCGT AGGTCAGATTGTAGG GCTACTGCACCA **GTTCCAATG**<br>TGACCGTGCGGTTCGG AAA **CAGAGATGA** CCGAACCGTACGGTCA AGGTGCGGTAGC **TT AC**<br>**CACGAAGTCAATAC** GCACCGTTGGATGCA **CCAGTTGTA** GGCACCGGTCCCGCAG AAA<br>**CAAGTCCAA** CTGCGGGATCGGTGCC AGCATCCAGCGGTGC CCCATGCGTGCA **GGATCTTGA**<br>CCGCCTATTGCCGTGG AAA **CAGTCTTAG** CCACGGCAGTAGGCGG AGCACGTATGGG **TT TA**<br>**CACGAAGTCAATAC** CAGCATTGCCTGGCA **CATACCTTG** GCAACAGGCAGGTCGC AAA<br>**GTTTCACAG** GCGACCTGTCTGTTGC AGCCAGGCGATGCTG GCTAGTACCCTA **TGAATGTCC**<br>GCTGCGTTATGGACTG AAA **GATAGATGC** CAGTCCATGACGCAGT AAGGGTGCTAGC **TT TAAT**<br>**CACGAAGTCAATAC** GCAACCTTTGACCTG **AAATAAA** GCATAGTGCGGACGTG AAA<br>**CTTACTTGC** CACGTCCGTACTATGC CAGGTCAGAGGTTGC CCAGGTGACGCA **GTCTGTAAG**<br>CCACGTCTCGTGACCG AAA **GTAGAGTCA** CGGTCACGGGACGTGG AGCGTCGCCTGG **TT TACT**<br>**CACGAAGTCAATAC** GGCATAGTCCGCTCA **GATAGGCAA** GCGCTCTCG **GTAA** CGAGAGCGC<br>AGAGCGGATTATGCC GCTCCTGTGGCA **CTAGTTACG** CCGTTGAAT **TTCG** ATTCAACGG<br>AGCCACGGGAGC **TT ATAT CACGAAGTCAATAC** CACCTCGTAGTTCCA **CTTTCATGC**<br>ATCTACAGGGACAGCG AAA **CATCACAAG** CGCTGTCCTTGTAGAT AGGAACTATGAGGTG<br>GCTACTTACGGA **CTCAGAATG** CCCTGGTTACGTCGGC AAA **GCATTCTCA**<br>GCCGACGTGACCAGGG ACCGTAGGTAGC **TT AATT CACGAAGTCAATAC** GCACCTTATCGAGGA<br>**CTCAAGTTG** GCCCTCGGGTCCCGTG AAA **GTGACAGAA** CACGGGACTCGAGGGC<br>ACCTCGATGAGGTGC GCACGTCCTGCA **TCGAGATAG** GCGGACGGCATGCGTG AAA<br>**GCTTGATGA** CACGCATGTCGTCCGC AGCAGGGCGTGC **TT ACTT CACGAAGTCAATAC**<br>**GTCCAACC TT CATGCTTACGACG** | 1566 |
| **3D-m-Plasmid**<br>Template for 3D-m tiles | **GTTCTAATACGACTCACTATA** GGGCTCGGAGCGGTA **CACTTCAGA**<br>CCGTAGTGTACGCATGCTCCGCAGGACGTTCTGCAGACCG AAA **GTTGATGTC**<br>CGGTCTGCGGAACGTCTTGCGGAGTATGCGTATACTACGG AGCCGCTCTGAGTCC<br>GCACCTGTTCCA **TGTCGATTG** GCCTGCATCGTCGTCG AAA **TCAACTGTC**<br>CGACGACGGTGCAGGT AGGAACGGGTGC **TT AT CACGAAGTCAATAC** GCCATCGGATGCGGA<br>**CACAGTATC** AGAGCCCAC **GTAA** GTGGGCTCT ACCGCATCTGATGGC CCAGGTGCGGCA<br>**GACATCAAC** TTAGCAGCG **TTCG** CGCTGCTAA AGCCGCGCCTGG **TT AA CACGAAGTCAATAC**<br>GCCTCGGAGCGTGCA **GTCAACTCA** GCGCAATGGCCTAATCTCATCTATGCTCCCGATCGGAACTG<br>AAA **TTGAAGACC** CAGTTCCGGTCGGGAGTATAGATGGGATTAGGTCATTGCGT<br>AGCACGCTTCGAGGC CCAGGGTCCGCA **CACTGTTCA** CCTGCGGGAGCCAGTG AAA<br>**CATCTATGC** CACTGGCTTCCGCAGG AGCGGATCCTGG **TT AC CACGAAGTCAATAC**<br>GCGTCTTGACCGGGA **CATCTAGAG** CCGCAACTGTTGCTTGGTACGGTATGCGATGCGACTATGTG<br>AAA **CTGATACTC** CACATAGTTGCATCGCGTACCGTATCAAGCAATAGTTGCGG<br>ACCCGGTCGAGACGC CCTAGGGACCGA **CACATGATC** TCCGGCTTGTAAACGG AAA<br>**CGCTAAGTA** CCGTTTACGAGCCGGA ACGGTCTCTAGG **TT TA CACGAAGTCAATAC**<br>GCCACCTG **TTCG** CAGGTGGC CCTCGCGCA **GGTCTTCAA** GGTCGCATGAGATCCG AAA<br>**GTCACATCA** CGGATCTCGTGCGACC AGCGCGAGG **TT TAAT CACGAAGTCAATAC** CCAGCGTG<br>**TTCG** CACGCTGG CCCTCGCCA **GAGTATCAG** CCGGAGCTCGTACCAG AAA **GATTGTACG**<br>CTGGTACGGGCTCCGG AGGCGAGGG **TT TACT CACGAAGTCAATAC** GCTACAGAGTGGGCA<br>**CGTGTAAGA** GCAGCGAGAGTCGTGG AAA **TTGCAAGAG** CCACGACTTTCGCTGC<br>AGCCCACTTTGTAGC GCAGCGCCAGCA **CTACATCTG** CCGGACGGGAGTACGC AAA<br>**CTTGTTCCA** GCGTACTCTCGTCCGG AGCTGGTGCTGC **TT ATAT CACGAAGTCAATAC**<br>CAGATGTCTGGACCA **CTGAGAATC** GCTAGGATTATGGCTG AAA **CTTTGTACC**<br>CAGCCATAGTCCTAGC AGGTCCAGGCATCTG CCGCTGCACTCA **CGTATGTTG**<br>ACCCTCAGGGCTTGCG AAA **GTTTGACAC** CGCAAGCCTTGAGGGT AGAGTGTAGCGG **TT AATT**<br>**CACGAAGTCAATAC** CGCAGTGCACCTCGA **GACATGGAA** CTGGTAGGACGAGGTG AAA<br>**GATCTAACG** CACCTCGTTCTACCAG ACGAGGTGTACTGCG GCTCAGCGGCAA **GTCAAGATG**<br>TCCCAGGTGAATGGAC AAA **TCTTCATCG** GTCCATTCGCCTGGGA ATGCCGTTGAGC **TT ACTT**<br>**CACGAAGTCAATAC GTCCAACC TT CATGCTTACGACG** | 1622 |

| Name (Note) | Sequence | length |
|---|---|---|
| **3D-n-Plasmid**<br>Template for 3D-n tiles | **GTTCTAATACGACTCACTATA** GGGCATGATCGAGCA **TGTGAATCC** GGCAACGCC **GTAA**<br>GGCGTTGCC AGCTCGATTATGCCC CCAGGTCGAGCA **GTACCTTAG** CCGCTTACC **TTCG**<br>GGTAAGCGG AGCTCGGCCTGG **TT AT CACGAAGTCAATAC** CCAGACGGGCTCGCA<br>**TCTCTACTG** TGCTCCGGGCCTCGTG AAA **GATCTCTAG** CACGAGGCTCGGAGCA<br>AGCGAGCCTGTCTGG CCGTGTCGACCA **TCAGTGAAG** GCTCGATGATGTGCAG AAA<br>**GATGTCTGA** CTGCACATTATCGAGC AGGTCGGCACGG **TT AA CACGAAGTCAATAC**<br>CCTATTTGAACCGCA **TACGATACC** CATCAGGTGCACCGTG AAA **TAGAACTGC**<br>CACGGTGCGCCTGATG AGCGGTTCGAATAGG GCTGCGAACGGA **TAGACATGC**<br>CTCTGGCTATGCGGTC AAA **CTTTCGTCA** GACCGCATGGCCAGAG ACCGTTTGCAGC **TT AC**<br>**CACGAAGTCAATAC** GCCTCAGG **TTCG** CCTGAGGC GGTCCGCGA **TTCCTGAAC**<br>GCGCTCCGGTAGCCAG AAA **CATGGATAC** CTGGCTACTGGAGCGC ACGCGGACC **TT TA**<br>**CACGAAGTCAATAC** GCCCTCGG **TTCG** CCGAGGGC CGGCGGTGA **CTAGAGCTA**<br>CCGACGTGCTCTCGCG AAA **CATTGGCTA** CGCGAGAGTACGTCGG ACACCGCCG **TT TAAT**<br>**CACGAAGTCAATAC** GGTCCGGA **TTCG** TCCGGACC GCCTCCGGA **CTTGCATAG**<br>CGTGGGTTCAAGCCGG AAA **CAAGAGTGA** CCGGCTTGGACCCACG ACCGGAGGC **TT TACT**<br>**CACGAAGTCAATAC** GCCTCAGG **TTCG** CCTGAGGC CGTCGCCCA **CAGTTAGGA**<br>GGGCAGCGGATCCCTG AAA **GCGATAGTA** CAGGGATCTGCTGCCC AGGGCGACG **TT ATAT**<br>**CACGAAGTCAATAC** GCCTCAGG **TTCG** CCTGAGGC GGTCCCGCA **CACTTAGTC**<br>GCGACCATGCGCTGTG AAA **CAATGAGTC** CACAGCGCGTGGTCGT AGCGGGACC **TT AATT**<br>**CACGAAGTCAATAC** GTCCAACC TT CATGCTTACGACG | 1087 |
